# Carborane–NHS esters as versatile reagents for amine-selective small-molecule modification, bioconjugation, and peptide-polymer assembly

**DOI:** 10.1101/2025.10.24.684456

**Authors:** Anže Jenko, Urban Barbič, Aljaž Renko, Ching-Pei Hsu, Dane Jemc, Špela Makuc, Lana Jamnik, Vera Župunski, Andrei Loas, Bradley L. Pentelute, Martin Gazvoda

**Author notes:** A.J and U.B. contributed equally.

## Abstract

Carboranes - hydrophobic, boron-rich clusters with unique structural properties - offer broad potential in materials science and biomedicine. They can mimic carbocyclic frameworks for designing advanced materials such as polymers and MOFs, and when incorporated into biomolecules, reduce local polarity while enriching boron content: a feature attractive for boron neutron capture therapy (BNCT). Here, we report the development of organic carborane reagents based on *N*-hydroxysuccinimide (NHS) esters serving as versatile platform for selective conjugation to amine residues, enabling functionalization of small molecules, peptides, and antibodies. We further introduce bifunctional carborane reagents bearing two NHS ester sites, which mediate inter- and intramolecular coupling as well as peptide-driven polymerization. The resulting carborane–peptide polymers, composed of fragments with distinct physicochemical features, represent a new class of hybrid materials. To demonstrate biomedical utility, therapeutic antibodies (trastuzumab, cetuximab, and daratumumab) were conjugated with these reagents, attaching up to 13 carboranes per antibody. Linker length was found to be critical for maximizing loading efficiency. nLC–MS/MS mapping of trastuzumab conjugates identified 15 preferential lysine modification sites, located outside CDR regions. Importantly, trastuzumab–carborane conjugates retained native HER2-targeting activity in BT-474 cell assays. These structures, integrating therapeutic antibody efficacy with boron delivery capability, hold promise as multimodal agents for BNCT.

## Introduction

Carboranes are unique boron-rich clusters whose stability and electronic properties underpin applications ranging from carbocyclic mimics in medicinal chemistry and functional motifs in polymers and metal–organic frameworks, to electron reservoirs in catalysis, and boron carriers for neutron capture therapy (BNCT) (Fig. 1a).^1,2,3^ Despite their broad potential, introducing carboranes into target compounds remains challenging, particularly within structurally complex systems such as biomolecules. ^4,5^ For example, peptide and protein carborane conjugates promise precise boron delivery for BNCT, but overcoming synthetic barriers is key to unlocking their full potential. ^6,7,8,9^

**Figure 1.**
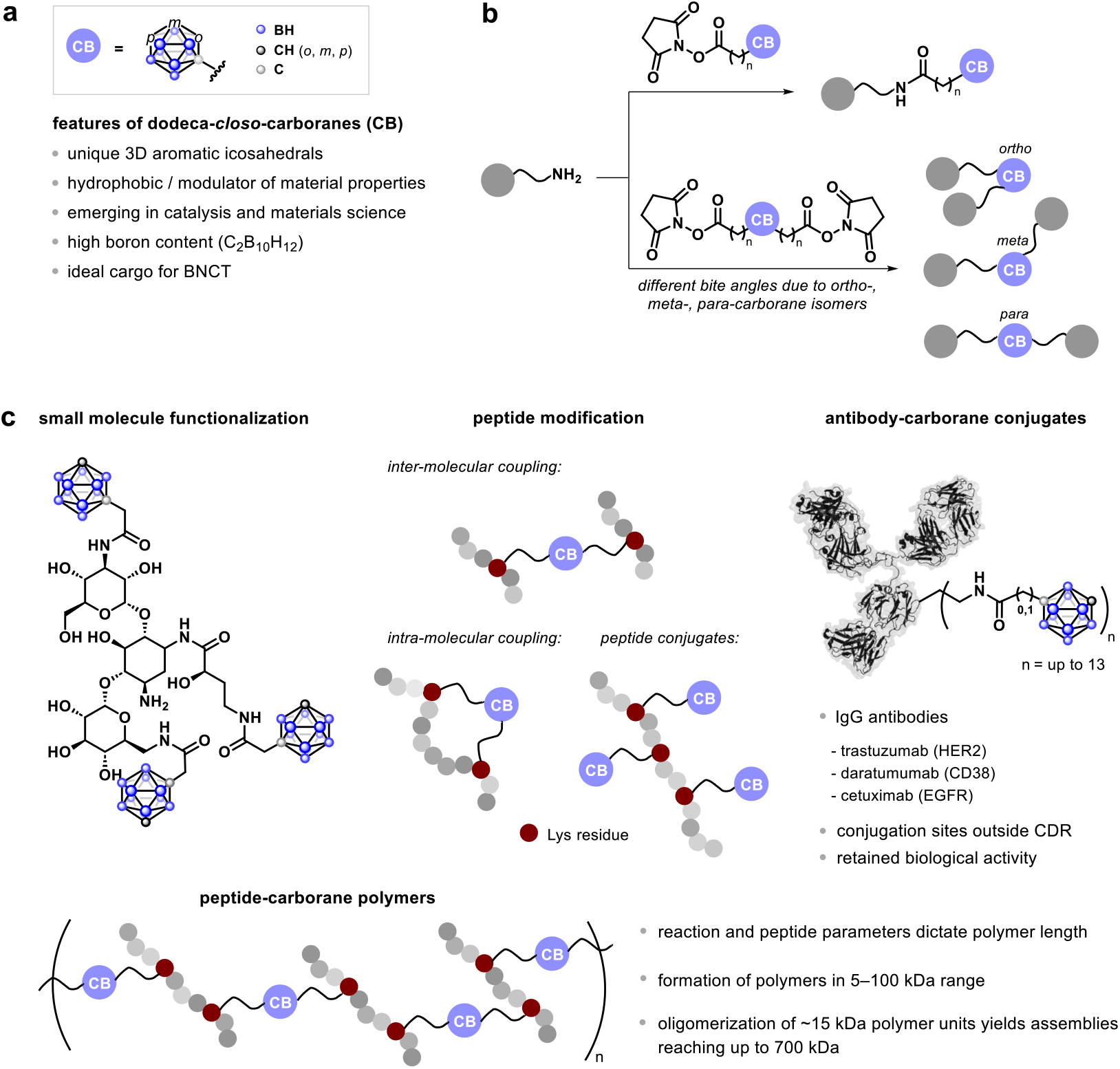
(a) Structure and general features of carboranes (dodeca-*closo*-carboranes). (b) Reactions of carborane–N-hydroxysuccinimide (NHS) esters enable functionalization of diverse structures through amine selective reactions. Additional NHS ester on carborane, forming bi-functional carborane–NHS esters, allow inter- and intra-molecular coupling, with substituent orientation depending on the carborane type (*ortho, meta, para*). (c) Developed reagents enable carborane introduction into small molecules, peptide functionalization (inter-/intra-molecular coupling), and antibody conjugation. Reagents allow installation of up to 13 carboranes per antibody, with conjugation occurring on lysine residues outside CDR regions as determined by nLC–MS/MS, while the antibodies retain their activity in cell-based assays. Under appropriate conditions, bifunctional carborane–NHS esters with peptides containing two Lys residues undergo polymerization, yielding novel type of polymers ranging from 10 kDa to 100 kDa, depending on the structure of monomers and reaction conditions.

Current strategies largely rely on solid-phase peptide synthesis using carborane-bearing noncanonical amino acids or late-stage acylation, both of which are hampered by side reactions, such as deboronation during Fmoc removal. ^10^ Alternative methods, including ribosomal incorporation of L-carboranylalanine, ^11^ cysteine borylation with PtII organometallic reagents, ^12^ or multistep conjugations to oxidized proteins, ^13,14^ have expanded the toolbox, however efficient late-stage modification of peptides and proteins in aqueous media remains scarce.

Beyond bioconjugation, carboranes have also been embedded into polymers,15 metal–organic frameworks (MOFs), ^16,^ and catalysts, ^17^ where they impart unique structural and electronic properties. These advances, however, often rely on specialized and case-specific syntheses. General and versatile methods for introducing carboranes into biomolecules and materials would therefore be highly valuable, both for advancing BNCT and for unlocking new opportunities in catalysis and materials design.

Following on recent palladium-mediated strategies for carborane installation into small molecules, peptides, and proteins based on the selective reactivity of thiols (cysteine) with palladium oxidative addition complexes,^18^ we sought to develop a complementary approach targeting amino groups. Amines are ubiquitous in small molecules and in the lysine side chains of peptides and proteins, making them attractive targets for functionalization. Among established reagents for amine-selective chemistry, *N*-hydroxysuccinimide (NHS) esters are especially versatile; ^19,20^ however, to our knowledge, carborane–NHS esters have not been reported yet.

Here we describe the development of carboranes functionalized with NHS esters (Fig. 1b) and demonstrate their broad application in modifying small molecules, peptides, and proteins (antibodies) (Fig. 1c). Notably, bis-NHS carboranes can act as interlinking fragments, enabling both inter- and intra-molecular peptide coupling, the latter of which can, for example, yield peptides with hydrophobic warheads. Moreover, under suitable conditions, these reagents drive the formation of carborane– peptide polymers, a completely new class of materials composed of structurally and chemically distinct fragments: hydrophobic carborane clusters and hydrophilic peptide domains (Fig. 1c).

## Results and Discussion

We have developed the synthesis of carborane–NHS ester reagents **1**–**7** (Fig. 2). Carborane with directly bound carboxylic group **7** was converted to the corresponding NHS ester **1** and isolated in 14% yield, owing to its lower stability (Fig. 2a). Following modified literature procedures, homologous **10** carboxylic acids of *meta*- and *para*-carboranes were accessed, and their subsequent reactions afforded *meta*- and *para*-carborane–NHS reagents **2** and **3**, in 50% and 87% yield, respectively (Fig. 2b). In contrast to **1**, the described structures **2** and **3** have a methylene linker between the carbonyl and NHS group, providing greater steric flexibility, which proved to be important for bioconjugation reactions (*vide infra*).

**Figure 2.**
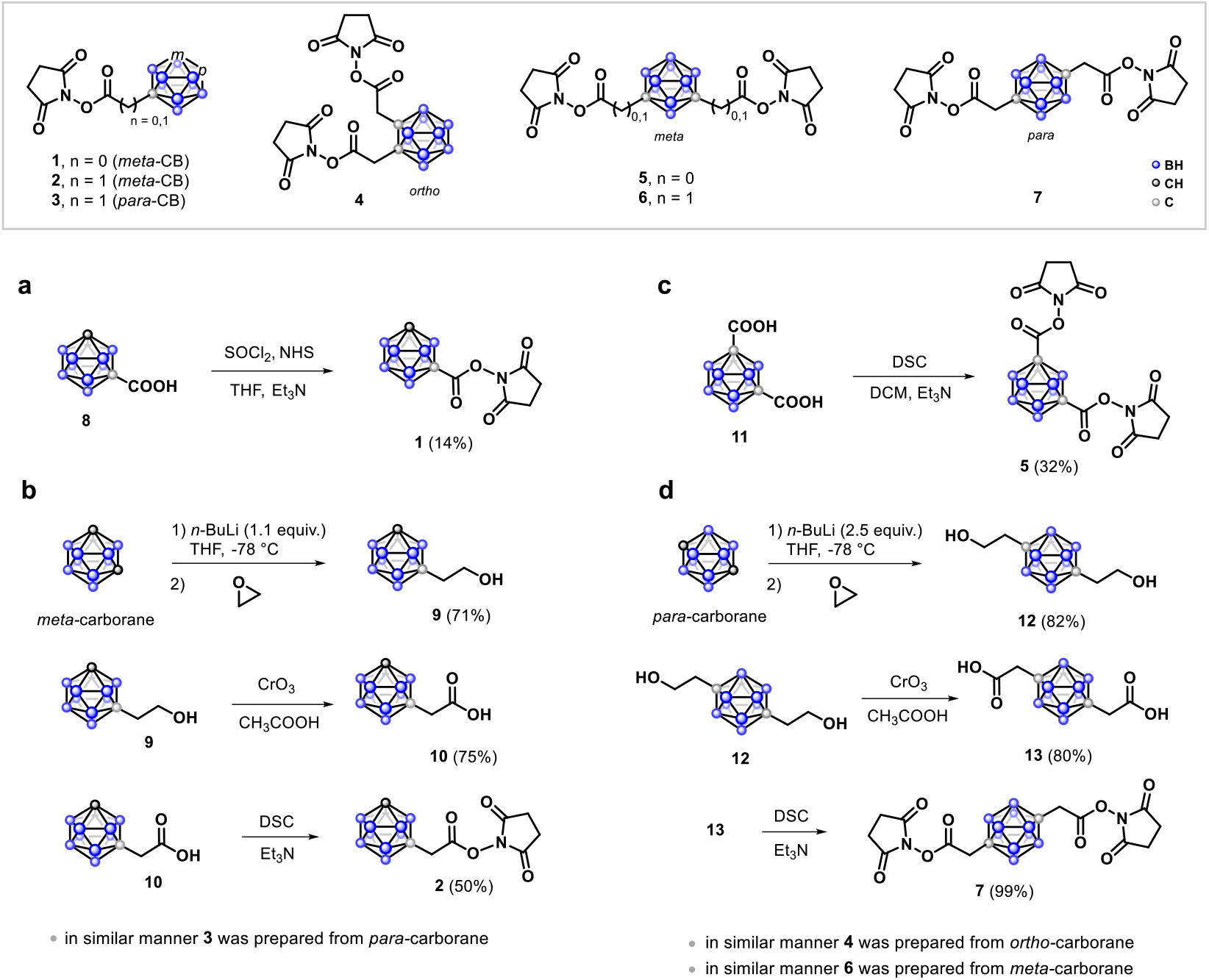
Developed carborane–NHS esters. Synthetic pathways for the preparation of (a, b) mono-NHS-functionalized carboranes **1**–**3** and (c, d) carboranes functionalized with two NHS ester groups **4**–**7** are shown. Isolated yields are indicated in parentheses. For more details on synthesis of **3, 4** and **6**, see the Supporting information.

To further increase the functionalization potential of these structures, we have prepared reagents with two NHS esters per carborane molecule **4**–**7**, which are suitable for both inter- and intra-molecular coupling (Fig. 2). In monofunctional carborane–NHS esters **1**–**3** the isomer mainly affects cage stability, whereas in bifunctional systems it also dictates the orientation of reactive groups. Given the 3D nature of car boranes (overall diameter of the icosahedral cage ∼5–6 Å; ∼1.5-times larger than benzene), controlling substituent position could be important. We therefore prepared *ortho*-, *meta*-, and *para*-carborane–bis-NHS esters **4**–**7**.

We synthesized NHS esters of *ortho*-, *meta*- and *para*-carborane **4, 6** and **7**, in which NHS ester is attached to the carborane via methylene linker. Additionally, bifunctional reagent of the *meta*-analog with two NHS esters directly attached to the carborane **5**. The latter was prepared from dicarboxylic acid **10** in 32 % isolated yield (Fig. 2c). The methylene-linked structures were synthesized analogously to **2**, but using excess *n*-BuLi and oxirane reagents to generate the corresponding alcohols, e.g., **11**, which were subsequently converted to bis-NHS esters, such as the *para*-carborane derivative **7** (Fig. 2d). In the same way, reagents **4** and **6** were obtained from *ortho*- and *meta*-carboranes, respectively. All developed reagents **1**–**7** were sufficiently stable for handling and short-term analysis under ambient conditions, however long-term storage was best achieved under an inert atmosphere at –20 °C.

Both type of developed reagents, having either one (Fig. 3a) or two NHS groups (Fig. 3b), proved suitable for amine-targeted reactions with small molecules and peptides. Reactions of amine-bearing small molecules with both types of reagents afforded carborane-functionalized products in nearly quantitative conversion, as determined by ^1^H NMR and HPLC–MS analyses. The corresponding small-molecule functionalized carboranes were subsequently isolated in 22–89% yields. Reactions of **1** with benzylamine, 3-phenylpropylamine and D-phenylalanine resulted in products **14, 15** and **17** in 59%, 38% and 89% isolated yields, respectively (Fig. 3c). In a similar way, the reaction of **2** with 3-phenylpropylamine afforded **16** in quantitative conversion as judged by 1H NMR and 39% isolated yield. Four equivalents of **1** reacted with amikacin, an antibiotic medication used for various bacterial infections^21^, to give near-quantitative conversion of the starting material into amikacin bearing two to four carboranes **18**–**20**, demonstrating potential of the developed reagents for late-stage functionalization and diversification.

**Figure 3.**
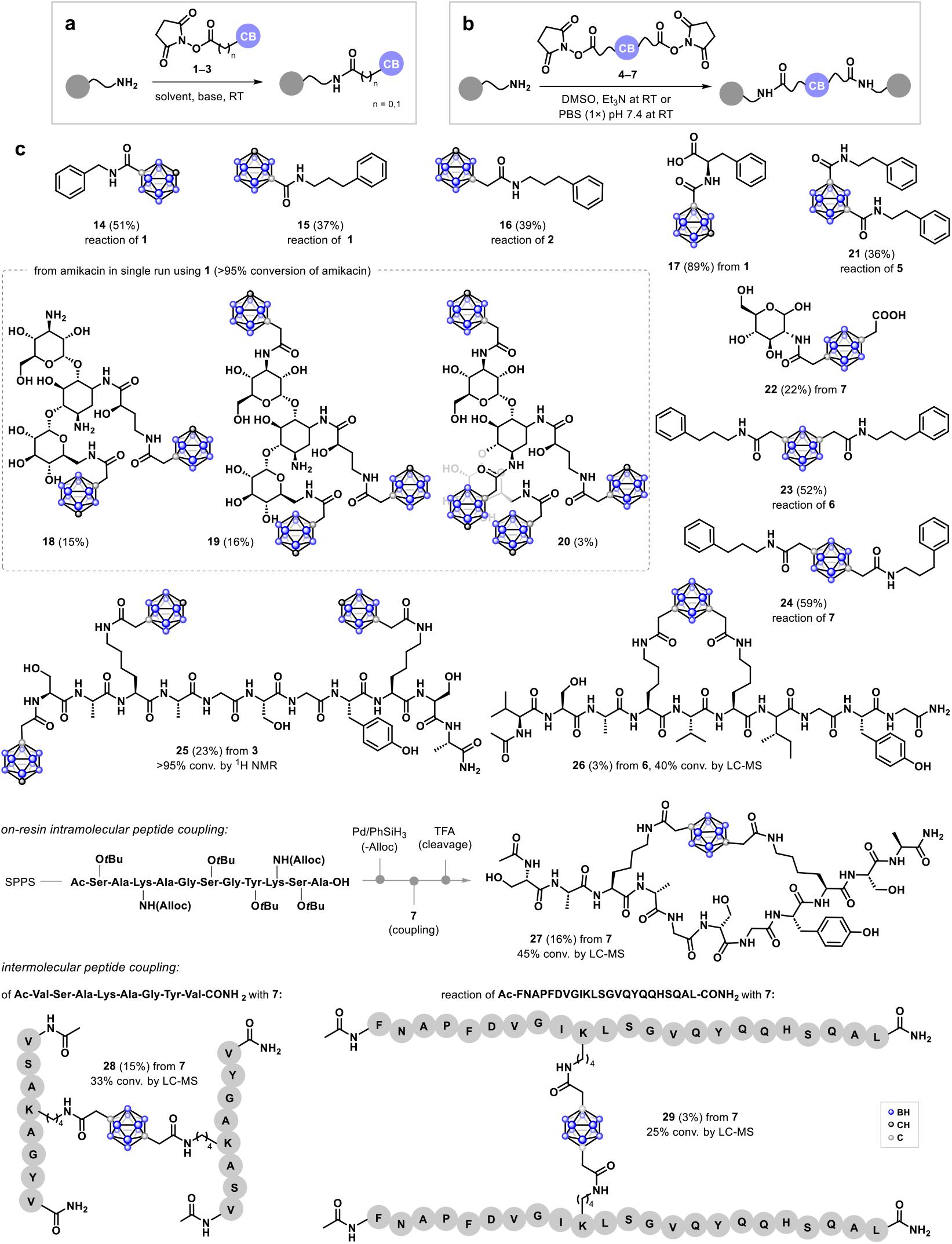
Reactions of amino residues of small molecules’ and peptides’ with carborane (a) mono- and (b) bi-functional NHS esters **1**–**7**. (c) Substrate scope resulting boron-rich small molecules and peptides, including inter-molecular and intra-molecular peptide coupling, the latter having potential to yield cyclic peptides. Isolated yields are indicated in parentheses.

Reactions of the bis-NHS carboranes **5, 6**, and **7** with 2-phenylethylamine or 3-phenylpropylamine afforded the bridged products **21, 23**, and **24** in 36%, 52%, and 59% isolated yields, respectively, with the carborane linking the two molecules. The reaction of **7** with glucosamine under basic conditions gave **22** in 22% isolated yield, with one NHS ester reacting with glucosamine and the other hydrolyzing in the basic medium.

Treatment of peptide H-Ser-Ala-Lys-Ala-Gly-Ser-Gly-Tyr-Lys-Ser-Ala-CONH_2_ (64 mM) bearing two Lys residues and a free N-terminal with 10 equiv. of **3** afforded conjugate **25** in quantitative conversion judged by ^1^H NMR and HPLC–MS, with 23% isolated yield. We investigated the intramolecular, linkage of two lysine residues within the N-capped Ac-Val-Ser-Ala-Lys-Val-Lys-Ile-Gly-Tyr-Gly-CONH_2_ peptide to generate a peptide with carborane hydrophobic warhead, thereby accessing intra-chain carborane-cyclized peptides. At a 14 mM concentration, the reaction gave about 40% conversion into **26** as determined by HPLC–MS, and the target product was isolated to confirm its structure.

To achieve more controlled assembly, we envisioned an on-resin coupling strategy^22^ using Alloc-protected Lys residues for selective deprotection and further modification. The resin-bound peptide Ac-Ser-Ala-Lys-Ala-Gly-Ser-Gly-Tyr-Lys-Ser-Ala-CONH–resin was prepared by SPPS using Alloc-protected Lys side chains, which were deprotected via the standard Pd/PhSiH_3_protocol. Subsequent overnight reaction with reagent **7**, followed by cleavage and global deprotection, afforded compound **27** in 45% conversion (HPLC–MS) and 16% isolated yield. To assess the effect of Lys spacing and NHS group positioning (*ortho, meta, para*) in the carborane reagent, we preliminarily examined the reaction of *ortho* analogue **4** with the Ac-VAGKGKGGIF-CONH–resin-bound peptide (Supporting information). Reagent **4**, containing adjacent NHS ester groups, yielded an intramolecularly cyclized product with a 2% conversion, as determined by HPLC–MS, suggesting that a greater spatial separation between the NHS esters is favorable for intramolecular coupling. However, further studies are required to correlate the spatial distance between amino (Lys) residues and the structure of the developed reagents to achieve efficient coupling.

Reactions of the N-capped 8-mer Ac-VSAKAGYV-CONH2 (144 mM) and N-capped obestatin, a ghrelin-derived peptide implicated in the modulation of feeding rhythm and hunger timing (45 mM), were carried out with reagent **7** in DMSO in the presence of Et_3_N. Both peptides contain a single lysine residue and afforded the intermolecular coupling products **28** and **29** in 33% and 25% conversion, respectively, as determined by HPLC–MS (Fig. 3c).

Carborane-based polymers, MOFs, and catalytic systems are emerging as a new class of functional materials, where the unique 3D geometry, hydrophobicity, and electronic richness of the cage impart properties unattainable with conventional organic or inorganic motifs. From electron-reservoir catalysts ultra-stable polyimides and ROMP-derived copolymers to water-stable, gas-selective MOFs, these architectures showcase the growing potential of carboranes to redefine the design landscape of advanced materials. ^23,24^

While exploring the intramolecular reaction to **26**, we observed formation of polymer-like byproducts. This suggested that, under appropriate conditions, larger polymers composed of two structurally distinct fragments - carborane and peptide - can be constructed (Fig. 4). To our knowledge, such hybrid architectures are unprecedented and represent a novel class of interesting materials.

**Figure 4.**
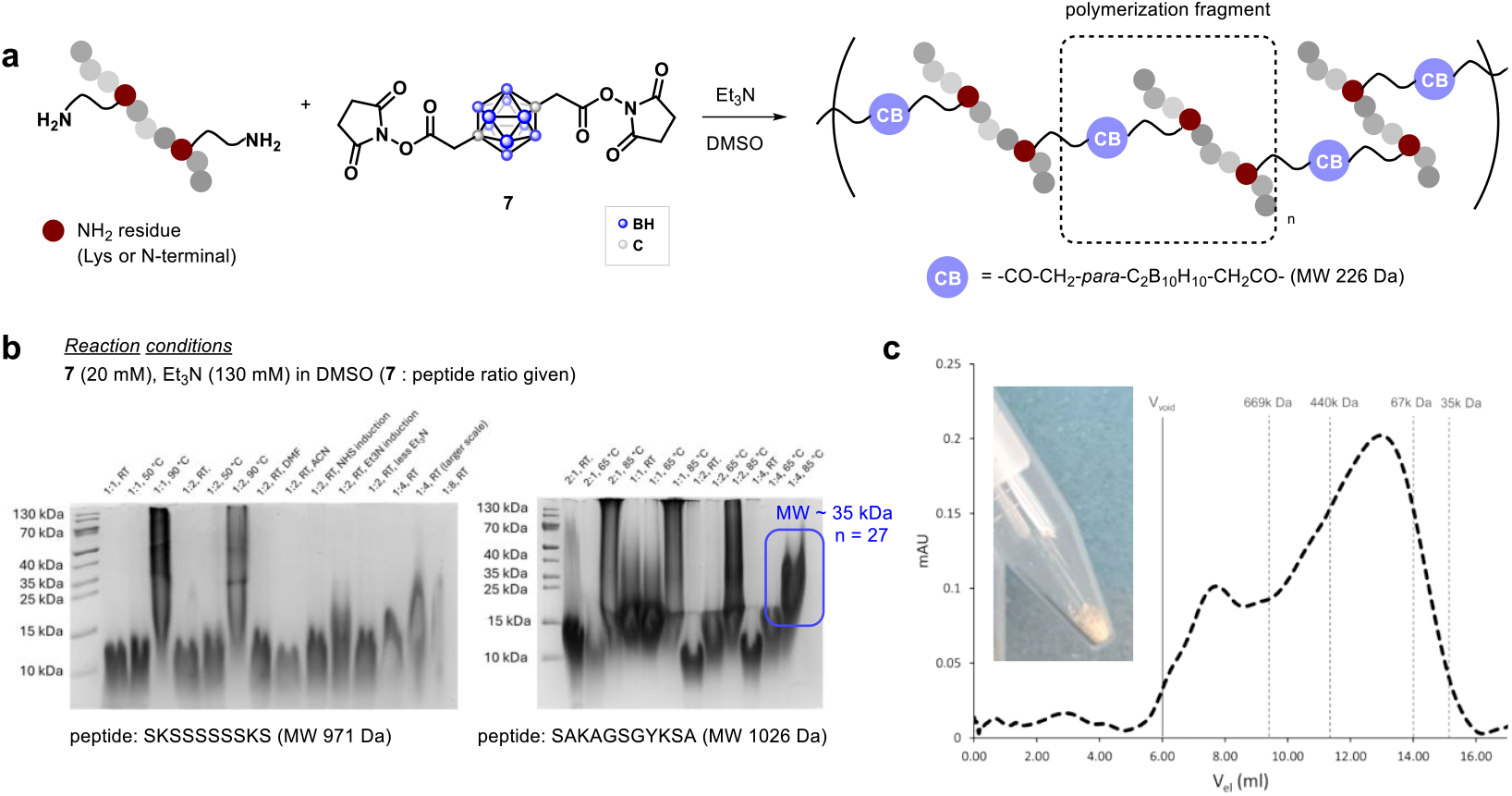
(a) Reaction scheme to construct peptide–carborane polymers. (b) Polymer length is tunable by reaction conditions and peptide structure. Screening reactions were performed on a small scale using 0.4 µmol of the limiting reagent (typically 20 µL reaction with 20 mM of **7** as limiting reagent). Prior to gel analysis, the compounds were incubated at 90 °C under reducing conditions to disrupt any potential non-covalent interactions. (c) Photo of an isolated carborane– peptide polymer, with a size-exclusion (SEC) chromatogram showing that the monomer polymer units undergo oligomerization in solution. The results indicate dynamic connectivity, with oligomers reaching up to ∼700 kDa (approximately 45 monomer units of about 15 kDa each). Most of the species, however, fall within the range from ∼70 kDa to ∼440 kDa to, corresponding to oligomers of 5 to 30 monomer units. Approximate elution positions of structures with different molecular weights are indicated, as determined by external standard analysis on a Superdex 200 10/300 GL column.

To investigate polymer formation, we employed two short model peptides, each containing two Lys residues, and examined how the ratio of bis-NHS carborane reagent **7** to peptide influenced the outcome (Fig. 4a). Optimization studies were carried out by systematically varying the reaction concentration (40–320 mM), reagent ratios, solvent (DMSO, DMF, or acetonitrile), temperature (22–90°C), and the order of reagent addition. Generally, reagent Et3N was added last to the mixture of peptide and **7**; however, we also investigated reactions in which **7** was introduced as the final reagent to initiate polymerization, to gain further insight into the polymerization process (Fig. 4b). Higher temperatures and larger excess of peptide relative to **7** favored the formation of longer polymers (Fig. 4b). The peptide sequence itself also influences the process; for example, while peptides of comparable length, such as SKSSSSSSKS and SAKAGSGYKSA, were both reactive, the latter reacted more efficiently, yielding polymers up to ∼35 kDa, corresponding to about 54 coupling events (n = 27). Under elevated temperature and near-equimolar peptide-to-**7** ratios, even longer polymers could be detected, although their formation was less controlled and produced heterogeneous mixtures (Fig. 4b).

The formed carborane–peptide polymers were stable under reducing conditions (90°C in the presence of DTT), to which they were subjected prior to SDS–PAGE analysis (Fig. 4b). Since each structurally distinct monomeric unit - i.e. carborane and peptide - of the constructed polymers is known to tend to oligomerize, ^25,26,27,28^ we investigated whether this behavior is retained in these hybrid structures. Size exclusion chromatography (SEC) revealed the formation of large oligomers of monomer units with molecular weights of up to approximately 700 kDa (about 45 polymer units of 15 kDa), with the majority of oligomers exhibiting molecular weights between 67 kDa and 440 kDa (oligomers of 5 to 30 polymer units).

We next evaluated the carborane–NHS reagents for antibody conjugation, transformation performed in almost completely aqueous media at moderate pH and temperature and low-µM antibody concentrations, to assess whether the reagents can transfer the carborane cage onto antibodies under these stringent conditions. IgG antibodies such as trastuzumab, cetuximab, and daratumumab contain approximately 85 lysine residues, of which about 40 are typically modifiable, including some located within CDR regions that are critical for activity. ^29^ Thus, in addition to overall conjugation efficiency, we were interested in determining whether the reagents targeted CDR or non-CDR lysines. Trastuzumab was chosen as a model antibody, while NHS ester **1**, featuring a rigid structure, and reagent **2**, containing a methylene linker between the carborane cage and NHS ester that imparts greater flexibility, served as model reagents. Reactions were carried out at 10 µM antibody concentration in PBS buffer (pH 7.4, 1×) at room temperature, with the NHS reagents dissolved in DMSO to address their low aqueous solubility, resulting in a final DMSO concentration of 5% to maintain protein stability (Fig. 5a)

**Figure 5.**
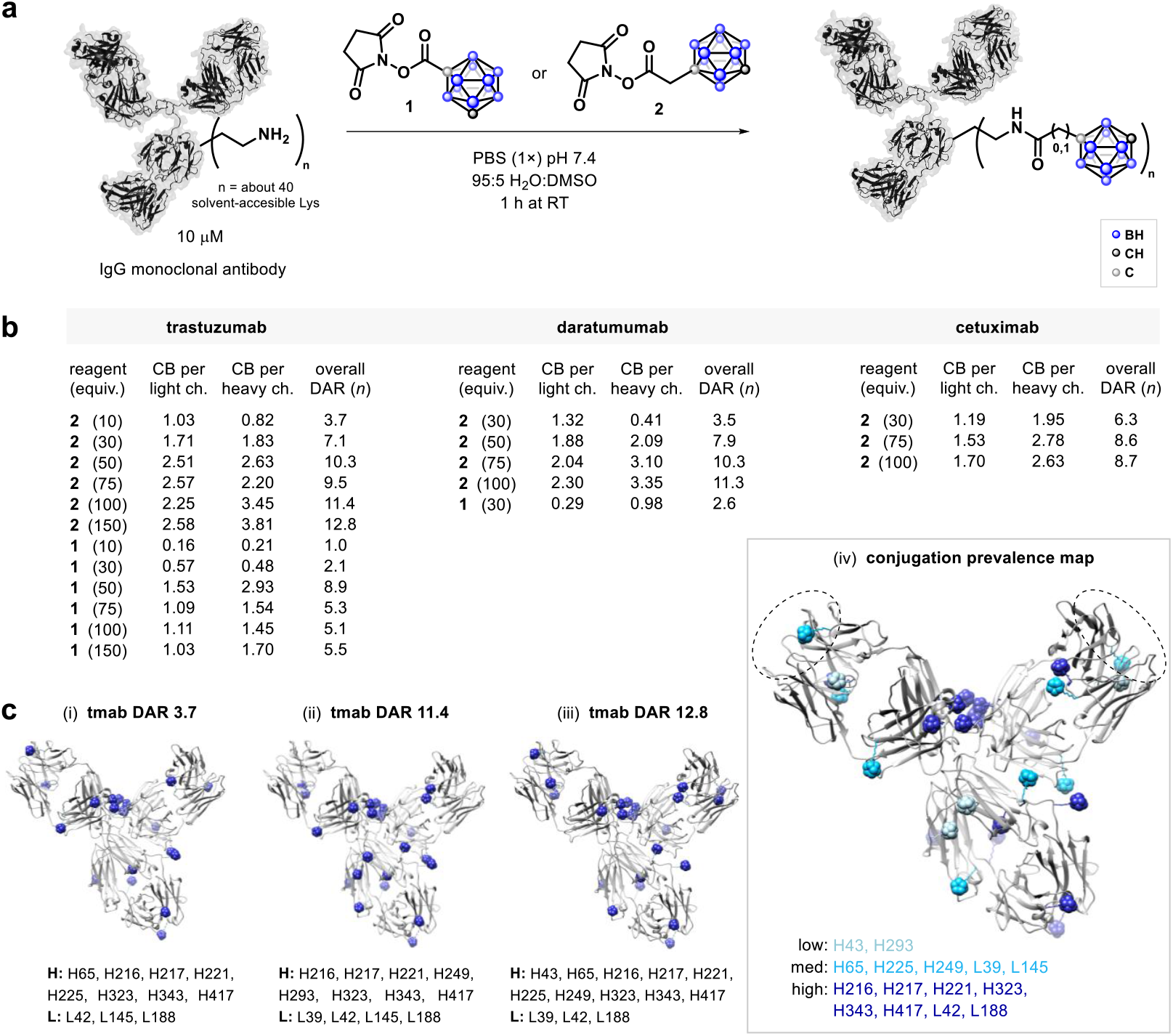
(a) Utilizing developed carborane–NHS reagents **1** and **2** for conjugation to antibodies. (b) Screening reaction conditions in respect to conjugation of light and heavy chains and overall DAR (drug-to-antibody ratio) according to LC– MS. (c) Mapping of modification sites as determined by nLC–MS/MS in trastuzumab with DAR (i) 3.7, (ii) 11.4, and (iii) 12.8, along with (iv) prevalence of Lys conjugation sites as based on statistical analysis of these results. Spherical blue shapes indicate carborane conjugation sites. The color scale in (iv), progressing from light to dark blue, reflects the relative prevalence of modification at each site. Prevalence was determined according to the number of reaction conditions (10, 100, and 150 equivalents of reagent) under which a given site was identified as modified. Sites consistently modified across all three conditions are depicted in the darkest blue, whereas those observed only under a single condition are represented in the lightest shade.; CDR regions are indicated with dashed circles (H: heavy chain, L: light chain).

Conjugation with reagent **2** enabled the installation of up to 13 carboranes per antibody (Fig. 5b). For example, 10, 30, and 50–100 equivalents of **2** yielded average DARs of 3, 7, and 9–11, respectively, while a maximum DAR (drug-to-antibody ratio) of 12.8 was achieved when using 150 equivalents.

Although yields were generally high (>50%), antibody recovery dropped by about 50% at the highest loading, and complete precipitation was observed when 200 equivalents of compound **2** were used, indicating limit of loading under these conditions. Reagent **1** proved less efficient, reaching a maximum DAR of ∼8 at 50 equivalents, while higher loadings led to precipitation and reduced overall conjugation efficiency. LC–MS analysis of reduced antibodies showed DAR values ranging from 1.0–2.6 for light chains and 0.8–3.8 for heavy chains (Fig. 5b). Similar trends were observed for daratumumab and cetuximab, demonstrating that despite its limitations at very high loadings, NHS ester **2** consistently delivers relatively high DARs across studied scaffolds, underscoring its potential for generating boron-rich antibodies.

To pinpoint the conjugation sites, we further analyzed trastuzumab conjugates at varying DAR levels,i.e. 3.7, 11.4, and 12.8, using nLC–MS/MS as a method for mapping (Fig. 5c, i–iii). In each case, we identified the conjugation positions within the heavy (H) and light (L) chains and then performed intersection statistical analysis to rank the conjugation sites from least (low) to most (high) probable (Fig. 5c). Fifteen lysine residues (H43, H293, H65, H225, H249, L39, L145, H216, H217, H221, H323, H343, H417, L42, L188) were identified as conjugation sites with varying prevalence, including two located within the CDR regions. Both showed only moderate likelihood of modification, suggesting that antigen binding could be largely preserved.

We further evaluated the prepared trastuzumab–carborane conjugates in cell-based assays to assess their functional activity. Notably, trastuzumab conjugates with DARs of 3.7 (37 boron atoms) and 9.5 (95 boron atoms) retained activity comparable to native trastuzumab in BT-474 cells at 50 nM (Fig. 6). These results indicate that, even at relatively high levels of conjugation, antigen recognition and downstream biological activity are largely preserved, highlighting the compatibility of the carborane modification with trastuzumab’s therapeutic function. Combing therapeutic profile of antibody with its simultaneous use as boron delivery agent for subsequent BNCT could establish such structures as multimodal agents for cancer treatment.

**Figure 6.**
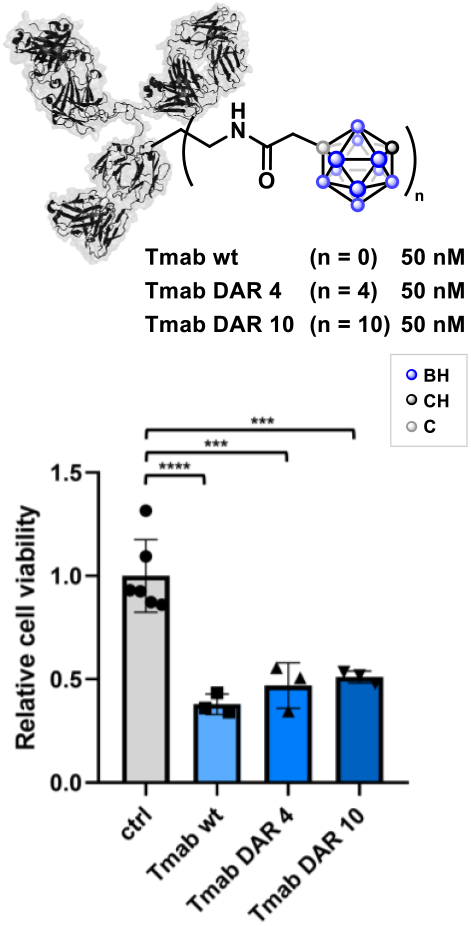
Carborane-conjugated trastuzumabs exhibited activity in BT-474 cells that was comparable to the wild-type form. Comparable inhibition of cell proliferation was observed for both Tmab conjugates with DAR of 4 and 10, corresponding to approximately 40 and 100 bound boron atoms, respectively. BT-474 cell lines were treated with the proteins at a concentration of 50 nM, and cell viability was assessed after 72 hours using relative luminescence from the CellTiter-Glo assay. All experiments were conducted in triplicate, with error bars representing variability. Statistical significance was evaluated (ns p > 0.05; *p < 0.05; **p < 0.01; ***p < 0.001; ****p < 0.0001) using one-way ANOVA, F(df, df) = X, p = Y, followed by Dunnett’s multiple comparisons test.

## Conclusion

In summary, we establish carborane–NHS esters as a versatile platform for amine-targeted functionalization enabling rapid and efficient preparation of small-molecule, peptide, and antibody– carborane conjugates. The bifunctional design of carborane–NHS esters allows both intra-molecular linkages and inter-molecular couplings, the latter being tunable toward polymer formation, as demonstrated in the preparation of peptide–carborane polymers - a novel class of materials. Importantly, the lysine conjugation strategy proved highly effective for generating antibody conjugates with exceptionally high boron loadings (DARs up to 12.8, corresponding to 128 boron atoms per antibody). Trastuzumab conjugates with DARs of 4 and 10 were evaluated *in vitro* cell-based assays and retained therapeutic activity comparable to the parent antibody, underscoring the compatibility of this modification with antibody function. Together, these findings highlight carborane–NHS esters as a powerful tool for constructing multimodal bioconjugates with potential utility as targeted boron delivery agents in anticancer therapy, combining established antibody mechanisms with boron delivery for BNCT. To our knowledge, described antibody conjugates represent the most boron-rich yet structurally defined and still biologically active biomolecules reported to date.

## Associated Content

Supporting Information containing additional experimental details and characterization data including 1H, 11B, and 13C NMR and mass spectra is available at xxxx

## Author Contributions

## Acknowledgments

UL FCCT authors acknowledge support from Slovenian Research Agency (project J1-60006, programme group P1-0230 and the Centre for Research Infrastructure at the University of Ljubljana, Faculty of Chemistry and Chemical Technology, which is part of the Network of Research and Infrastructural Centres UL (MRIC UL) and is financially supported by the Slovenian Research and Innovation Agency (Infrastructure programme No. I0-0022). M.G. acknowledges support from International Myeloma Foundation (Brian D. Novis Junior Grant). A.J. acknowledges Young Researcher Grant from Slovenian Research Agency. We thank Dr. Damijana Urankar from the Research Infrastructure Centre at the Faculty of Chemistry and Chemical Technology, University of Ljubljana for HRMS analyses of small molecules.

## References

1 Carboranes. Grimes, R. N. 3rd Edition, Academic Press, 2016.

2 Stockmann, P.; Gozzi, M.; Kuhnert, R.; Sárosi, M. B.; Hey-Hawkins, E. New keys for old locks: carborane-containing drugs as platforms for mechanism-based therapies. Chem. Soc. Rev. 2019, 48, 3497–3512.

3 Scholz, M.; Hey-Hawkins, E. Carbaboranes as Pharmacophores: Properties, Synthesis, and Application Strategies. Chem. Rev. 2011, 111, 7035–7062.

4 Grams, R. J.; Santos, W. L.; Scorei, I. R.; Abad-García, A.; Rosenblum, C. A.; Bita, A.; Cerecetto, H.; Viñas, C.; Soriano-Ursúa, M. A. The Rise of Boron-Containing Compounds: Advancements in Synthesis, Medicinal Chemistry, and Emerging Pharmacology. Chem. Rev. 2024, 124, 2441–2511.

5 Marfavi, A.; Kavianpour, P.; Rendina, L. M. Carboranes in drug discovery, chemical biology and molecular imaging. Nat. Rev. Chem. 2022, 6, 486–504.

6 Hu, K.; Yang, Z.; Zhang, L.; Xie, L.; Wang, L.; Xu, H.; Josephson, L.; Liang, S. H.; Zhang, M.-R. Boron agents for neutron capture therapy. Coord. Chem. Rev. 2020, 405, 213139.

7 Barth, R. F.; Mi, P.; Yang, W. Boron delivery agents for neutron capture therapy of cancer. Cancer Commun. 2018, 38, 35.

8 Dymova, M. A.; Taskaev, S. Y.; Richter, V. A.; Kuligina, E. V. Boron neutron capture therapy: Current status and future perspectives. Cancer Commun. 2020, 40, 406–421.

9 Luo, T.; Huang, W.; Chu, F.; Zhu, T.; Feng, B.; Huang, S.; Hou, J.; Zhu, L.; Zhu, S.; Zeng, W. The Dawn of a New Era: Tumor-Targeting Boron Agents for Neutron Capture Therapy. Mol. Pharm. 2023, 20, 4942–4970

10 Worm, D. J.; Hoppenz, P.; Els-Heindl, S.; Kellert, M.; Kuhnert, R.; Saretz, S.; Köbberling, J.; Riedl, B.; Hey-Hawkins, E.; Beck-Sickinger, A. G. Selective Neuropeptide Y Conjugates with Maximized Carborane Loading as Promising Boron Delivery Agents for Boron Neutron Capture Therapy. J. Med. Chem. 2020, 63, 2358–2371.

11 Yin, Y.; Ochi, N.; Craven, T. W.; Baker, D.; Takigawa, N.; Suga, H. De Novo Carborane-Containing Macrocyclic Peptides Targeting Human Epidermal Growth Factor Receptor. J. Am. Chem. Soc. 2019, 141, 19193–19197.

12 Waddington, M. A.; Zheng, X.; Stauber, J. M.; Hakim Moully, E.; Montgomery, H. R.; Saleh, L. M. A.; Král, P.; Spokoyny, A. M. An Organometallic Strategy for Cysteine Borylation. J. Am. Chem. Soc. 2021, 143, 8661–8668.

13 Wu, G.; Barth, R. F.; Yang, W.; Chatterjee, M.; Tjarks, W.; Ciesielski, M. J.; Fenstermaker, R. A. Site-Specific Conjugation of Boron-Containing Dendrimers to Anti-EGF Receptor Monoclonal Antibody Cetuximab (IMC-C225) and Its Evaluation as a Potential Delivery Agent for Neutron Capture Therapy. Bioconjug. Chem. 2004, 15, 185–194.

14 Yang, W.; Barth, R. F.; Wu, G.; Kawabata, S.; Sferra, T. J.; Bandyopadhyaya, A. K.; Tjarks, W.; Ferketich, A. K.; Moeschberger, M. L.; Binns, P. J.; Riley, K. J.; Coderre, J. A.; Ciesielski, M. J.; Fenstermaker, R. A.; Wikstrand, C. J. Molecular Targeting and Treatment of EGFRvIII-Positive Gliomas Using Boronated Monoclonal Antibody L8A4. Clin. Cancer Res. 2006, 12, 3792–3802.

15 Zhang, X.; Rendina, L. M.; Müllner, M. Carborane-Containing Polymers: Synthesis, Properties, and Applications. ACS Polym. Au 2024, 4, 7–33.

16 Meng, Y.; Lin, X.; Huang, J.; Zhang, L. Recent Advances in Carborane-Based Crystalline Porous Materials. Molecules 2024, 29, 3916.

17 Bawari, D.; Toami, D.; Jaiswal, K.; Dobrovetsky, R. Hydrogen splitting at a single phosphorus centre and its use for hydrogenation. Nat. Chem. 2024, 16, 1261–1266.

18 Gazvoda, M.; Dhanjee, H. H.; Rodriguez, J.; Brown, J. S.; Farquhar, C.; Truex, N. L.; Loas, A.; Buchwald, S. L.; Pentelute, B. L. Palladium-Mediated Incorporation of Carboranes into Small Molecules, Peptides, and Proteins. J. Am. Chem. Soc. 2022, 144, 7852–7860.

19 Spicer, C. D.; Pashuck, E. T.; Stevens, M. M. Achieving Controlled Biomolecule–Biomaterial Conjugation. Chem. Rev. 2018, 118, 7702–7743.

20 Haque, M.; Forte, N.; Baker, J. R. Site-selective lysine conjugation methods and applications towards antibody–drug conjugates. Chem. Commun. 2021, 57, 10689–10702.

21 Seely, S. M.; Parajuli, N. P.; De Tarafder, A.; Ge, X.; Sanyal, S.; Gagnon, M. G. Molecular basis of the pleiotropic effects by the antibiotic amikacin on the ribosome. Nat. Commun. 2023, 14, 4666.

22 Rossler, S.; Grob, N. M.; Buchwald, S. L.; Pentelute, B. L. Abiotic peptides as carriers of information for the encoding of small-molecule library synthesis. Science 2023, 379, 939–945.

23 Fisher, S. P.; Tomich, A. W.; Lovera, S. O.; Kleinsasser, J. F.; Guo, J.; Asay, M. J.; Nelson, H. M.; Lavallo, V. Nonclassical Applications of closo-Carborane Anions: From Main Group Chemistry and Catalysis to Energy Storage. Chem. Rev. 2019, 119, 8262–8290.

24 Xia, Q.; Zhang, J.; Chen, X.; Cheng, C.; Chu, D.; Tang, x.; Li, H.; Cui, Y. Synthesis, structure and property of boron-based metal–organic materials. Coord. Chem. Rev. 2021, 435, 213783.

25 Klingen, T. J.; Hepburn Jr., D.R. Investigation of γ-ray induced polymer formation in the carboranes—VI: Phase effects in the oligomerization of 1-allyl-o-carborane. J. Inorg. Nucl. Chem. 1975, 37, 1343–1346.

26 Cao, N.; Huang, K.; Xie, J.; Wang, H.; Shi, X. Self-assembly of peptides: The acceleration by molecular dynamics simulations and machine learning. Nano Today 2024, 55, 102160.

27 Hashimoto, K.; Panchenko, A. P. Mechanisms of protein oligomerization, the critical role of insertions and deletions in maintaining different oligomeric states. Proc. Natl. Acad. Sci. 2010, 107, 20352–20357.

28 Liu, C.; Luo, J. Protein Oligomer Engineering: A New Frontier for Studying Protein Structure, Function, and Toxicity. Angew. Chem. Int. Ed. 2023, 62, e202216480.

29 Walsh, S. J.; Bargh, J. D.; Dannheim, F. D.; Hanby, A. R.; Seki, H.; Counsell, A. J.; Ou, X.; Fowler, E.; Ashman, N.; Takada, Y.; Isidro-Llobet, A.; Parker, J. S.; Carroll, J. S.; Spring, D. R. Site-selective modification strategies in antibody–drug conjugates. Chem. Soc. Rev. 2021, 50, 1305–1353.

