## supporting information for "Carborane–NHS esters as versatile reagents for amine-selective small-molecule modification, bioconjugation, and peptide-polymer assembly"

### General information

#### General experimental information

All reactions were carried out in oven-dried glassware under a positive pressure of nitrogen using standard Schlenk techniques, unless otherwise stated. Reactions were typically performed in 5-, 10-, 25-, 50-mL, or 100-mL round-bottom flasks equipped with magnetic stir bars and sealed with rubber septa. Glassware was dried overnight in an oven at 130 °C prior to use. After drying, the flasks were sealed with rubber septa, pierced with a needle connected to the Schlenk line, and allowed to cool under a flow of nitrogen or argon. Reagents were introduced under nitrogen, followed by evacuation (15–30 min, depending on reagent sensitivity) and backfilling with inert gas. Reactions conducted at elevated temperatures were heated by submerging the reaction vessel in a preheated oil bath or an IKA heating/stirring block.

Thin-layer chromatography (TLC) was performed on aluminum-backed silica gel plates (Fluka Analytical or Merck) containing a fluorescent indicator ($\lambda$ = 254 nm). Visualization was achieved under UV light (CAMAG) or by staining with an aqueous potassium permanganate solution prepared from $KMnO_4$ (1.5 g), $K_2CO_3$ (10 g), and 10 % NaOH (1.25 mL) in 200 mL of water, followed by heating with a hot air gun.

Preparative column chromatography was carried out on silica gel (Fluka Silica Gel 60, 220–240 mesh). Automated flash chromatography was performed on an Interchim PuriFlash XS520Plus system using reverse-phase C18 columns (PF-15C18HP-F0025 or PF-15C18HP-F0004, Interchim) with MeCN/$H_2O$ gradients containing 0.1 % TFA. Fractions were collected based on UV detection at 220 nm, with thresholds of 10 or 5 mAU, using 25 mL test tubes.

#### General reagent information

Work-up solvents HPLC-grade solvents for chromatography were purchased from Honeywell, Carlo Erba and J. T. Baker. Deuterated chloroform ($CDCl_3$) and DMSO-$d_6$ were obtained from Euriso-top and were used without further purification. Dry solvents (acetonitrile (MeCN), dichloromethane (DCM), diethyl ether, tetrahydrofuran (THF), and toluene) were obtained via an MBraun MB SPS-5 solvent purification system, or were obtained from Sigma-Aldrich, e.g. hexane.

Reagents were obtained from commercial suppliers and used without further purification unless otherwise noted. Dicarba-*closo*-dodecaboranes were obtained from Sigma-Aldrich, Alfa Aesar, and Katchem. *N*-Hydroxysuccinimide (NHS), benzylamine, 3-phenylpropylamine, L-phenylalanine, and D(+)-glucosamine hydrochloride were obtained from Fluka. *N,N'*-Diisopropylcarbodiimide (DIC), trifluoroacetic acid (TFA), ethyl cyanohydroxyiminoacetate (Oxyma), and phenylsilane were purchased from Abcr. 1-Ethyl-3-(3-dimethylaminopropyl)carbodiimide (EDCI) was obtained from Carl ROTH. *n*-Butyllithium (*n*-BuLi), ethylene oxide, triethylamine ($Et_3N$), and *N,N'*-disuccinimidyl carbonate (DSC) were obtained from Sigma-Aldrich. Tetrakis(triphenylphosphine)palladium(0) was obtained from Fluorochem. Trastuzumab, cetuximab, and daratumumab were obtained from Obrnuta faza.

#### General analytical information

Melting points (Mp) were determined using a Kofler microscope fitted with a Leica Galen III hot stage; values are reported without correction.

Infrared (IR) spectra were obtained on a Bruker Alpha FT-IR spectrometer equipped with a Platinum ATR unit.

$^1$H, $^{13}$C{$^1$H}, and $^{11}$B{$^1$H} NMR spectra were recorded on a Bruker Avance III 500 MHz spectrometer (operating at 500 MHz, 126 MHz and 161 MHz) or Bruker Avance NEO 600 MHz NMR (operating at 600 MHz, 151 MHz and 193 MHz) at 298 K or 302. Chemical shifts (δ) are reported in ppm. Proton chemical shifts were referenced to the residual $CHCl_3$ signal in $CDCl_3$ (δ = 7.26 ppm) or DMSO-$d_5$ in DMSO-$d_6$ (δ = 2.50 ppm). Carbon signals were referenced to δ = 77.2 ppm ($CDCl_3$) and δ = 39.5 ppm (DMSO-$d_6$). Coupling constants (J) are given in Hz. Signal multiplicities are abbreviated as: s (singlet), d (doublet), t (triplet), dd (doublet of doublets), q (quartet), quint (quintet), m (multiplet), and br (broad).

High-resolution mass spectra (HRMS) were acquired on an Agilent 6224 TOF LC/MS instrument using electrospray ionization (ESI) under atmospheric pressure.

Analytical HPLC was carried out on an Agilent Infinity II 1260 LC system using a reverse-phase Atlantis C18 column (5 µm, 4.6 × 250 mm). Elution was performed with a gradient of acetonitrile (ACN) in water (both containing 0.1% trifluoroacetic acid, TFA): 20–80% ACN over 30 min or over 20 min, followed by 90% ACN for 3 min, at a flow rate of 0.75 mL/min. The column was re-equilibrated with 20% ACN for 5 min. Isocratic analyses were also conducted using ACN/$H_2O$ (50:50, 0.1% TFA) for 20 min. Compound purity was assessed by UV detection at 220 nm and expressed as the percentage of the compound peak area relative to the total integrated area.

HPLC–MS analyses were performed on an AZURA LC system (KNAUER) coupled to an Advion Expression CMS-L single quadrupole mass spectrometer (ESI, ~$10^{-3}$ mbar). Separation was achieved using a Eurospher II 100-5 C18 reverse-phase column (Vertex Plus, 125 × 4 mm ID) with a guard column. A gradient of 20–100% ACN in water (both with 0.1% formic acid) was applied over 10 min, followed by 2 min at 100% ACN. The column was then re-equilibrated at 20% ACN for 2 min. UV detection was carried out at 220 nm.

LC-MS Analysis. LC-MS chromatograms and corresponding mass spectra of antibodies were acquired using an Agilent 6550 ESI-Q-TOF mass spectrometer as described below. The mobile phases consisted of solvent A (0.1% formic acid in $H_2O$) and solvent B (0.1% formic acid in acetonitrile). Chromatographic separation was performed on a ZORBAX 300SB-C3 column (2.1 × 150 mm, 5 µm) maintained at 40 °C, using a gradient of 1–61% solvent B over 0–8 min at a flow rate of 0.5 mL min$^{-1}$. Mass spectrometric detection was carried out in positive electrospray ionization (ESI) mode with an extended dynamic range over m/z 100–3000. The drying gas temperature was set to 290 °C with a flow rate of 14 L min$^{-1}$, and the nebulizer pressure was maintained at 50 psig. The capillary, nozzle, fragmentor and octopole RF voltages were set to 5500 V, 2000 V, 380 V and 750 V, respectively. A reference calibration solution (mass 922.009798) was continuously introduced during analysis. Data were processed using Agilent MassHunter Workstation BioConfirm Software v10.0. Protein deconvolution was performed using a maximum entropy algorithm. Chromatograms shown were exported directly from BioConfirm Software.

nLC-MS/MS analysis of antibody samples from enzymatic digest were done on a Thermo Fisher Orbitrap Fusion Eclipse Tribrid Mass Spectrometer with an EASY-Spray source using a Thermo Fisher EASY-nLC 1200 System and Acclaim™ PepMap™ 100 C18 trap columns (20 mm × 75 µm, 3 µm particle size, 100 Å pore size, PN164946) and Acclaim™ PepMap™ RSLC C18 HPLC columns (150 mm × 50 µm, 2 µm particle size, 100 Å pore size, PN ES901). LC was performed with solvent A = $H_2O$ + 0.1% FA and solvent B = 80% MeCN + 0.1% FA + 19.9% $H_2O$ prepared with LiChrosolv® water and MeCN suitable for MS (Millipore Sigma) and Optima™ LC/MS grade formic acid (Thermo Fisher Scientific). Chromatography was performed at 40 °C with a flow rate of 300 nL/min using the following gradient: 1–51% B (0–90 min), 51–90% B (90–100 min), 90% B (100–110 min), with MS acquisition from 0–90 min in a data-dependent method. Full MS cycle time = 3 s. Detector type = Orbitrap.

#### Experimental procedures and characterization data

##### Preparation and characteriztaion of small molecules (Fig. 2 and Fig. 3)

1,7-Dicarba-*closo*-dodecaboran-1-carboxylic acid (**8**)

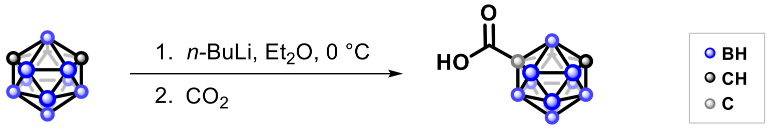

**8** (92%)

Compound **8** was prepared according to a modified literature procedure.[1,2] The reaction was carried out under an inert nitrogen atmosphere using standard techniques as decscribed in general information. A 100 mL oven-dried round-bottom flask equipped with a magnetic stir bar was connected to a Schlenk line with a hose, evacuated, and backfilled with nitrogen. While under a positive pressure of nitrogen, the tube was then uncapped and a compound *meta*-CB (288 mg, 2.0 mmol) was added followed by the addition of dry diethyl ether (20 mL), freshly dispensed from an MBraun MB SPS5 solvent purification system with the aid of a syringe. The reaction mixture was cooled to –78 °C in an ethanol bath (liquid nitrogen). While maintaining this temperature, *n*-BuLi (2.5 M in hexanes; 0.96 mL, 2.4 mmol) was added dropwise via syringe. The mixture was stirred at –78 °C for 20 min, after which finely ground dry ice ($CO_2$: 1.0 g) was added directly to the flask. The reaction mixture was maintained at –78 °C for an additional 30 min and then allowed to warm up to room temperature over 1 h. The solvent was removed under reduced pressure, and the residue was dissolved in 15 mL of water and transferred to a separatory funnel. The aqueous layer was washed with petroleum ether (2 × 15 mL). The combined aqueous layers were acidified to pH 4 using 2 M HCl (aq) and extracted with petroleum ether (3 × 30 mL). The organic layers were combined, dried over anhydrous $Na_2SO_4$, filtered, and concentrated under reduced pressure using a rotary evaporator. Crude product **8** (346 mg, 92%) was sufficiently pure and was used directly in the next synthetic step without further purification.

**IR** ($cm^{-1}$): 3065, 2869, 2611, 1711, 1415, 1273, 905, 717.

**$^1$H NMR** (500 MHz, $CDCl_3$): δ 3.05 (br, 1H), 3.4–1.6 (m, 10H).

NMR data were in agreement with the literature.[1]

1-(*N*-succinimidyl carbonate)-1,7-dicarba-*closo*-dodecaboran (**1**)

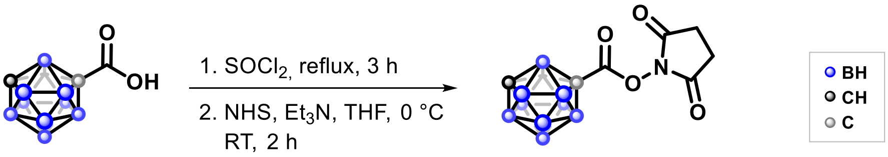

**8** **1** (14%)

Compound **1** was prepared in two steps, following modified literature procedures.[3,4] The reaction was carried out under argon atmosphere. To a 50 mL oven-dried round-bottom flask containing compound **8** (326 mg, 1.73 mmol) was added thionyl chloride (16 mL, 220 mmol). The reaction mixture was heated to reflux under argon for 3 h, then cooled to room temperature. Excess thionyl chloride was removed under reduced pressure using a rotary evaporator. The resulting residue was dissolved in dry tetrahydrofuran (9.6 mL) and cooled to 0 °C. *N*-hydroxysuccinimide (NHS, 406 mg, 3.53 mmol) and triethylamine ($Et_3N$, 0.486 mL, 3.53 mmol) were added, and the mixture was stirred at room temperature for 2 h. The solvent was evaporated under reduced pressure, and the residue was dissolved in dichloromethane (DCM, 10 mL). The organic layer was washed with water (3 × 10 mL) and saturated aqueous NaCl (1 × 10 mL), dried over anhydrous $Na_2SO_4$, filtered, and concentrated under reduced pressure. The crude product was purified by silicagel column chromatography (100% DCM) to yield compound **1** as a white solid (70 mg, 14%).

**R*f*** 0.3 (DCM)

**Mp** 66–68 °C

**IR** ($cm^{-1}$): 3066, 2611, 1694, 1423, 1211, 1077, 814, 651.

**$^1$H NMR** (500 MHz, $CDCl_3$): δ 3.40–1.70 (m, 10H, protons of carborane cage), 3.10 (br, 1H, $CH_{carboran}$), 2.83 (br, 4H, $CH_2$-$CH_2$).

**$^{13}$C NMR** (126 MHz, $CDCl_3$): δ 167.7 (C=O, CON), 158.3 (C=O, COO), 67.4 ($C_{carboran}$), 55.2 ($C_{carboran}$H), 25.5 ($CH_2$-$CH_2$).

**$^{11}$B NMR** (161 MHz, $CDCl_3$): δ -4.08, -5.13, -6.11, -9.76, -10.66, -12.38, -13.43, -14.83, -15.97.

**HRMS** (*m/z*): $[M+H]^+$ calculated for $C_7H_{16}B_{10}NO_4^+$, 286.2078; found, 286.2079.

1,2-Dicarba-*closo*-dodecaboran-1-ethanol (**SI-1**) and 1,2-dicarba-*closo*-dodecaboran-1,2-diethanol (**SI-2**):

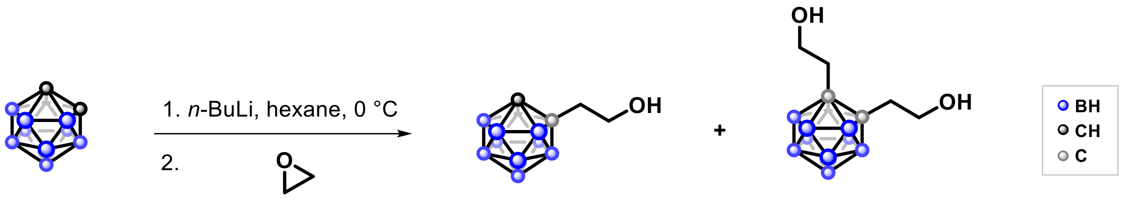

**SI-1** (61%) **SI-2** (27%)

Compounds **SI-1** and **SI-2**, similary as *meta*- and *para*-carborane analogues described below, were synthesized following a modified literature procedure.[5] All reactions were performed under an inert nitrogen atmosphere using standard Schlenk techniques. In a 25-mL oven-dried round-bottom flask equipped with a magnetic stir bar and connected to a Schlenk line, the system was evacuated and backfilled with nitrogen. Under positive nitrogen pressure, the flask was uncapped and *ortho*-carborane (864 mg, 6 mmol) was added, followed by dry hexane (3.5 mL). The reaction mixture was cooled to 0 °C, and *n*-BuLi (1.6 M in hexane, 5.63 mL, 9 mmol) was added dropwise. After the addition was complete, the reaction mixture was allowed to warm to room temperature and stirred for 30 min. The mixture was then cooled again to 0 °C, and a solution of ethylene oxide in THF (2.5–3.3 M, 3.6 mL, 9 mmol) was added dropwise. The reaction was stirred at room temperature for 5 h under a continuous nitrogen flow. Subsequently, methanol (1.5 mL) and acetic acid (2 drops) were added via syringe. Then, water (10 mL) was added, and the reaction mixture was transferred to a separatory funnel and extracted with diethyl ether (3 × 15 mL). The combined organic layers were dried over anhydrous $Na_2SO_4$, filtered, and concentrated under reduced pressure using a rotary evaporator. The crude residue, as confirmed by $^1$H NMR, contained a mixture of **SI-1** and **SI-2**. The mixture was purified by column chromatography (first with petroleum ether/ethyl acetate = 25:3, then with petroleum ether/ethyl acetate = 1:1), affording the individual isomers **SI-1** (687 mg, 61%) and **SI-2** (371 mg, 27%) as white solids.

**SI-1**

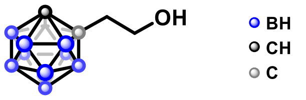

**$^{1}$H NMR** (500 MHz, $CDCl_3$): δ 3.99 (br, 1H), 3.83 (td, *J* = 4.6, 16.2 Hz, 2H), 2.9–1.6 (m, 10H), 2.49 (t, *J* = 5.9 Hz, 2H). NMR data were in agreement with the literature.[5]

**SI-2**

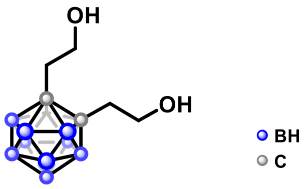

**$^{1}$H NMR** (500 MHz, $CDCl_3$): δ 3.87 (td, *J* = 5.1, 18.4 Hz, 4H), 2.8–1.7 (m, 10H), 2.54 (t, *J* = 6.7 Hz, 4H). NMR data were in agreement with the literature.[5]

**9** and **SI-3** were prepared from *meta*-carborane in the same manner as described for **SI-1** and **SI-2**: *m*-carborane: 576 mg, 4 mmol; hexane: 3 mL; *n*-BuLi: 6.25 mL, 10 mmol; ethylene oxide: 4 mL, 10 mmol.

1,7-Dicarba-*closo*-dodecaboran-1-ethanol (**9**)

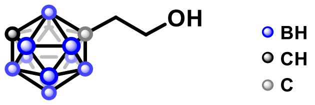

136 mg (18%)

**$^{1}$H NMR** (500 MHz, $CDCl_3$): δ 3.64 (t, *J* = 6.9 Hz, 2H), 2.94 (br, 1H), 2.23 (t, *J* = 6.9 Hz, 2H). NMR data were in agreement with those from the literature.[5]

1,7-Dicarba-*closo*-dodecaboran-1,7-diethanol (**SI-3**)

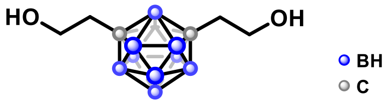

483 mg (52%)

**$^{1}$H NMR** (500 MHz, $CDCl_3$): δ 3.64 (t, *J* = 6.9 Hz, 4H), 2.22 (t, *J* = 6.9 Hz, 4H). NMR data were in agreement with the literature.[5]

**SI-4** and **12** were prepared from *para*-carborane in the same manner as described for synthesis of **9** and **SI-3**: *para*-carborane: 500 mg, 3.47 mmol; hexane: 2 mL; *n*-BuLi: 2.81 mL, 4.51 mmol; ethylene oxide: 1.81 mL, 4.51 mmol.

1,12-Dicarba-*closo*-dodecaboran-1-ethanol (**SI-4**)

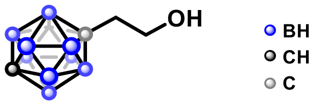

300 mg (46%)

**$^1$H NMR** (500 MHz, $CDCl_3$): δ 3.47 (t, $J$ = 6.9 Hz, 2H), 2.8–1.7 (m, 10H), 2.68 (br, 1H), 1.92 (t, $J$ = 6.9 Hz, 2H). NMR data were in agreement with the literature.[5]

1,12-Dicarba-*closo*-dodecaboran-1,12-diethanol (**12**)

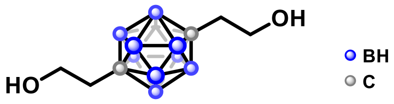

274 mg (34%)

**$^1$H NMR** (500 MHz, $CDCl_3$): δ 3.46 (td, $J$ = 5.8, 19.7 Hz, 4H), 2.9–1.7 (m, 10H), 1.92 (t, $J$ = 7.0 Hz, 4H). $^1$H NMR shows the presence of residual solvents. NMR data were in agreement with the literature.[5]

1,7-Dicarba-*closo*-dodecaboran-1-acetic acid (**10**)

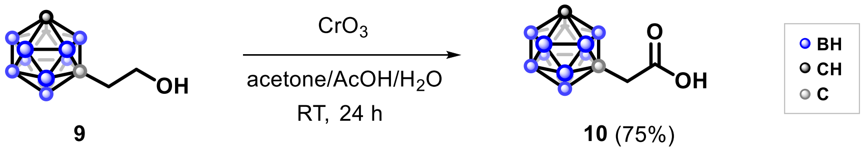

Compound **10** was synthesized following a modified literature procedure.[5] A 100 mL round-bottom flask containing compound **9** (188 mg, 1 mmol) was charged with acetone (7.5 mL) and cooled to 0 °C. A solution of chromium(VI) oxide ($CrO_3$, 700 mg, 7 mmol) in a mixture of acetic acid (25 mL) and water (25 mL) was added dropwise to the stirred solution. The reaction mixture was then allowed to warm to room temperature and stirred overnight. Upon completion, the mixture was extracted with diethyl ether (3 × 40 mL). The combined organic layers were acidified to pH 4 with 2 M HCl (aq) and washed with saturated aqueous NaCl (1 × 50 mL). The organic phase was dried over anhydrous $Na_2SO_4$, filtered, and concentrated under reduced pressure using a rotary evaporator. Compound **10** was obtained as a white solid (146 mg, 75%) and was used directly in the subsequent reaction without further purification.

**$^1$H NMR** (500 MHz, $CDCl_3$): δ 3.00 (1H, s), 2.96 (2H, s). NMR data were in agreement with the literature.[5]

1,12-Dicarba-*closo*-dodecaboran-1-acetic acid (**SI-5**)

Compound **SI-5** was prepared in the same manner as acid **10**: **SI-4**: 177 mg, 0.94 mmol; acetone: 7 mL; $CrO_3$: 678 mg, 6.8 mmol, water: 24 mL; acetic acid: 20 mL.

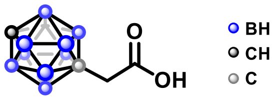

153 mg, 81%

**$^1$H NMR** (500 MHz, $CDCl_3$): δ 2.88–1.64 (m, 10H), 2.72 (br, 1H), 2.63 (s, 2H). NMR data were in agreement with the literature.[7]

1-(2,5-dioxopyrolidin-1-yl)acetate-1,7-dicarba-*closo*-dodecaborane (**2**)

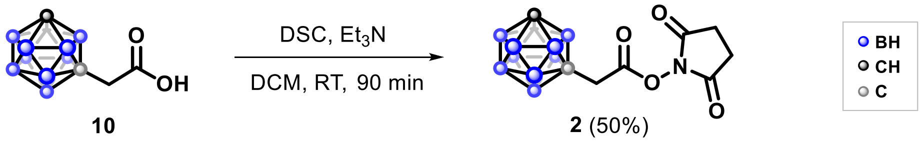

**10** **2** (50%)

Compound **2** was synthesized following a modified literature procedure.[8] The reaction was carried out under an inert nitrogen atmosphere using standard Schlenk techniques. To a 25 mL oven-dried round-bottom flask containing compound 15 (146 mg, 0.75 mmol) were added dry dichloromethane (3 mL), *N,N'*-disuccinimidyl carbonate (DSC; 330 mg, 1.29 mmol), and triethylamine ($Et_3N$; 0.105 mL, 0.75 mmol). The mixture was stirred at room temperature for 90 min. Subsequently, dichloromethane (10 mL) was added, and the reaction mixture was transferred to a separatory funnel. The organic phase was washed with 1 M HCl (aq) (2 × 10 mL), dried over anhydrous $Na_2SO_4$, filtered, and concentrated under reduced pressure using a rotary evaporator. The crude residue was purified by column chromatography, affording compound **2** as a white solid (147 mg, 50%).

**Mp** 121–125 °C

**IR** ($cm^{-1}$): 2960, 2596, 1810, 1777, 1740, 1202, 1117, 1061, 725, 644.

**$^1$H NMR (**500 MHz, $CDCl_3$): δ 3.19 (br, 2H, -$CH_2$-), 3.2–1.7 (protons of carborane cage), 3.01 (br, 1H, $CH_{carborane}$), 2.85 (br, 4H, $CH_2$-$CH_2$).

**$^{13}$C NMR** (126 MHz, $CDCl_3$): δ 168.5 (C=O, CON), 163.0 (C=O, COO), 66.9 ($C_{carborane}$), 55.6 ($C_{carborane}$H), 38.5 (-$CH_2$-), 25.6 ($CH_2$-$CH_2$).

**$^{11}$B NMR** (160 MHz, $CDCl_3$): δ -2.91, -3.91, -8.41, -9.77, -10.76, -12.59, -13.63, -14.53, - 15.66.

**HRMS** (*m*/*z*): $[M+H]^+$ calculated for $C_8H_{18}B_{10}NO_4^+$: 300.2234; found: 300.2236.

1-(2,5-dioxopyrolidin-1-yl)acetate-1,12-dicarba-*closo*-dodecaborane (**3**)

Carborane NHS ester **3** was prepared in the same manner as described for compound **2**: **SI-5**: 40 mg, 0.20 mmol; DSC: 55 mg, 0.21 mmol; $Et_3N$: 23 µL, 0.16 mmol; DCM: 2 mL.

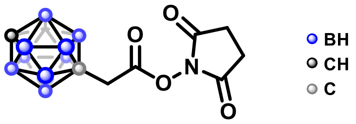

65 mg, 87%

**Mp** 111.4–114.8 °C

**IR** ($cm^{-1}$): 3058, 2931, 2607, 1823, 1732, 1366, 1202, 1066, 730, 644.

**$^1$H NMR** (500 MHz, $CDCl_3$): δ 2.89 (s, 2H, -$CH_2$-), 2.9–1.6 (protons of carborane cage), 2.85 (br, 4H, $CH_2$-$CH_2$), 2.74 (br, 1H, $CH_{carborane}$).

**$^{13}$C NMR** (126 MHz, $CDCl_3$): δ 168.4 (C=O, CON), 162.7 (C=O, COO), 75.0 ($C_{carborane}$), 59.3 ($C_{carborane}$H), 40.2 (-$CH_2$-), 25.6 ($CH_2$-$CH_2$).

**$^{11}$B NMR** (160 MHz, $CDCl_3$): δ -11.80, -12.64, -14.12, -15.17.

**HRMS** (m/z): $[M+H]^+$ calculated $C_8H_{18}B_{10}NO_4^+$: 300.2234; found: 300.2241.

1,7-Dicarba-*closo*-dodecaboran-1,7-biscarboxylic acid (**11**)

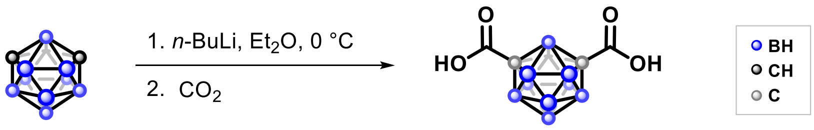

**11** (quant.)

Compound **11** was prepared according to a modified literature procedure.[6] The reaction was carried out under an inert nitrogen atmosphere using standard Schlenk techniques. A 100 mL oven-dried round-bottom flask equipped with a magnetic stir bar and connected to a Schlenk line was evacuated and backfilled with nitrogen. Under a positive nitrogen pressure, the flask was uncapped, and *meta*-carborane (432 mg, 3.0 mmol) was added, followed by dry diethyl ether (30 mL), obtained from a solvent purification system using a syringe. The solution was cooled to 0 °C, and *n*-BuLi (1.6 M in hexanes, 4.0 mL, 6.39 mmol) was added dropwise via syringe. The reaction mixture was stirred at 0 °C for 1 h, then allowed to warm to room temperature and stirred for an additional 30 min. Subsequently, carbon dioxide gas ($CO_2$) was bubbled through the solution at room temperature for 1 h. The solvent was removed under reduced pressure, and water (15 mL) was added to the residue. The aqueous layer was transferred to a separatory funnel and washed with petroleum ether (2 × 15 mL). The aqueous phase was then acidified to pH 4 with 2 M HCl (aq) and extracted with petroleum ether (3 × 30 mL). The combined organic extracts were dried over anhydrous $Na_2SO_4$, filtered, and concentrated under reduced pressure using a rotary evaporator. The crude compound **11** (696 mg, 99%) was obtained as a white solid and used directly in the subsequent synthetic step without further purification.

**IR** ($cm^{-1}$): 2872, 2607, 1716, 1410, 1268, 719.

**$^1$H NMR** (500 MHz, $CDCl_3$): δ 4.19 (br, 2H), 3.1–1.7 (m, 10H, protons of carborane cage). $^1$H NMR shows the presence of residual solvents. NMR data were in agreement with the literature.[7]

1,2-(*N*-succinimidyl dicarbonate)-1,2-dicarba-*closo*-dodecaboran (**7**)

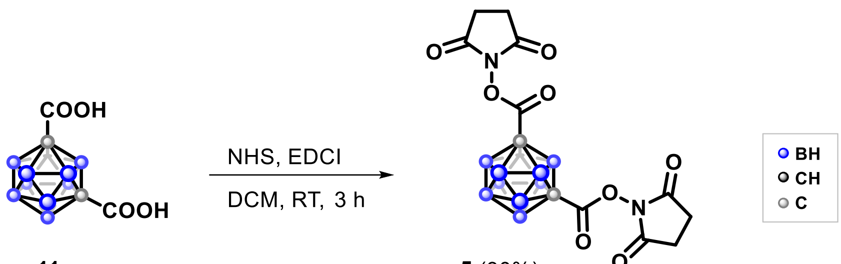

**11** **5** (30%)

Compound **5** was synthesized via a modified literature procedure.[8] The reaction was carried out under an inert nitrogen atmosphere using standard Schlenk techniques. To a 25 mL oven-dried round-bottom flask containing compound **11** (44 mg, 0.19 mmol) and *N*-hydroxysuccinimide (NHS; 53 mg, 0.46 mmol) were added dry dichloromethane (10 mL) and *N*-(3-dimethylaminopropyl)-*N′*-ethylcarbodiimide hydrochloride (EDCI; 87 mg, 0.456 mmol). The reaction mixture was stirred at room temperature for 3 h. Then dichloromethane (10 mL) was added, and the mixture was transferred to a separatory funnel. The organic layer was washed with 1 M HCl (aq) (3 × 10 mL), dried over anhydrous $Na_2SO_4$, filtered, and concentrated under reduced pressure using a rotary evaporator. The crude residue was purified by column chromatography (petroleum ether/ethyl acetate = 1:1), affording compound **5** as a white solid (24 mg, 30%).

**IR** ($cm^{-1}$): 2952, 2619, 1815, 1786, 1741, 1351, 1192, 1095, 1056, 989, 807.

**$^1$H NMR** (500 MHz, $CDCl_3$): δ 2.85 (br, 8H, $CH_2$-$CH_2$), 3.1–1.7 (m, 10H, protons of carborane cage).

**$^{13}$C NMR** (126 MHz $CDCl_3$): δ 167.6 (C=O, CON), 157.6 (C=O, COO), 25.7 ($CH_2$-$CH_2$), carborane carbon resonances are not visible.

**HRMS** (*m/z*): $[M+HCOO]^-$ calcd for $C_{13}H_{19}B_{10}N_2O_{10}^-$ 472.2012 (corresponds to second calcd most abundand *m*/*z*), found: 472.43282 (represents the third most abundant species, relative to the base peak).

<u>Comment</u>: In some cases, the HRMS-determined *m*/*z* values corresponded to the second most abundant isotopic peak rather than the base peak. This was particularly observed in samples containing multiple carborane moieties, which produced complex isotopic patterns due to the presence of ten boron atoms (with natural isotopic abundances of approximately 20% $^{10}$B and 80% $^{11}$B). The resulting overlap of isotopic envelopes from more than one carborane cage contributed to the overall spectral complexity. The structure of compound **5** was further confirmed indirectly through its use in the synthesis of compound **21**.

1,12-Dicarba-*closo*-dodecaboran-1,12-diacetic acid (**13**)

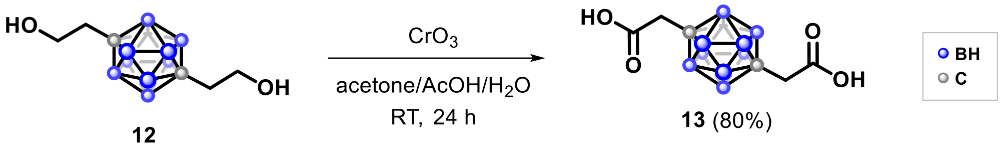

Compound **13** was synthesized via a modified literature procedure.[5] To a 100-mL round-bottom flask containing compound **12** (274 mg, 1.18 mmol), acetone (11 mL) was added, and the solution was cooled to 0 °C. To this mixture a solution of chromium(VI) oxide ($CrO_3$, 826 mg, 8.26 mmol) in a mixture of acetic acid (25 mL) and water (30 mL) was added dropwise. The reaction was then stirred at room temperature overnight. The reaction mixture was extracted with diethyl ether (3 × 40 mL). The combined organic layers were acidified to pH 4 using 4 M HCl (aq) and washed with saturated aqueous NaCl solution (2 × 40 mL). The organic phase was dried over anhydrous $Na_2SO_4$, filtered, and concentrated under reduced pressure using a rotary evaporator. Crude compound **13** (244 mg, 80%) was obtained as a white solid and used in the next step without further purification.

**$^1$H NMR** (500 MHz, $CDCl_3$): δ 3.2–1.7 (protons of carborane cage), 2.53 (s, 4H). NMR data were in agreement with the literature.[5]

1,7-Dicarba-*closo*-dodecaboran-1,7-diacetic acid (**SI-6**)

Diacid **SI-6** was prepared in the same manner as described for compound **13**: **SI-3**: 234 mg, 1 mmol; acetone: 12 mL; $CrO_3$: 750 mg, 7.5 mmol, water: 25 mL; acetic acid: 25 mL.

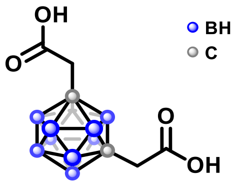

190 mg, 73%.

**$^1$H NMR** (500 MHz, $CDCl_3$): δ 2.91 (s, 4H). NMR data were in agreement with the literature.[5]

1,12-Bis(2,5-dioxopyrolidin-1-yl)acetate-1,12-dicarba-*closo*-dodecaborane (**7**)

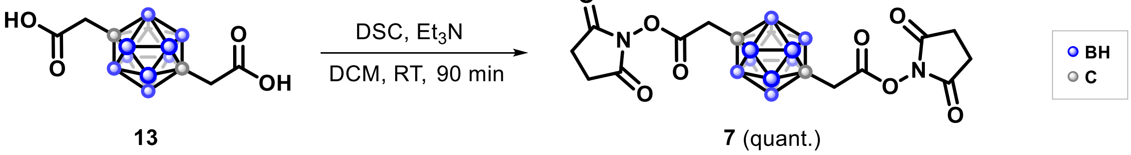

Compound **7** was prepared by modified literature procedure.[9] The reaction was carried out under an inert atmosphere using standard Schlenk techniques. To a 25-mL round-bottom flask containing compound **13** (362 mg, 1.4 mmol) were added dry dichloromethane (6 mL), *N,N'*-disuccinimidyl carbonate (DSC; 860 mg, 3.4 mmol), and dry triethylamine ($Et_3N$; 0.390 mL, 2.8 mmol). The reaction mixture was stirred at room temperature for 90 min. Additional dichloromethane (10 mL) was then added, and the mixture was transferred to a separatory funnel. The organic layer was washed with 1

M HCl (aq) (2 × 10 mL), dried over anhydrous $Na_2SO_4$, filtered, and concentrated under reduced pressure. Compound **9** was obtained as a white solid (637 mg, 99%).

**Mp** 248.6–250.1 °C

**IR** ($cm^{-1}$): 2959, 2608, 1811, 1778, 1738, 1358, 1201, 1123, 1064, 648.

**$^1$H NMR** (500 MHz, $CDCl_3$): δ 3.0–1.9 (m, 10H, protons of carborane cage), 2.89 (br, 4H, $CH_2$), 2.84 (br, 8H, $CH_2$-$CH_2$).

**$^1$H NMR** (500 MHz DMSO-$d_6$): δ 3.17 (br, 4H, $CH_2$), 2.80 (br, 8H, $CH_2$-$CH_2$).

**$^{13}$C NMR** (126 MHz DMSO-$d_6$): δ 170.2 (C=O, CON), 163.9 (C=O, COO), 72.7 ($C_{carboran}$), 38.5 ($CH_2$), 25.9 ($CH_2$-$CH_2$).

**$^{11}$B NMR** (161 MHz, DMSO-$d_6$): δ -11.91, -12.92.

**HRMS-ESI+** (*m/z*): $[M+H]^+$ calculated $C_{14}H_{23}B_{10}N_2O_8^+$, 455.2453; found, 455.2459.

**HRMS-ESI-** (*m/z*): $[M-H]^-$ calculated for $C_{14}H_{21}B_{10}N_2O_8^-$, 453.2306; found, 453.2316.

In the same manner **6** was prepared: **SI-6**: 170 mg, 0.65 mmol; DSC: 400 mg, 1.56 mmol, dichloromethane: 3 mL; $Et_3N$: 0.2 mL, 0.65 mmol.

1,7-Bis(2,5-dioxopyrolidin-1-yl)acetate-1,7-dicarba-*closo*-dodecaborane (**6**)

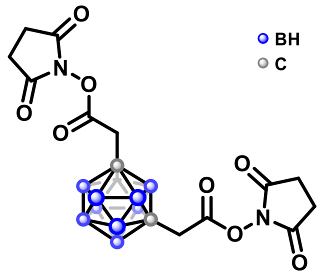

140 mg; 47%

**Mp** 165–173 °C

**IR** ($cm^{-1}$): 2604, 1812, 1782, 1732, 1362, 1200, 1097, 1063, 1047, 644.

**$^1$H NMR** (500 MHz, $CDCl_3$): δ 3.6–1.8 (protons of carborane cage), 3.21 (br, 4H, $CH_2$), 2.84 (br, 8H, $CH_2$-$CH_2$).

**$^{13}$C NMR** (151 MHz, $CDCl_3$): δ 168.4 (C=O, CON), 162.8 (C=O, COO), 67.5 ($C_{carboran}$), 38.5 ($CH_2$), 25.6 ($CH_2$-$CH_2$).

**$^{11}$B NMR** (160 MHz, $CDCl_3$): δ -5.7, -10.0, -10.86, -12.28, -13.45.

**HRMS** (*m/z*): $[M+H]^+$ calculated for $C_{14}H_{23}B_{10}N_2O_8^+$: 455.2452; found: 455.2465.

1,2-Bis(2,5-dioxopyrolidin-1-yl)acetate-1,2-dicarba-*closo*-dodecaborane (**6**)

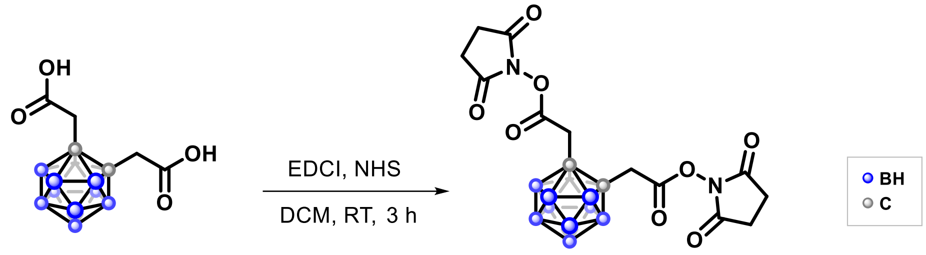

**SI-7** **4** (10%)

Compound **4** was synthesized via a modified literature procedure.[8] The reaction was carried out under an inert atmosphere using standard Schlenk techniques. To a 25 mL round-bottom flask containing compound **SI-7** (36 mg, 0.14 mmol) and *N*-hydroxysuccinimide (NHS; 48 mg, 0.41 mmol) were added dry dichloromethane (2 mL) and *N*-ethyl-*N′*-(3-dimethylaminopropyl)carbodiimide hydrochloride (EDCI; 79 mg, 0.41 mmol). The reaction mixture was stirred at room temperature for 3 h. Additional dichloromethane (10 mL) was then added, and the mixture was transferred to a separatory funnel. The organic layer was washed with 1 M HCl (aq) (3 × 10 mL), dried over anhydrous $Na_2SO_4$, filtered, and concentrated under reduced pressure using rotary evaporator. The crude product was purified by reverse-phase flash chromatography (ACN/$H_2O$ + 0.1% TFA) to afford compound **4** as a white solid (6 mg, 10%).

**IR** ($cm^{-1}$): 2963, 2591, 1712, 1259, 1200, 1067, 1022, 797, 645.

**$^1$H NMR** (600 MHz, $CDCl_3$): δ 3.32 (br, 4H, -$CH_2$-), 2.84 (br, 8H, $CH_2$-$CH_2$).

**$^{13}$C NMR** (151 MHz, $CDCl_3$): δ 179.6 (C=O, COO), 169.2 (C=O, CON), 69.7 ($C_{carboran}$), 41.5 ($CH_2$), 25.6 ($CH_2$-$CH_2$).

**$^{11}$B NMR** (160 MHz, $CDCl_3$) δ -4.09, -5.01, -8.01, -8.95, -10.59, -11.61.

Comment: Compound **4** was found to be relatively unstable, as indicated by the $^1$H and $^{13}$C NMR spectra, which showed partial degradation of **4** back to **SI-7** and NHS.

*N*-(Benzyl)-1,7-dicarba-*closo*-dodecaboran-1-carboxamide (**25**)

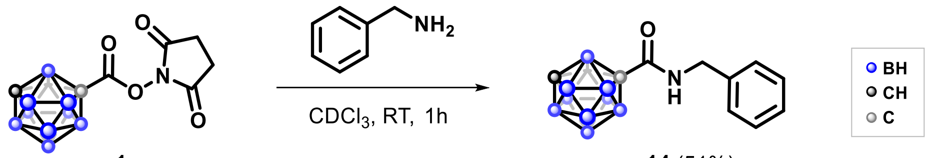

**1** **14** (51%)

Compound **1** (10 mg, 0.035 mmol) was dissolved in $CDCl_3$ (1 mL), and benzylamine (35 µL, 0.32 mmol) was added at room temperature. The mixture was stirred at room temperature for 1 h. The crude product was purified by column chromatography (gradient: A = petroleum ether; B = 1:1 petroleum ether/ethyl acetate) to afford compound **14** as a white solid (5 mg, 51%).

**Mp** 101–103 °C

**IR** ($cm^{-1}$): 3343, 3059, 2929, 2605, 2567, 1668, 1526, 1451, 1282, 1259, 714.

**$^1$H NMR** (500 MHz, $CDCl_3$): δ 7.37–7.28 (m, 3H, 3×$H_{aromatic}$), 7.21–7.16 (m, 2H, 2×$H_{aromatic}$), 6.08 (br, 1H, NH), 4.39 (d, *J* = 5.7 Hz, 2H, $CH_2$), 3.04 (br, 1H, $CH_{carboran}$) 3.4–1.6 (m, 10H, protons of carborane cage).

**$^{13}$C NMR** (126 MHz, $CDCl_3$): δ 160.3 (C=O), 136.8 ($C_{aromatic}$), 128.9 ($C_{aromatic}$), 127.9 ($C_{aromatic}$), 127.4 ($C_{aromatic}$), 75.4 ($C_{carboran}$), 54.9 ($C_{carboran}$H), 44.7 ($CH_2$).

**$^{11}$B NMR** (161 MHz, $CDCl_3$): δ -5.3, -6.4, -7.9, -10.4, -11.2, -12.1, -12.6, -13.6, -15.0, -16.1.

**HRMS** (*m/z*): $[M+H]^+$ calculated for $C_{10}H_{20}B_{10}NO^+$, 278.2543; found, 278.2542.

*N*-(3-phenylpropyl)-1,7-dicarba-*closo*-dodecaboran-1-carboxamide (**15**)

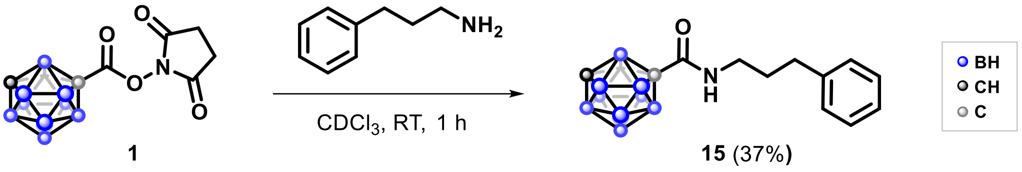

Compound **1** (10 mg, 0.035 mmol) was dissolved in $CDCl_3$ (1 mL), and 3-phenyl-1-propylamine (25 µL, 0.176 mmol) was added. The mixture was stirred at room temperature for 1 h. The crude product was purified by filtration through a short plug of silica gel (eluent: DCM) to afford compound **15** as a white solid (4 mg, 37%).

**Mp** 91–92 °C

**IR** ($cm^{-1}$): 3342, 3025, 2953, 2590, 1661, 1531, 1453, 1287, 1264, 1074, 1029, 714.

**$^1$H NMR** (500 MHz, $CDCl_3$): δ 7.31–7.27 (m, 2H, 2×$H_{aromatic}$), 7.22–7.19 (m, 1H, 1×$H_{aromatic}$), 7.15–7.14 (m, 2H, 2×$H_{aromatic}$), 5.77 (br s, 1H, NH), 3.22 (q, *J* = 7.0 Hz, 2H, $CH_2$), 3.01 (br s, 1H, $CH_{carboran}$), 3.0–1.6 (m, 10H, protons of carborane cage), 2.61 (t, *J* = 7.5 Hz, 2H, $CH_2$), 1.82 (quint, *J* = 7.5 Hz, 2H, $CH_2$).

**$^{13}$C NMR** (126 MHz $CDCl_3$): δ 160.2 (CON), 140.8 ($C_{aromatic}$), 128.6 ($C_{aromatic}$), 128.3 ($C_{aromatic}$), 126.2 ($C_{aromatic}$), 75.6 ($C_{carboran}$), 54.9 ($C_{carboran}$H), 40.4 ($CH_2$), 33.1 ($CH_2$), 30.6 ($CH_2$).

**$^{11}$B NMR** (161 MHz, $CDCl_3$): δ -5.27, -6.28, -6.91, -7.96, -10.36, -11.22, -12.09, -12.61, -13.66, -15.02, -16.16.

**$^{15}$N NMR** ($^1$H–$^{15}$N HMBC, $CDCl_3$): δ +110.

**HRMS** (*m/z*): $[M+H]^+$ calculated for $C_{12}H_{24}B_{10}NO^+$, 306.2856; found, 306.2857.

*N*-(3-Phenylpropyl)-1-dicarba-*closo*-dodecaborane-1-acetamide (**16**)

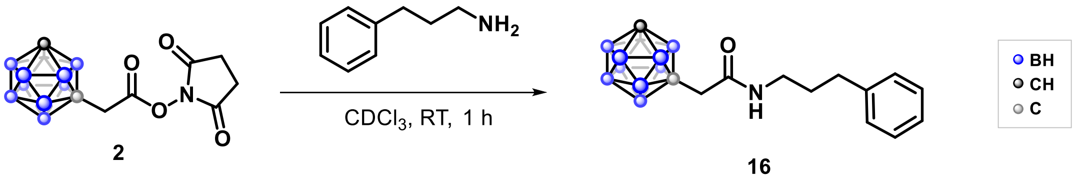

Compound **2** (12 mg, 0.04 mmol) was dissolved in $CDCl_3$ (1 mL), and 3-phenyl-1-propylamine (25 µL, 0.175 mmol) was added. The mixture was stirred at room temperature for 1 h. The crude product was purified by filtration through a short plug of silica gel (eluent: DCM) to afford compound **16** as a white solid (5 mg, 39%).

**Mp** 100–103 °C.

**IR** ($cm^{-1}$): 3247, 3044, 2961, 2592, 1632, 1572, 1282, 1088, 1015, 796, 730, 694.

**$^1$H NMR** (500 MHz, $CDCl_3$): δ 7.31–7.28 (m, 2H, 2×$H_{aromatic}$), 7.22–7.17 (m, 3H, 3×$H_{aromatic}$), 5.35 (br s, 1H, NH), 3.29–3.26 (m, 2H, -$CH_2$-), 2.96 (br s, 1H, $CH_{carborane}$), 3.0–1.6 (m, 10H, protons of carborane cage), 2.70–2.63 (m, 4H, -$CH_2$-), 1.86 (quint, $J$ = 7.4 Hz, 2H, -$CH_2$-).

**$^{13}$C NMR** (126 MHz, $CDCl_3$): δ 165.5 (CON), 140.1 ($C_{aromatic}$), 127.5($C_{aromatic}$), 127.3 ($C_{aromatic}$), 125.1 ($C_{aromatic}$), 69.4 ($C_{carboran}$), 54.4 ($C_{carboran}$H), 43.5 ($CH_2$), 38.5 ($CH_2$), 32.3 ($CH_2$), 29.9 ($CH_2$).

**$^{11}$B NMR** (160 MHz, $CDCl_3$): δ -3.7, -10.0, -13.3, -15.1.

**HRMS** (m/z): $[M+H]^+$ calculated for $C_{13}H_{25}B_{10}NO^+$: 321.2867; found: 321.2983.

*N*-(1,7-Dicarba-*closo*-dodecaboranylcarbonyl)-D-phenylalanine (**17**)

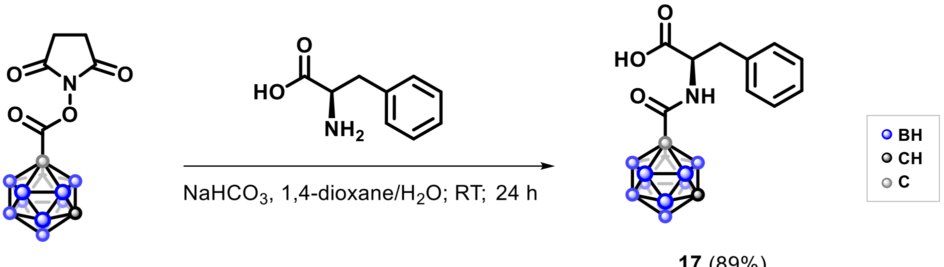

2

**17** (89%)

To a solution of compound **1** (23 mg, 0.08 mmol) in 1,4-dioxane (1 mL) in a 25-mL reaction flask, was added a solution of $NaHCO_3$ (21 mg, 0.256 mmol) and D-phenylalanine (26.4 mg, 0.16 mmol) in a $H_2O$/1,4-dioxane mixture (8 mL, 3:5 v/v). The reaction mixture was stirred at room temperature overnight. The reaction was then acidified to pH 4 with 4 M HCl (aq) (0.2 mL). The aqueous layer was extracted with ethyl acetate (3 × 15 mL), and the combined organic layers were washed with saturated NaCl (aq) (1 × 15 mL), dried over anhydrous $Na_2SO_4$, filtered, and concentrated under reduced pressure. Compound **17** was obtained as a solid (24 mg, 89%).

**Mp** 215–217°C

**IR** ($cm^{-1}$): 2922, 2852, 2602, 1721, 1672, 1515, 1408, 1215, 1096, 732, 698.

**$^1$H NMR** (500 MHz, $CDCl_3$): δ 7.34–7.29 (m, 3H, 3×$H_{aromatic}$), 7.10–7.08 (m, 2H, 2×$H_{aromatic}$), 6.19 (d, $J$ = 7.1 Hz, 1H, NH), 4.75–4.71 (m, 1H, CH), 3.22 (dd, $J$ = 14.0, 8.5 Hz; 1H, CH), 3.10 (dd, $J$ = 14.0, 6.0 Hz, 1H, CH), 3.02 (br, 1H, $CH_{carboran}$), 3.0–1.6 (m, 10H, protons of carborane cage).

**$^{13}$C NMR** (126 MHz, $CDCl_3$): δ 174.7 (C=O, COOH), 159.9 (C=O, CON), 134.7 ($C_{aromatic}$), 129.3 ($C_{aromatic}$), 128.8($C_{aromatic}$), 127.6 ($C_{aromatic}$), 54.9 ($C_{carboran}$H), 53.9 (CH), 36.9 ($CH_2$), 31.0 ($C_{carboran}$).

**$^{11}$B NMR** (161 MHz, $CDCl_3$): δ -4.02, -5.17, -6.16, -10.16, -11.07, -12.59, -13.62, -15.02, -16.16.

**HRMS** (*m/z*): $[M+H]^+$ calculated for $C_{12}H_{22}B_{10}NO_3^+$, 336.2598; found, 336.2602.

Synthesis of amikacin carborane conjugates **18**–**20**

Amikacin (40 mg, 0.068 mmol) was dissolved in DMSO (1.2 mL), and compound **2** (25 mg, 0.084 mmol) was added to the solution, in one portion, at room temperature. The reaction mixture was stirred at room temperature overnight (18 h). HPLC-MS indicated no remaining starting material and the reaction mixture was loaded onto and purified by reverse-phase flash chromatography (ACN/$H_2O$ + 0.1% TFA) to afford amikacin carborane conjugates **18**, **19** and **20**.

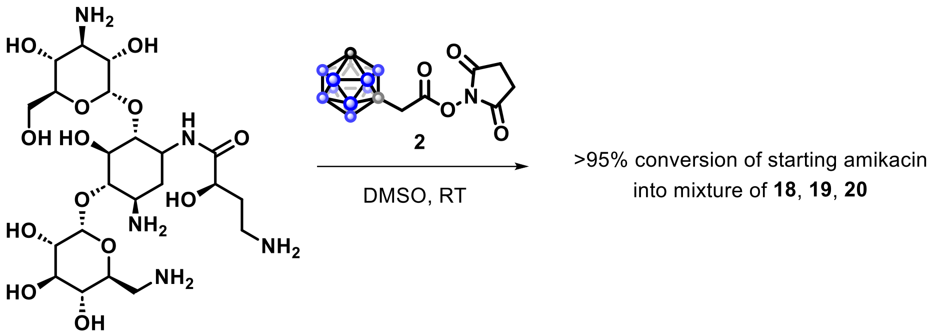

**18**:

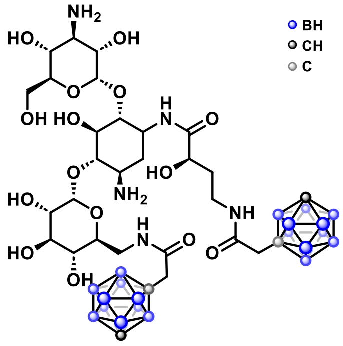

10 mg, 15%;

**$^{1}$H NMR** (500 MHz, DMSO-*d*$_6$): complex spectrum.

**$^{13}$C NMR** (126 MHz, DMSO-*d*$_6$): δ 174.3, 167.1, 166.7, 101.8, 98.2, 91.2, 80.6, 75.5, 73.4, 73.1, 72.8, 72.3, 71.9, 71.7, 70.0, 69.8, 61.0, 56.9, 56.9, 55.3, 50.2, 49.3, 43.1, 42.7, 40.8, 36.3, 34.2. Three carbon resonances are missing or overlapping.

**HRMS** (*m/z*): $[M-H]^-$ calculated for $C_{30}H_{66}B_{20}N_5O_{15}^-$, 952.6567; found, 952.6585.

**19**:

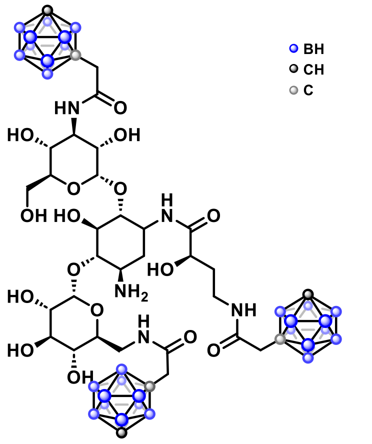

12 mg, 16%.

**$^{1}$H NMR** (500 MHz, DMSO-*d6*): complex spectrum.

**$^{13}$C NMR** (126 MHz, DMSO-*d6*): δ 174.5, 167.8, 167.2, 166.8, 98.1, 80.6, 79.6, 79.4, 79.2, 74.7, 73.3, 72.7, 72.3, 72.2, 71.8, 71.7, 70.7, 69.7, 67.7, 60.6, 56.9, 55.0, 49.3, 43.6, 43.0, 42.7, 36.1, 34.0. Six carbon resonances are missing or overlapping.

**HRMS** (*m/z*): [M-H]$^-$ calculated for $C_{34}H_{78}B_{30}N_5O_{16}^-$, 1137.8422; found, 1137.8435.

**20**:

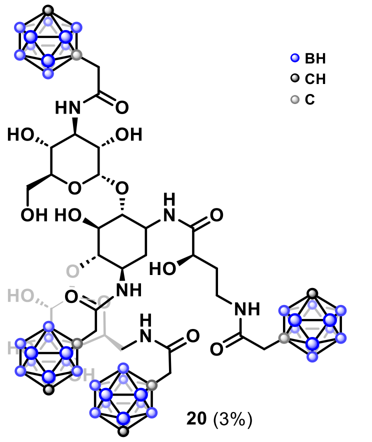

**20** (3%)

3 mg, 3%

**$^{1}$H and $^{13}$C NMR**: complex spectra.

**HRMS** (*m/z*): [M-H]$^-$ calculated for $C_{38}H_{90}B_{40}N_5O_{17}^-$, 1322.0314; found, 1322.0292.

*N*-(3-phenylpropyl)-1,7-dicarba-*closo*-dodecaboran-1-carboxamide (**21**)

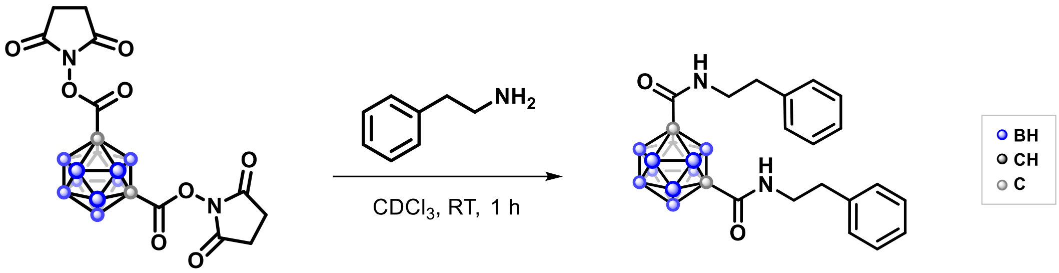

**5** **21** (36%)

Compound **5** (10.5 mg, 0.025 mmol) was dissolved in $CDCl_3$ (2 mL), and 2-phenyl-1-ethylamine (20 µL, 0.16 mmol) was added. The mixture was stirred at room temperature for 1 h. The crude product was purified by filtration through a short plug of silica gel (eluent: ethyl acetate, then acetone) to afford compound **21** as a colorless oil (4 mg, 36%).

**IR** ($cm^{-1}$): 3368,, 2928, 2600, 1670, 1514, 1454, 1265, 1030, 742, 698.

**$^1$H NMR** (500 MHz, $CDCl_3$): δ 7.35–7.30 (m, 4H, 4×$H_{aromatic}$), 7.28–7.24 (m, 2H, 2×$H_{aromatic}$), 7.16–7.12 (m, 4H, 4×$H_{aromatic}$), 5.76 (br, 2H, 2×NH), 3.47–3.44 (4H, $CH_2$), 2.78 (t, $J$ = 6.7 Hz, 4H, $CH_2$).

**$^{13}$C NMR** (126 MHz $CDCl_3$): δ 159.7 (CON), 138.1 ($C_{aromatic}$), 129.0 ($C_{aromatic}$), 128.9 ($C_{aromatic}$), 127.0 ($C_{aromatic}$), 75.2 ($C_{carboran}$), 42.2 ($CH_2$), 35.2 ($CH_2$).

**$^{11}$B NMR** (161 MHz, $CDCl_3$): δ -5.17, -6.21, -11.12, -13.70, -14.87, -16.16.

**HRMS** (*m/z*): $[M+H]^+$ calculated for $C_{20}H_{31}B_{10}N_2O_2^+$, 439.3383; found, 439.3394.

Synhesis of compound **22**

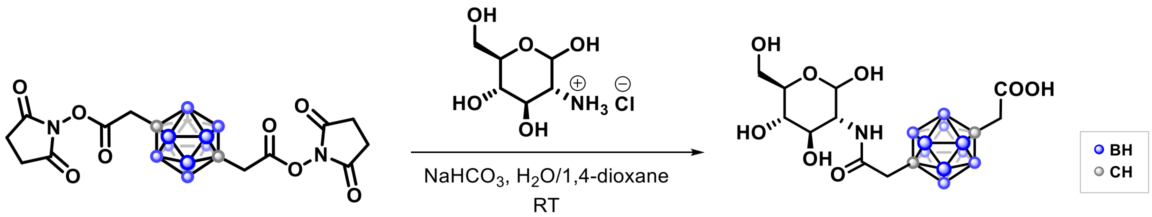

**7** **22** (22%)

To a 10 mL flask containing glucosamine hydrochloride (47 mg, 0.22 mmol) in an aqueous $NaHCO_3$ (55.4 mg, 0.66 mmol) solution (2 mL) was added compound **7** (25 mg, 0.055 mmol). 1,4-Dioxane was added dropwise until the solution became clear. The reaction mixture was stirred at room temperature overnight. The reaction was then acidified to pH 4 with 2 M HCl (aq) (0.3 mL). The aqueous layer was extracted with ethyl acetate (3 × 15 mL), and the combined organic layers were washed with saturated NaCl (aq) (10 mL), dried over anhydrous $Na_2SO_4$, filtered, and concentrated under reduced pressure using rotary evaporator. The crude product was purified by reverse-phase flash chromatography (ACN/Milli-Q + 0.1% TFA) to afford compound **22** as a white solid (5 mg, 22%).

Comment: Along with the desired coupling reaction with glucosamine, hydrolysis of the other NHS ester occurred (before coupling of another molecule of glucosamine) under the aqueous conditions, yielding the corresponding carboxylic acid.

$^{1}$**H NMR** (500 MHz, DMSO-$d_6$): δ 7.61 (d, 1H), 6.39 (br, 1H), 4.79 (br, 1H), 3.59 (d, 1H), 3.55–3.47 (m, 5H), 3.02 (t, 2H), 2.31 (d, 3H).

$^{13}$**C NMR** (126 MHz, DMSO-$d_6$): δ 169.2, 166.6, 91.0, 72.5, 71.6, 70.78, 61.6, 54.8, 43.5, 43.1, 31.2. One carbon resonance is missing or overlapping.

$^{11}$**B NMR** (161 MHz, DMSO-$d_6$): δ -12.00, -12.99.

**HRMS** (*m/z*): $[M+H]^+$ calculated for $C_{12}H_{28}B_{10}NO_8^+$, 422.2813; found, 422.2825.

*N*-(3-Phenylpropyl)-1,7-dicarba-*closo*-dodecaborane-1,7-bis(acetamide) (**23**)

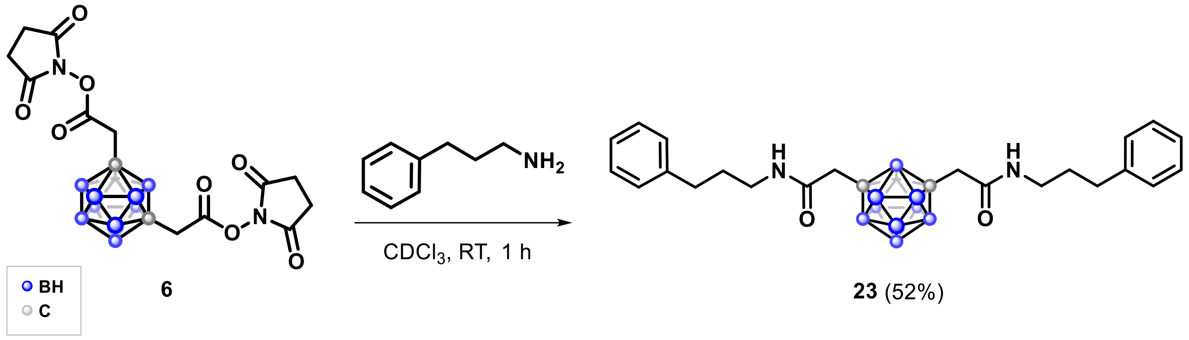

Compound **6** (7 mg, 0.015 mmol) was dissolved in $CDCl_3$ (1 mL), and 3-phenyl-1-propylamine (29 μL, 0.2 mmol) was added. The mixture was stirred at room temperature for 1 h. The crude product was purified by filtration through a short plug of silica gel (gradient elution: 100% DCM, DCM/ACN = 1/1, 100% ACN) to afford compound **23** as a colorless gel (4 mg, 52%).

**IR** ($cm^{-1}$): 3292, 3084, 2937, 2594, 1649, 1555, 1453, 1283, 740, 698.

$^{1}$**H NMR** (600 MHz, $CDCl_3$): δ 7.30–7.28 (m, 4H, 4×$H_{aromatic}$), 7.21–7.17 (m, 6H, 6×$H_{aromatic}$), 5.38 (br s, 2H, 2×NH), 3.27–3.24 (m, 4H, $CH_2$), 2.67–2.64 (m, 8H, $CH_2$), 1.87–1.82 (m, 4H, $CH_2$).

$^{13}$**C NMR** (151 MHz, $CDCl_3$): δ 166.5 (C=O, CON), 141.2 ($C_{aromatic}$), 128.6 ($C_{aromatic}$), 128.4 ($C_{aromatic}$), 126.1 ($C_{aromatic}$), 70.5 ($C_{carborane}$), 44.4 ($CH_2$), 39.6 ($CH_2$), 33.3($CH_2$), 30.9 ($CH_2$).

$^{11}$**B NMR** (160 MHz, $CDCl_3$): δ -5.69, -10.00, -10.86, -12.28, -13.45.

$^{15}$**N NMR** ($^{1}$H– $^{15}$N HMBC, $CDCl_3$): δ +120.

**HRMS** (m/z): $[M+H]^+$ calculated for $C_{24}H_{39}B_{10}N_2O_2^+$: 496.3937; found: 496.3989.

*N*-(3-Phenylpropyl)-1,12-dicarba-*closo*-dodecaborane-1,12-bis(acetamide) (**24**)

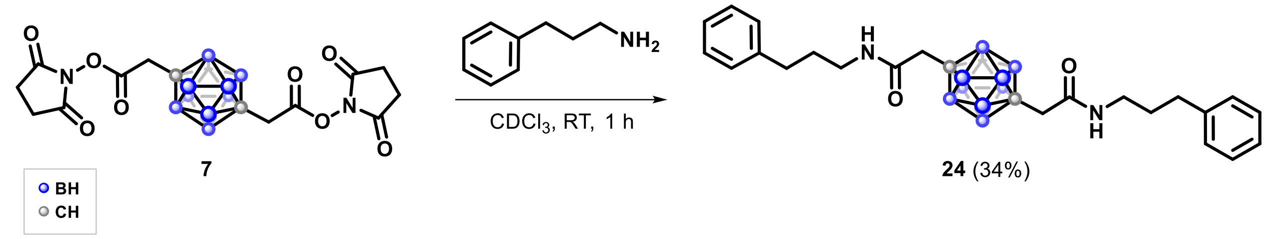

7

**24** (34%)

Compound **7** (10 mg, 0.022 mmol) was dissolved in $CDCl_3$ (1 mL), and 3-phenyl-1-propylamine (25 µL, 0.176 mmol) was added at room temperature. The mixture was stirred at room temperature for 1 h, and the crude product was purified by filtration through a short plug of silica gel (eluent: DCM) to afford compound **24** as an off-white oil (17 mg, 34%).

**IR** ($cm^{-1}$): 3294, 3085, 3026, 2926, 2856, 2599, 1649, 1549, 1496, 1453, 1437, 1365, 1091, 1045, 739, 697.

**$^1$H NMR** (500 MHz, $CDCl_3$): δ 7.31–7.27 (m, 4H, 4×$H_{aromatic}$), 7.22–7.15 (m, 6H, 6×$H_{aromatic}$), 5.23 (br s, 2H, 2×NH), 3.22–3.20 (m, 4H, $CH_2$), 2.64 (m, 4H, $CH_2$), 2.36 (s, 4H, $CH_2$) 1.84–1.80 (m, 4H, $CH_2$).

**$^{13}$C NMR** (126 MHz, $CDCl_3$): δ 166.5 (C=O, CON), 141.2 ($C_{aromatic}$), 128.6 ($C_{aromatic}$), 128.4 ($C_{aromatic}$), 126.1 ($C_{aromatic}$), 45.3 ($C_{carborane}$), 45.2 ($CH_2$), 39.5 ($CH_2$), 33.4 ($CH_2$), 30.9 ($CH_2$).

**$^{11}$B NMR** (161 MHz, $CDCl_3$): δ -11.92, -12.97.

**HRMS** (*m/z*): $[M+H]^+$ calculated for $C_{24}H_{39}B_{10}N_2O_2^+$, 495.4010; found, 495.4006.

#### Peptide preparation and their reactions with carborane NHS reagents (Fig. 3)

##### General procdures for peptides synthesis

Peptides were synthesized either manually using a pressurized manifold or on an automated Liberty Blue 2.0 peptide synthesizer (CEM) via standard Fmoc/tBu solid-phase peptide synthesis (SPPS). The syntheses were performed in DMF using Fmoc-protected amino acids, DIC/Oxyma as the coupling system, and piperidine for Fmoc deprotection. Rink Amide AM resin (Sigma-Aldrich, loading capacity 0.6 mmol/g.) was used as the solid support. Fmoc-protected amino acids were obtained from BLD-Pharm or Fluorochem, including: Fmoc-Ala-OH, Fmoc-Asn(Trt)-OH, Fmoc-Asp(OtBu)-OH, Fmoc-Gln(Trt)-OH, Fmoc-Gly-OH, Fmoc-His(Trt)-OH, Fmoc-Ile-OH, Fmoc-Leu-OH, Fmoc-Lys(Alloc)-OH, Fmoc-Lys(Boc)-OH, Fmoc-Phe-OH, Fmoc-Pro-OH, Fmoc-Ser(tBu)-OH, Fmoc-Tyr(tBu)-OH, and Fmoc-Val-OH.

For N-terminal acetylation, the N-terminus was capped with acetic anhydride after Fmoc deprotection of the final amino acid residue. Before cleavage and global deprotection, the peptide-resin was washed with DCM and dried under vacuum for 10–15 min. Cleavage from the resin and global deprotection were performed in a plastic vial placed on a vacuum collection chamber. A TFA/TIS/$H_2O$ (95:2.5:2.5 v/v/v) cleavage cocktail was then added, and the mixture was agitated on a mechanical shaker for 1–2 h at room temperature. After cleavage, the resin was filtered off, and the filtrate was evaporated under inert gas. The crude peptide was precipitated with cold $Et_2O$ (0 °C), centrifuged at 11k rpm for 5 min, and washed with cold $Et_2O$ once more. The resulting solid was dried under vacuum and analyzed by HPLC–MS. Crude peptides were purified by reverse-phase flash chromatography using a puriFlash XS520Plus system. Collected fractions were analyzed by HPLC–MS, combined, flash-frozen in liquid nitrogen, and lyophilized using a FreeZone (LABCONCO) freeze dryer. The final peptides were characterized by HPLC–MS and analytical RP-HPLC.

General procedure for On-resin Alloc deprotection: Two 5–10 mL reaction flasks were cooled under argon and sealed with septa connected to a Schlenk line under a slight nitrogen overpressure. Dry dichloromethane (1 mL) was added to one flask and 4 mL to the other. The first solution was charged with phenylsilane (24 equiv. relative to resin) and added to the resin (placed in a syringe within the vacuum chamber). The second flask was charged with $Pd(PPh_3)_4$ (0.6 equiv) and then added to the resin. The mixture was gently agitated for 30 min with occasional mixing and intermittent argon flushing. The solution was then removed under vacuum on the manifold. The described procedure was repeated twice. The resin was finally washed thoroughly with dichloromethane and dried under inert atmosphere.

##### Ac-Ser-Ala-Lys-Ala-Gly-Ser-Gly-Tyr-Lys-Ser-Ala-$CONH_2$ (Peptide A)

Rink Amide resin: 167 mg (0.1 mmol scale).

For the synthesis of Peptide A, the following amino acids were used in sequence from the C- to the N-terminus: Fmoc-Ala-OH, Fmoc-Ser(tBu)-OH, Fmoc-Lys(Alloc)-OH, Fmoc-Tyr(tBu)-OH, Fmoc-Gly-OH, Fmoc-Ser(tBu)-OH, Fmoc-Gly-OH, Fmoc-Ala-OH, Fmoc-Lys(Alloc)-OH, Fmoc-Ala-OH, Fmoc-Ser(tBu)-OH.

Alloc deprotection, N-capping, cleavage from resin and global deprotection, along with purification were performed as described in General procedures for peptide synthesis.

**Ac-Val-Ser-Ala-Lys-Val-Lys-Ile-Gly-Tyr-Gly-$CONH_2$ (Peptide B)**

Rink Amide AM: 333 mg (0.2 mmol scale).

For the synthesis of Peptide B, the following amino acids were used in sequence from the C- to the N-terminus: Fmoc-Gly-OH, Fmoc-Tyr(tBu)-OH, Fmoc-Gly-OH, Fmoc-Ile-OH, Fmoc-Lys(Boc)-OH, Fmoc-Val-OH, Fmoc-Lys(Boc)-OH, Fmoc-Ala-OH, Fmoc-Ser(*t*Bu)-OH, Fmoc-Val-OH.

N-capping, cleavage from resin and global deprotection, along with purification were performed as described in General procedures for peptide synthesis.

#### HPLC

Agilent Infinity II 1260 LC system, Atlantis C18 column (5 µm, 4.6 × 250 mm). Elution was performed with a gradient of acetonitrile (ACN) and water (both containing 0.1% TFA): 20–80% ACN over 20 min, at a flow rate of 0.75 mL/min. Injection 5 µL of 1 mg/mL solution of peptide.

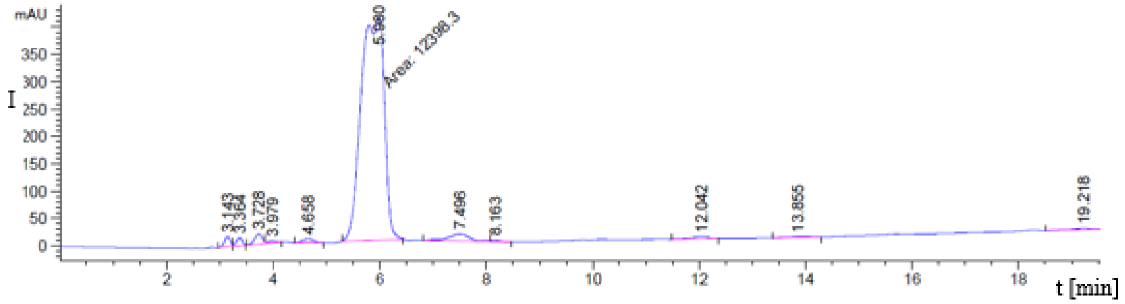

**Figure S1**. HPLC trace of the purified Peptide B (y-axis: absorbance, wavelength = 220 nm; x-axis: time in minutes).

**MS-ESI** (*m*/*z*):

[M+H]+ calculated for $C_{49}H_{84}N_{13}O_{13}^{+}$, 1062.6; found, 1062.7.

$[M+2H]^{2+}$ calculated for $C_{49}H_{85}N_{13}O_{13}^{2+}$, 531.8; found, 532.0.

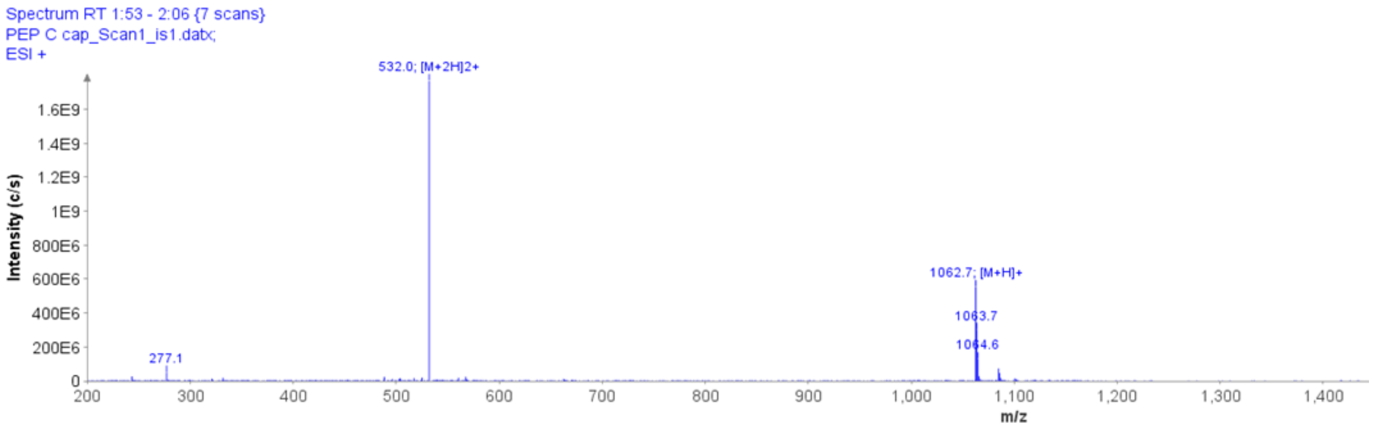

**Figure S2**. MS spectrum of purified the purified Peptide B (y-axis: total ion count (TIC), x-axis: *m*/*z*).

**Ac-Val-Ser-Ala-Lys-Ala-Gly-Tyr-Val-CONH2 (Peptide C)**

Rink Amide AM: 167 mg (0.1 mmol scale).

For the synthesis of Peptide C, the following amino acids were used in sequence from the C- to the N-terminus: Fmoc-Val-OH, Fmoc-Tyr(tBu)-OH, Fmoc-Gly-OH, Fmoc-Ala-OH, Fmoc-Lys(Boc)-OH, Fmoc-Ala-OH, Fmoc-Ser(tBu)-OH, Fmoc-Val-OH.

N-capping, cleavage from resin and global deprotection, along with purification were performed as described in General procedures for peptide synthesis.

**HPLC-MS:**

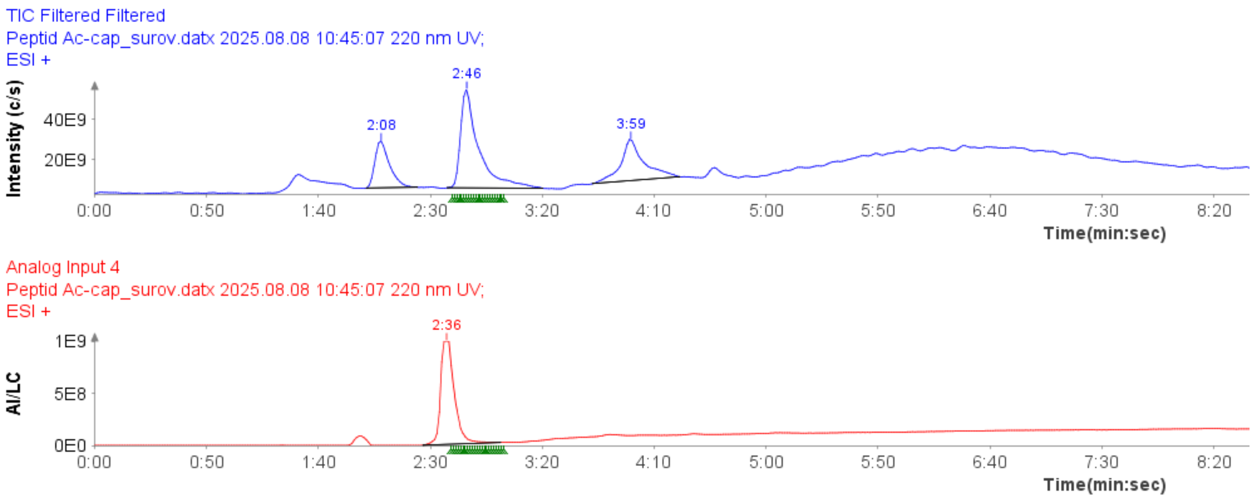

**Figure S3.** HPLC-MS of Peptide C, TIC (blue) and HPLC trace (red).

**MS-ESI** (*m*/*z*):

[M+H]+ calculated for $C_{38}H_{63}N_{10}O_{11}^{+}$, 835.4; found, 835.6.

$[M+2H]^{2+}$ calculated for $C_{38}H_{64}N_{10}O_{11}^{2+}$, 418.2; found, 418.4.

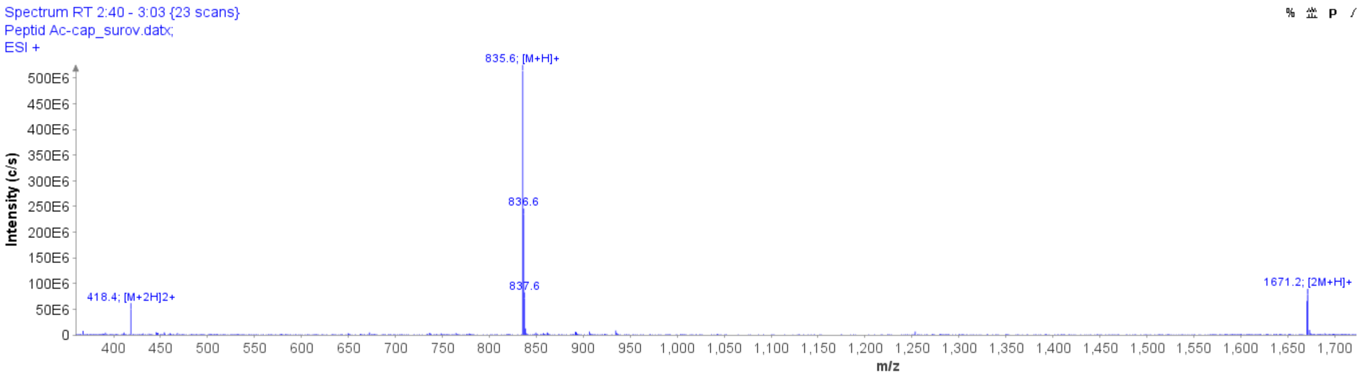

**Figure S4.** MS spectrum of purified the Peptide C (y-axis: total ion count (TIC), x-axis: *m*/*z*) of the above HPLC-MS analysis.

**Ac-FNAPFDVGIKLSGVQYQQHSQAL-CONH2 (Peptide D)**

Rink Amide AM: 167 mg (0.1 mmol scale).

For the synthesis of Peptide D, the following amino acids were used in sequence from the C- to the N-terminus: Fmoc-Leu-OH, Fmoc-Ala-OH, Fmoc-Gln(Trt)-OH, Fmoc-Ser(tBu)-OH, Fmoc-His(Trt)-OH, Fmoc-Gln(Trt)-OH, Fmoc-Gln(Trt)-OH, Fmoc-Tyr(tBu)-OH, Fmoc-Gln(Trt)-OH, Fmoc-Val-OH, Fmoc-Gly-OH, Fmoc-Ser(tBu)-OH, Fmoc-Leu-OH, Fmoc-Lys(Boc)-OH, Fmoc-Ile-OH, Fmoc-Gly-OH, Fmoc-Val-OH, Fmoc-Asp(OtBu)-OH, Fmoc-Phe-OH, Fmoc-Pro-OH, Fmoc-Ala-OH, Fmoc-Asn(Trt)-OH, Fmoc-Phe-OH.

N-capping, cleavage from resin and global deprotection, along with purification were performed as described in General procedures for peptide synthesis.

**HPLC-MS**

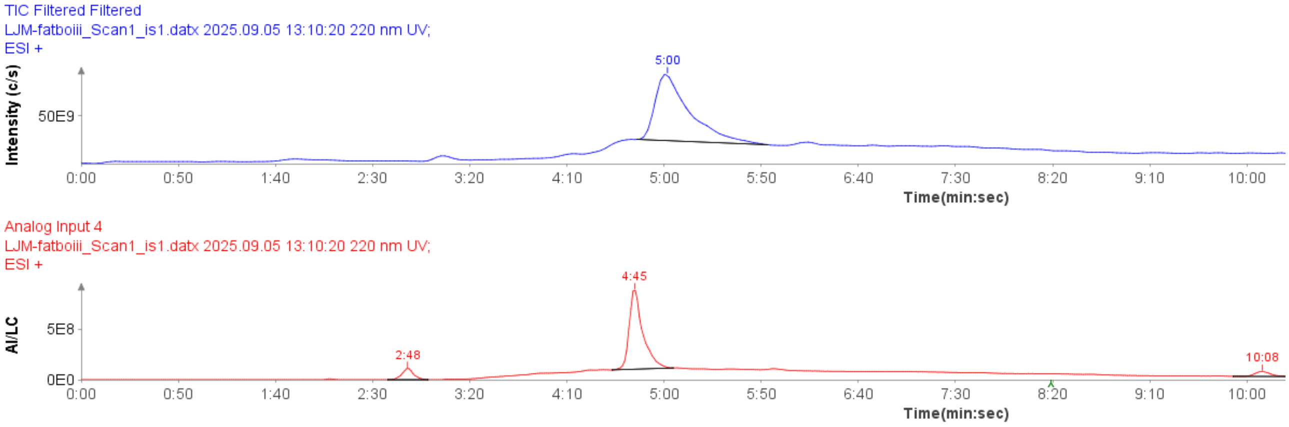

**Figure S5.** HPLC-MS of purified Peptide D, TIC (blue) and HPLC trace (red).

**MS-ESI** (*m*/*z*):

$[M+2H]^{2+}$ calculated for $C_{118}H_{180}N_{32}O_{34}{}^{2+}$, 1295.2; found, 1295.7.

$[M+3H]^{3+}$ calculated for $C_{118}H_{181}N_{32}O_{34}{}^{3+}$, 863.8; found, 864.1.

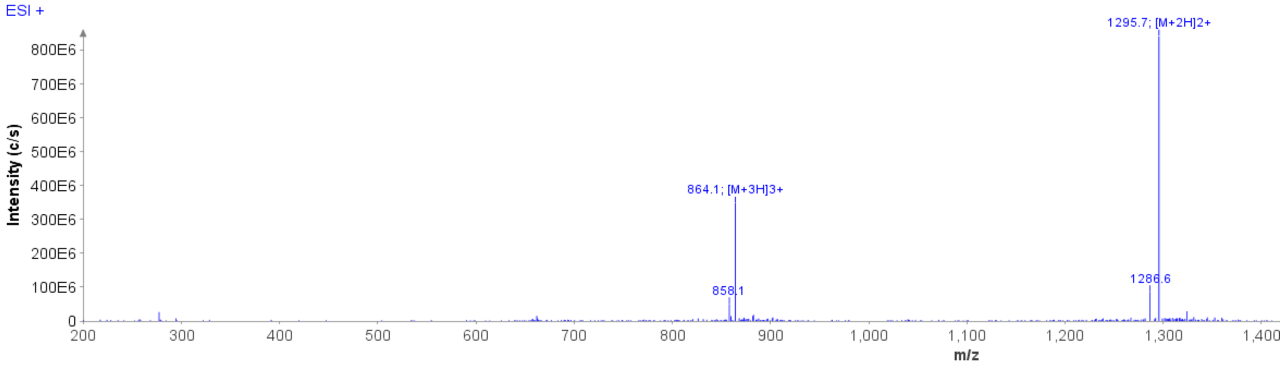

**Figure S6.** MS spectrum of purified the Peptide D (y-axis: total ion count (TIC), x-axis: *m*/*z*) of the above HPLC-MS analysis.

[M+H]+ calculated for $C_{118}H_{179}N_{32}O_{34}^{+}$, 2589.3; found (deconvoluted) 2589.3.

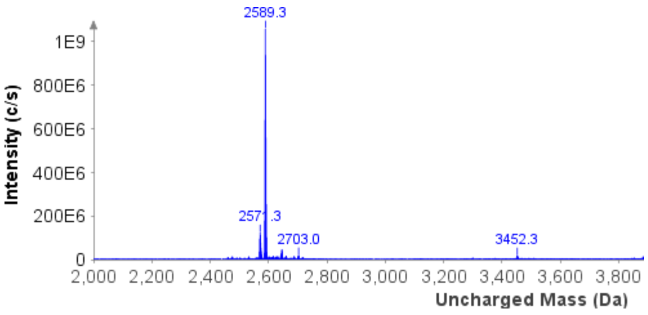

**Figure S7.** Deconvoluted MS spectrum of purified the Peptide D.

##### H-Ser-Ala-Lys-Ala-Gly-Ser-Gly-Tyr-Lys-Ser-Ala-CONH$_2$ (Peptide E)

Rink Amide AM: 167 mg (0.1 mmol scale).

For the synthesis of Peptide E, the following amino acids were used in sequence from the C- to the N-terminus: Fmoc-Ala-OH, Fmoc-Ser(tBu)-OH, Fmoc-Lys(Boc)-OH, Fmoc-Tyr(tBu)-OH, Fmoc-Gly-OH, Fmoc-Ser(tBu)-OH, Fmoc-Gly-OH, Fmoc-Ala-OH, Fmoc-Lys(Boc)-OH, Fmoc-Ala-OH, Fmoc-Ser(tBu)-OH.

Cleavage from resin and global deprotection, along with purification were performed as described in General procedures for peptide synthesis.

##### HPLC-MS

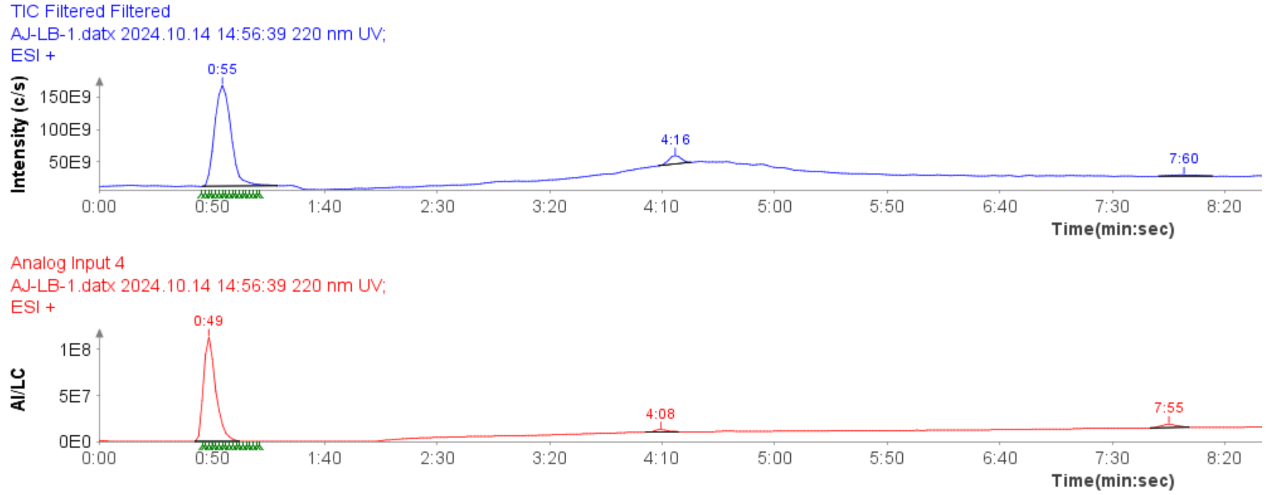

**Figure S8.** HPLC-MS of purified Peptide D, TIC (blue) and HPLC trace (red).

**MS-ESI** (*m*/*z*):

$[M+H]^{+}$ calculated for $C_{43}H_{73}N_{14}O_{15}^{+}$, 1025.5; found, 1025.4.

$[M+2H]^{2+}$ calculated for $C_{43}H_{74}N_{14}O_{15}^{2+}$, 513.3; found, 513.3.

$[M+3H]^{3+}$ calculated for $C_{43}H_{75}N_{14}O_{15}^{3+}$, 342.5; found, 342.6.

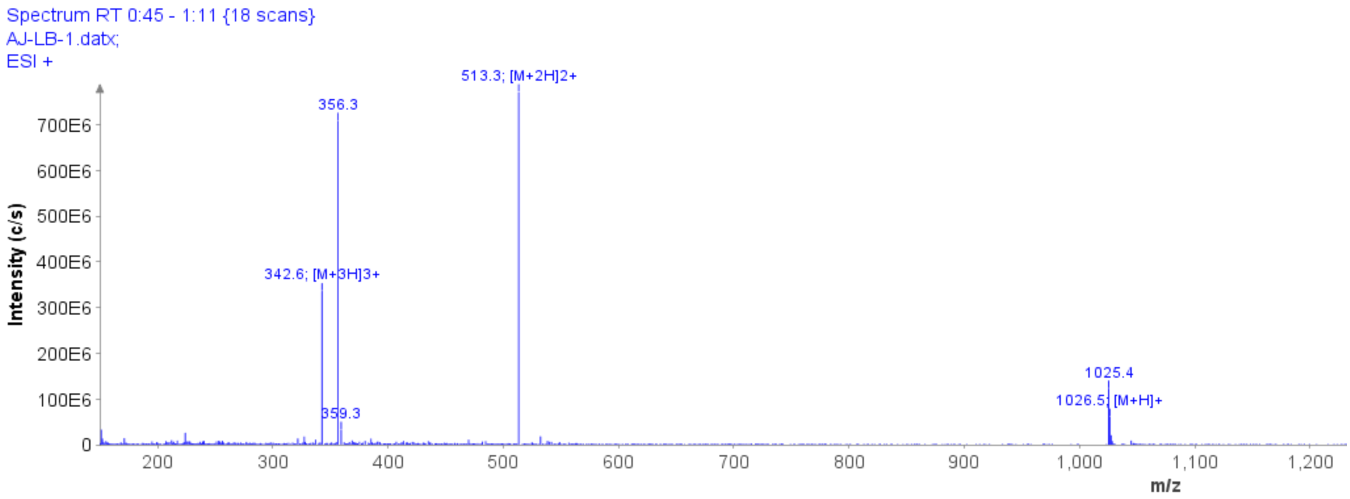

**Figure S9.** MS spectrum of purified the Peptide E (y-axis: total ion count (TIC), x-axis: *m*/*z*) of the above HPLC-MS analysis.

**Ac-Val-Ala-Gly-Lys-Gly-Lys-Gly-Gly-Ile-Phe-OH (Peptide F)**

Rink Amide AM: 167 mg (0.1 mmol scale).

For the synthesis of Peptide F, the following amino acids were used in sequence from the C- to the N-terminus: Fmoc-Phe-OH, Fmoc-Ile-OH, Fmoc-Gly, Fmoc-Gly-OH, Fmoc-Lys(Alloc)-OH, Fmoc-Gly-OH, Fmoc-Lys(Alloc)-OH, Fmoc-Gly-OH, Fmoc-Ala-OH, Fmoc-Val-OH.

For N-capping, Alloc deprotection and cleavage from resin and global deprotection, along with purification were performed as described in General procedures for peptide synthesis.

Synthesis of peptide **25**

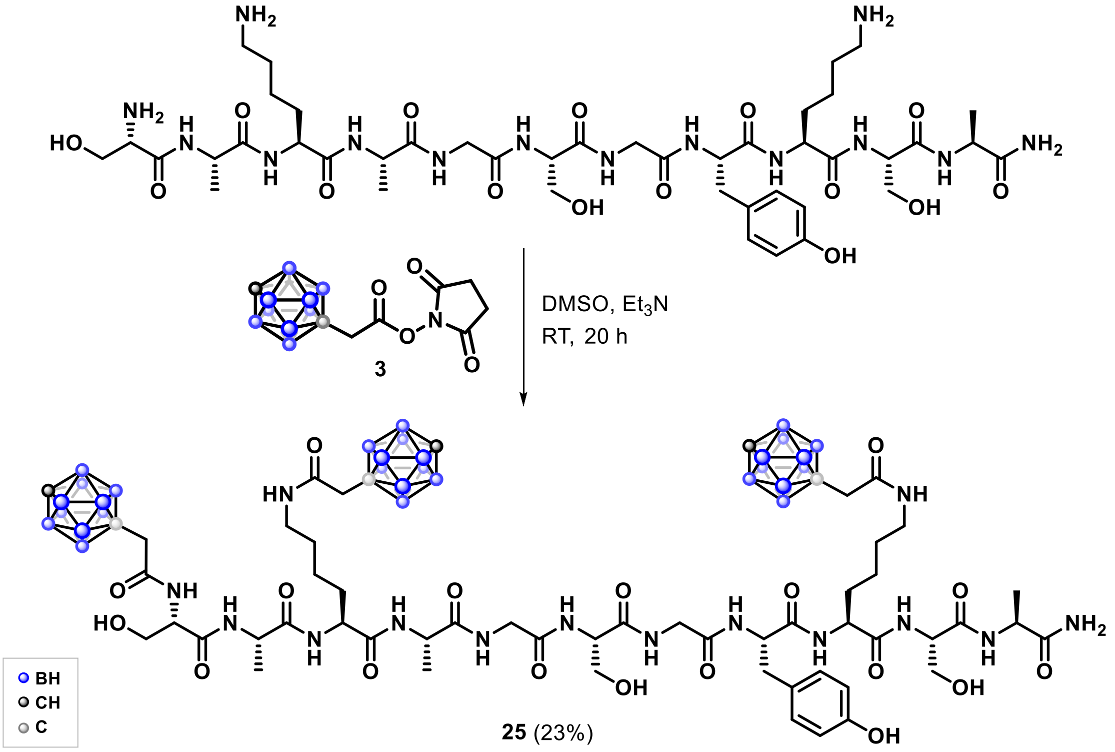

A solution of Peptide E (33 mg, 0.032 mmol) and compound **3** (29 mg, 0.097 mmol) in DMSO-$d_6$ (0.5 mL) was treated with $Et_3N$ (13.5 µL, 0.096 mmol). The reaction mixture was stirred at room temperature overnight. The crude product was purified by reverse-phase flash chromatography. Fractions were analyzed by LC–MS, and those containing the desired product were flash-frozen in liquid nitrogen and lyophilized to afford compound **25** as a white solid (11.4 mg, 23%).

**$^1$H NMR** and **$^{13}$C NMR** complex spectra.

**$^{11}$B NMR** (161 MHz, DMSO-$d_6$): δ -11.91, -12.76, -14.65, -15.50.

**HRMS** (*m/z*): $[M+2H]^{2+}$ calculated for $C_{55}H_{110}B_{30}N_{14}O_{18}^{2+}$, 790.0542; found, 790.0021.

Synthesis of peptide **26**

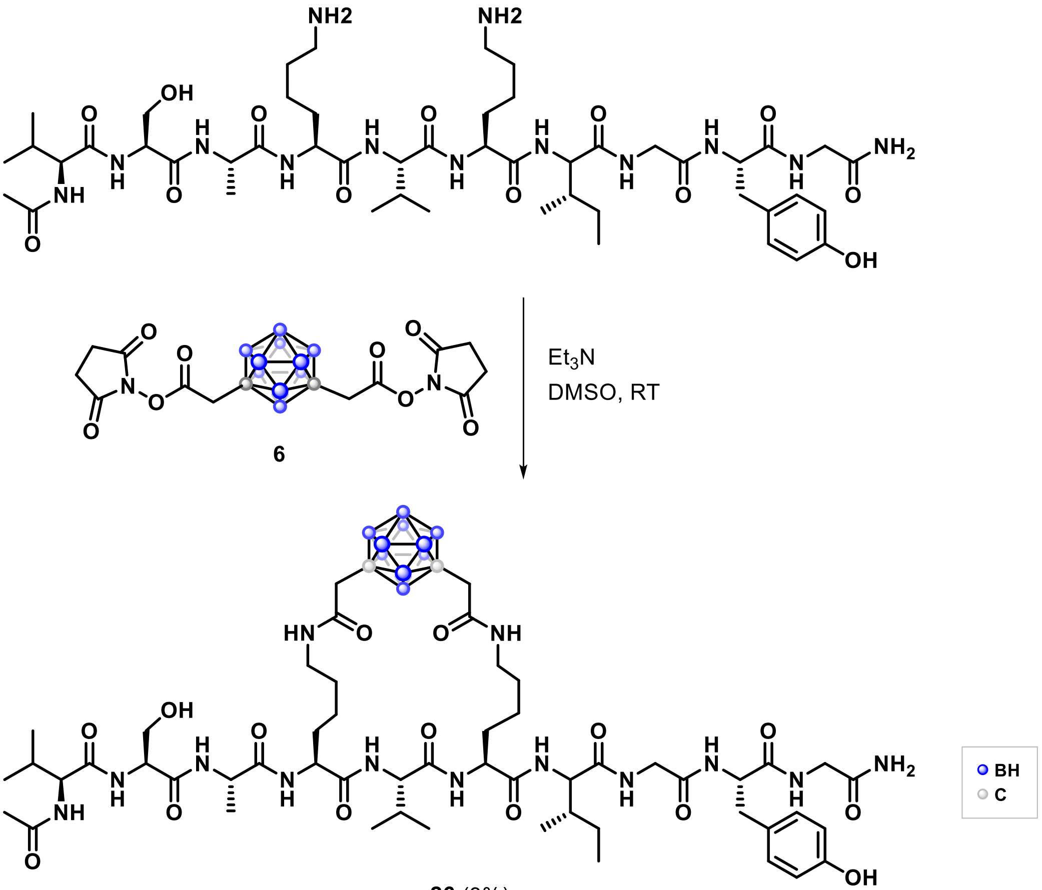

**26** (3%)

Peptide B (30 mg, 0.028 mmol) was dissolved in DMSO (2 mL). Compound **6** (25 mg, 0.055 mmol) and $Et_3N$ (10 µL, 0.072 mmol) were added sequentially at room temperature. The reaction mixture was stirred at room temperature for 30 min, after which HPLC-MS analysis indicated about 40% conversion to the target peptide. The reaction mixture was purified by a 25-gram reverse-phase C18 chromatography using an ACN/Milli-Q water gradient containing 0.1% TFA (ACN 20% → 90% over 30 min). Fractions were analyzed by HPLC–MS, and those containing the desired peptide were combined, flash-frozen in liquid nitrogen, and lyophilized to afford the target peptide as a white solid (1.0 mg, 3%).

**HPLC-MS** analyses with AZURA LC system (KNAUER) coupled to an Advion Expression CMS-L single quadrupole mass spectrometer. RP column Eurospher II 100-5 C18, 4 × 125 mm, flow rate 0.75 mL/min, Milli-Q water/acetonitrile gradient, 15 min.

Crude product: 15 µL of sample injected at a concentration of 1 mg/mL.

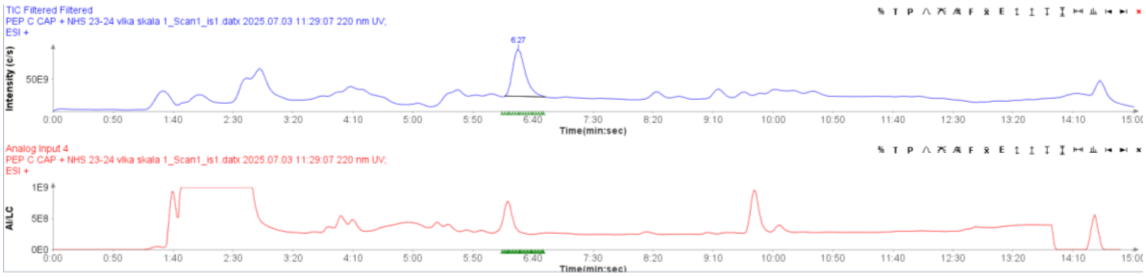

**Figure S10.** HPLC-MS of crude reaction mixture of peptide **26**, TIC (blue) and HPLC trace (red).

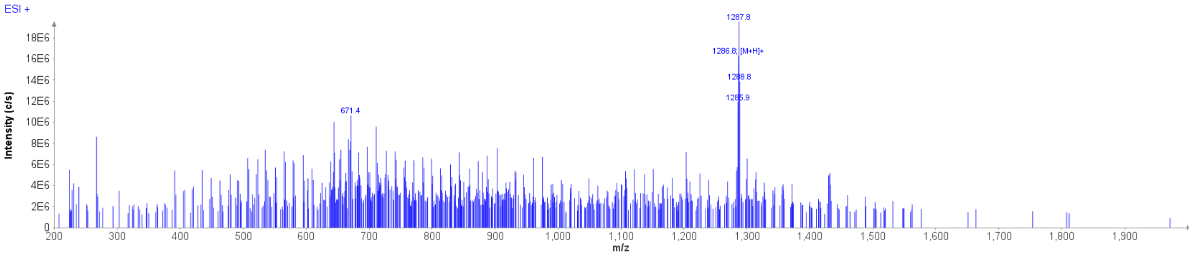

**Figure S11.** MS spectrum of crude reaction mixture of peptide **26** (y-axis: total ion count (TIC), x-axis: *m*/*z*) of the above HPLC-MS analysis.

**HRMS** (*m/z*):

$[M+H]^+$ calculated $C_{55}H_{96}B_{10}N_{13}O_{15}^+$, 1286.8147; found, 1286.8159.

$[M+2H]^{2+}$ calculated $C_{55}H_{97}B_{10}N_{13}O_{15}^{2+}$, 643.9110; found, 643.9138.

Synthesis of peptide **27**

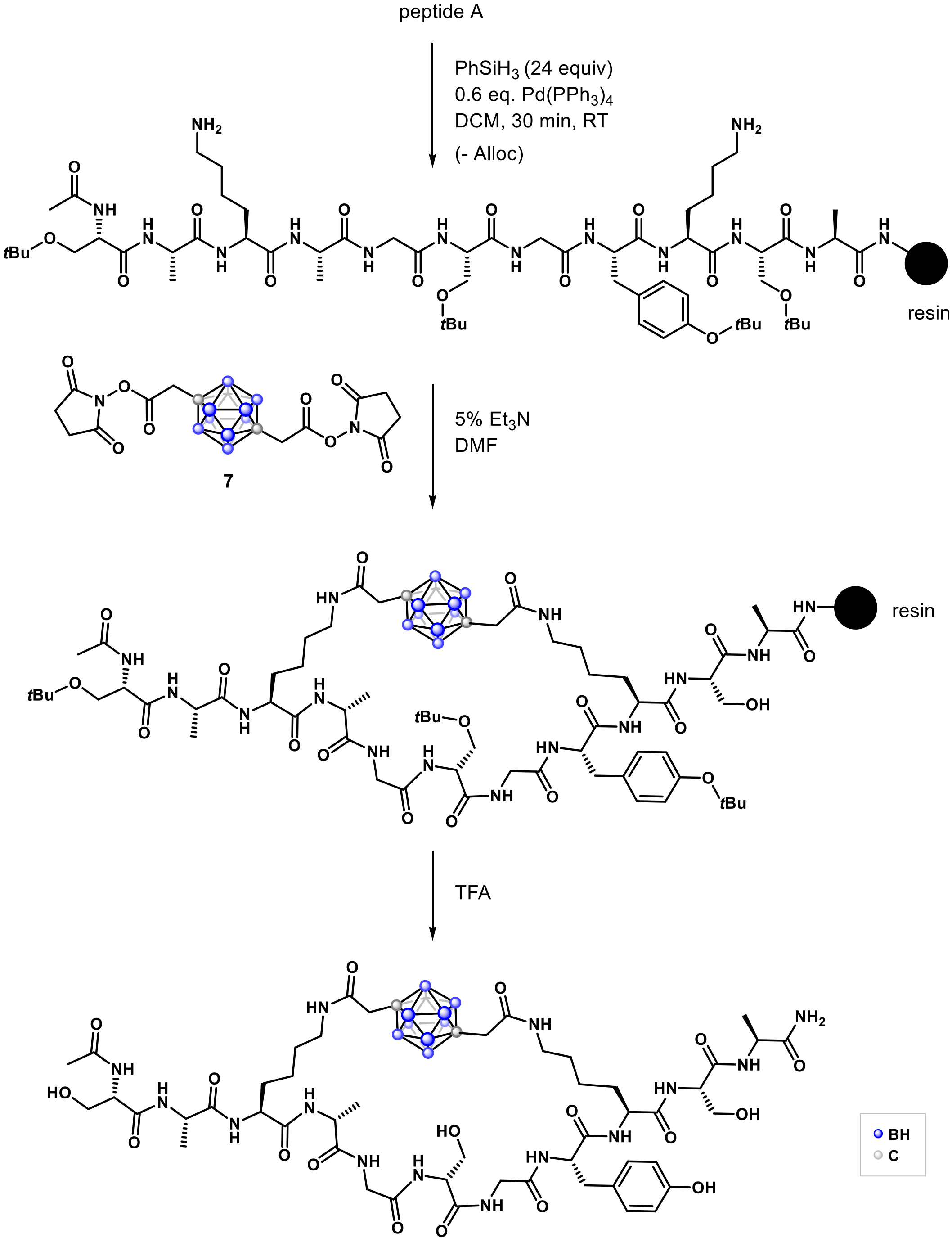

Resin-bound Peptide A (loading: 0.025 mmol), bearing deprotected lysine ε-amino groups and protected side chains on all other residues, was swollen in DMF (5 mL) for 15 min in 8 mL tube on a vaccum manifold. The solvent was removed under vacuum, and a solution of reagent **7** (23 mg, 0.05 mmol) in DMF (4.75 mL) was added to the resin, followed by $Et_3N$ (0.25 mL, 1.8 mmol, 5%). The reaction mixture was gently mixed and allowed to proceed overnight at room temperature in a closed tube. The solvent was then removed under vacuum, and the peptide was cleaved from the resin. Cleavage from resin and side-chain protecting groups were removed using a TFA-based cleavage cocktail (see General procedure). The crude peptide was purified by reverse-phase flash chromatography (C18, gradient: MeCN/Milli-Q water containing 0.1% TFA; MeCN 30% → 90% over

30 min), the fractions were analyzed with HPLC-MS, the fractions containing target peptide were combined, flashed freezed in liquid nitrogen and lyophilized to afford the desired **27** as a white solid (5 mg, 16%).

**HPLC-MS** analyses with AZURA LC system (KNAUER) coupled to an Advion Expression CMS-L single quadrupole mass spectrometer. RP column Eurospher II 100-5 C18, 4 × 125 mm, flow rate 0.75 mL/min, water/acetonitrile gradient, 15 min.

Crude product: 15 µL of sample injected at a concentration of about 1 mg/mL.

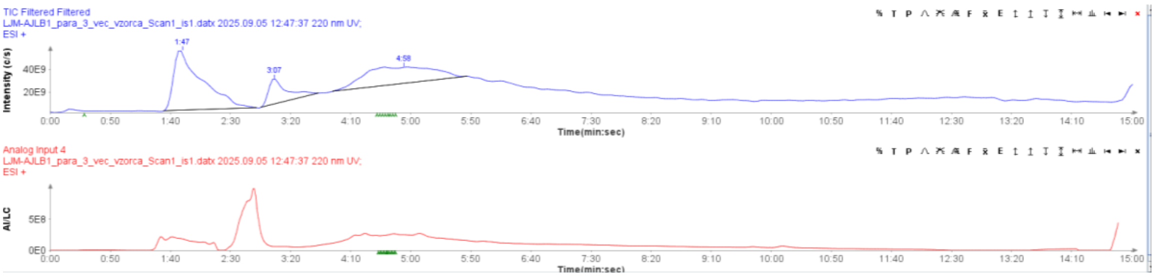

**Figure S12.** HPLC-MS of crude reaction mixture of peptide **27**, TIC (blue) and HPLC trace (red).

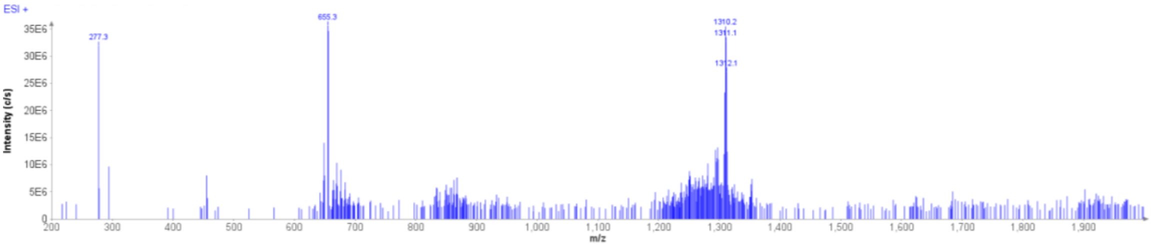

**Figure S13.** MS spectrum of crude reaction mixture of peptide **27** (y-axis: total ion count (TIC), x-axis: *m*/*z*) of the above HPLC-MS analysis.

Purified product:

**HRMS** (*m/z*): $[M+NH_4]^+$ calculated for $C_{51}H_{90}B_{10}N_{15}O_{18}^+$, 1309.7550; found, 1309.7459.

Preliminarily attempts of intramolecular coupling of Ac-VAGKGKGGIF-CONH–resin-bound peptide with *ortho*-carborane bis-NHS ester **4**

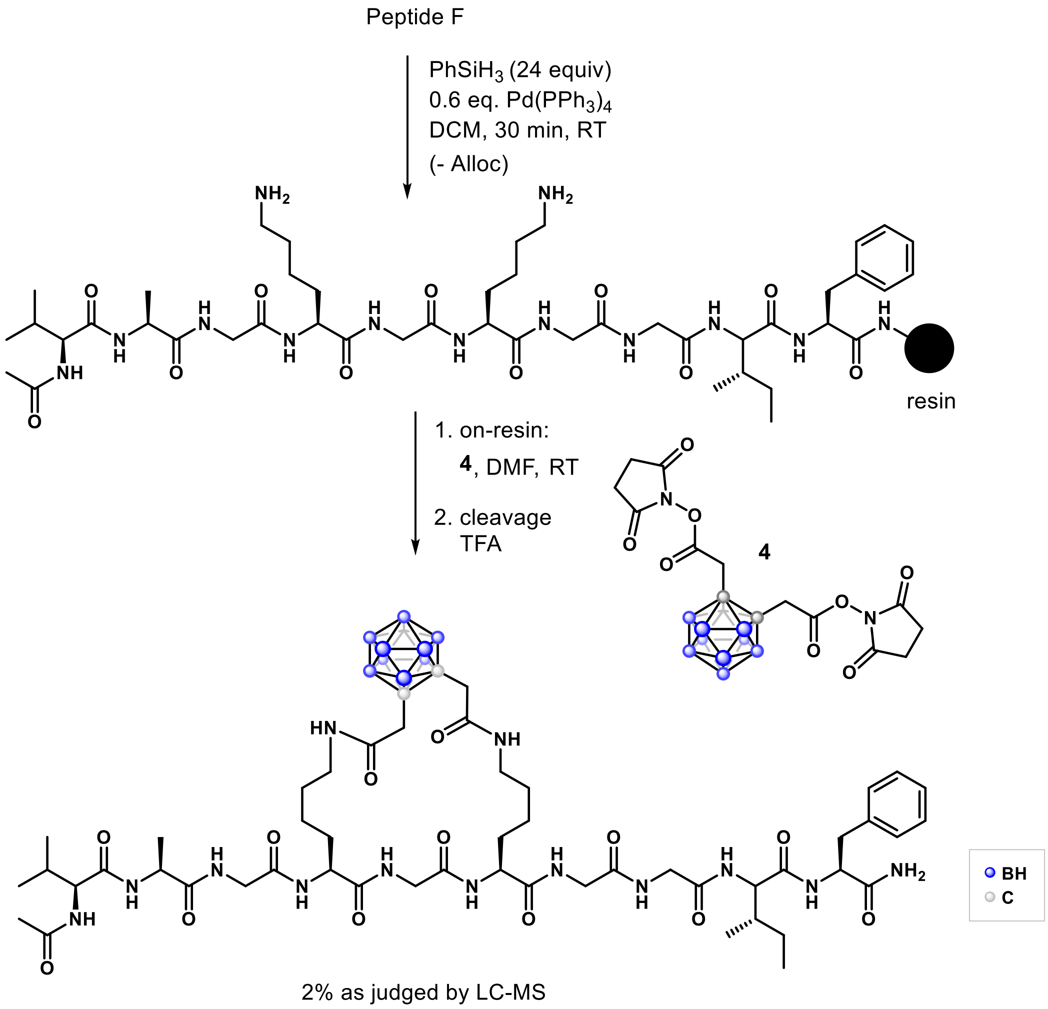

Resin-bound N-acylated Peptide F (0.025 mmol), bearing deprotected lysine ε-amino groups, was swollen in DMF (5 mL) for 15 min. The solvent was removed under vacuum, and a solution of reagent **4** (11 mg, 0.025 mmol, 2 equiv) in DMF (4.75 mL) was added to the resin, followed by $Et_3N$ (0.25 mL, 1.8 mmol, 5%). The reaction mixture was gently agitated and allowed to proceed overnight at room temperature. After completion, the solvent was removed under vacuum, and the peptide was cleaved from the solid support, along with side-chain protecting groups removal, using a TFA-based cleavage cocktail. The crude peptide was analyzed by LC-MS.

**MS-ESI** (m/z):

$[M+H]^+$ calculated for $C_{51}H_{89}B_{10}N_{13}O_{13}^+$, 1199.8; found, 1200.2.

$[M+2H]^{2+}$ calculated for $C_{51}H_{90}B_{10}N_{13}O_{13}^{2+}$, 599.9; found, 600.8.

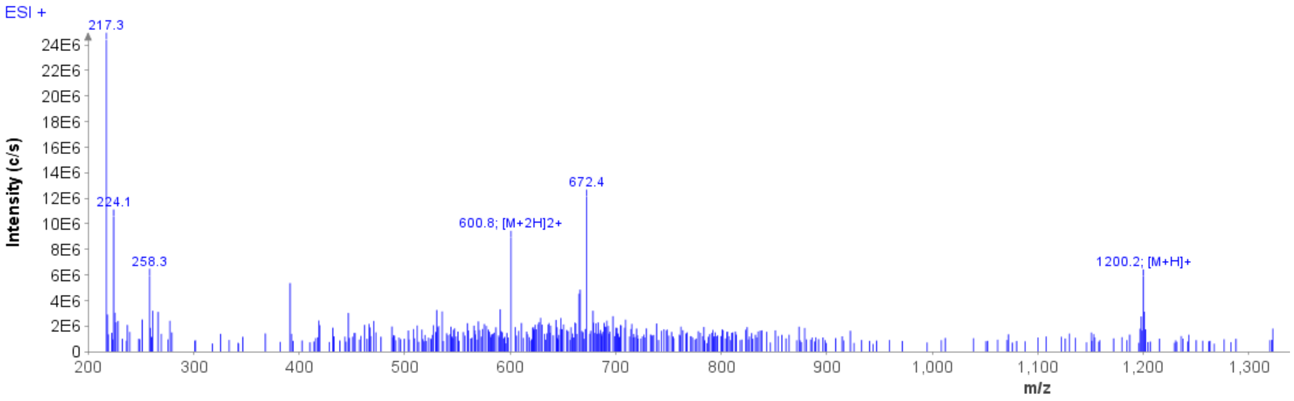

**Figure S14.** MS spectrum of crude reaction mixture of peptide **27** (y-axis: total ion count (TIC), x-axis: *m*/*z*) of the above HPLC-MS analysis.

Synthesis of peptide **28**

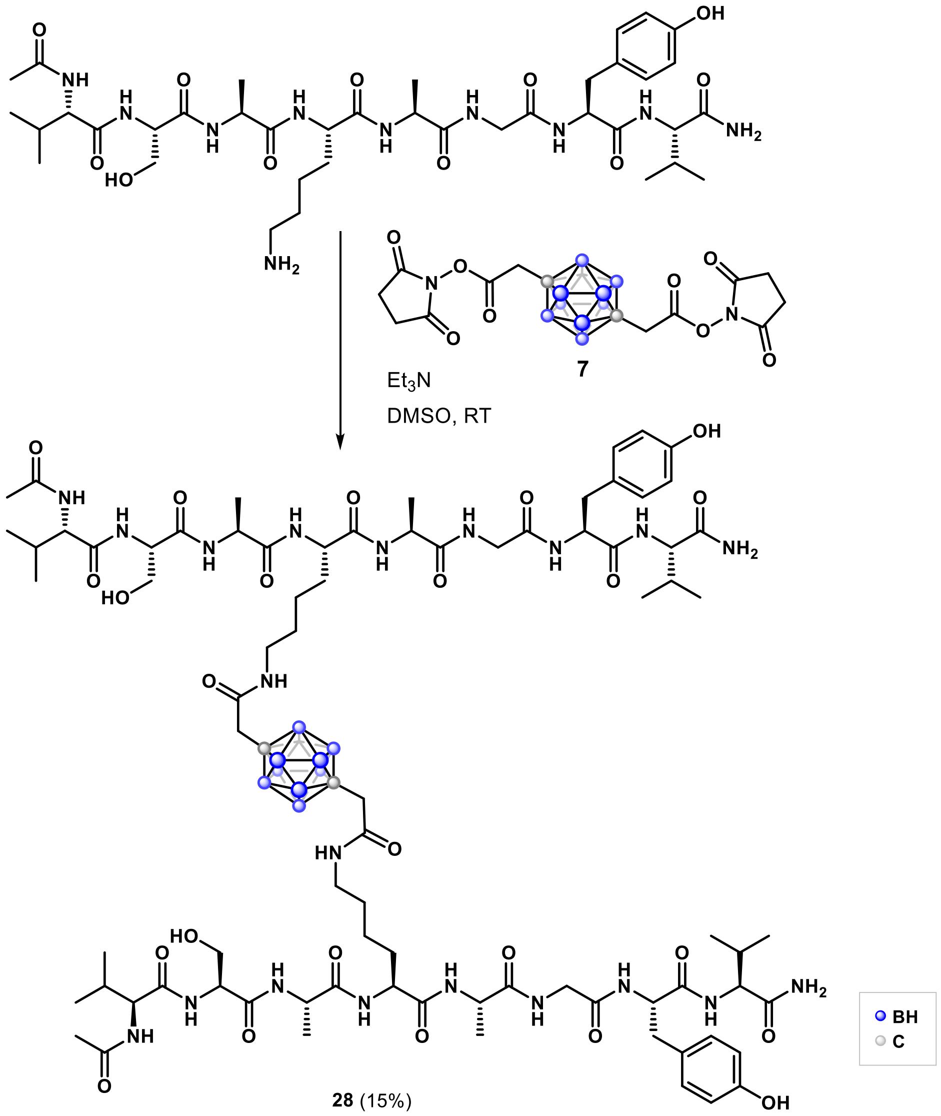

Peptide C (60 mg, 0.072 mmol) was dissolved in DMSO (0.5 mL). Compound **7** (8.2 mg, 0.018 mmol) and $Et_3N$ (20 µL, 0.144 mmol) were added sequentially at room temperature, and the reaction mixture was gently agitated and allowed to proceed overnight at room temperature . The reaction mixture was then purified by reverse-phase flash chromatography (C18, MeCN/Milli-Q water containing 0.1% TFA; gradient: 30 → 90% MeCN over 30 min). Fractions were analyzed by HPLC-MS, and those containing the target peptide were combined, flash-frozen in liquid nitrogen, and lyophilized to afford the peptide **28** as a white solid (5 mg, 15%).

**$^{1}$H NMR** (500 MHz, DMSO-*d*$_6$): Complex spectrum.

**$^{13}$C NMR** (126 MHz, DMSO-*d*$_6$): δ 173.2, 172.8, 172.6, 171.8, 171.7, 171.3, 170.4, 170.0, 168.9, 166.4, 156.2, 130.6, 128.2, 115.3, 62.1, 58.4, 58.0, 55.4, 54.6, 49.0, 48.8, 42.3, 39.0, 37.1, 31.7, 30.9, 29.0, 23.2, 23.0, 19.7, 19.7, 18.6, 18.5, 18.5, 18.3, 9.1. Five carbon resonances are missing or overlapping.

**$^{11}$B NMR** (161 MHz, DMSO-*d*$_6$): δ -12.0, -12.8.

**HRMS** (*m/z*):

$[M+H]^+$ calculated for $C_{82}H_{137}B_{10}N_{20}O_{24}^+$, 1894.1112; found, 1894.1142.

$[M+2H]^{2+}$ calculated for $C_{82}H_{138}B_{10}N_{20}O_{24}^{2+}$, 947.5593; found, 947.5619.

**HPLC:** Atlantis C18 column (5 µm, 4.6 × 250 mm); isocratic ACN/Milli-Q water (50:50, 0.1% TFA) for 20 min, flow 0.75 mL/min. Compound purity was assessed by UV detection at 220 nm and expressed as the percentage of the compound peak area relative to the total integrated area.

Peptide **28** in pure form, concentration: 1 mg/mL, injected 20 µL.

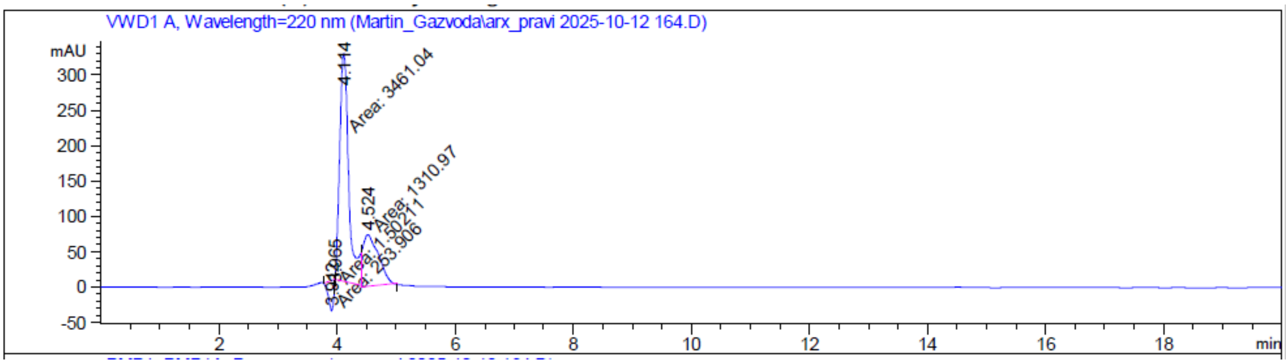

Signal 1: VWD1 A, Wavelength=220 nm

| Peak # | RetTime [min] | Type | Width [min] | Area [mAU*s] | Height [mAU] | Area % |
|---|---|---|---|---|---|---|
| 1 | 3.912 | MM N | 0.0999 | 253.90579 | 42.35684 | 5.0504 |
| 2 | 3.965 | MP N | 2.41e-3 | 1.50211 | 10.40051 | 0.0299 |
| 3 | 4.114 | PM | 0.1794 | 3461.04297 | 321.47989 | 68.8433 |
| 4 | 4.524 | MM | 0.2985 | 1310.97058 | 73.19446 | 26.0764 |
| Totals : | | | | 5027.42145 | 447.43169 | |

**Figure S15**. HPLC trace of the purified peptide **28** (y-axis: absorbance, wavelength = 220 nm; x-axis: time in minutes).

Synthesis of peptide **29**

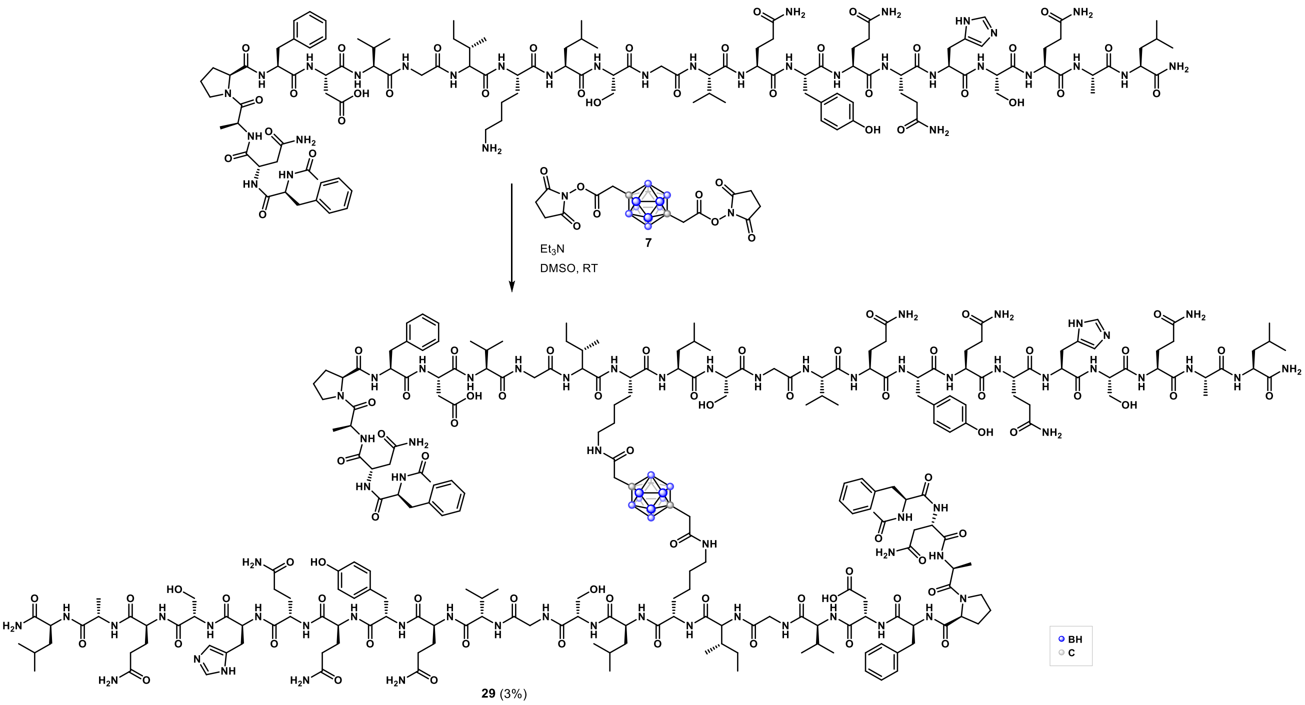

Peptid **D** (170 mg, 0.066 mmol) was dissolved in DMSO (1.5 mL). To this solution was added compound **7** (15 mg, 0.033 mmol) and $Et_3N$ (18.4 µL, 0.132 mmol). The reaction mixture was gently mixed and allowed to proceed overnight at room temperature after which it was purified by reverse-phase flash chromatography (C18, MeCN/Milli-Q water containing 0.1% TFA; gradient: 30 → 90% MeCN over 30 min). Fractions were analyzed by HPLC-MS, and those containing the target peptide were combined, flash-frozen in liquid nitrogen, and lyophilized to afford the peptide **29** as a white solid (6 mg, 3%).

**HPLC-MS** analyses with AZURA LC system (KNAUER) coupled to an Advion Expression CMS-L single quadrupole mass spectrometer. RP column Eurospher II 100-5 C18, 4 × 125 mm, flow rate 0.75 mL/min, water/acetonitrile gradient, 15 min.

Crude product: 15 µL of sample injected at a concentration of 1 mg/mL.

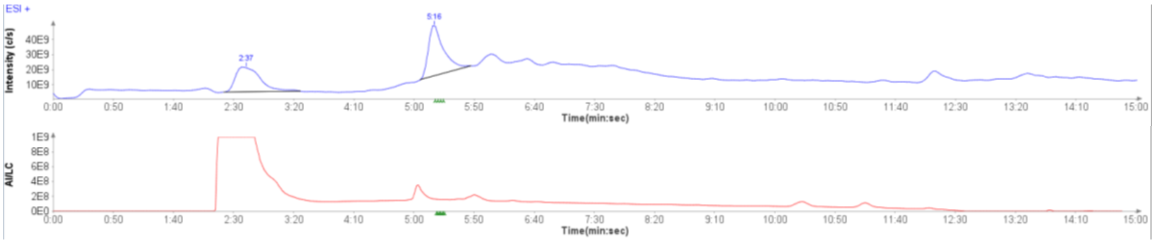

**Figure S16.** HPLC-MS of crude reaction mixture of peptide **29**, TIC (blue) and HPLC trace (red).

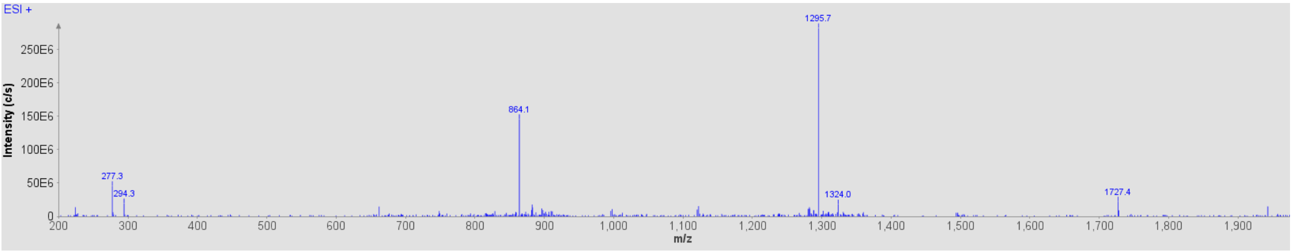

**Figure S17.** MS spectrum of crude reaction mixture of peptide **29** (y-axis: total ion count (TIC), x-axis: *m*/*z*) of the above HPLC-MS analysis.

Purified product after reverse-phase chromatography: 15 µL of sample injected at a concentration of 1 mg/mL:

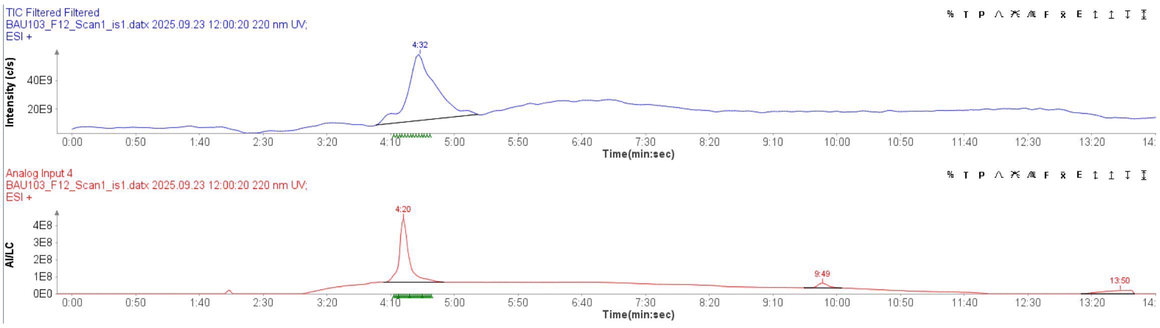

**Figure S18.** HPLC-MS of purified peptide **29**, TIC (blue) and HPLC trace (red).

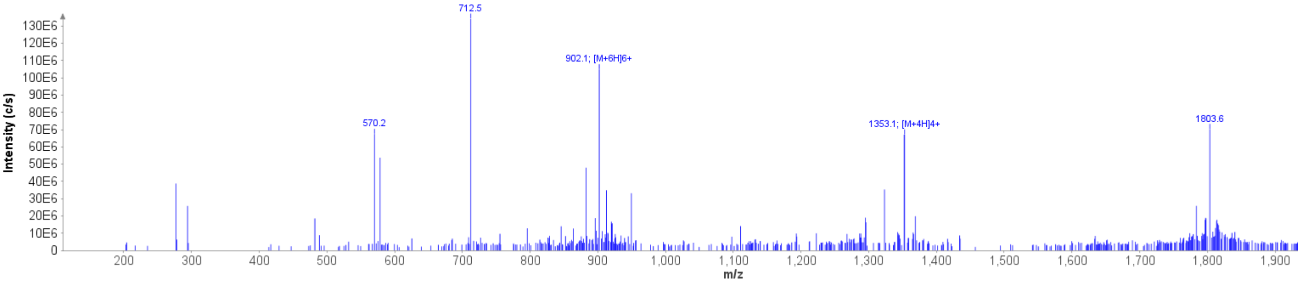

**Figure S19.** MS spectrum of purified peptide **29** (y-axis: total ion count (TIC), x-axis: *m*/*z*) of the above HPLC-MS analysis.

$[M+H]^+$ calculated for $C_{242}H_{369}B_{10}N_{64}O_{70}^+$, 5403.1; found (deconvoluted) 5407.3

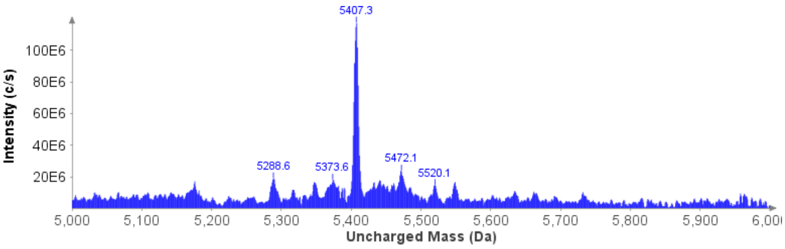

**Figure S20.** Deconvoluted MS spectrum of purified the peptide **29**.

Comment: Deamidation of Asn/Gln residues is a well-known side reaction that may occured and account for the difference between the calculated and experimentally observed masses (+4 Da) in case of **29**.

**HRMS** (*m/z*):

$[M+4H]^{4+}$ calculated $C_{242}H_{372}B_{10}N_{64}O_{70}^{4+}$, 1351.9570; found, 1352.1824.

$[M+6H]^{6+}$ calculated $C_{242}H_{374}B_{10}N_{64}O_{70}^{6+}$, 901.6404; found, 901.7914.

**HPLC:** Atlantis C18 column (5 µm, 4.6 × 250 mm); isocratic ACN/$H_2O$ (50:50, 0.1% TFA) for 20 min, flow 0.75 mL/min. Peptide concentration: 1 mg/mL, injected 20 µL. Compound purity was assessed by UV detection at 220 nm and expressed as the percentage of the compound peak area relative to the total integrated area.

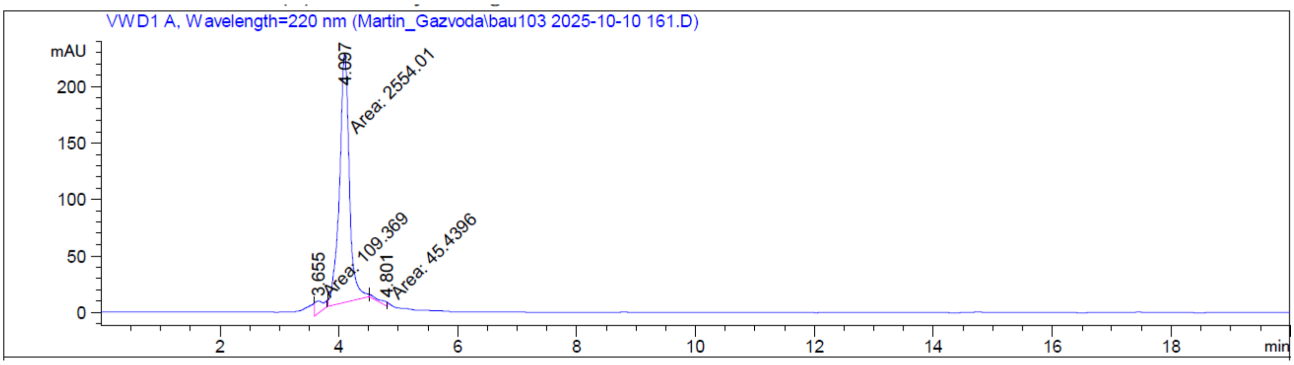

Signal 1: VWD1 A, Wavelength=220 nm

| Peak # | RetTime [min] | Type | Width [min] | Area [mAU*s] | Height [mAU] | Area % |
|---|---|---|---|---|---|---|
| 1 | 3.655 | MM | 0.1703 | 109.36893 | 10.70110 | 4.0375 |
| 2 | 4.097 | MM | 0.1934 | 2554.01050 | 220.08865 | 94.2850 |
| 3 | 4.801 | MM | 0.2138 | 45.43961 | 3.54193 | 1.6775 |
| Totals : | | | | 2708.81904 | 234.33168 | |

**Figure S21**. HPLC trace of the purified peptide **29** (y-axis: absorbance, wavelength = 220 nm; x-axis: time in minutes).

#### Carborane-peptide polymer assembly using carborane bi-NHS esters (Fig. 4)

##### Preliminary testing of polymerization reactions

The bifunctional NHS reagent **7** (MW 454 g/mol) and the model peptide H-SKSSSSSSKS-$CONH_2$ (MW 970 g/mol) were first evaluated for their potential to form peptide-carborane polymers.

To investigate the influence of reaction stoichiometry on polymer length, three reaction conditions were tested, varying the ratio of peptide to reagent **7**. The reactions were performed in DMSO-$d_6$ at room temperature, and the progress was monitored by recording the $^1$H NMR spectrum one hour after initiation. The resulting polymer length was analyzed by SDS-PAGE.

Each reaction was carried out using 12 mg of reagent **7** (0.026 mmol, c = 50 mM) in 0.5 mL of DMSO-$d_6$, with $Et_3N$ (7 µL, 0.05 mmol, c = 100 mM) as base. Different equivalents of the peptide were used - 50 mM (1 equiv.), 25 mM (0.5 equiv.), and 10 mM (0.2 equiv.) - to assess the effect of the peptide-to-reagent ratio on polymer length.

**Table S1**: Preliminary attempts of polymer formation:

| entry | **7** : peptide (ratio) | **7** | peptide | $Et_3N$ | solvent | temperature |
|---|---|---|---|---|---|---|
| 1 | 1:1 | 50 mM | 50 mM | 100 mM | DMSO | RT |
| 2 | 2:1 | 50 mM | 25 mM | 100 mM | DMSO | RT |
| 3 | 4:1 | 50 mM | 10 mM | 100 mM | DMSO | RT |

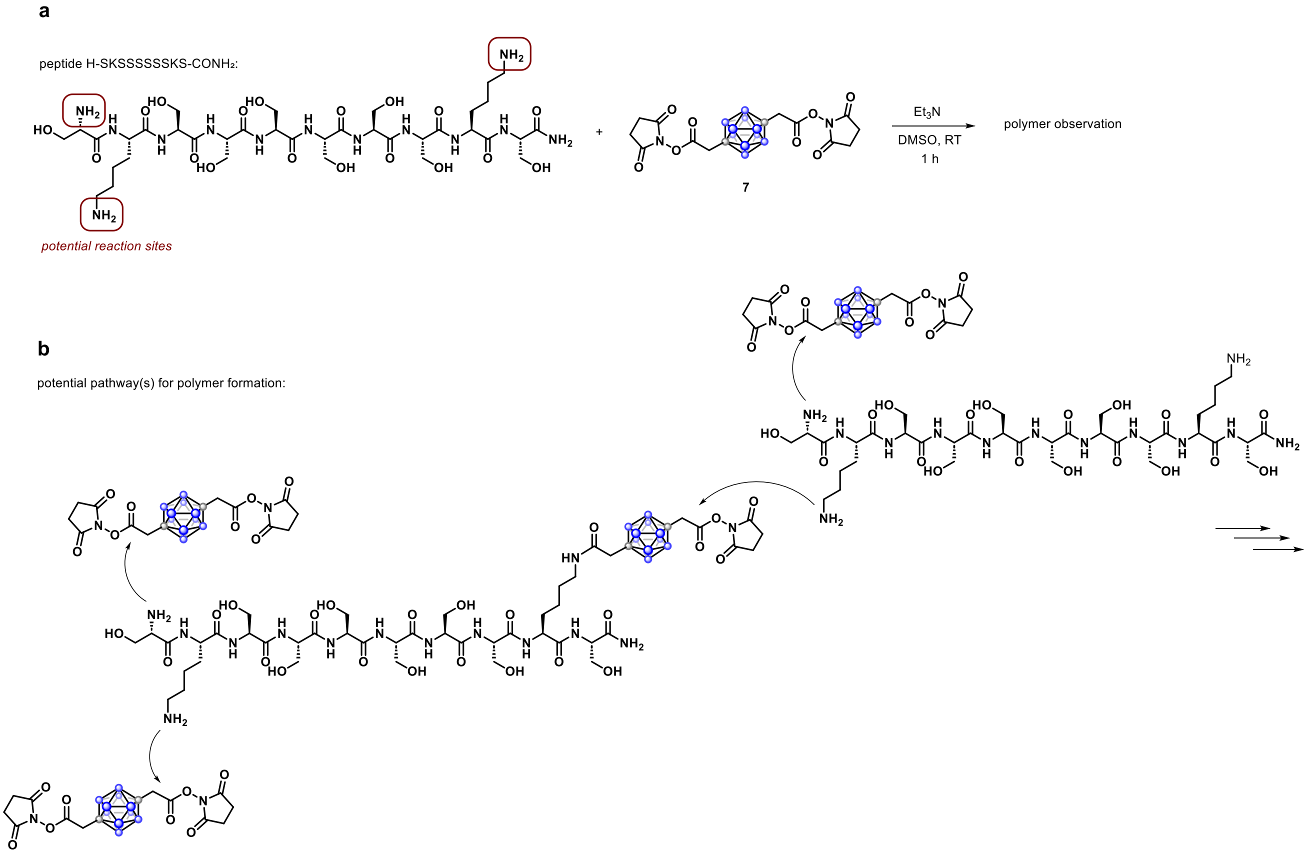

**Figure S22**: a, Schematic representation of the model reaction for polymer assembly using reagent 7 and peptide H-SKSSSSSSKS-$CONH_2$ in DMSO at room temperature. **b**, The peptide H-SKSSSSSSKS-$CONH_2$ contains two lysine residues and an unprotected N-terminus, providing three potential reactive sites for coupling with reagent 7, enabling polymer branching.

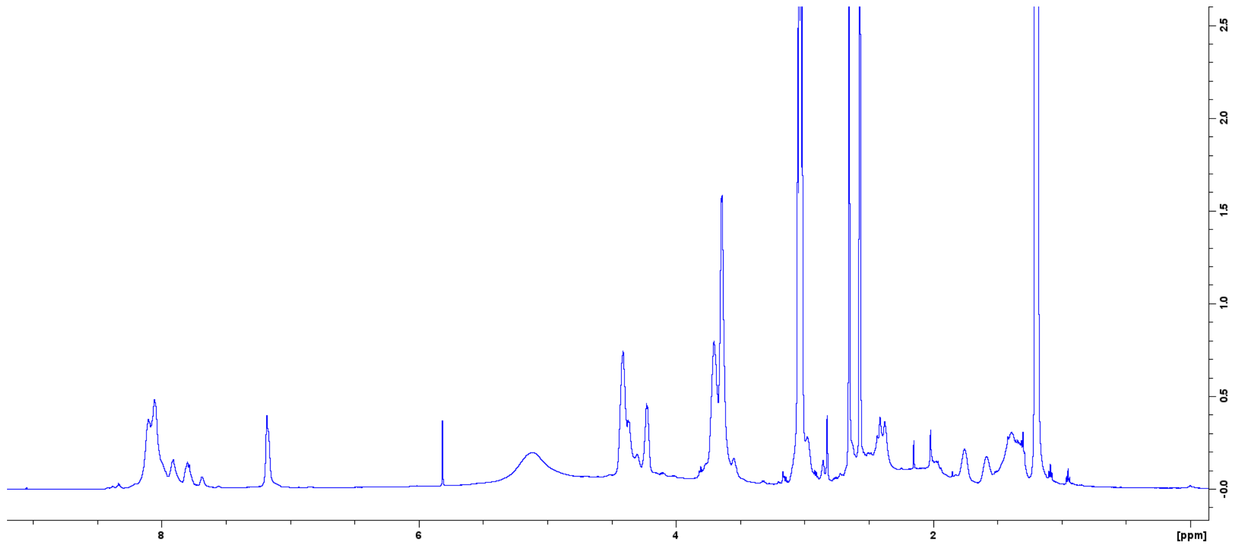

**Figure S23**: $^{1}$H NMR spectrum of the reaction described in Table S1, entry 1, after 1 hour. The solution remained transparent, no precipitation was observed. $^{1}$H NMR spectrum exhibited broad, non-discrete resonances, indicative of polymer formation. 1H NMR spectra of the reactions corresponding to entries 2 and 3 were similar to the spectrum above.

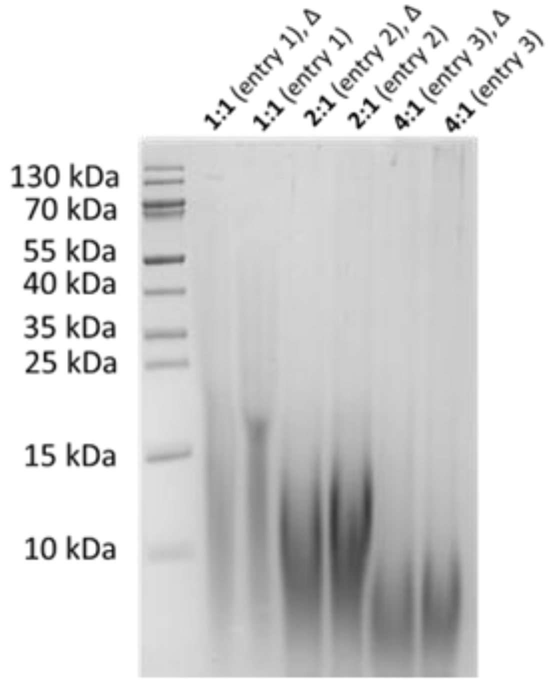

**Figure S24**. SDS-PAGE analysis of preliminary polymerization reactions performed in DMSO using a 16% SDS–Tricine gel. Various reaction conditions were tested by altering the molar ratio of the reagent **7** to model peptide (1:1, 2:1, and 4:1). Each sample was mixed with the appropriate volume of SDS loading buffer containing a reducing agent. Samples marked with Δ were heated at 95 °C for 10 minutes prior to loading. A volume of 5 µL of each sample was loaded per well

**Further screening of reaction parameters**

The results of preliminary polymerization reactions indicated that an increased excess of peptide favored the formation of longer polymer chains. We therefore sought to further evaluate this observation by increasing the peptide ratio in subsequent optimization screening reactions.

Further polymerization screening assays were performed using peptides H-SAKAGSGYKSA-$CONH_2$ and H-SKSSSSSSKS-$CONH_2$. Similary as above, to initiate polymerization, appropriate amounts of the bifunctional reagent **7** and base ($Et_3N$) were added to each peptide solution. Stock solutions of the peptides and NHS ester **7** were prepared at 0.2 M in DMSO, and $Et_3N$ was prepared as a 1.3 M stock solution in DMSO.

Reaction mixtures (20 µL total volume) were assembled in 1.5 mL microcentrifuge tubes. Unless stated otherwise, reagents were added in the following order: solvent, peptide, NHS ester **7** and finally $Et_3N$. The final concentrations of all reaction components are provided in Supplementary Tables 1 and 2. Reactions were incubated overnight at room temperature or under heating conditions: 50 °C or 90 °C for H-SKSSSSSSKS-$CONH_2$, and 65 °C or 85 °C for H-SAKAGSGYKSA-$CONH_2$ - using a sand bath.

**Table S2**. Reaction conditions while testing polymerization conditions for peptide H-SKSSSSSSKS-$CONH_2$

| **7** : peptide (ratio) | **7** | peptide | $Et_3N$ | solvent | temperature |
|---|---|---|---|---|---|
| 1:2 | 20 mM | 40 mM | 65 mM | DMSO | RT |
| 1:2 | 20 mM | 40 mM | 130 mM | DMSO | RT |
| 1:4 | 20 mM | 80 mM | 130 mM | DMSO | RT |
| 1:8 | 20 mM | 160 mM | 260 mM | DMSO | RT |
| 1:4 | 40 mM | 160 mM | 260 mM | DMSO | RT |
| 1:2 | 20 mM | 40 mM | 130 mM | DMSO | 50 °C |
| 1:2 | 20 mM | 40 mM | 130 mM | DMSO | 90 °C |
| 1:2 | 20 mM | 40 mM | 130 mM | DMF | RT |
| 1:2 | 20 mM | 40 mM | 130 mM | MeCN | RT |
| 1:1 | 40 mM | 40 mM | 130 mM | DMSO | RT |
| 1:1 | 40 mM | 40 mM | 130 mM | DMSO | 50 °C |
| 1:1 | 40 mM | 40 mM | 130 mM | DMSO | 90 °C |
| 1:2 (NHS induction) | 20 mM | 40 mM | 130 mM | DMSO | RT |
| 1:2 ($Et_3N$ induction) | 20 mM | 40 mM | 130 mM | DMSO | RT |

**Table S3**. Reaction conditions while testing polymerization conditions for peptide H–SAKAGSGYKSA–$CONH_2$

| **7** : peptid (ratio) | **7** | peptide | $Et_3N$ | solvent | temperature |
|---|---|---|---|---|---|
| 2:1 | 80 mM | 40 mM | 260 mM | DMSO | RT |
| 2:1 | 80 mM | 40 mM | 260 mM | DMSO | 65 °C |
| 2:1 | 80 mM | 40 mM | 260 mM | DMSO | 85 °C |
| 1:1 | 40 mM | 40 mM | 260 mM | DMSO | RT |
| 1:1 | 40 mM | 40 mM | 260 mM | DMSO | 65 °C |
| 1:1 | 40 mM | 40 mM | 260 mM | DMSO | 85 °C |

| | | | | | |
|---|---|---|---|---|---|
| 1:2 | 20 mM | 40 mM | 130 mM | DMSO | RT |
| 1:2 | 20 mM | 40 mM | 130 mM | DMSO | 65 °C |
| 1:2 | 20 mM | 40 mM | 130 mM | DMSO | 85 °C |
| 1:4 | 20 mM | 80 mM | 130 mM | DMSO | RT |
| 1:4 | 20 mM | 80 mM | 130 mM | DMSO | 65 °C |
| 1:4 | 20 mM | 80 mM | 130 mM | DMSO | 85 °C |

After incubation, an appropriate volume of SDS loading buffer was added directly to the reaction mixtures, which were then heated for 5 min at 95 °C. The samples were subsequently analyzed by SDS–PAGE using 16% Tricine gels to evaluate polymer formation.

Preliminary experiments using the longer peptide (H–SKAQGWNYSGYAWASQNWQSGKA–$CONH_2$) produced reaction products that were insoluble even under strongly denaturing conditions (2% SDS, 6 M urea, and 100 mM DTT) (Fig. S25). As a result, these products did not migrate in either SDS–PAGE or SDS–AGE (0.8% agarose) gels, indicating the formation of highly aggregated or cross-linked polymeric species that remained insoluble under the tested conditions.

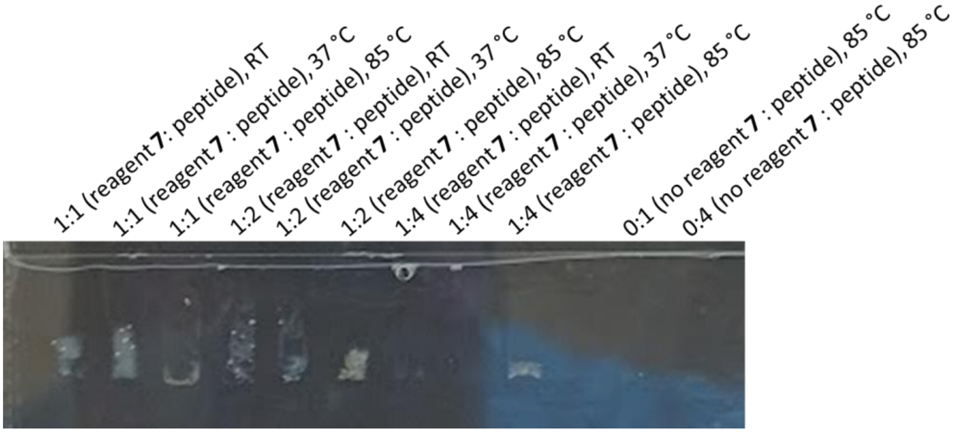

**Figure S25**. Evaluation of polymerization attempts with longer peptide on SDS-PAGE 16% Tricine gel.

**Size-Exclusion Chromatography (SEC) (Fig. 4c)**

For large-scale preparation, the reaction between peptide H–SAKAGSGYKSA–$CONH_2$ and H–SKSSSSSSKS–$CONH_2$ was carried out at a 4:1 molar ratio in a total volume of 460 µL DMSO under the same conditions described above (Supporting Table 1 and 2). The resulting mixture was analyzed by size-exclusion chromatography (SEC) using a Superdex® 200 Increase 10/300 GL column (Cytiva). Prior to injection, the reaction mixture was diluted 1:100 in SEC buffer and centrifuged at 10,000 × g for 10 min at room temperature to remove insoluble material. The clarified supernatant was then loaded onto the column. The SEC buffer consisted of 100 mM sodium phosphate and 300 mM NaCl at pH 7.2, and chromatographic profiles were monitored by UV absorbance.

For the peptide H–SAKAGSGYKSA–$CONH_2$, which was prepared on a larger scale, purification was also attempted using reverse-phase liquid chromatography (RP-LC). Automated flash chromatography was performed on an Interchim PuriFlash XS520Plus system equipped with reverse-phase C18 column (PF-15C18HP-F0004, Interchim) and employing MeCN/$H_2O$ gradients containing 0.1% TFA. Fractions were collected based on UV absorbance at 220 nm into 25 mL test tubes, then lyophilized and reconstituted in water to a final concentration of 10 mg/mL. An appropriate volume of SDS loading buffer was subsequently added, and the samples were heated at 95 °C for 10 minutes. The resulting samples were analyzed by SDS–PAGE using 16% Tricine gels.

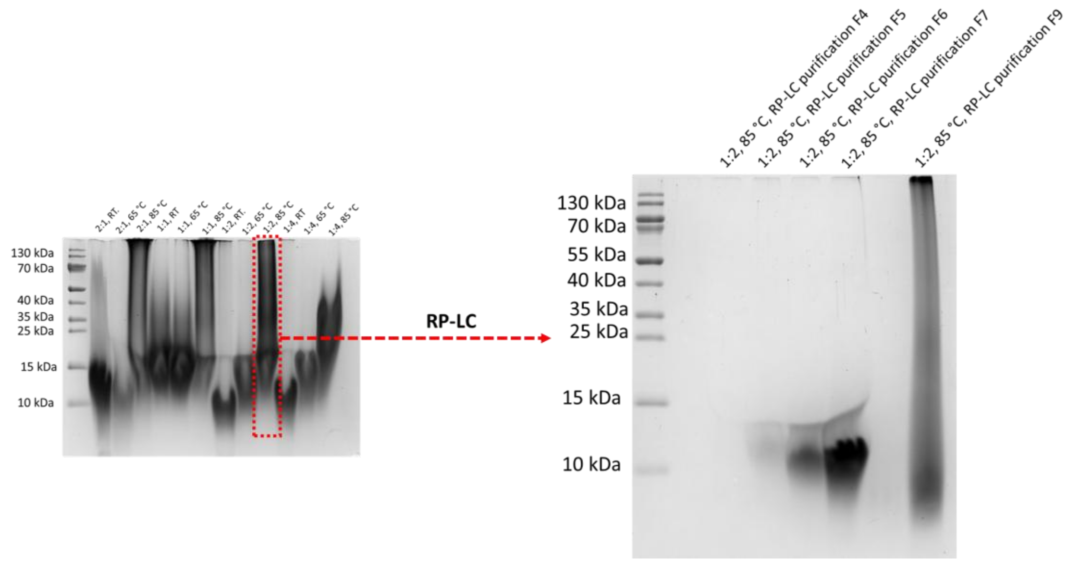

**Figure S26**. Analysis of samples after RP-LC purification.

#### Antibody conjugation (Fig. 5)

##### General procedure for antibody conjugation with carborane NHS esters (Fig. 5a)

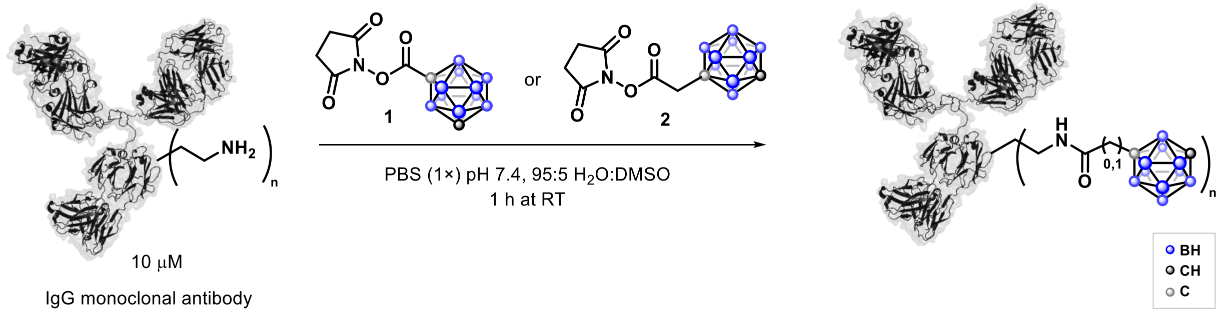

In a transparent polypropylene 0.6 mL Eppendorf tube, an antibody solution (10.5 μM in PBS, pH 7.4, 1×; 95 μL) was prepared at room temperature. To this solution, 5 μL of NHS ester **1** or **2** in DMSO (2, 6, 10, 15, 20, 30, or 40 mM) was added, corresponding to 10, 30, 50, 75, 100, 150, and 200 equivalents of NHS ester relative to the antibody. The mixture was mixed thoroughly by pipetting up and down 15 times. The final reaction conditions were 10 μM antibody, 100–2000 μM NHS ester, and 5% DMSO.

The reaction was allowed to proceed for 1 h at room temperature. Buffer exchange was then performed as follows: the reaction mixture was transferred to a 30 kDa Amicon Ultra 0.5 mL centrifugal filter. The original Eppendorf tube was washed with PBS (pH 7.4, 1×; 2 × 75 μL), and the washings were combined on the filter, which was centrifuged (4 min, 10,000 × rcf). PBS (pH 7.4, 1×; 250 μL) was added to the remaining solution on filter (~100 μL), followed by centrifugation (4 min, 10,000 × rcf). This dilution and centrifugation step was repeated four additional times. The residual liquid on the filter was collected, yielding approximately 100 μL of antibody solution in PBS (pH 7.4, 1×). The concentration of the reduced trastuzumab was determined by UV absorbance at 280 nm ($\varepsilon_{280}$ = 225,000 $M^{-1}$ $cm^{-1}$) using a NanoDrop spectrophotometer or microplate reader. Daratumumab, cetuximab, and trastuzumab were evaluated, with trastuzumab chosen as the model antibody substrate. When using NHS ester **2** at up to 100 equivalents, no significant protein loss was observed; the final antibody concentration corresponded to >65% yield (loss likely due to handling). At 150 equivalents of **2**, approximately 50% yield (~5 μM of conjugated tmab) was obtained, and LC–MS analysis indicated an average drug-to-antibody ratio (DAR) of 12.8 carboranes (128 boron atoms) per trastuzumab. When 200 equivalents of **2** were used, noticeable precipitation occurred, leaving negligible protein in solution. In contrast, with NHS ester **1**, which has a more rigid structure, precipitation occurred at lower reagent equivalents - already at 75 equivalents - with optimal conjugation efficiency observed at 50 equivalents (see Figure 5 in the manuscript and LC-MS analysis below).

The same trend was observed for conjugation of NHS esters **1** and **2** with daratumumab and cetuximab (Figure 5).

##### LC–MS analysis of antibody conjugates and DAR determination (Fig. 5b)

For LC–MS analysis, an aliquot of the reaction mixture was taken after workup as described above and analyzed on an Agilent 6550 LC-MS spectrometer using Zorbax 300SB-C3 column, as described in the General information section.

Sample preparation: An aliquot (5 µL) of the reaction mixture was reduced with DTT in ammonium bicarbonate buffer. Specifically, 5 µL of the reaction mixture were added to 44 µL of 50 mM ammonium bicarbonate buffer, followed by 1 µL of a 1 M DTT stock solution (final DTT concentration: 20 mM). Both reagents were prepared in Milli-Q water. The mixture was pipetted up and down 15 times and incubated for 30–60 min at 40 °C. After incubation, the solution was transferred to a 10 kDa Amicon Ultra 0.5 mL centrifugal filter. The parent Eppendorf tube was rinsed with PBS (pH 7.4 1×; 2 × 75 µL), and the washings were combined on the filter, which was centrifuged (4 min, 10,000 × rcf). PBS (pH 7.4, 1×; 250 µL) was added to the remaining solution (~100 µL), followed by centrifugation (4 min, 10,000 × rcf). This dilution and centrifugation step was repeated two additional times. The remaining liquid on the filter (~100 µL) constituted the reduced antibody solution in PBS (pH 7.4, 1×). A 7 µL of this solution (corresponding to about 500 ng of protein) was injected for LC–MS analysis.

**LC-MS analysis of conjugation reactions (Fig. 5b)**

Data were processed using Agilent MassHunter Workstation BioConfirm Software (v10.0). Protein deconvolution was performed using the maximum entropy algorithm. Conjugation efficiency was calculated from the mass differences between the unmodified antibody light and heavy chains and their corresponding modified forms. The expected mass shifts were determined based on the mass of the attached NHS ester fragments (**1** or **2**) minus one proton, accounting for amide bond formation.

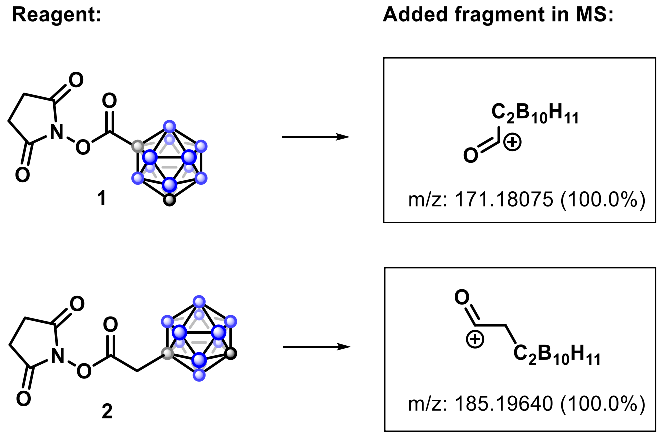

#### Example of Trastuzumab (Tmab) reaction for determining conjugation efficiency

LC chromatogram: MS spectrum: Deconvoluted MS spectrum:

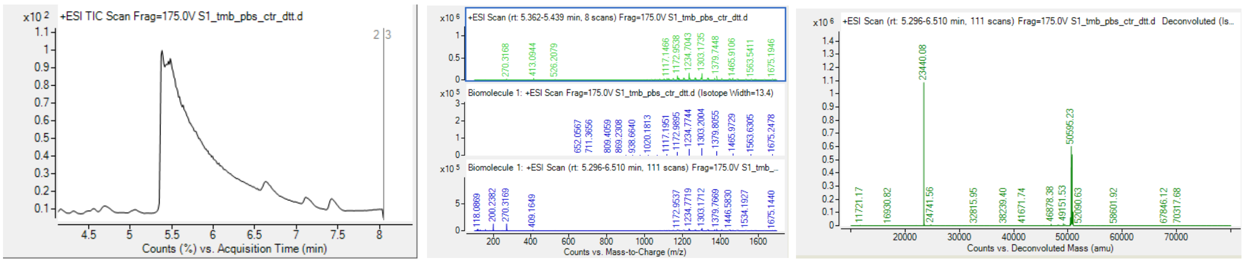

**Figure S27**. LC chromatogram, MS spectrum, and deconvoluted MS spectrum of native trastuzumab.

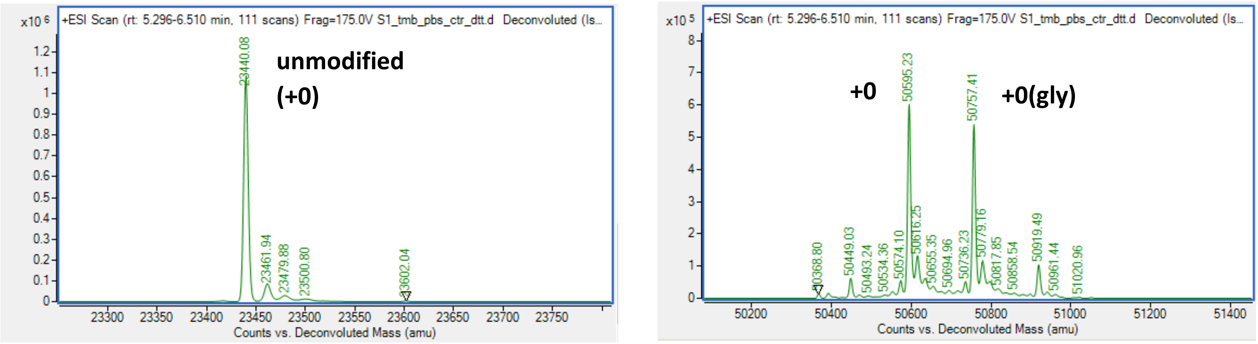

**Figure S28**. Deconvoluted MS of WT Tmab, the light chain and heavy chain. The Tmab light chain corresponds to a molecular weight of 23,440 Da. The Tmab heavy chain shows two main deconvoluted masses at 50,595 Da and 50,757 Da, corresponding to the non-glycosylated (unmodified, +0) and glycosylated (unmodified-gly, +0(gly)) heavy-chain forms, respectively. The mass difference of approximately +162 Da is consistent with the addition of a hexose residue (e.g., galactose or glucose), which is characteristic of antibody Fc-region N-glycosylation. These two species are commonly observed for recombinant IgG1 antibodies such as trastuzumab.[10,11]

#### Reaction of Tmab (10 µM) with NHS reagent 2 (30 equiv.):

LC chromatogram: MS spectrum: Deconvoluted MS spectrum:

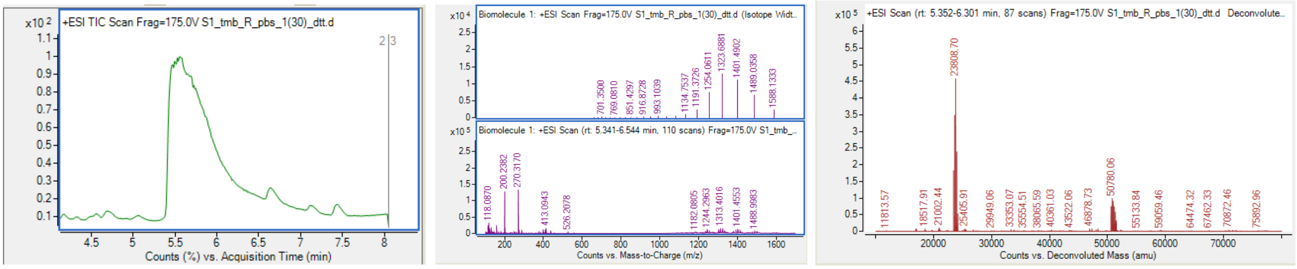

**Figure S29**. LC chromatogram, MS spectrum, and deconvoluted MS spectrum.

Deconvoluted MS spectrum of light chain:

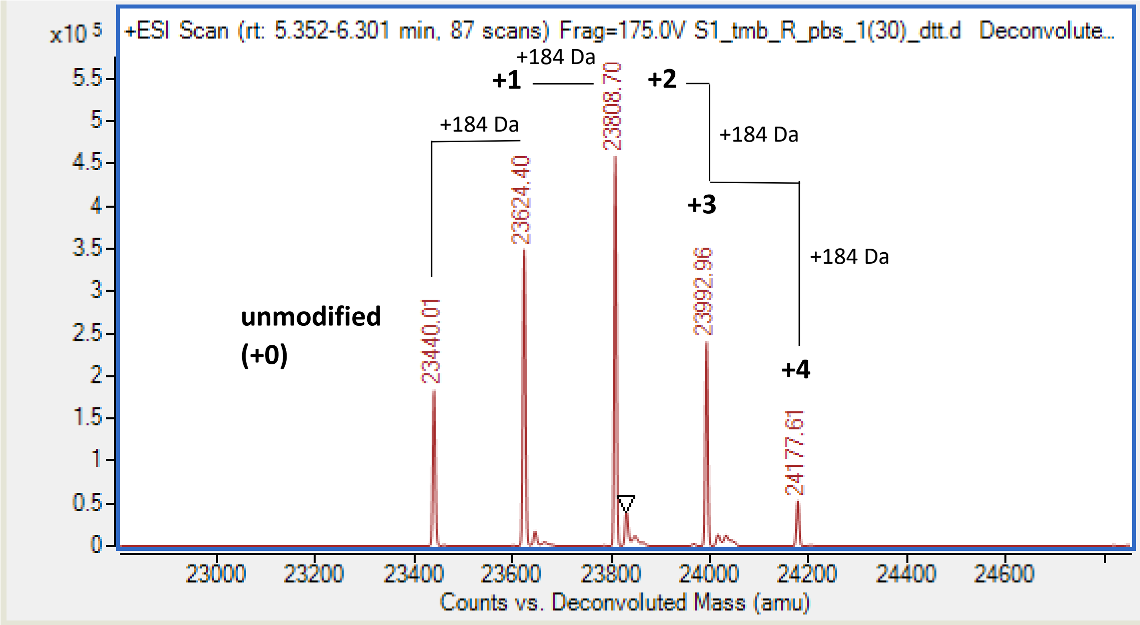

**Figure S30**. Example of conjugation efficiency determination from a deconvoluted light chain MS spectrum. "+1", "+2", etc., denote the number of conjugation events, calculated from the mass difference between the native antibody and the corresponding modified species. Each successive addition reflects the incorporation of one conjugation fragment from reagent **2** (with a mass increase corresponding to the added fragment minus one proton due to amide bond formation). For example:

+1 modification: 23624.4 – 23440.0 Da = 184.4 Da → one fragment added from reagent **2**;

+2 modification: 23808.7 – 23440.0 Da = 368.7 Da → two fragments added from reagent **2**. etc.

Deconvoluted MS spectrum of heavy chain:

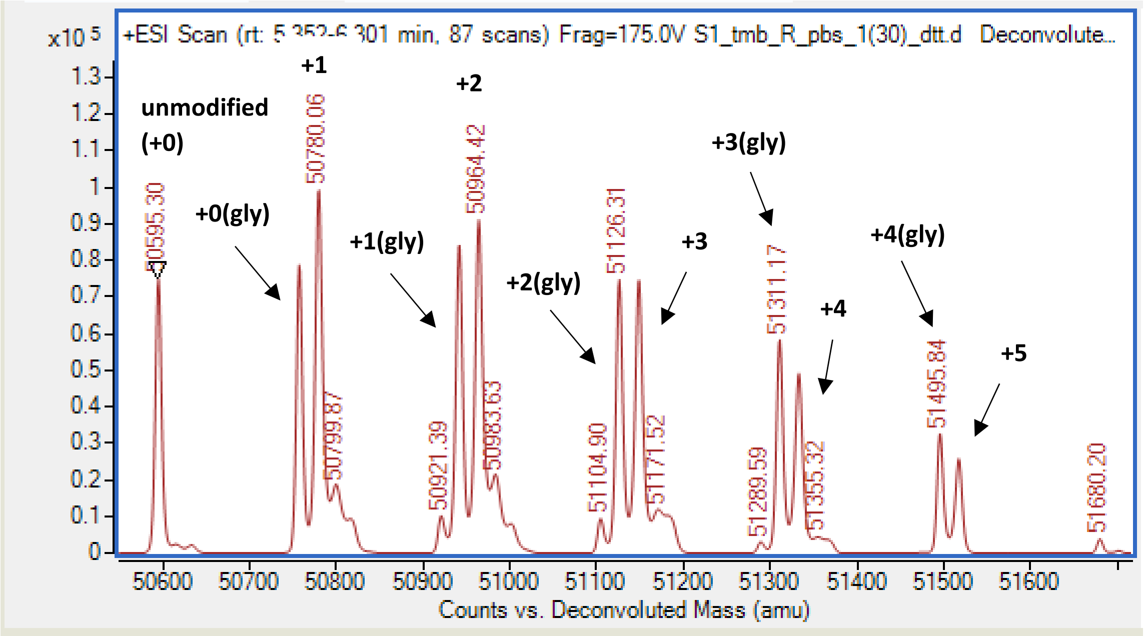

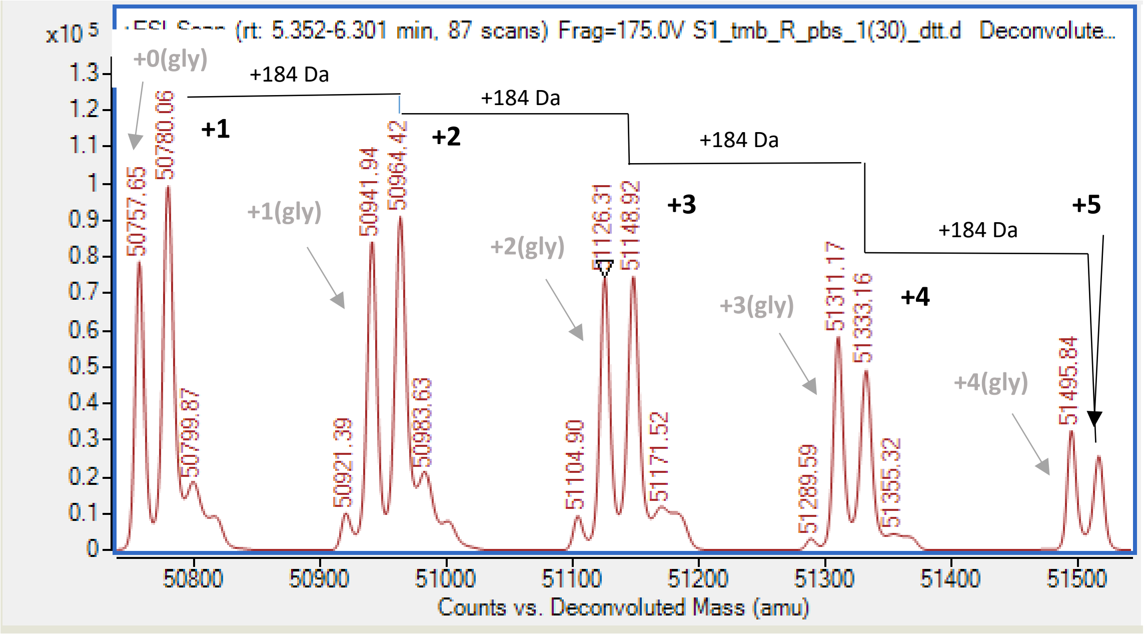

**Figure S31.** Example of conjugation efficiency determination from a deconvoluted heavy chain MS spectrum. The conjugation efficiency of the heavy chain was determined in the same manner as described above for the light chain. In this case, two sets of signals corresponding to the unmodified heavy chain of trastuzumab (unmodified, 50,595 Da; black series above) and the glycosylated form (50,757 Da; grey series) were observed and analyzed in the same manner. An approximate average between the non-glycosylated and glycosylated conjugate chains (0, +1, +2, …) was used to estimate the indicated ratio for the DAR calculation in the table below.

**Calcucation of light and heavy chains DAR, and overall DAR**

The drug-to-antibody ratio (DAR) for the light and heavy chains was calculated from the relative intensity ratios for mass signals obtained from the deconvoluted MS spectra. The intensities of the unmodified and modified species were normalized for each chain separately to determine the extent of conjugation. Each light- or heavy-chain modification corresponds to the number of bound fragments, as identified from the mass shifts and their relative intensities (see above). The contribution for each species was calculated as (number of modifications × relative intensity), and the DAR for each chain was obtained by dividing the total contribution by the total summed intensity for that chain.

The overall DAR was then calculated as: DAR (overall) = 2×DAR (light) + 2×DAR (heavy)

**Table S4.** Calcualtion of DAR in example of reaction of Tmab (10 μM) with NHS reagent **2** (30 equiv.)

| Light ch. modif. | Ratio (from MS) | Contribution | Heavy ch. modif. | Ratio (from MS) | Contribution |
|---|---|---|---|---|---|
| 0 | 2 | 0 | 0 | 1.6 | 0 |
| +1 | 3.5 | 3.5 | +1 | 1.9 | 1.9 |
| +2 | 4.6 | 9.2 | +2 | 1.7 | 3.4 |
| +3 | 2.5 | 7.5 | +3 | 1.3 | 3.9 |
| +4 | 0.6 | 2.4 | +4 | 0.8 | 3.2 |
| +5 | 0 | 0 | +5 | 0.3 | 1.5 |
| +6 | 0 | 0 | +6 | 0 | 0 |
| | | | +7 | | |
| total | 13.2 | 22.6 | total | 7.6 | 13.9 |
| DAR light = total contribution / total ratio | | | | | |
| DAR light single | | 1.712121 | DAR heavy single | | 1.828947 |

DAR overall:  7.082

**Reaction of trastuzumab with 150 equiv. of 2 (tmab conjugate with DAR 12.8)**

LC chromatogram: MS spectrum: Deconvoluted MS spectrum:

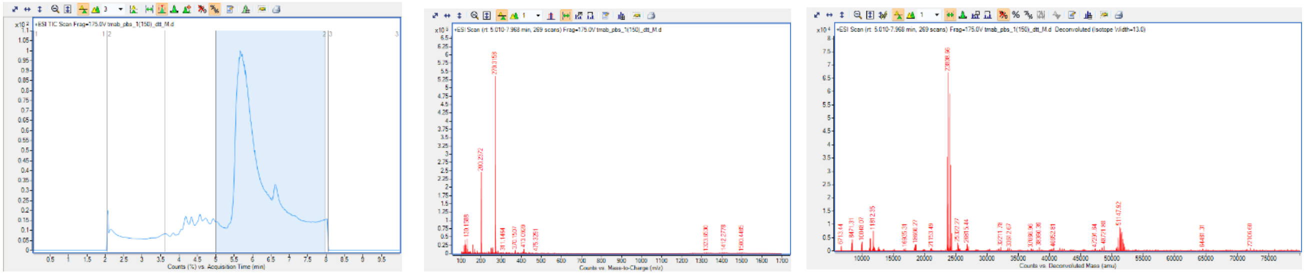

**Figure S32**. LC chromatogram, MS spectrum, and deconvoluted MS spectrum.

Deconvoluted MS spectrum of light chain:

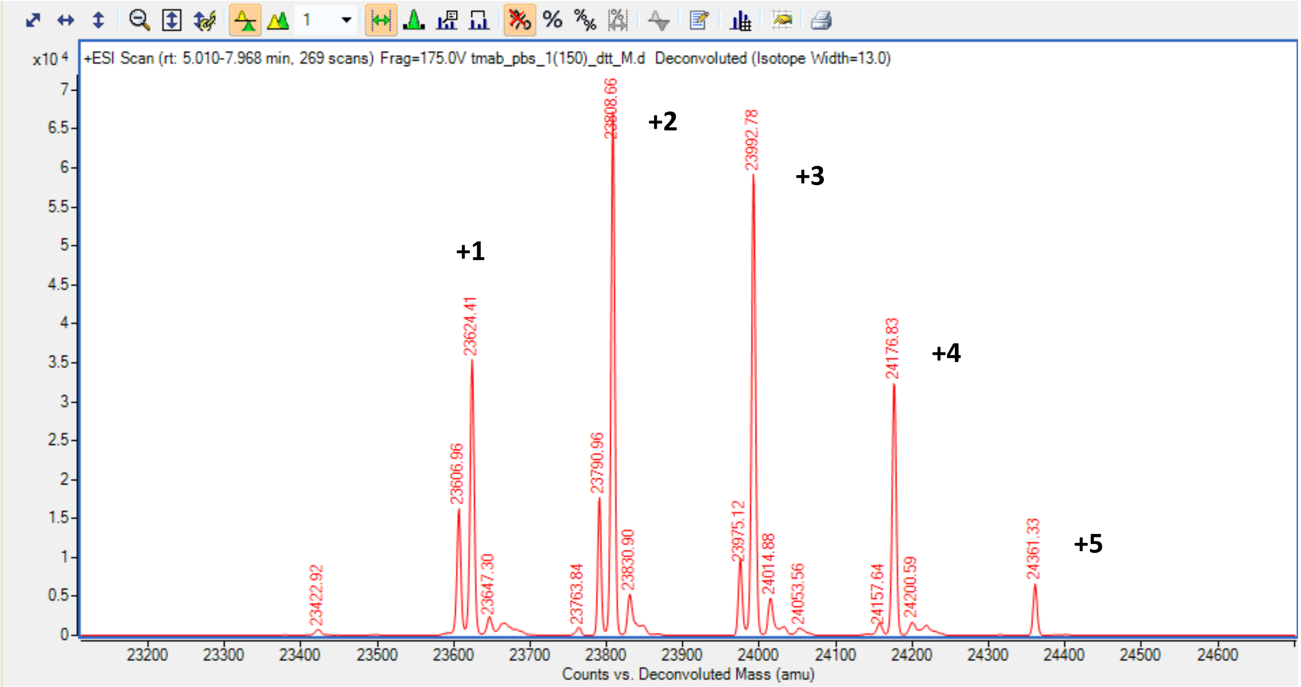

**Figure S33**. Conjugation efficiency determination from a deconvoluted light chain MS spectrum. “+1”, “+2”, etc., denote the number of conjugation events, calculated from the mass difference between the native antibody and the corresponding modified species (for more detailed calculations see Figure S29).

Deconvoluted MS spectrum of heavy chain:

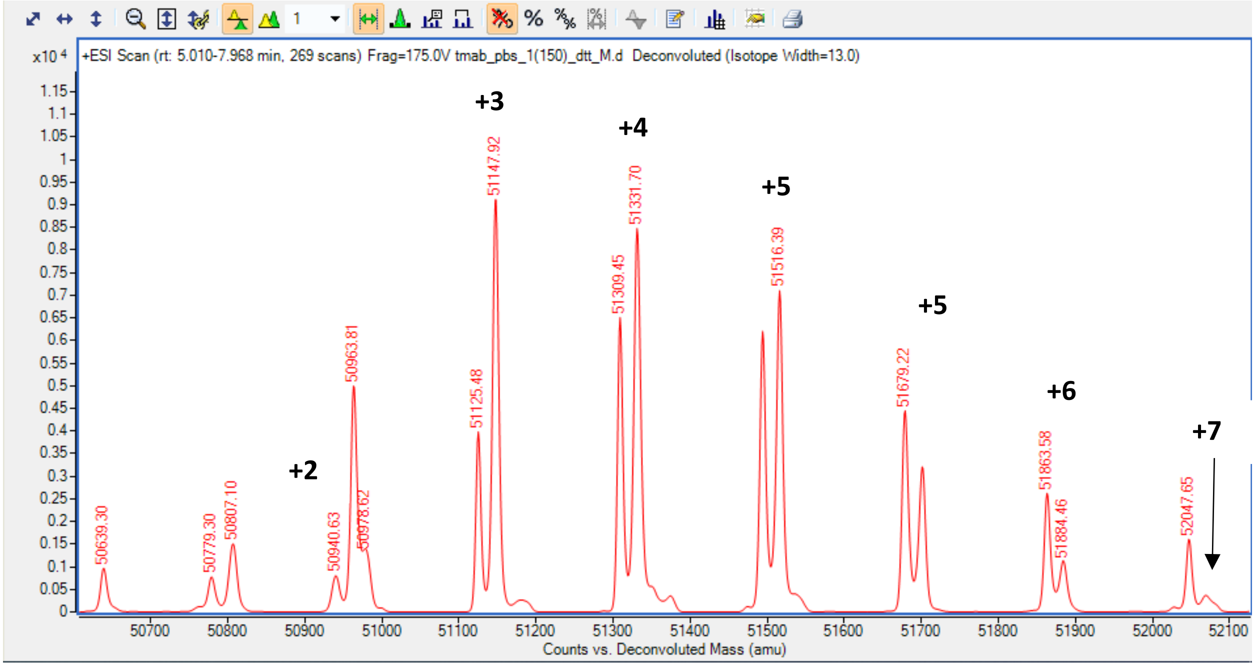

**Figure S34**. Conjugation efficiency determination from a deconvoluted heavy chain MS spectrum. “+2”, “+3”, etc., denote the number of conjugation events, calculated from the mass difference between the native antibody and the corresponding modified species (for more detailed calculations see Figure S29).

**Table S5**. Calcualtion of DAR in example of reaction of Tmab (10 μM) with NHS reagent **2** (150 equiv.)

| light | ratio | contribution | heavy | ratio | contribution |
|---|---|---|---|---|---|
| 0 | 0 | 0 | 0 | 0 | 0 |
| 1 | 3.5 | 3.5 | 1 | 0.05 | 0.05 |
| 2 | 6.5 | 13 | 2 | 0.5 | 1 |
| 3 | 6 | 18 | 3 | 0.9 | 2.7 |
| 4 | 3.5 | 14 | 4 | 0.85 | 3.4 |
| 5 | 0.75 | 3.75 | 5 | 0.7 | 3.5 |
| 6 | | 0 | 6 | 0.35 | 2.1 |
| | | | 7 | 0.1 | 0.7 |
| total | 20.25 | 52.25 | total | 3.35 | 12.75 |

dar light = total contribution / total ratio

dar light single 2.580247 DAR heavy single 3.80597

DAR overall: 12.7724

In this manner, the conjugation efficiency was determined for all conjugation reactions.

**Results of conjugation reations (Fig. 5b)**

**trastuzumab (tmab)**

| **Reactions in PBS (pH 7.4, 1×)** | | | | |
|---|---|---|---|---|
| Reagent | equiv | Light DAR | Heavy DAR | Overall DAR |
| **2** | 10 | 1.034483 | 0.818182 | 3.705329 |
| **2** | 30 | 1.712121 | 1.828947 | 7.082137 |
| **2** | 50 | 2.512821 | 2.632911 | 10.29146 |
| **2** | 75 | 2.566667 | 2.197674 | 9.528682 |
| **2** | 100 | 2.250554 | 3.454545 | 11.4102 |
| **2** | 150 | 2.580247 | 3.80597 | 12.77243 |
| **1** | 10 | 0.16 | 0.212 | 0.744242 |
| **1** | 30 | 0.568862 | 0.47619 | 2.090106 |
| **1** | 50 | 1.533333 | 2.934783 | 8.936232 |
| **1** | 75 | 1.090909 | 1.542857 | 5.267532 |
| **1** | 100 | 1.105882 | 1.445783 | 5.103331 |
| **1** | 150 | 1.026738 | 1.697183 | 5.447842 |

| **Reactions in Histide (pH 7.4) buffer** | | | | |
|---|---|---|---|---|
| Reagent | equiv | Light DAR | Heavy DAR | Overall DAR |
| **2** | 10 | 0.2 | 0.1 | 0.6 |
| **2** | 100 | 0.78125 | 0.459459 | 2.481419 |

Although histidine buffer was expected to be unsuitable due to the nucleophilicity of its imidazole group (which turned to be beneficial for Pd catalyzed conjugation[12]),[13] we tested it nonetheless and confirmed poor NHS ester coupling efficiency.

**Daratumumab (dar)**

| Reagent | equiv | Light DAR | Heavy DAR | DAR |
|---|---|---|---|---|
| **2** | 30 | 1.323944 | 0.411348 | 3.470582 |
| **2** | 50 | 1.88 | 2.089172 | 7.938344 |
| **2** | 75 | 2.039735 | 3.103208 | 10.28589 |
| **2** | 100 | 2.301587 | 3.343582 | 11.29034 |
| **1** | 30 | 0.292818 | 0.983333 | 2.552302 |

**Cetuximab (cet)**

| Reagent | equiv | Light DAR | Heavy DAR | DAR |
|---|---|---|---|---|
| **2** | 30 | 1.189189 | 1.95 | 6.278378 |
| **2** | 75 | 1.527778 | 2.777778 | 8.611111 |
| **2** | 100 | 1.702128 | 2.634783 | 8.673821 |

#### Mapping of conjugation sites in antibody conjugates (Fig. 5c)

Mapping of conjugation sites was performed by nLC–MS/MS analysis on a Thermo Fisher Orbitrap Fusion Eclipse Tribrid Mass Spectrometer. Antibody samples were digested enzymatically with trypsin, and the resulting peptides were analyzed under the conditions described in the General Information section. The tryptic digestion protocol was adapted from previously reported method for trastuzumab sequencing.[14]

Briefly, native trastuzumab and its conjugates (DAR = 3.7, 11.4, and 12.8, as determined by LC-MS) were digested with sequence-grade trypsin (Trypsin/Lys-C Mix, Mass Spec Grade from Promega) at a 1:10 (w/w) enzyme-to-tmab ratio. Trastuzumab samples were first prepared in a concentration of 3 µM, and 1 µL of corresponding Tmab conjugate solution was added to 15 µL of denaturing reagent (6 M guanidine·HCl in 50 mM ammonium bicarbonate, at pH 8) followed by 2 µL of 100 mM DTT. The mixture was incubated at 95 °C for 5 min and then cooled to room temperature. Subsequently, 3 µL of 100 mM iodoacetamide were added, and the mixture was incubated in the dark for 20 min. Trypsin (5 µL; 1:10 w/w) was then added to this solution, and digestion was carried out at room temperature for 3 h, followed by the addition of an additional 5 µL of trypsin for overnight digestion (20 h) at room temperature. After digestion, 5 µL of concentrated HCl and 200 µL of 0.1% TFA in water were added to quench the reaction. Samples were desalted using ZipTip C18 tips and eluted three times with 5 µL of 0.1% TFA in a 1:1 $H_2O$/ACN mixture, and further diluted to 10 µL in $H_2O$/ACN with 0.1% TFA - 1 µL of this solution was injected for nLC–MS/MS analysis. ThermoFisher Xcalibur software package and PEAKS Studio 8.5 were used for data analysis with carborane conjugation fragment *m/z* 184.1886 (carborane fragment added: 185.1964 – 1.008 (proton) from amide bond formation event) being added to the listo of PTM.

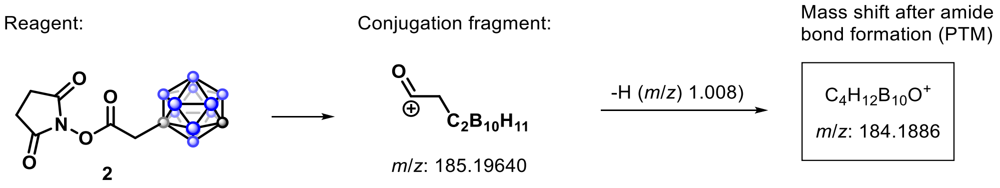

Alignment of all three antibodies and labeling of all (complementarity-determining region) CDR loops: sites without asterisks indicate variable positions. CDR regions were analyzed for trastuzumab (tmab), and daratumumab (dara), for cetuximab (cet) the CDR is based on the alignment.[15]

In the sequence, the heavy chain is analyzed first and then the light chain:

**Table S6**. Sequence alignment of trastuzumab (tmab), daratumumab (dara), and cetuximab (cetux)

```
CLUSTAL 2.1 multiple sequence alignment

tmab            EVQLVESGGGLVQPGGSLRLSCAASGFNIKDTYIHWVRQAPGKGLEWVARIYPTNGYTRY
dara            EVQLLESGGGLVQPGGSLRLSCAVSGFTFNSFAMSWVRQAPGKGLEWVSAISGSGGGTYY
cetux           QVQLKQSGPGLVQPSQSLSITCTVSGFSLTNYGVHWVRQSPGKGLEWLGVIWS-GGNTDY
                :*** :** *****. ** ::*:.***.:..  : ****:*******:. *   .* * *

tmab            ADSVKGRFTISADTSKNTAYLQMNSLRAEDTAVYYCSR--WGGDGFYAMDYWGQGTLVTV
dara            ADSVKGRFTISRDNSKNTLYLQMNSLRAEDTAVYFCAKDKILWFGEPVFDYWGQGTLVTV
cetux           NTPFTSRLSINKDNSKSQVFFKMNSLQSNDTAIYYCAR--ALTYYDYEFAYWGQGTLVTV
                  ....*::*. *.**.  :::****:::***:*:*::          : **********

tmab            SSASTKGPSVFPLAPSSKSTSGGTAALGCLVKDYFPEPVTVSWNSGALTSGVHTFPAVLQ
dara            SSASTKGPSVFPLAPSSKSTSGGTAALGCLVKDYFPEPVTVSWNSGALTSGVHTFPAVLQ
cetux           SAASTKGPSVFPLAPSSKSTSGGTAALGCLVKDYFPEPVTVSWNSGALTSGVHTFPAVLQ
                *:**********************************************************

tmab            SSGLYSLSSVVTVPSSSLGTQTYICNVNHKPSNTKVDKKVEPKSCDKTHTCPPCPAPELL
dara            SSGLYSLSSVVTVPSSSLGTQTYICNVNHKPSNTKVDKRVEPKSCDKTHTCPPCPAPELL
cetux           SSGLYSLSSVVTVPSSSLGTQTYICNVNHKPSNTKVDKRVEPKSCDKTHTCPPCPAPELL
                **************************************:*********************

tmab            GGPSVFLFPPKPKDTLMISRTPEVTCVVVDVSHEDPEVKFNWYVDGVEVHNAKTKPREEQ
dara            GGPSVFLFPPKPKDTLMISRTPEVTCVVVDVSHEDPEVKFNWYVDGVEVHNAKTKPREEQ
cetux           GGPSVFLFPPKPKDTLMISRTPEVTCVVVDVSHEDPEVKFNWYVDGVEVHNAKTKPREEQ
                ************************************************************

tmab            YNSTYRVVSVLTVLHQDWLNGKEYKCKVSNKALPAPIEKTISKAKGQPREPQVYTLPPSR
dara            YNSTYRVVSVLTVLHQDWLNGKEYKCKVSNKALPAPIEKTISKAKGQPREPQVYTLPPSR
cetux           YNSTYRVVSVLTVLHQDWLNGKEYKCKVSNKALPAPIEKTISKAKGQPREPQVYTLPPSR
                ************************************************************

tmab            EEMTKNQVSLTCLVKGFYPSDIAVEWESNGQPENNYKTTPPVLDSDGSFFLYSKLTVDKS
dara            EEMTKNQVSLTCLVKGFYPSDIAVEWESNGQPENNYKTTPPVLDSDGSFFLYSKLTVDKS
cetux           EEMTKNQVSLTCLVKGFYPSDIAVEWESNGQPENNYKTTPPVLDSDGSFFLYSKLTVDKS
                ************************************************************

tmab            RWQQGNVFSCSVMHEALHNHYTQKSLSLSPG-DIQMTQSPSSLSASVGDRVTITCRASQD
dara            RWQQGNVFSCSVMHEALHNHYTQKSLSLSPGKEIVLTQSPATLSLSPGERATLSCRASQS
cetux           RWQQGNVFSCSVMHEALHNHYTQKSLSLSPGKDILLTQSPVILSVSPGERVSFSCRASQS
                ******************************* :* :****  ** * *:*.:::*****.

tmab            VNTAVAWYQQKPGKAPKLLIYSASFLYSGVPSRFSGSRSGTDFTLTISSLQPEDFATYYC
dara            VSSYLAWYQQKPGQAPRLLIYDASNRATGIPARFSGSGSGTDFTLTISSLEPEDFAVYYC
cetux           IGTNIHWYQQRTNGSPRLLIKYASESISGIPSRFSGSGSGTDFTLSINSVESEDIADYYC
                :.: : ****:.. :*:***  **   :*:*:***** *******:*.*::.**:* ***

tmab            QQHYTTPPTFGQGTKVEIKRTVAAPSVFIFPPSDEQLKSGTASVVCLLNNFYPREAKVQW
dara            QQRSNWPPTFGQGTKVEIKRTVAAPSVFIFPPSDEQLKSGTASVVCLLNNFYPREAKVQW
cetux           QQNNNWPTTFGAGTKLELKRTVAAPSVFIFPPSDEQLKSGTASVVCLLNNFYPREAKVQW
                **. . *.*** ***:*:******************************************
```

```
tmab            KVDNALQSGNSQESVTEQDSKDSTYSLSSTLTLSKADYEKHKVYACEVTHQGLSSPVTKS
dara            KVDNALQSGNSQESVTEQDSKDSTYSLSSTLTLSKADYEKHKVYACEVTHQGLSSPVTKS
cetux           KVDNALQSGNSQESVTEQDSKDSTYSLSSTLTLSKADYEKHKVYACEVTHQGLSSPVTKS
                ************************************************************

tmab            FNRGEC
dara            FNRGEC
cetux           FNRGEC
```

**Trastuzumab (tmab)**

Wt:

EVQLVESGGGLVQPGGSLRLSCAASGFNIKDTYIHWVRQAPGKGLEWVARIYPTNGYTRYA
DSVKGRFTISADTSKNTAYLQMNSLRAEDTAVYYCSRWGGDGFYAMDYWGQGTLVTVSS
ASTKGPSVFPLAPSSKSTSGGTAALGCLVKDYFPEPVTVSWNSGALTSGVHTFPAVLQSSGL
YSLSSVVTVPSSSLGTQTYICNVNHKPSNTKVDKKVEPKSCDKTHTCPPCPAPELLGGPSVFL
FPPKPKDTLMISRTPEVTCVVVDVSHEDPEVKFNWYVDGVEVHNAKTKPREEQYNSTYRV
VSVLTVLHQDWLNGKEYKCKVSNKALPAPIEKTISKAKGQPREPQVYTLPPSREEMTKNQV
SLTCLVKGFYPSDIAVEWESNGQPENNYKTTPPVLDSDGSFFLYSKLTVDKSRWQQGNVFS
CSVMHEALHNHYTQKSLSLSPGDIQMTQSPSSLSASVGDRVTITCRASQDVNTAVAWYQQK
PGKAPKLLIYSASFLYSGVPSRFSGSRSGTDFTLTISSLQPEDFATYYCQQHYTTPPTFGQGTK
VEIKRTVAAPSVFIFPPSDEQLKSGTASVVCLLNNFYPREAKVQWKVDNALQSGNSQESVTE
QDSKDSTYSLSSTLTLSKADYEKHKVYACEVTHQGLSSPVTKSFNRGEC

Modified positions:

C96, C147, C264, C370, C428, C537, C583, C643 (cysteines are modified by iodoacetamide during protein denaturation process)

Legend (color code the same for all analyses):

CDR region

Underlined region represents light chain of trastzumab

Peptide detected in unmodified antibody

**Trastuzumab DAR 3.7** (reaction of tmab with 10 equiv. of NHS ester **2**, Fig. 5b)

EVQLVESGGGLVQPGGSLRLSCAASGFNIKDTYIHWVRQAPGKGLEWVARIYPTNGYTRYA
DSVKGRFTISADTSKNTAYLQMNSLRAEDTAVYYCSRWGGDGFYAMDYWGQGTLVTVSS
ASTKGPSVFPLAPSSKSTSGGTAALGCLVKDYFPEPVTVSWNSGALTSGVHTFPAVLQSSGL
YSLSSVVTVPSSSLGTQTYICNVNHKPSNTKVDKKVEPKSCDKTHTCPPCPAPELL**GGPSVF**

LFPPKPKDTLMISRTPEVTCVVVDVSHEDPEVKFNWYVDGVEVHNAKTKPREEQYNSTYRV
VSVLTVLHQDWLNGKEYKCKVSNKALPAPIEKTISKAKGQPREPQVYTLPPSREEMTKNQV
SLTCLVKGFYPSDIAVEWESNGQPENNYKTTPPVLDSDGSFFLYSKLTVDKSRWQQGNVFS
CSVMHEALHNHYTQKSLSLSPGDIQMTQSPSSLSASVGDRVTITCRASQDVNTAVAWYQQK
PGKAPKLLIYSASFLYSGVPSRFSGSRSGTDFTLTISSLQPEDFATYYCQQHYTTPPTFGQGTK
VEIKRTVAAPSVFIFPPSDEQLKSGTASVVCLLNNFYPREAKVQWKVDNALQSGNSQESVTE
QDSKDSTYSLSSTLTLSKADYEKHKVYACEVTHQGLSSPVTKSFNRGEC

Modified positions:

C147, C232, C264, C324, C370, C428, C537, C583, C643, C96

K: H65, H216, H217, H221, H225, H323, H343, H417, L42, L145, L188

Color legend (the same applies to all other analyses):

Underlined region represents light chain of trastzumab

CDR region

Peptide detected in unmodified (WT) antibody

Peptide that is only detected in the modified antibody but not in the unmodified antibody

**Peptide that is only detected in the unmodified antibody but not in the modified antibody**

Peptide is not detected

Carborane modification site detected

**Trastuzumab DAR 11.4** (reaction of tmab with 100 equiv. of NHS ester **2**, Fig. 5b)

EVQLVESGGGLVQPGGSLRLSCAASGFNIKDTYIHWVRQAPGKGLEWVARIYPTNGYTRYA
DSVKGRFTISADTSKNTAYLQMNSLRAEDTAVYYCSRWGGDGFYAMDYWGQGTLVTVSS
ASTKGPSVFPLAPSSKSTSGGTAALGCLVKDYFPEPVTVSWNSGALTSGVHTFPAVLQSSGL
YSLSSVVTVPSSSLGTQTYICNVNHKPSNTKVDKKVEPKSCDKTHTCPPCPAPELL**GGPSVF**
LFPPKPKDTLMISR**TPEVTCVVVDVSHEDPEVK**FNWYVDGVEVHNAKTKPREEQYNSTYR
VVSVLTVLHQDWLNGKEYKCKVSNKALPAPIEKTISKAKGQPREPQVYTLPPSREEMTKNQ
VSLTCLVKGFYPSDIAVEWESNGQPENNYKTTPPVLDSDGSFFLYSKLTVDKSRWQQGNVF
SCSVMHEALHNHYTQKSLSLSPGDIQMTQSPSSLSASVGDRVTITCRASQDVNTAVAWYQQ
KPGKAPKLLIYSASFLYSGVPSRFSGSRSGTDFTLTISSLQPEDFATYYCQQHYTTPPTFGQGT
KVEIKRTVAAPSVFIFPPSDEQLKSGTASVVCLLNNFYPREAKVQWKVDNALQSGNSQESVT
EQDSKDSTYSLSSTLTLSKADYEKHKVYACEVTHQGLSSPVTKSFNRGEC

Modified positions:

C147, C232, C324, C370, C428, C537, C583, C643, C96

K: H216, H, H221, H249, H293, H323, H343, H417, L39, L42, L145, L188

**Trastuzumab DAR 12.8** (reaction of tmab with 150 equiv. of NHS ester **2**, Fig. 5b)

EVQLVESGGGLVQPGGSLRLSCAASGFNIKDTYIHWVRQAPGKGLEWVARIYPTNGYTRYADSVKGRFTISADTSKNTAYLQMNSLRAEDTAVYYCSRWGGDGFYAMDYWGQGTLVTVSS**ASTK**GPSVFPLAPSSKSTSGGTAALGCLVKDYFPEPVTVSWNSGALTSGVHTFPAVLQSSGLYSLSSVVTVPSSSLGTQTYICNVNHKPSNTKVDKKVEPKSCDKTHTCPPCPAPELL**GGPSVF**LFPPKPKDTLMISRTPEVTCVVVDVSHEDPEVKFNWYVDGVEVHNAKTKPREEQYNSTYRVVSVLTVLHQDWLNGKEYKCKVSNKALPAPIEKTISKAKGQPREPQVYTLPPSREEMTKNQVSLTCLVKGFYPSDIAVEWESNGQPENNYKTTPPVLDSDGSFFLYSKLTVDKSRWQQGNVFSCSVMHEALHNHYTQKSLSLSPGDIQMTQSPSSLSASVGDRVTITCRASQDVNTAVAWYQQKPGKAPKLLIYSASFLYSGVPSRFSGSRSGTDFTLTISSLQPEDFATYYCQQHYTTPPTFGQGTKVEIKRTVAAPSVFIFPPSDEQLKSGTASVVCLLNNFYPREAKVQWKVDNALQSGNSQESVTEQDSKDSTYSLSSTLTLSKADYEKHKVYACEVTHQGLSSPVTKSFNRGEC

Modified positions:

C147, C232, C264, C324, C370, C428, C537, C583, C643, C96

K: H43, H65, H216, H217, H221, H225, H249, H323, H343, H417, L39488, L42, L188

#### Cell assays (Fig. 6)

BT-474 cells (ATCC, HTB-20, Lot: 70050151) were cultured in RPMI-1640 GlutaMAX medium (Gibco) supplemented with 10% fetal bovine serum (Gibco), and 1% penicillin-streptomycin (Gibco). Cells were maintained in a humidified incubator at 37 °C with 5% $CO_2$. For subculturing, cells were detached using trypsin-EDTA (Gibco).

A total of 10,000 cells in 150 µL of culture medium were seeded into each well of a 96-well microtiter plate and allowed to adhere for 20 h. Subsequently, the cells were treated with either modified or unmodified antibodies and incubated for an additional 20 h, after which cell viability was assessed using the MTT assay. For this assay, 17 µL of MTT reagent (5 mg/mL solution in 1× PBS; Thiazolyl Blue Tetrazolium Bromide, M5655, Sigma) was added to each well, followed by incubation for 3 h. Plates were then centrifuged (1000 g, 10 min, 4 °C), and the culture medium was carefully removed. The resulting formazan crystals were dissolved in 170 µL of solubilization solution (4 mM HCl and 0.1% Triton X-100 in isopropanol) by repeated pipetting until complete dissolution.

Absorbance was measured at 570 nm with background correction at 690 nm. The corrected absorbance values were further adjusted by subtracting the blank (wells without cells). Cell survival rates were normalized to untreated controls. Data analysis was performed using GraphPad Prism with one-way ANOVA, followed by Dunnett's multiple comparisons test. Statistical significance was denoted as follows: *$p < 0.05$; **$p < 0.01$; ***$p < 0.001$; ****$p < 0.0001$.

#### Copies of IR spectra

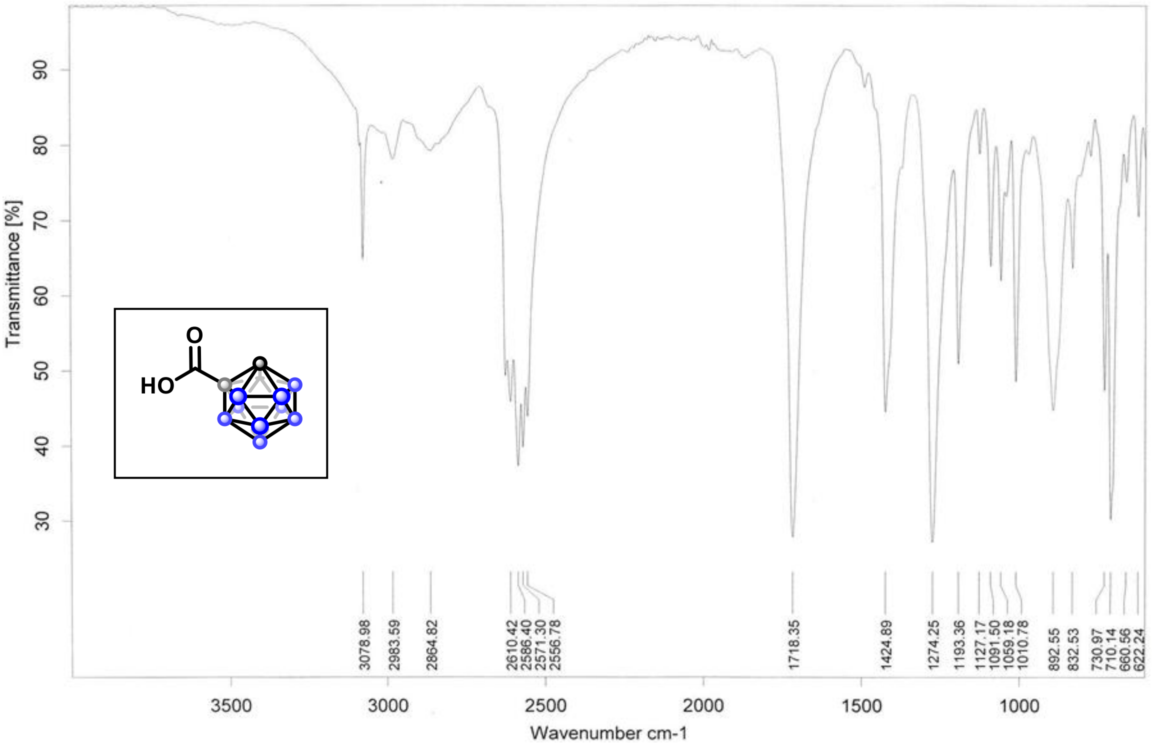

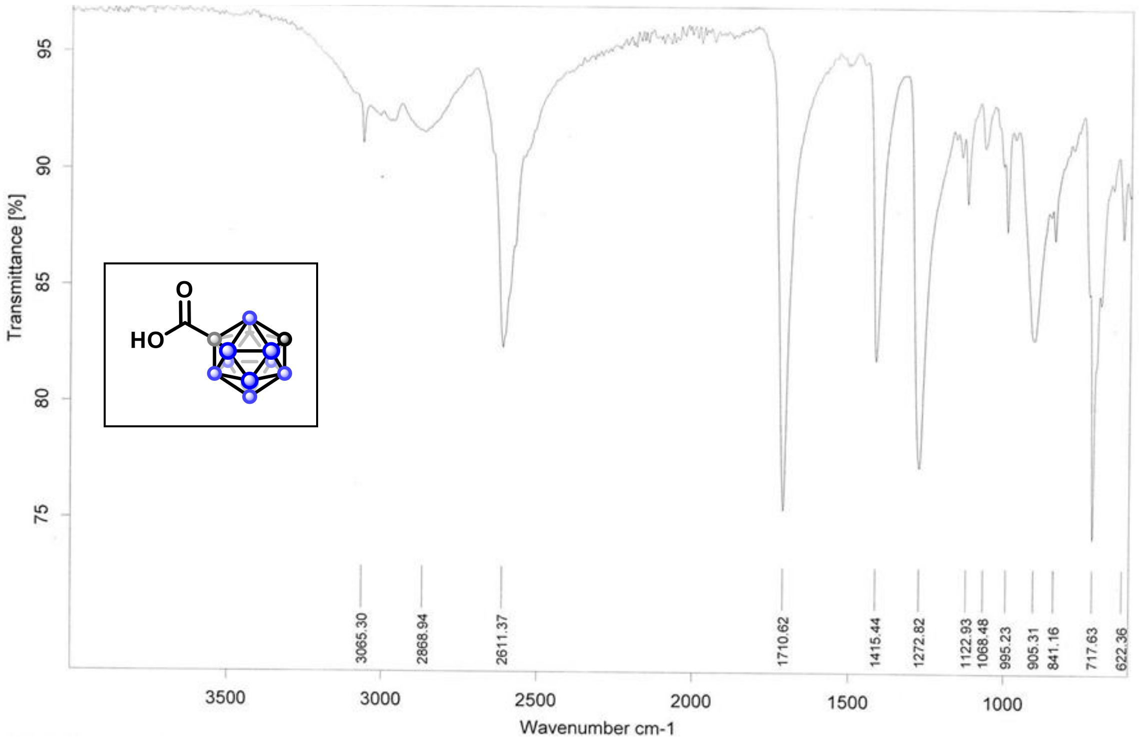

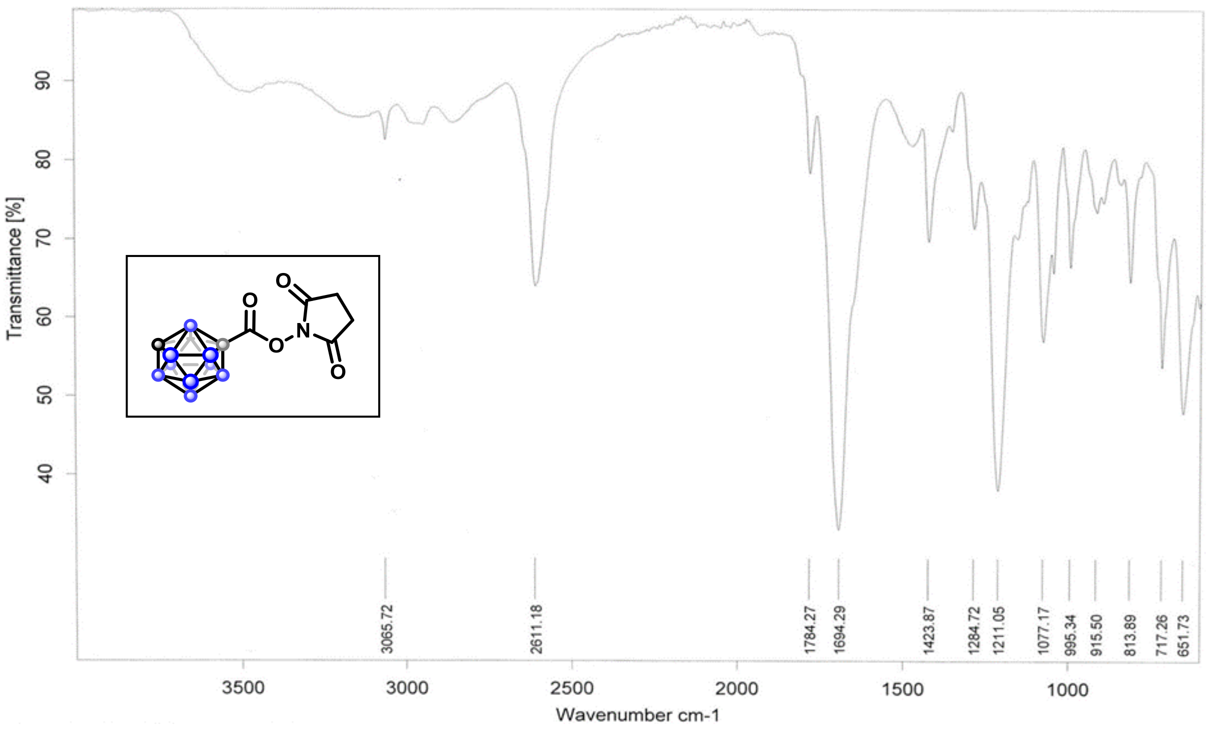

Transmittance [%]
Wavenumber cm-1
3065.72
2611.18
1784.27
1694.29
1423.87
1284.72
1211.05
1077.17
995.34
915.50
813.89
717.26
651.73

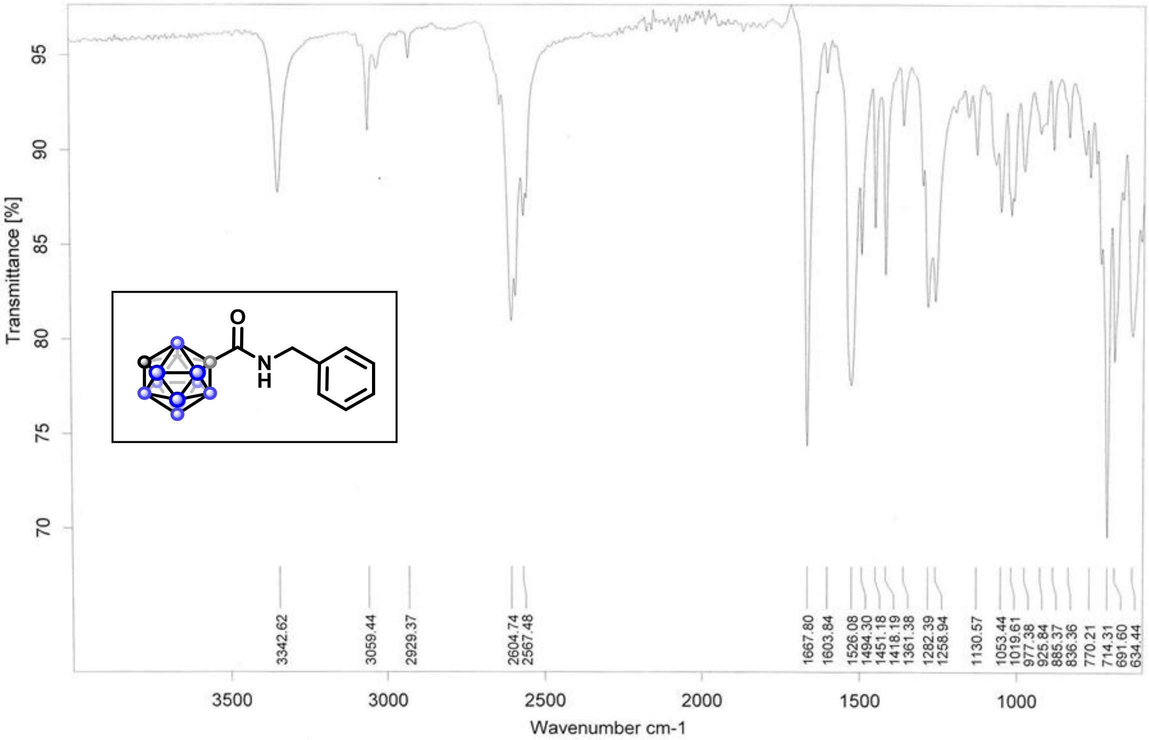

Transmittance [%]
Wavenumber cm-1
3342.62
3059.44
2929.37
2604.74
2567.48
1667.80
1603.84
1526.08
1494.30
1451.18
1418.19
1361.38
1282.39
1258.94
1130.57
1053.44
1019.61
977.38
925.84
885.37
836.36
770.21
714.31
691.60
634.44

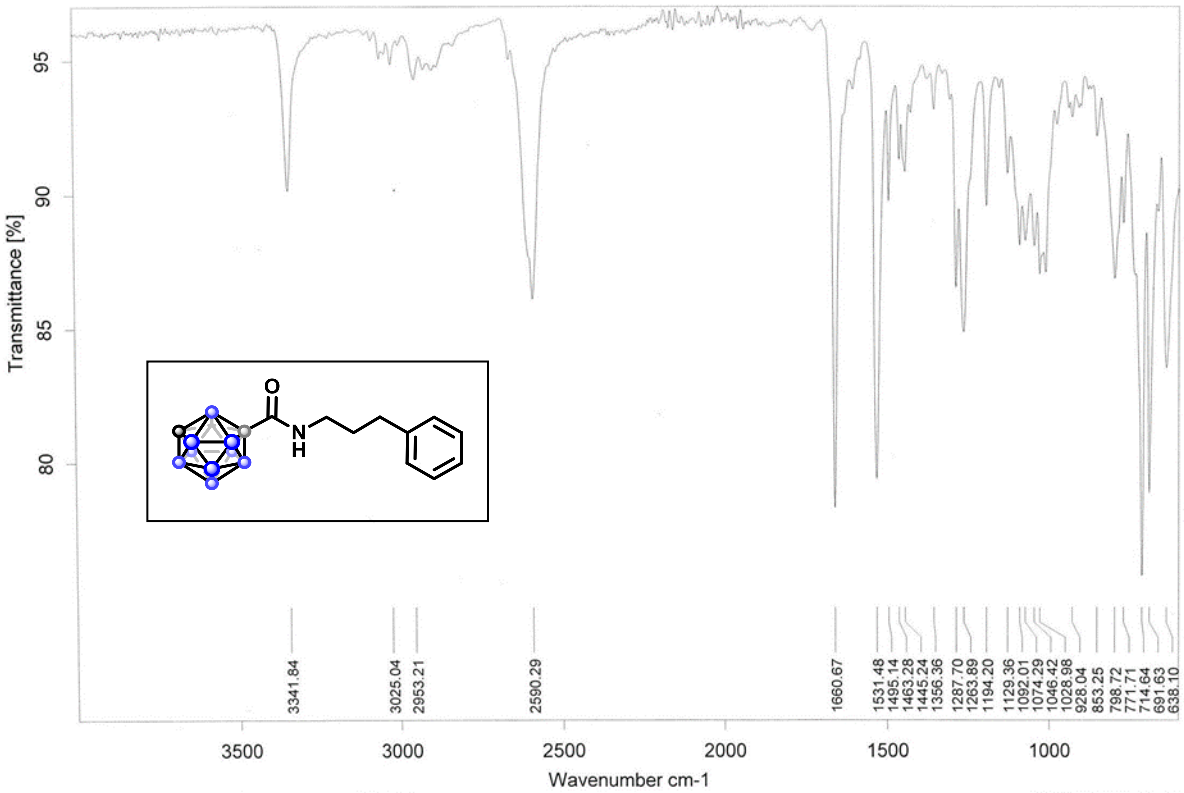

Transmittance [%]
95
90
85
80
O
N
H
3341.84
3025.04
2953.21
2590.29
1660.67
1531.48
1495.14
1463.28
1445.24
1356.36
1287.70
1263.89
1194.20
1129.36
1092.01
1074.29
1046.42
1028.98
928.04
853.25
798.72
771.71
714.64
691.63
638.10
3500
3000
2500
2000
1500
1000
Wavenumber cm-1

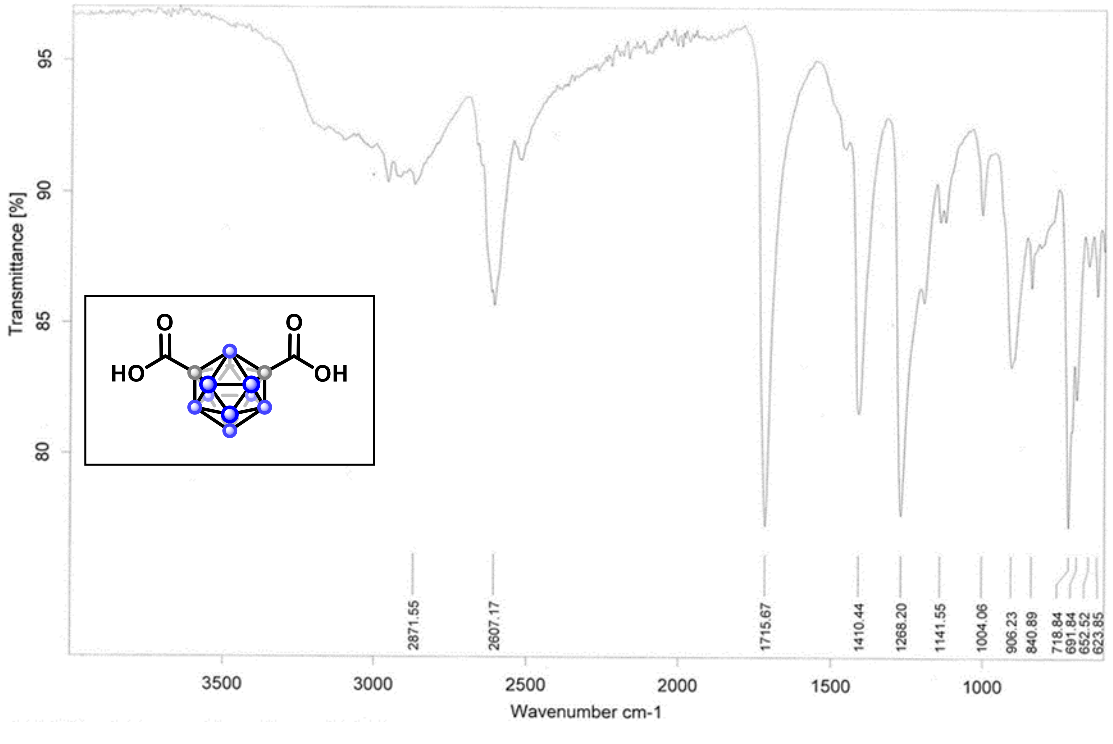

Transmittance [%]
95
90
85
80
O
HO
O
OH
2871.55
2607.17
1715.67
1410.44
1268.20
1141.55
1004.06
906.23
840.89
718.84
691.84
652.52
623.85
3500
3000
2500
2000
1500
1000
Wavenumber cm-1

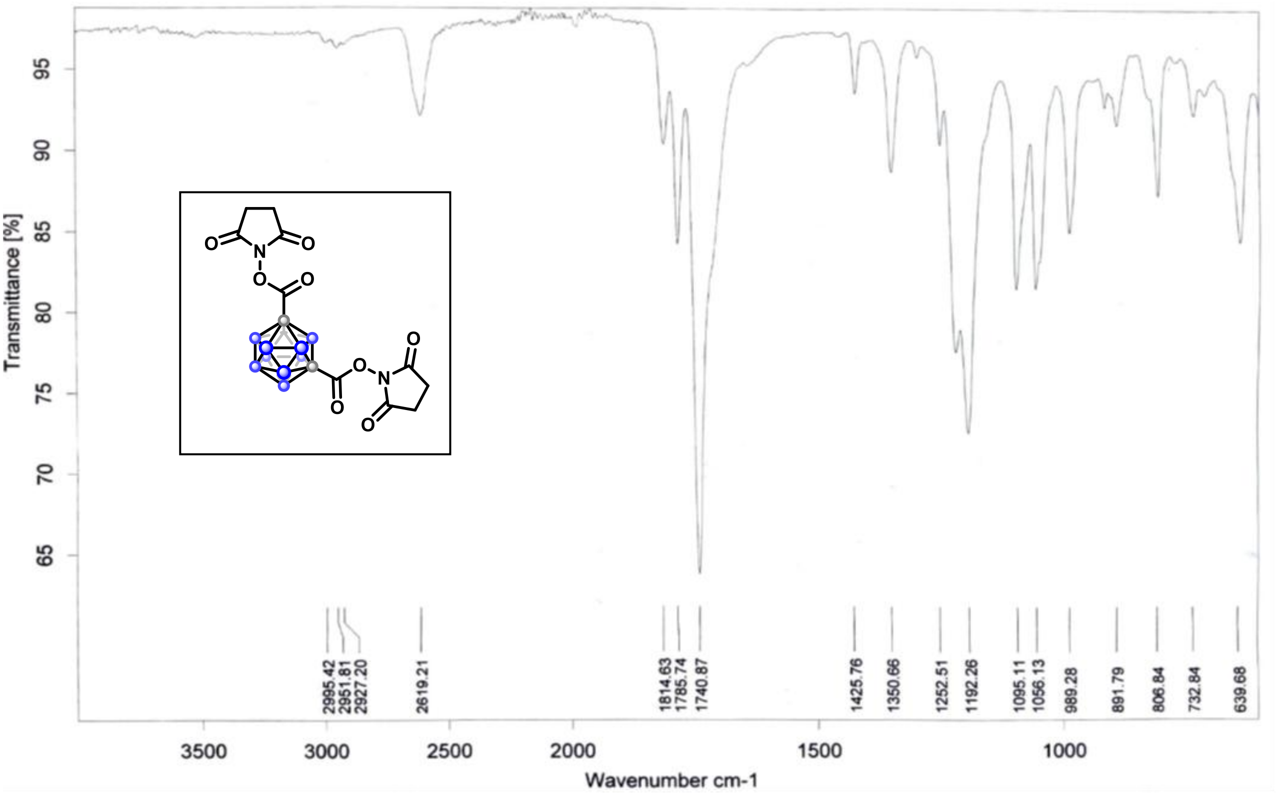
Transmittance [%]
Wavenumber cm-1
2995.42
2951.81
2927.20
2619.21
1814.63
1785.74
1740.87
1425.76
1350.66
1252.51
1192.26
1095.11
1056.13
989.28
891.79
806.84
732.84
639.68

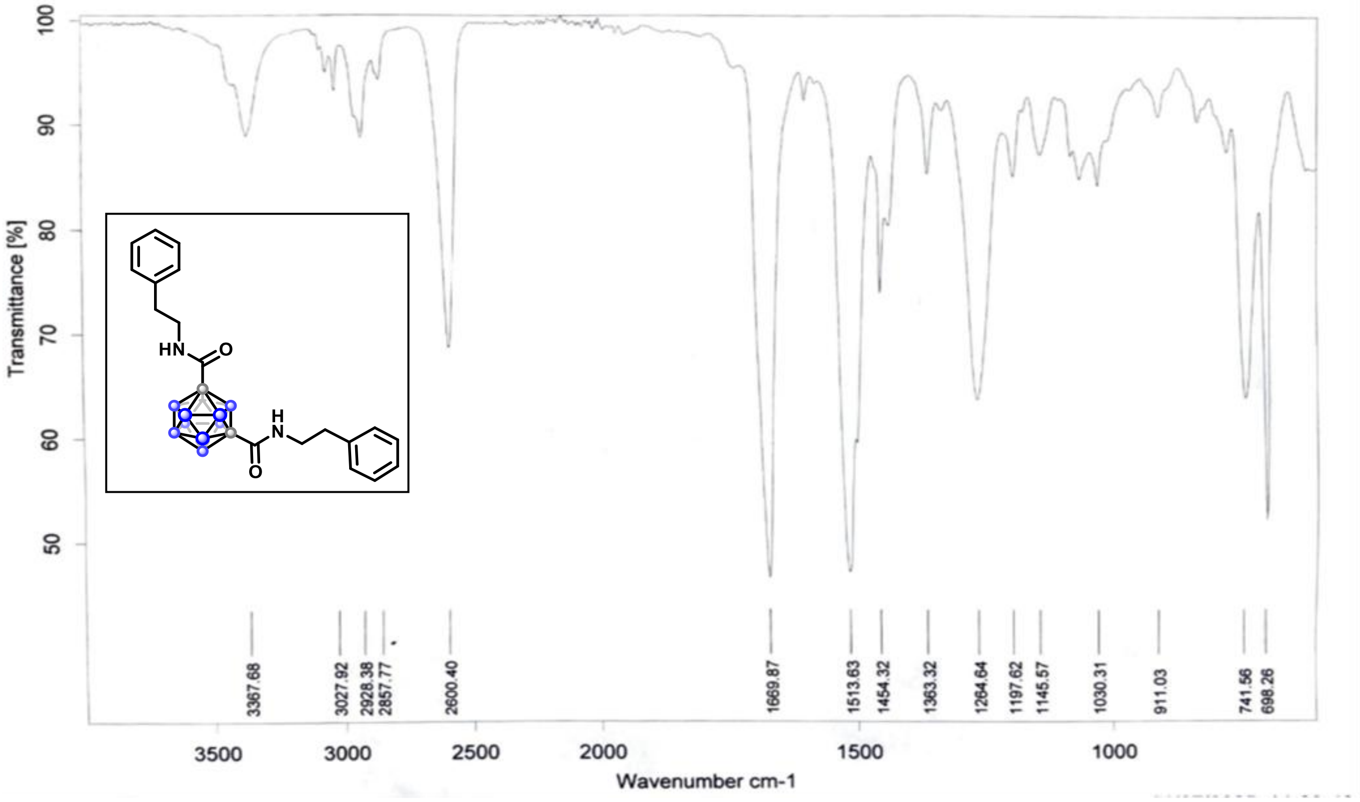
Transmittance [%]
Wavenumber cm-1
3367.68
3027.92
2928.38
2857.77
2600.40
1669.87
1513.63
1454.32
1363.32
1264.64
1197.62
1145.57
1030.31
911.03
741.56
698.26

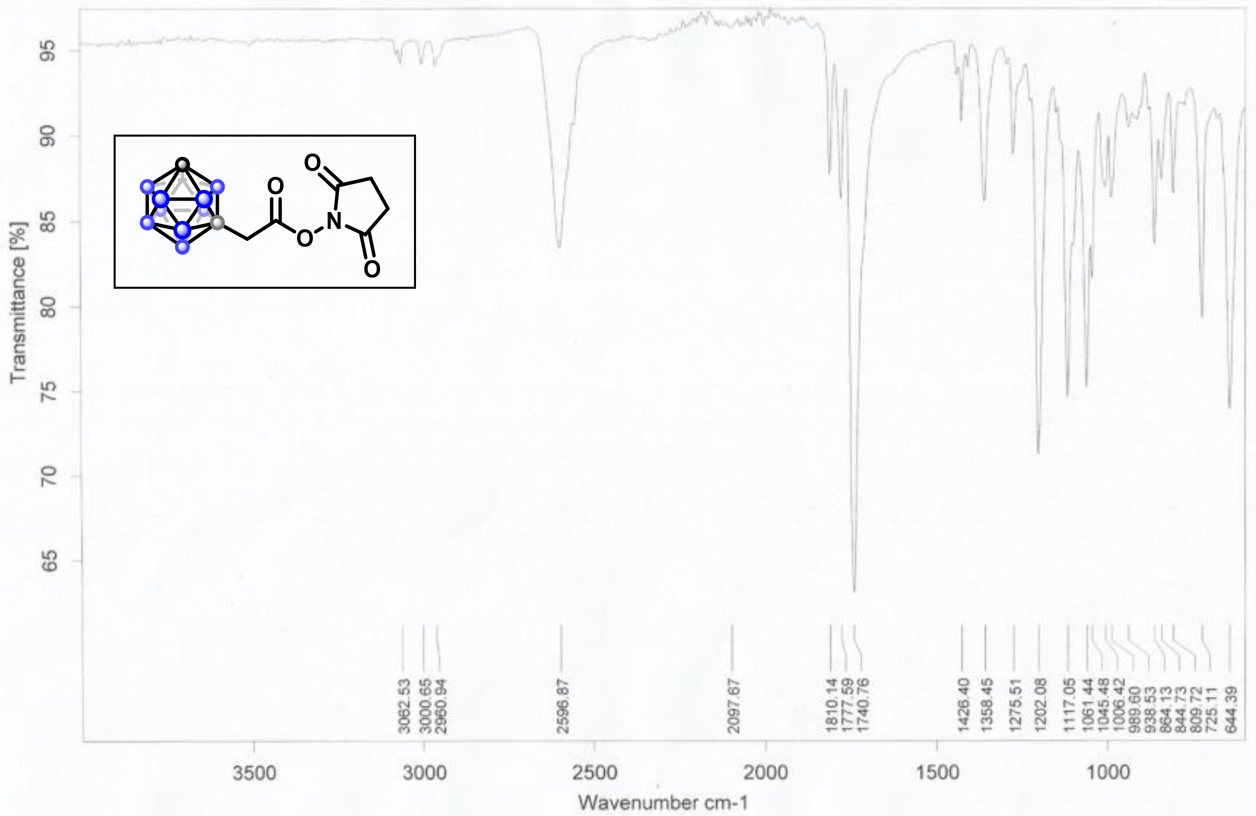
Transmittance [%]
95
90
85
80
75
70
65
3062.53
3000.65
2960.94
2596.87
2097.67
1810.14
1777.59
1740.76
1426.40
1358.45
1275.51
1202.08
1117.05
1061.44
1045.48
1006.42
989.60
938.53
864.13
844.73
809.72
725.11
644.39
3500
3000
2500
2000
1500
1000
Wavenumber cm-1

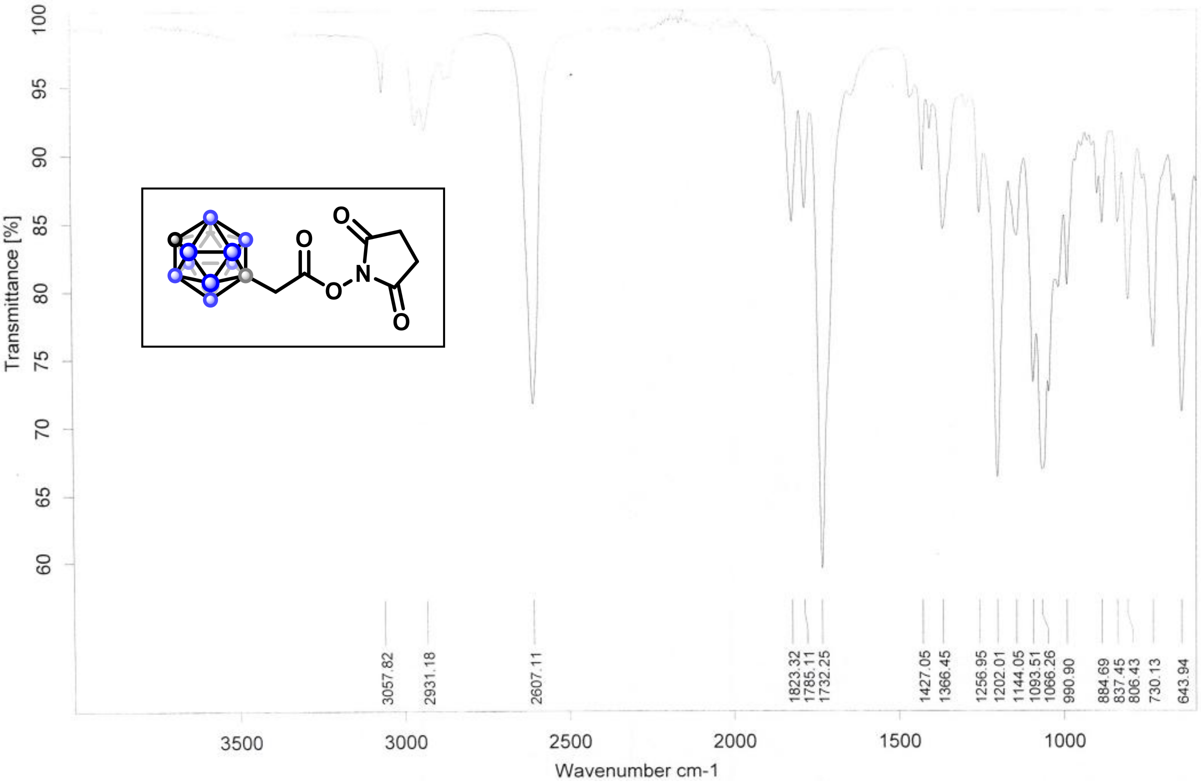
Transmittance [%]
100
95
90
85
80
75
70
65
60
3057.82
2931.18
2607.11
1823.32
1785.11
1732.25
1427.05
1366.45
1256.95
1202.01
1144.05
1093.51
1066.26
990.90
884.69
837.45
806.43
730.13
643.94
3500
3000
2500
2000
1500
1000
Wavenumber cm-1

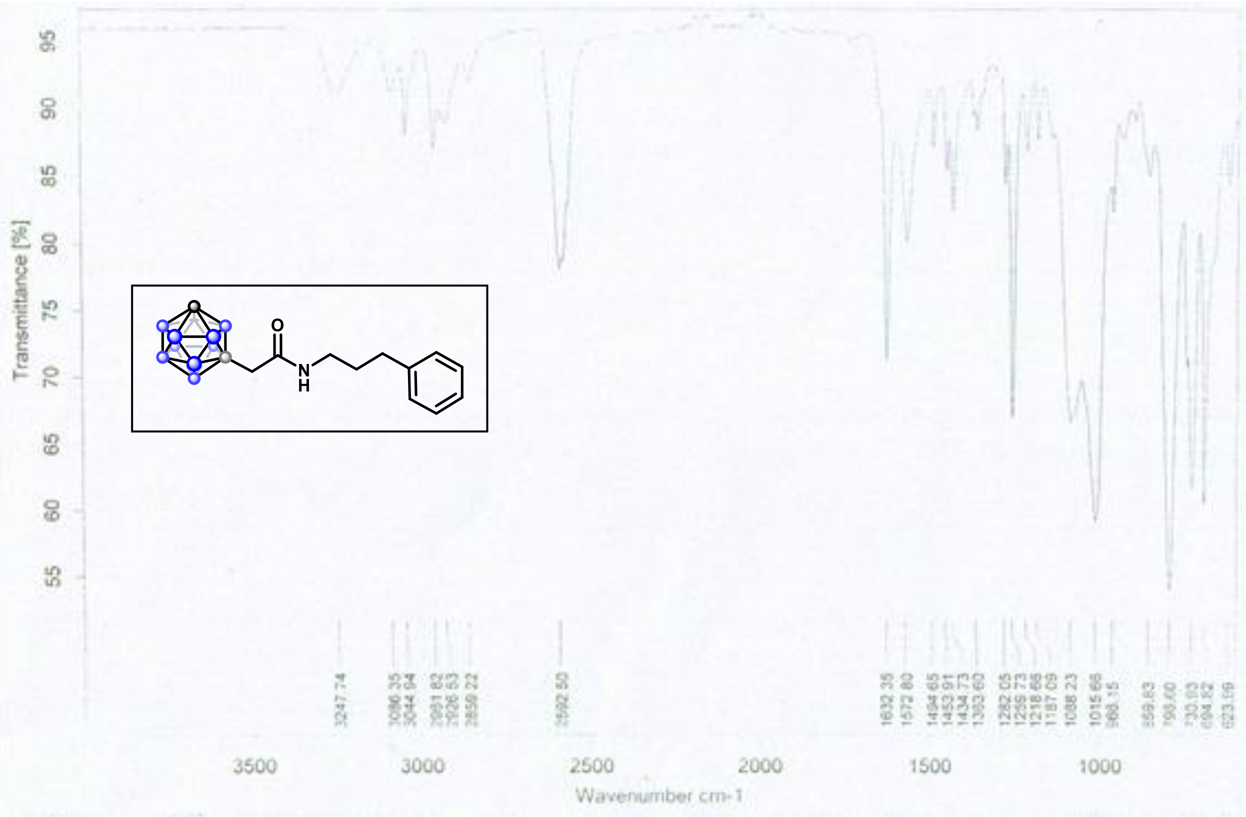

Transmittance [%]
95
90
85
80
75
70
65
60
55
O
N
H
3500
3000
2500
2000
1500
1000
Wavenumber cm-1

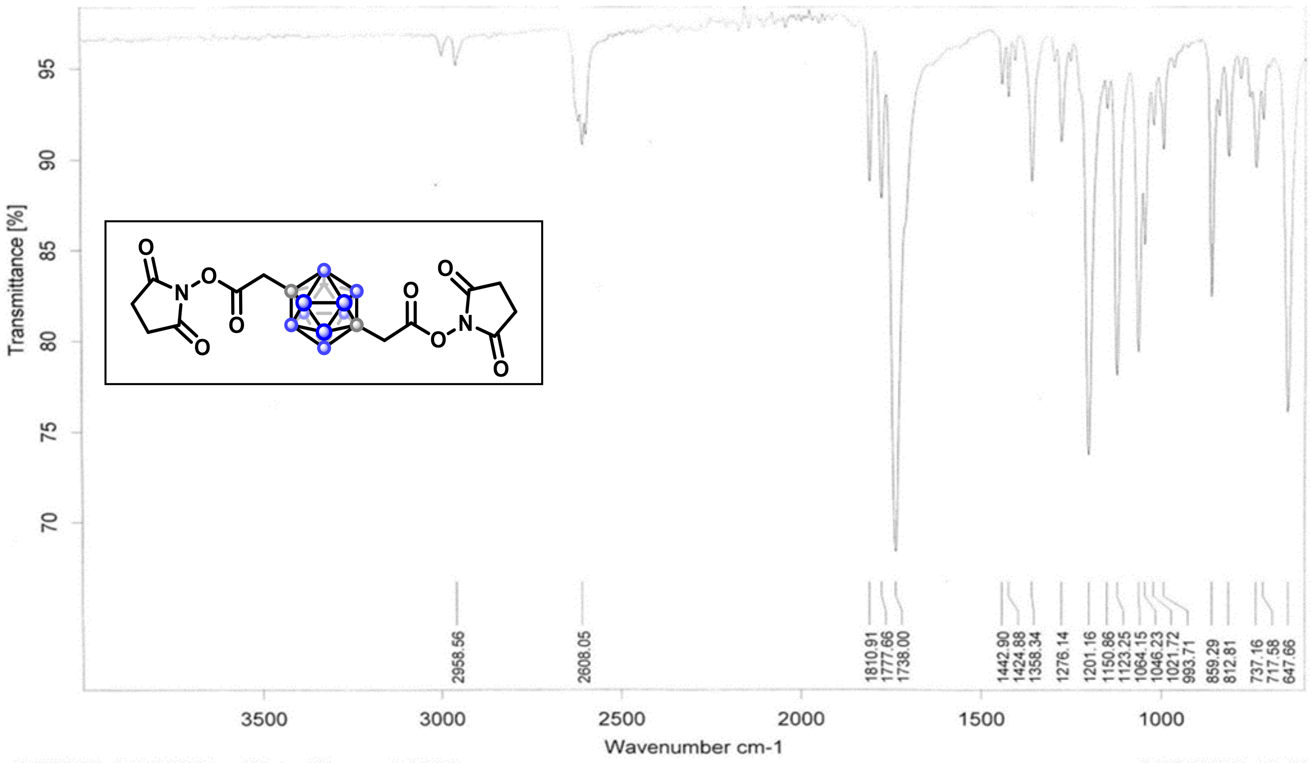

Transmittance [%]
95
90
85
80
75
70
O
N
O
O
O
O
O
O
N
O
2958.56
2608.05
1810.91
1777.66
1738.00
1442.90
1424.88
1358.34
1276.14
1201.16
1150.86
1123.25
1064.15
1046.23
1021.72
993.71
859.29
812.81
737.16
717.58
647.66
3500
3000
2500
2000
1500
1000
Wavenumber cm-1

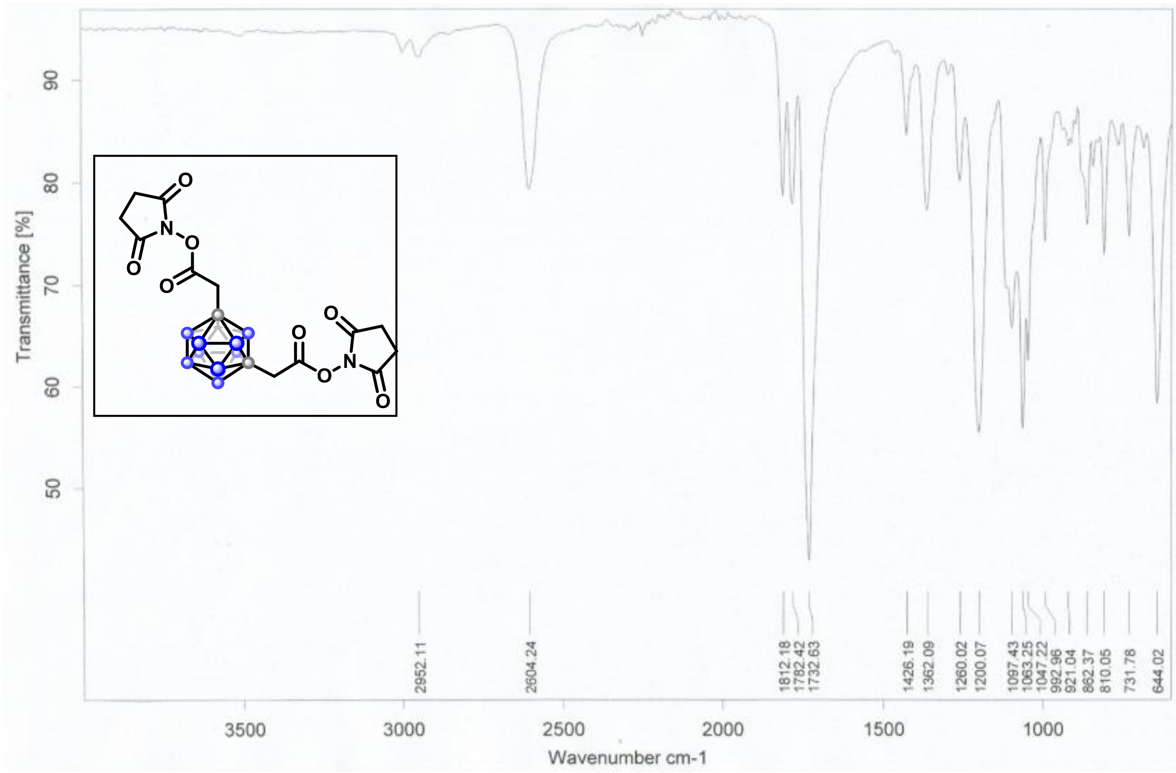

Transmittance [%]
Wavenumber cm-1
2952.11
2604.24
1812.18
1782.42
1732.63
1426.19
1362.09
1260.02
1200.07
1097.43
1063.25
1047.22
992.96
921.04
862.37
810.05
731.78
644.02
O
N

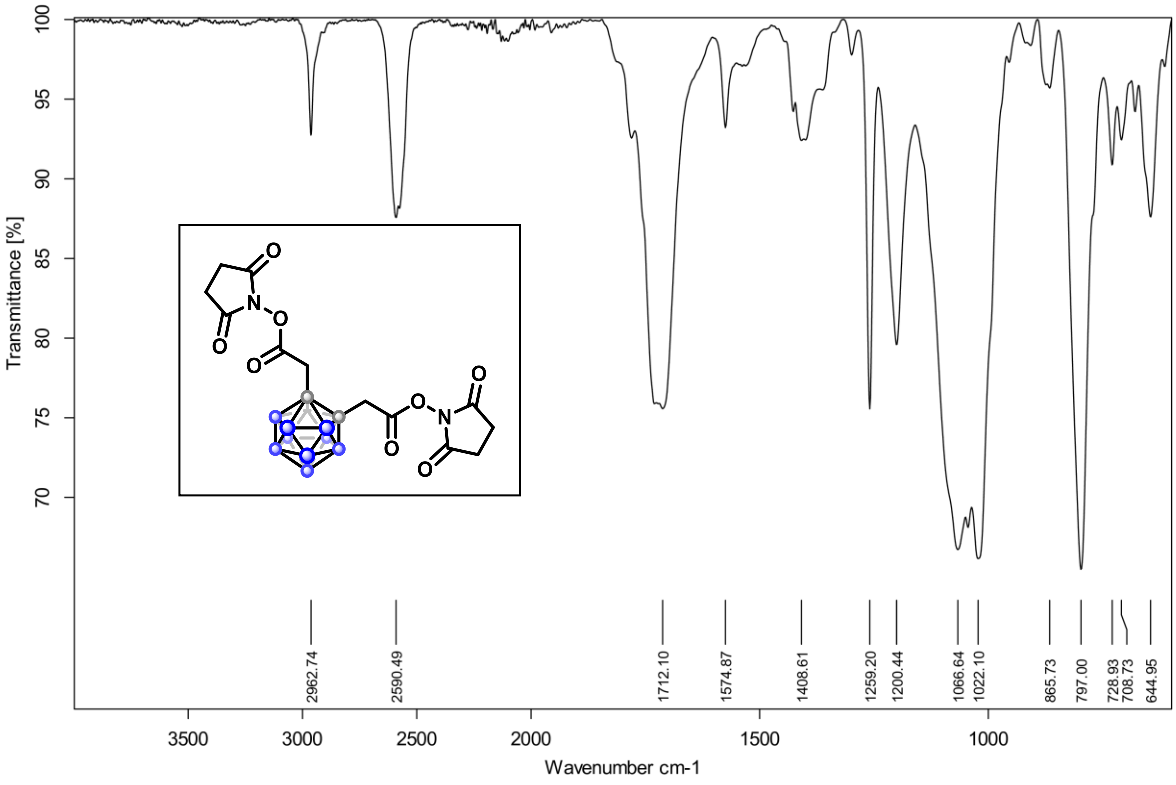

Transmittance [%]
Wavenumber cm-1
2962.74
2590.49
1712.10
1574.87
1408.61
1259.20
1200.44
1066.64
1022.10
865.73
797.00
728.93
708.73
644.95
O
N

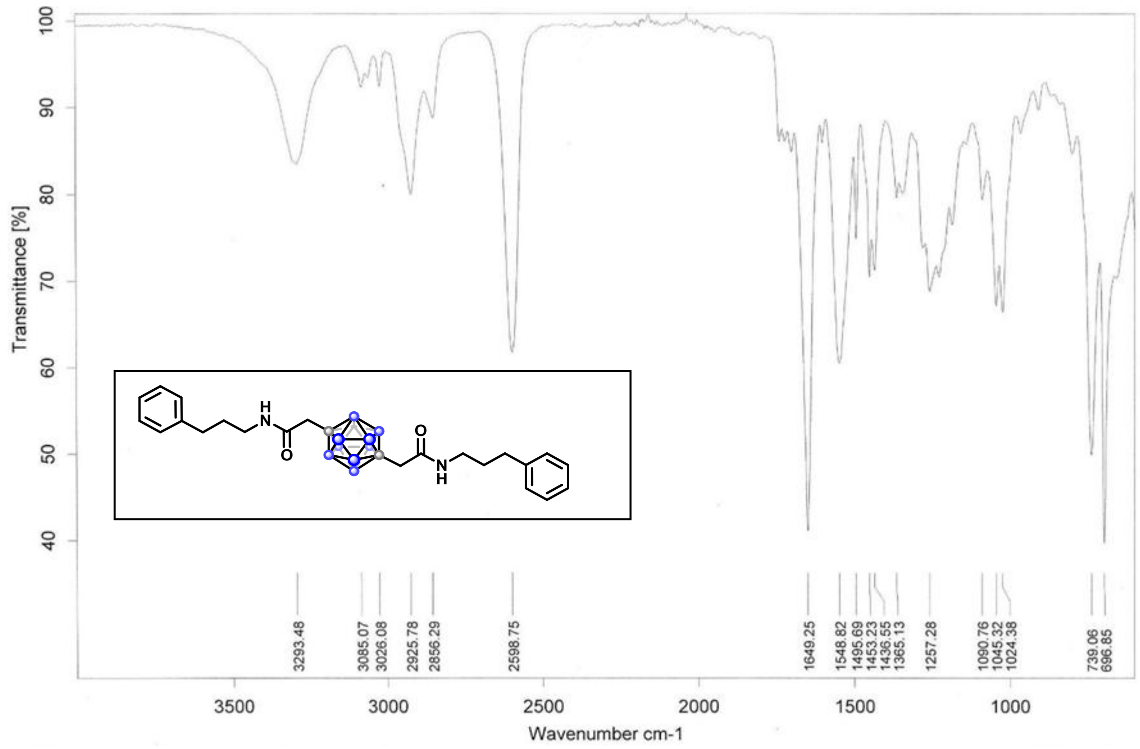
Transmittance [%]
Wavenumber cm-1
3293.48
3085.07
3026.08
2925.78
2856.29
2598.75
1649.25
1548.82
1495.69
1453.23
1436.55
1365.13
1257.28
1090.76
1045.32
1024.38
739.06
696.85

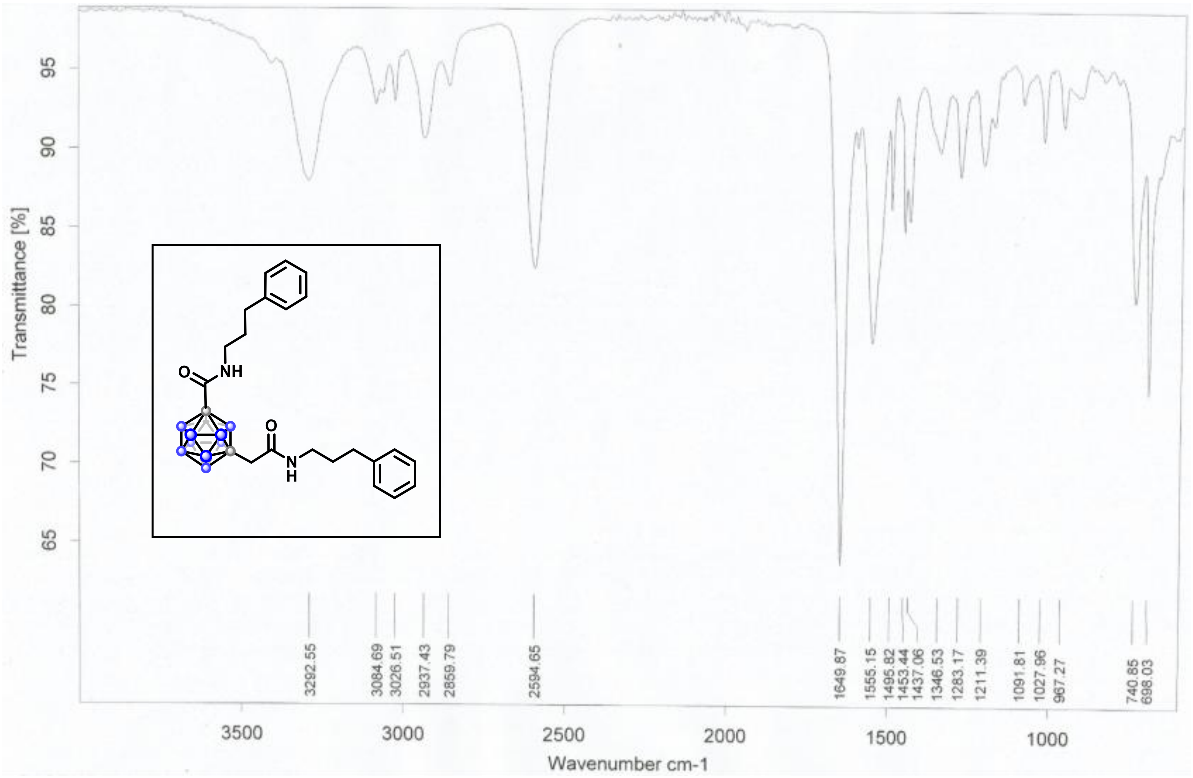
Transmittance [%]
Wavenumber cm-1
3292.55
3084.69
3026.51
2937.43
2859.79
2594.65
1649.87
1555.15
1495.82
1453.44
1437.06
1346.53
1283.17
1211.39
1091.81
1027.96
967.27
740.85
698.03

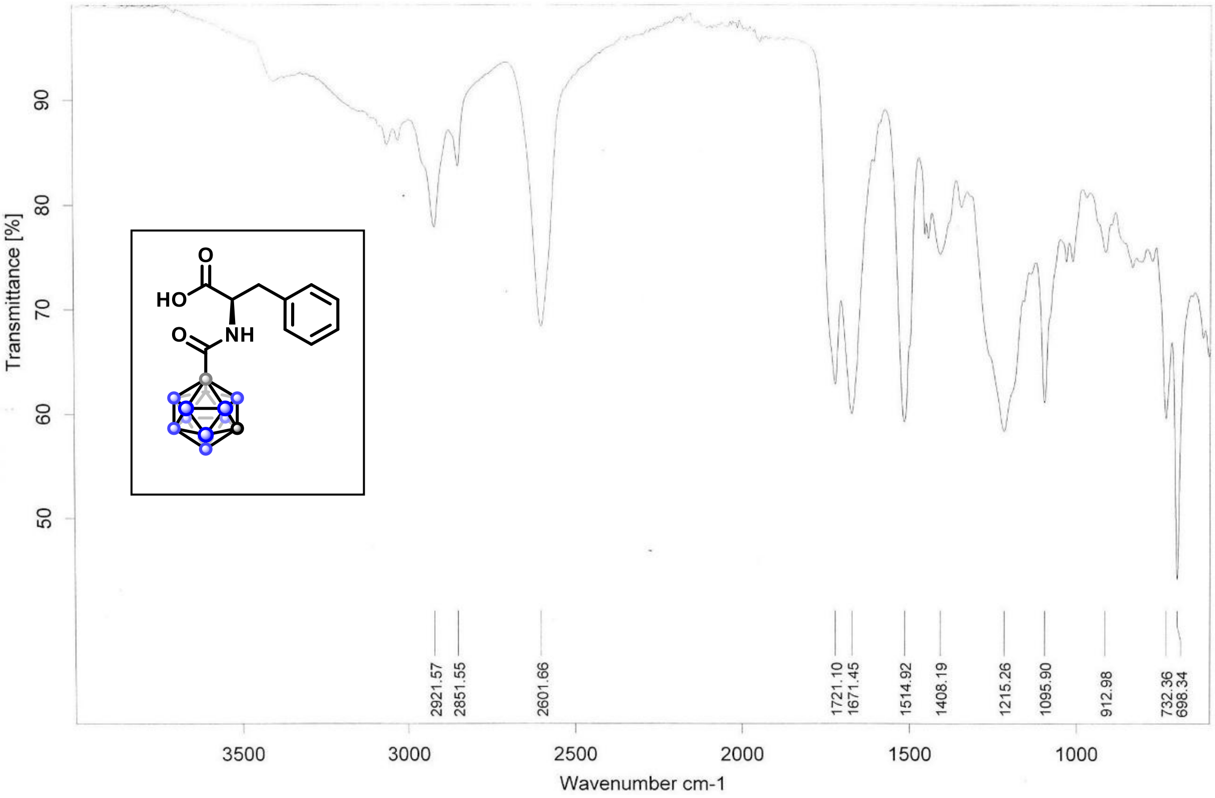
Transmittance [%]
90
80
70
60
50
HO
O
NH
2921.57
2851.55
2601.66
1721.10
1671.45
1514.92
1408.19
1215.26
1095.90
912.98
732.36
698.34
3500
3000
2500
2000
1500
1000
Wavenumber cm-1

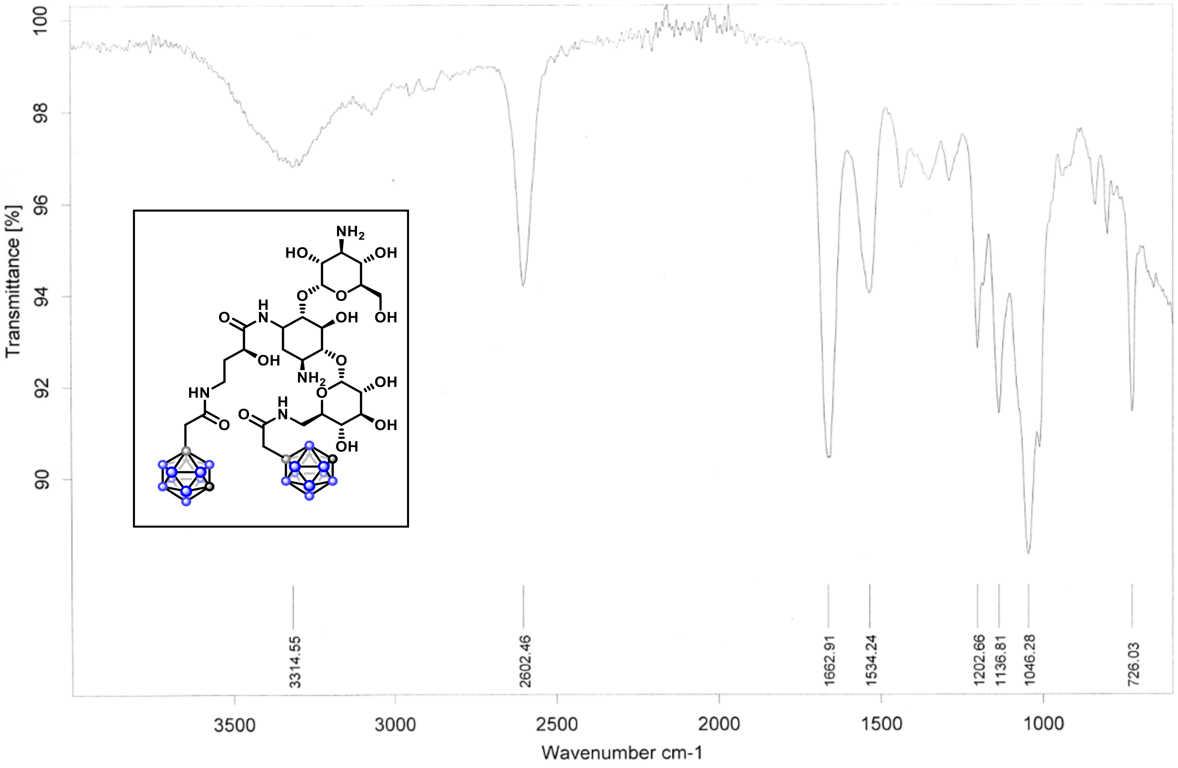
Transmittance [%]
100
98
96
94
92
90
$NH_2$
HO
OH
OH
OH
$NH_2$
OH
OH
OH
HN
O
H
N
3314.55
2602.46
1662.91
1534.24
1202.66
1136.81
1046.28
726.03
3500
3000
2500
2000
1500
1000
Wavenumber cm-1

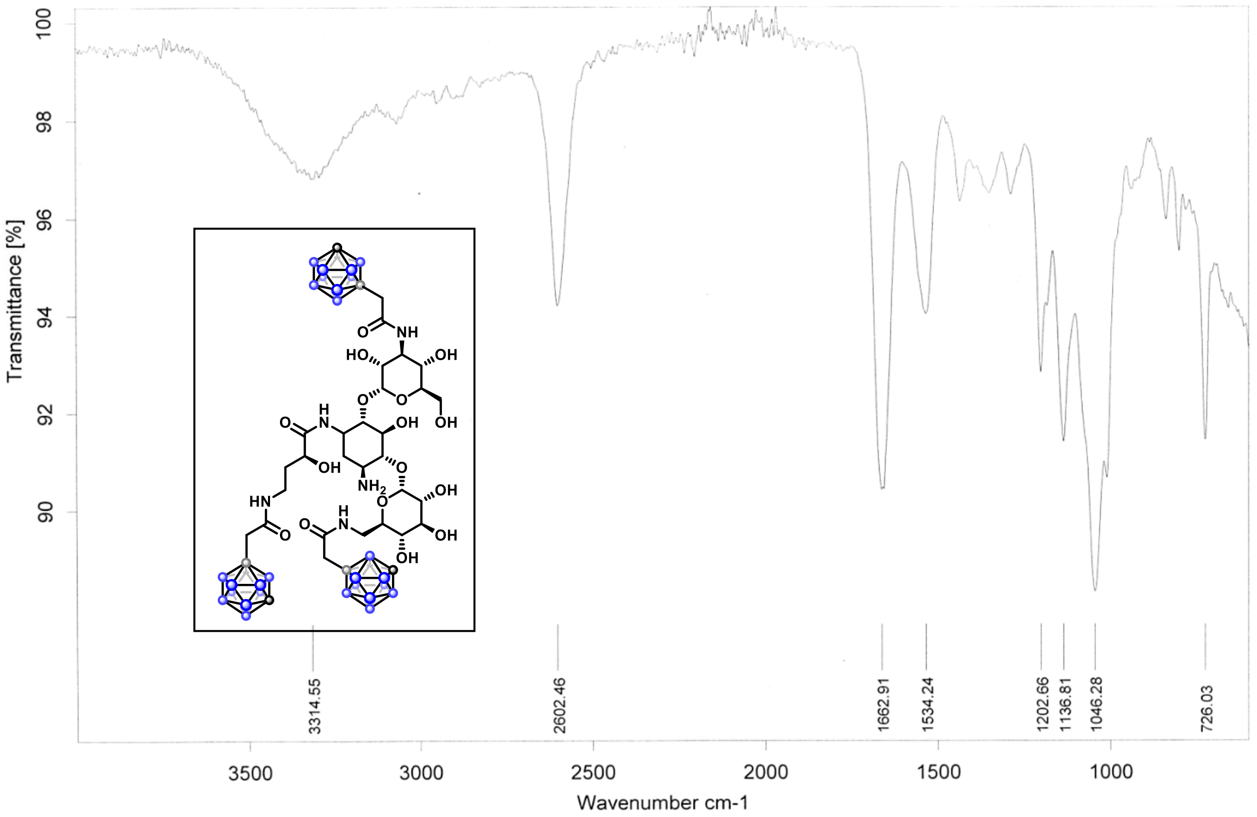

Transmittance [%]
Wavenumber cm-1
3314.55
2602.46
1662.91
1534.24
1202.66
1136.81
1046.28
726.03

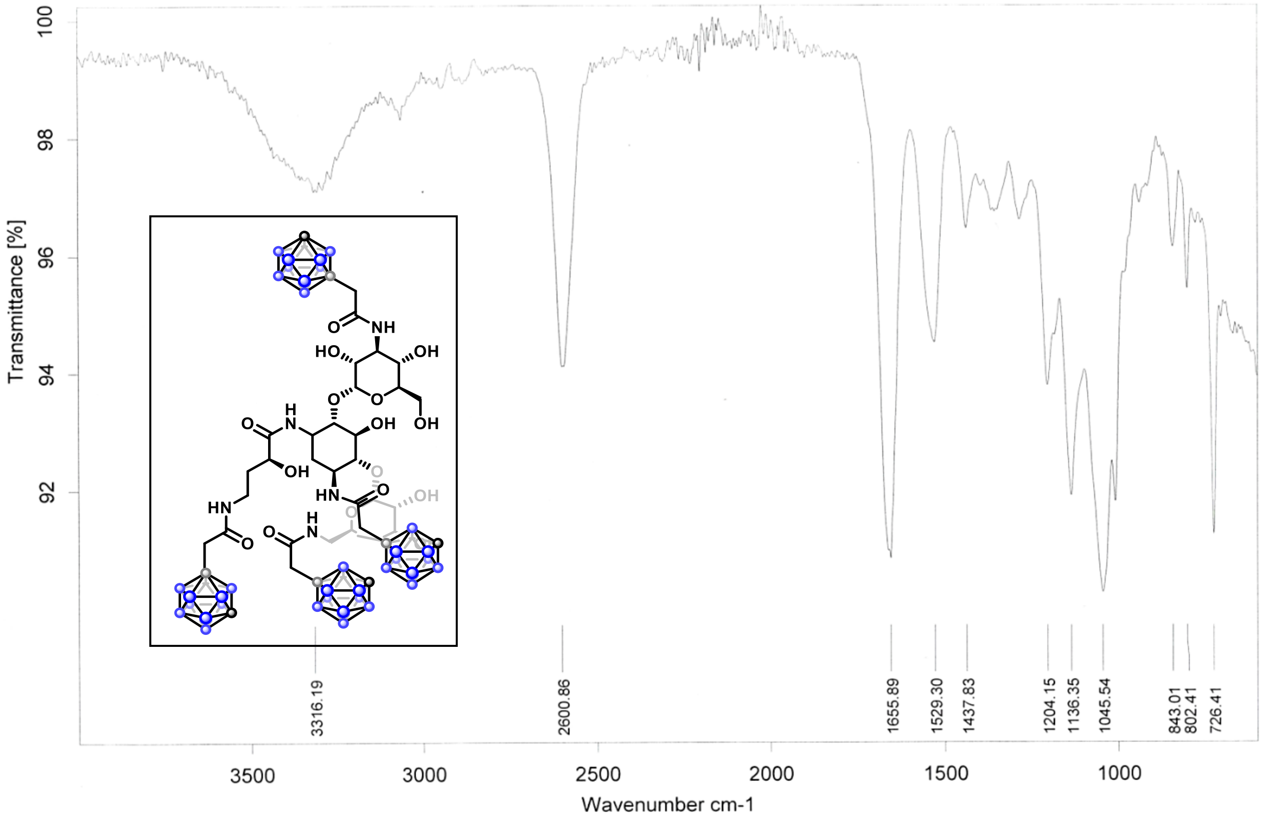

Transmittance [%]
Wavenumber cm-1
3316.19
2600.86
1655.89
1529.30
1437.83
1204.15
1136.35
1045.54
843.01
802.41
726.41

#### Copies of NMR spectra

$^{1}H$ NMR (500 MHz, $CDCl_3$)

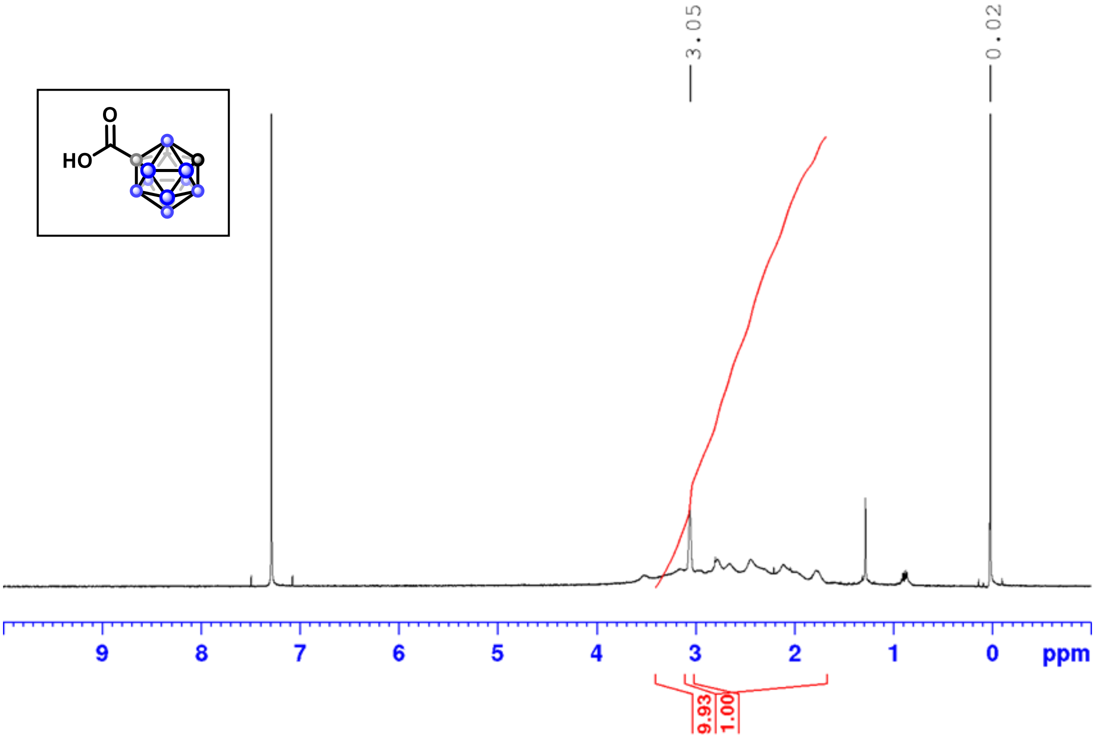

$^{1}H$ NMR (500 MHz, $CDCl_3$)

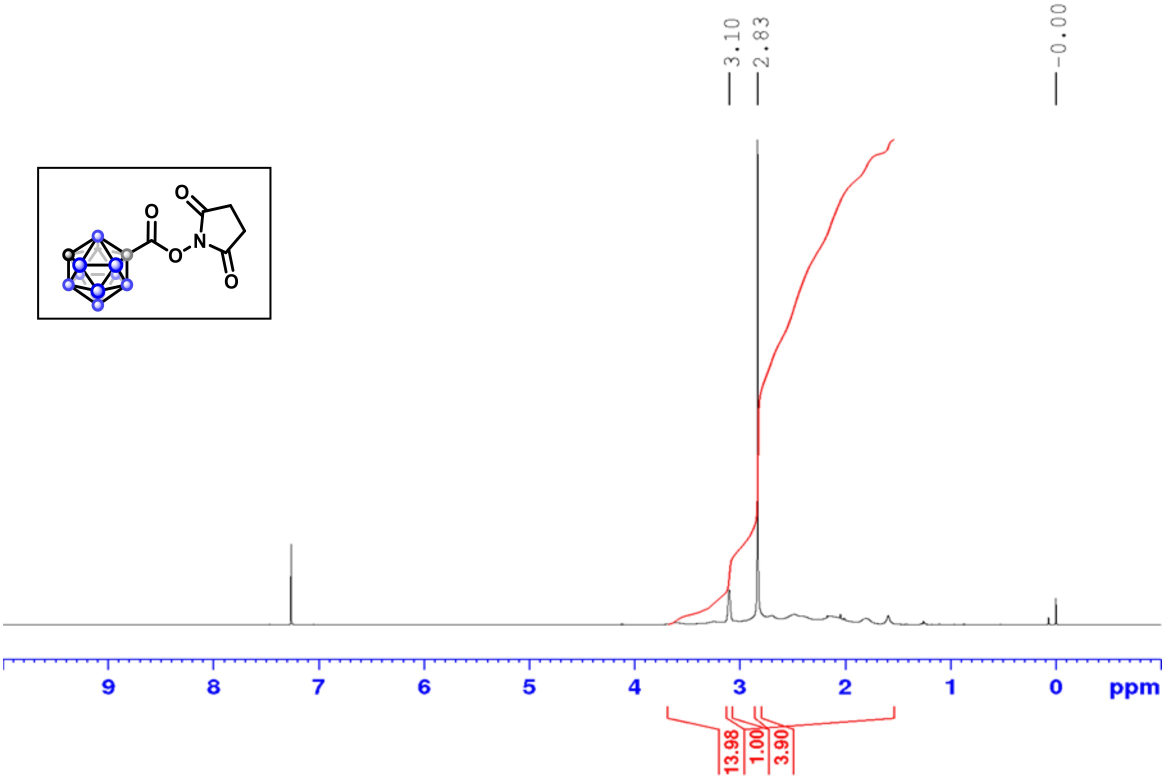

$^{13}C$ NMR (126 MHz, $CDCl_3$)

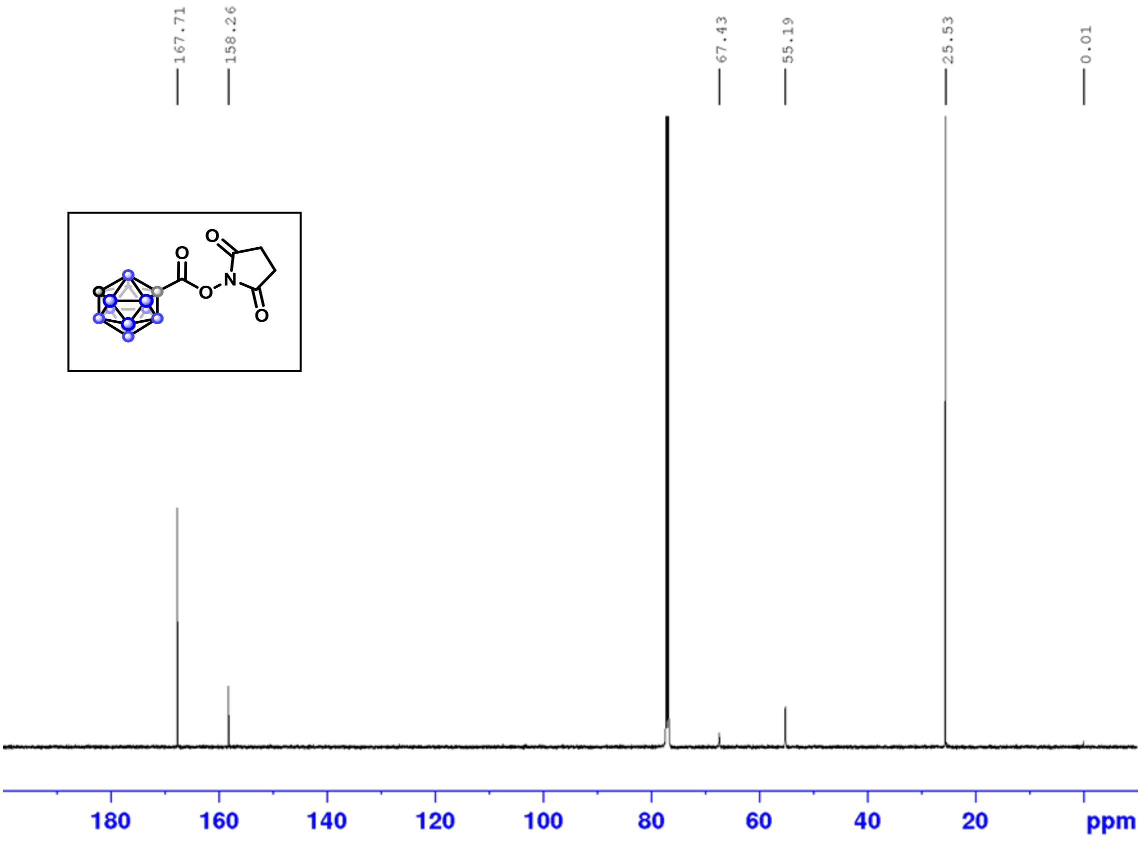

$^{11}B$ NMR (160 MHz, $CDCl_3$)

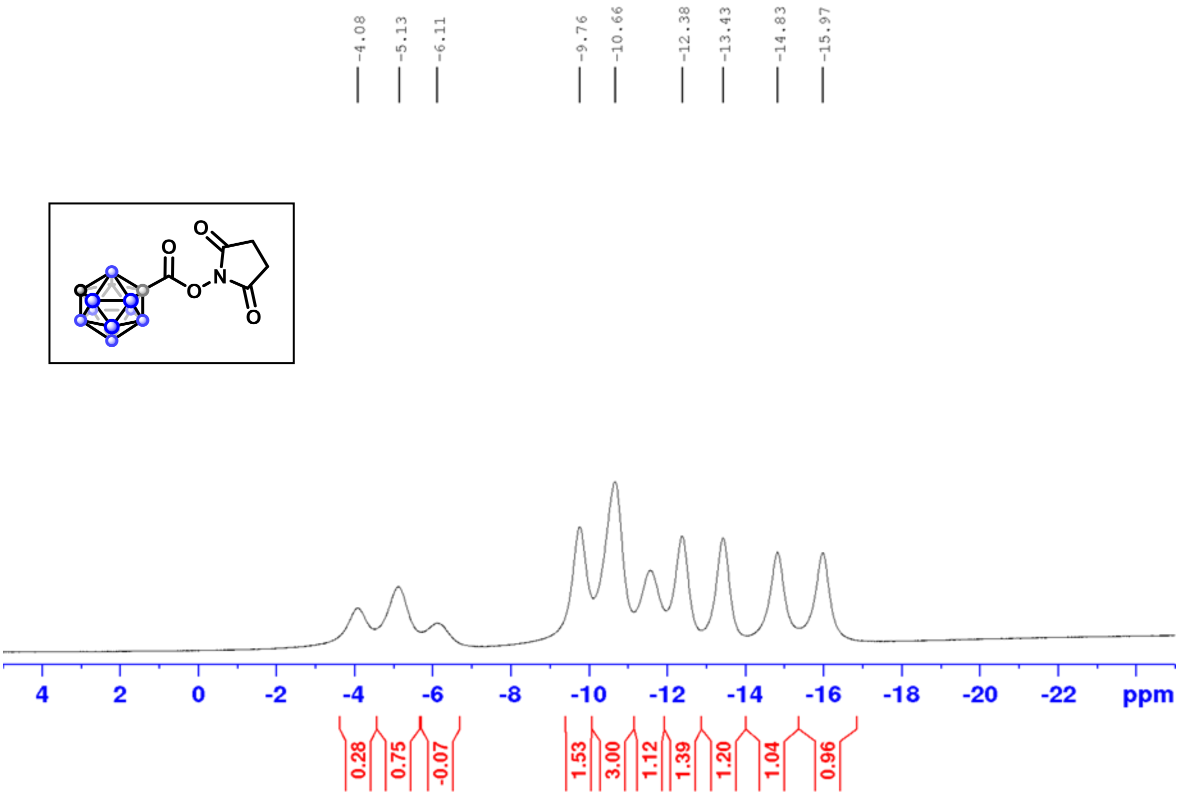

$^1H$-$^1H$ COSY

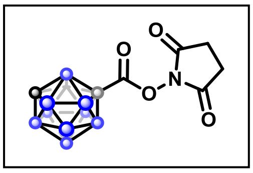

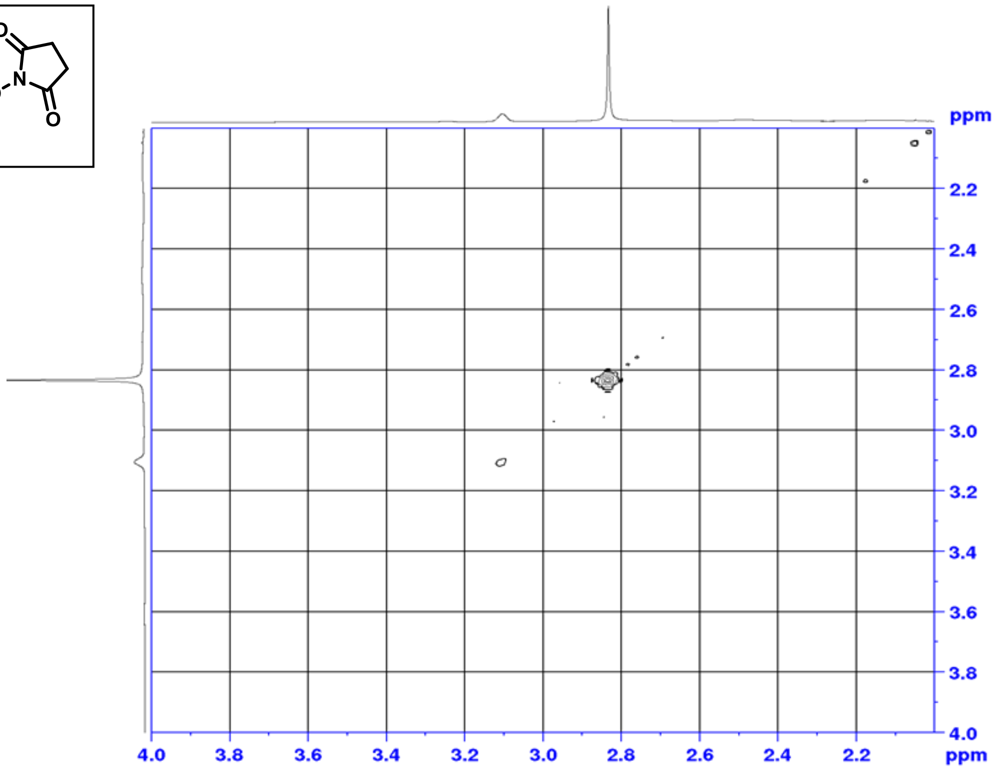

$^1H$-$^{13}C$ HSQC

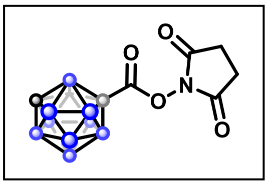

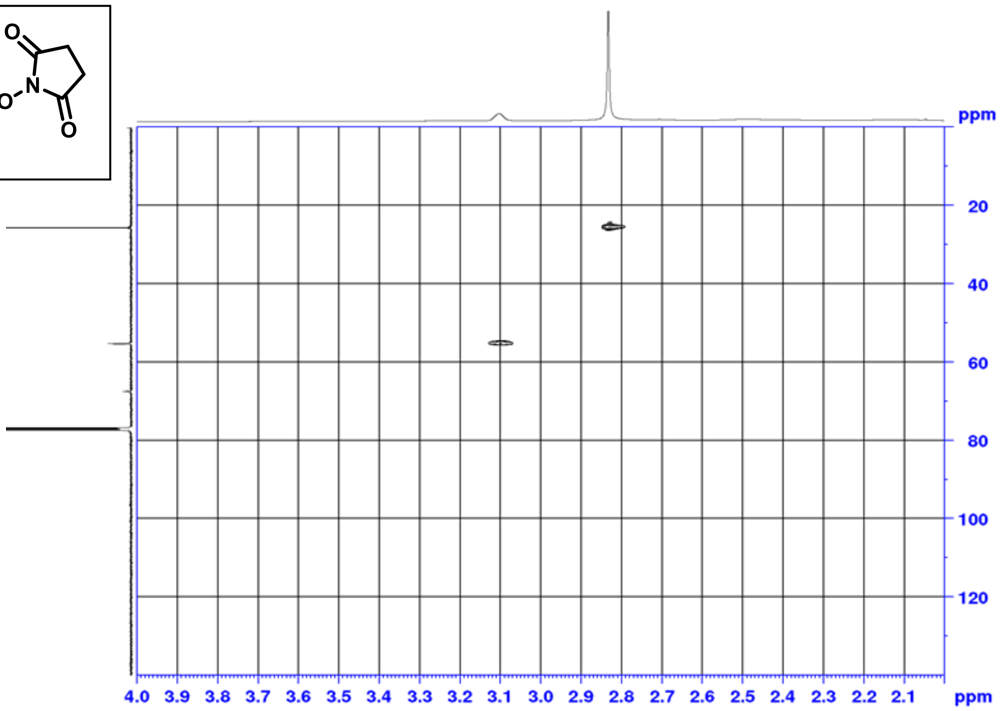

$^{1}H$-$^{13}C$ HMBC

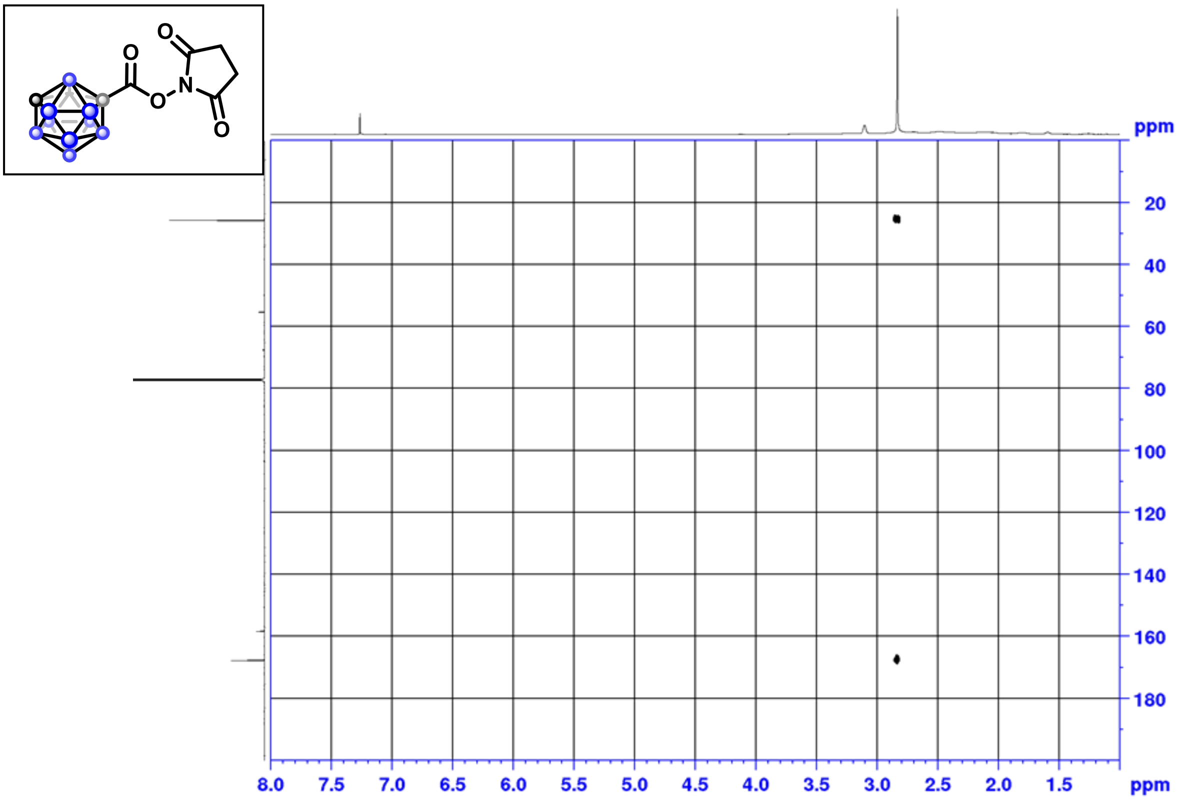

$^{1}H$ NMR (500 MHz, $CDCl_3$)

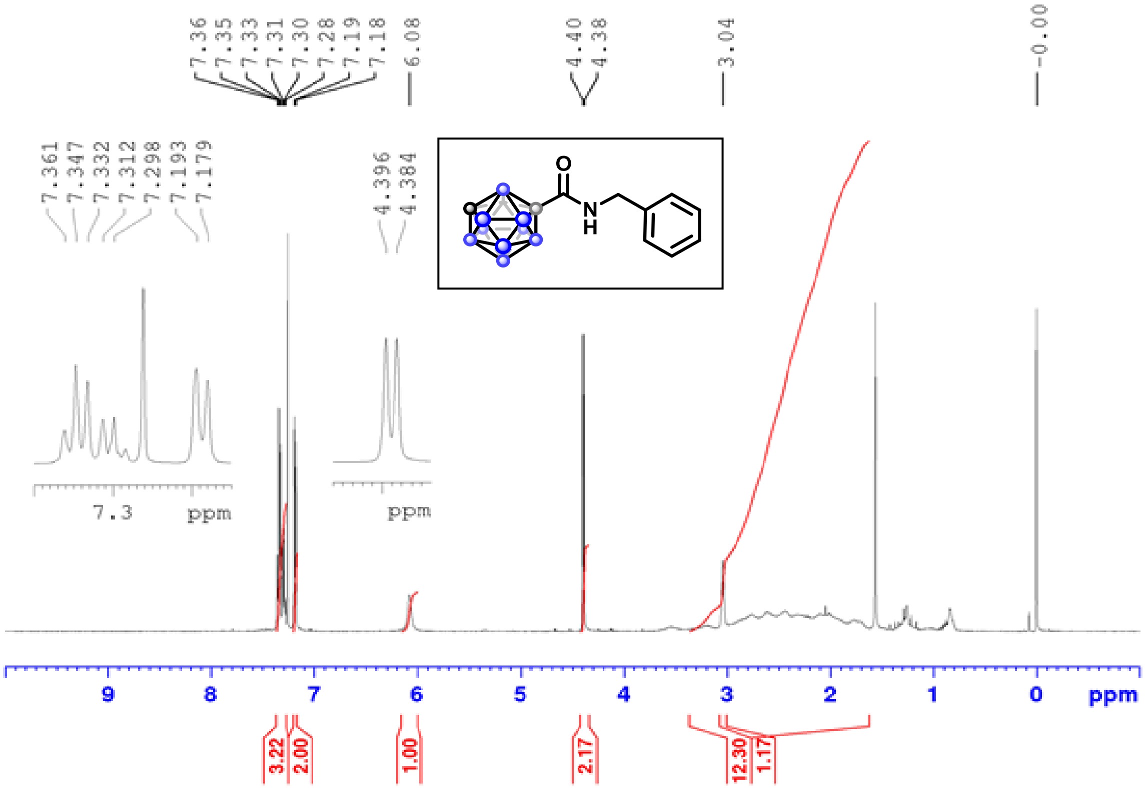

$^{13}C$ NMR (126 MHz, $CDCl_3$)

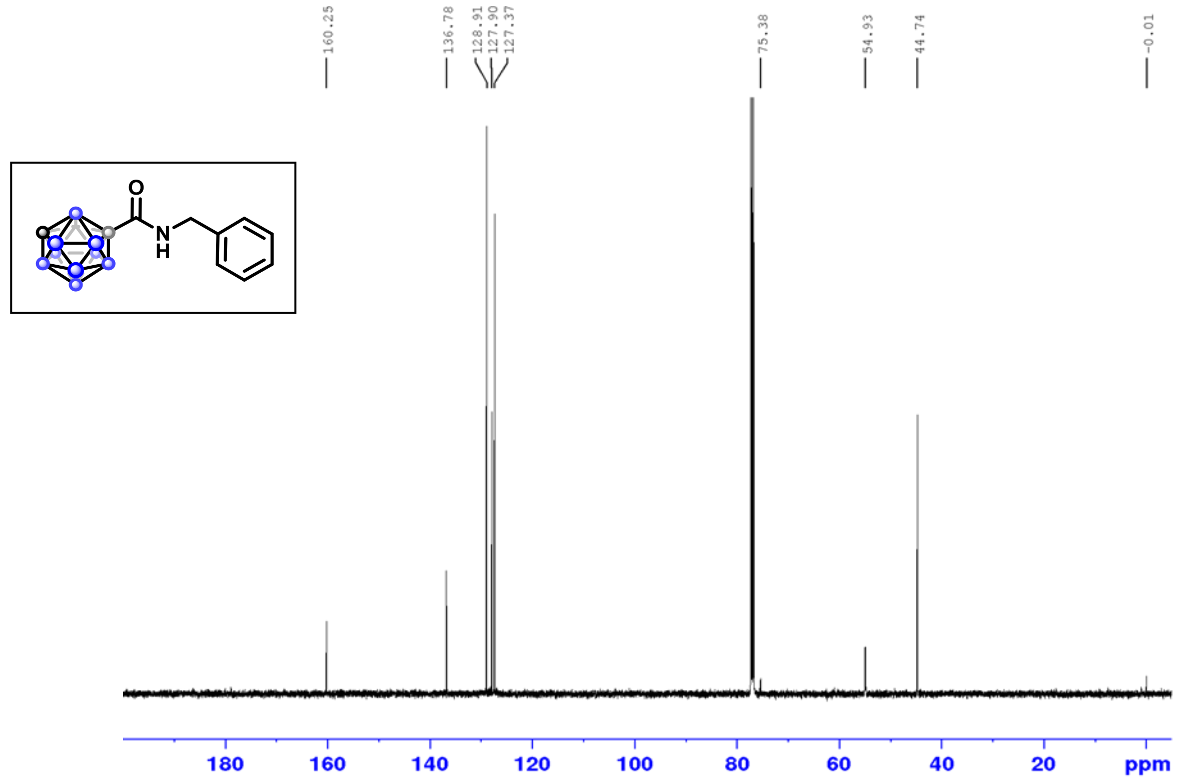

$^{11}B$ NMR (160 MHz, $CDCl_3$)

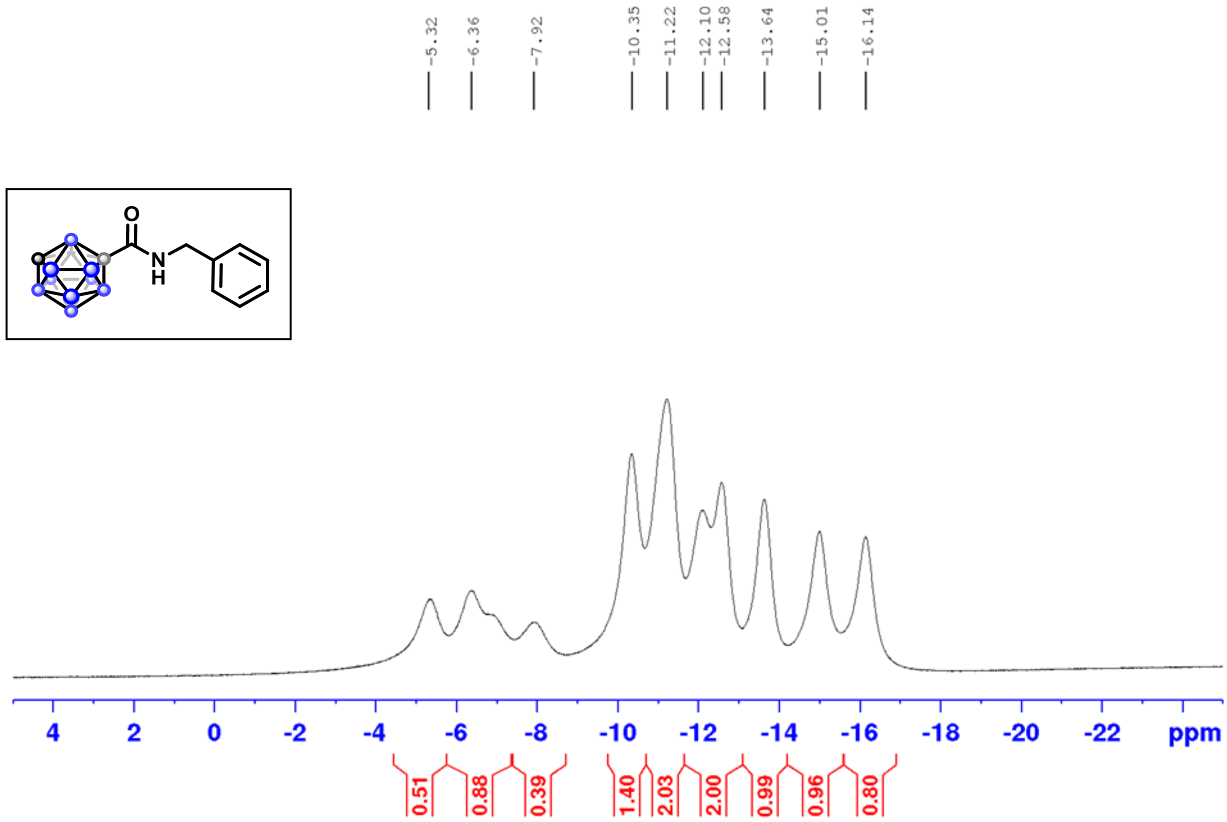

$^{1}H$-$^{1}H$ COSY

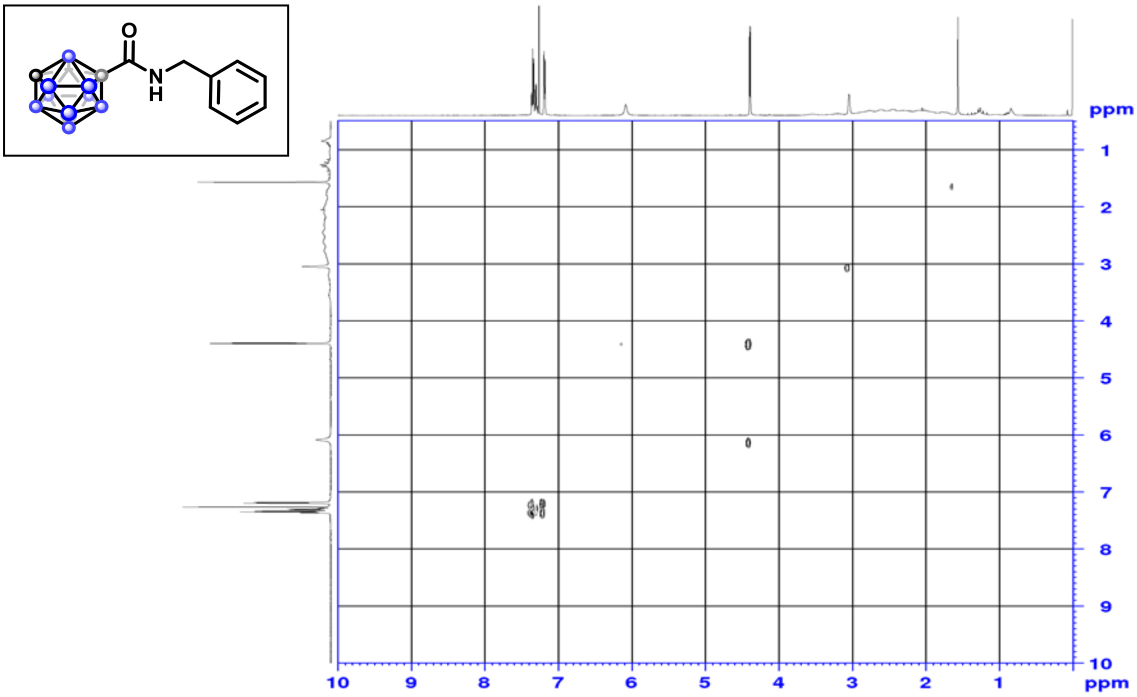

$^{1}H$-$^{13}C$ HSQC

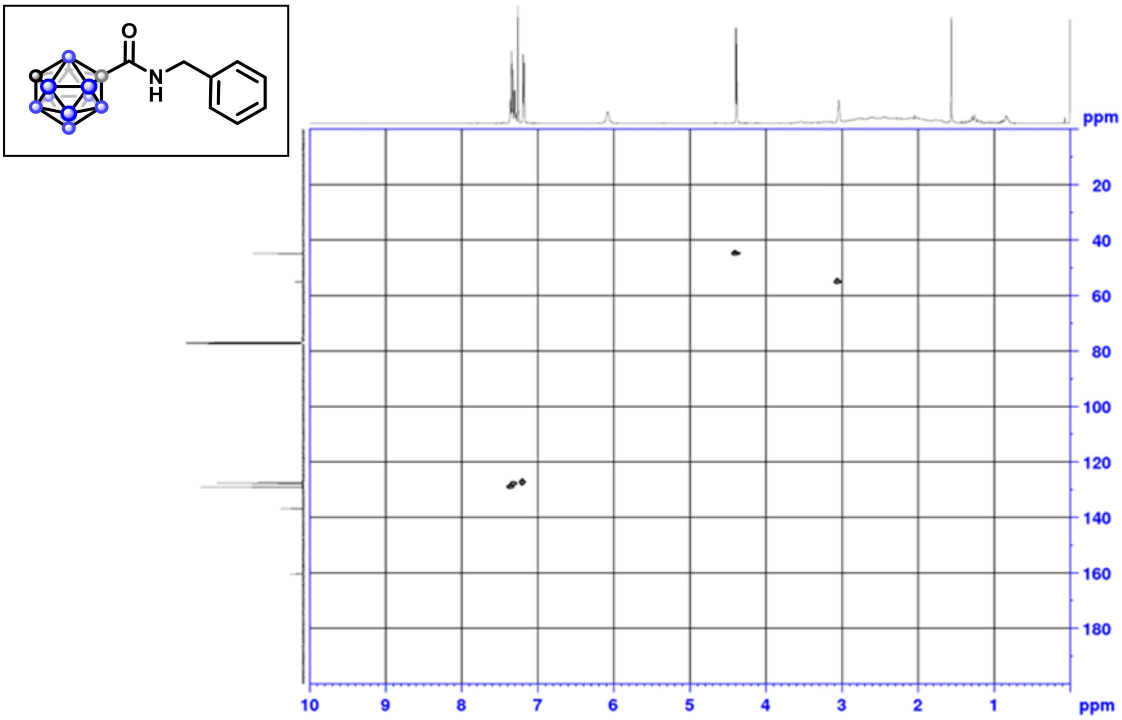

$^{1}$H NMR (500 MHz, $CDCl_3$)

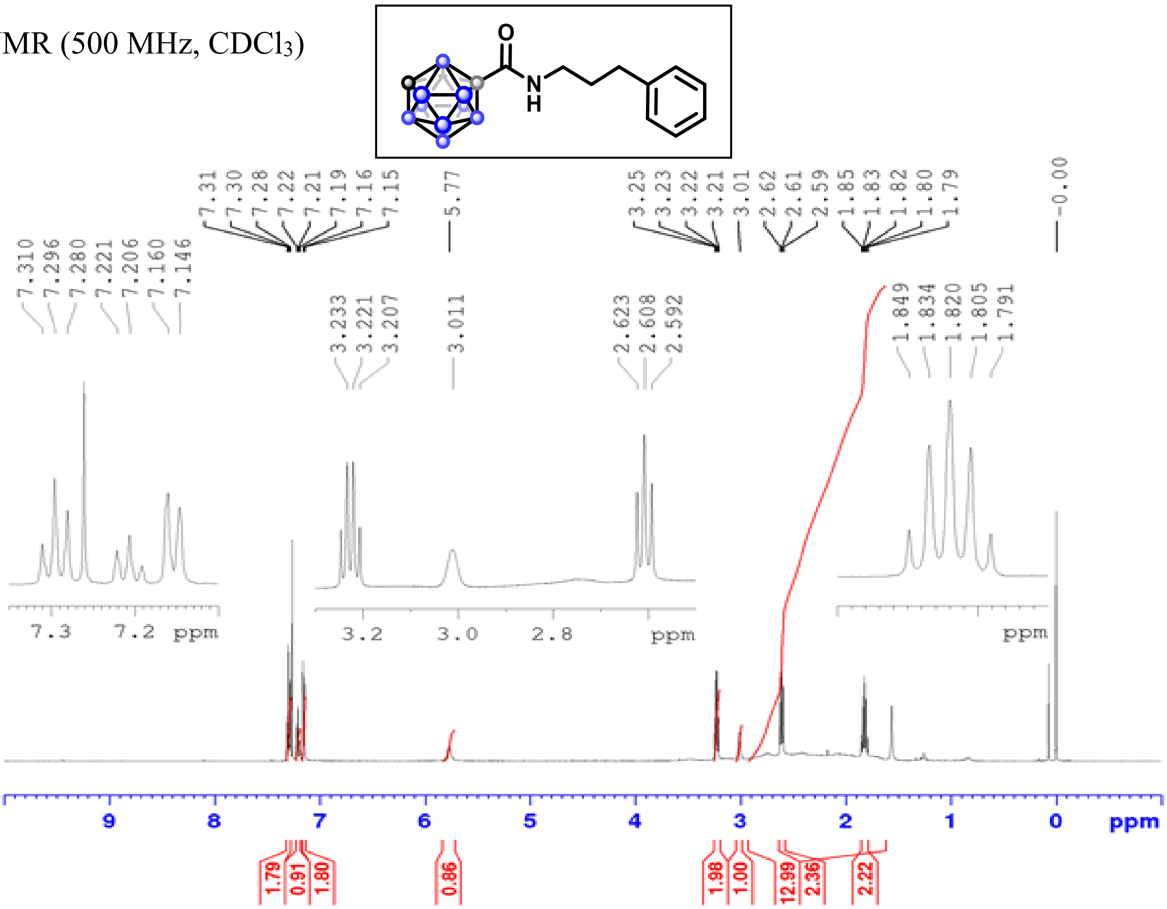

$^{13}$C NMR (126 MHz, $CDCl_3$)

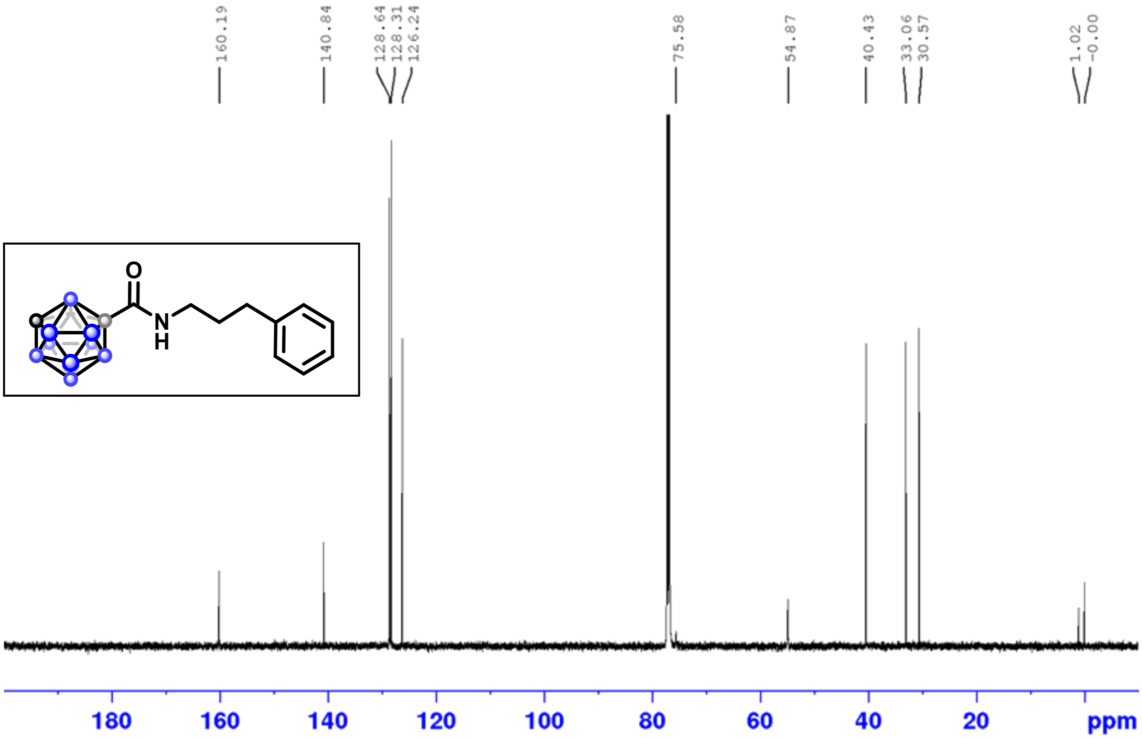

$^{11}B$ NMR (160 MHz, $CDCl_3$)

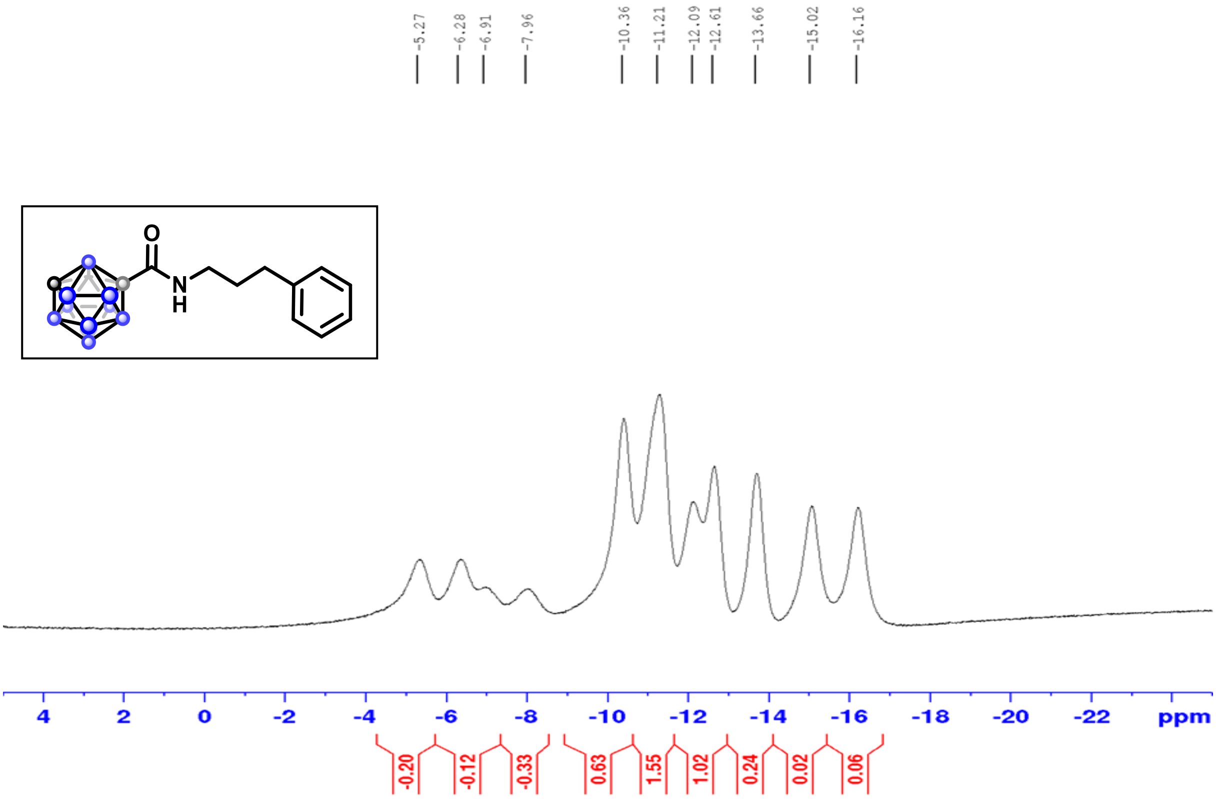

$^1H$-$^1H$ COSY

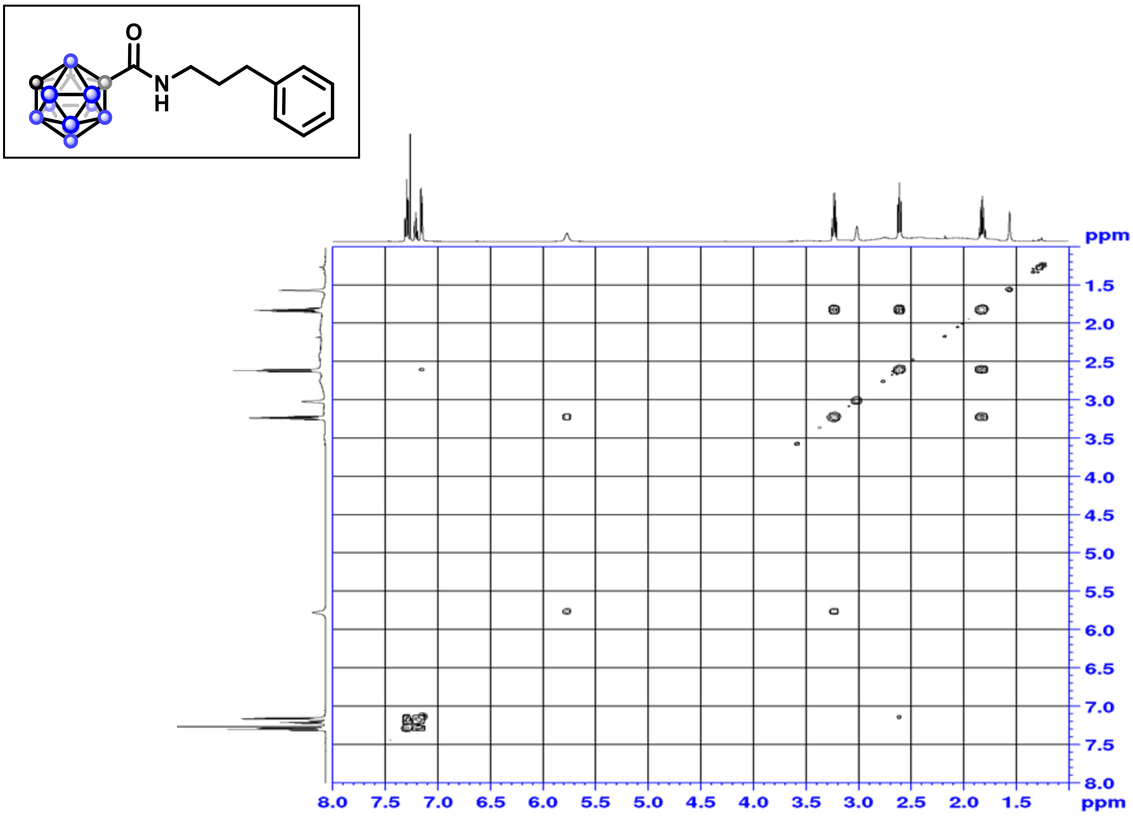

$^{1}H$-$^{13}C$ HSQC

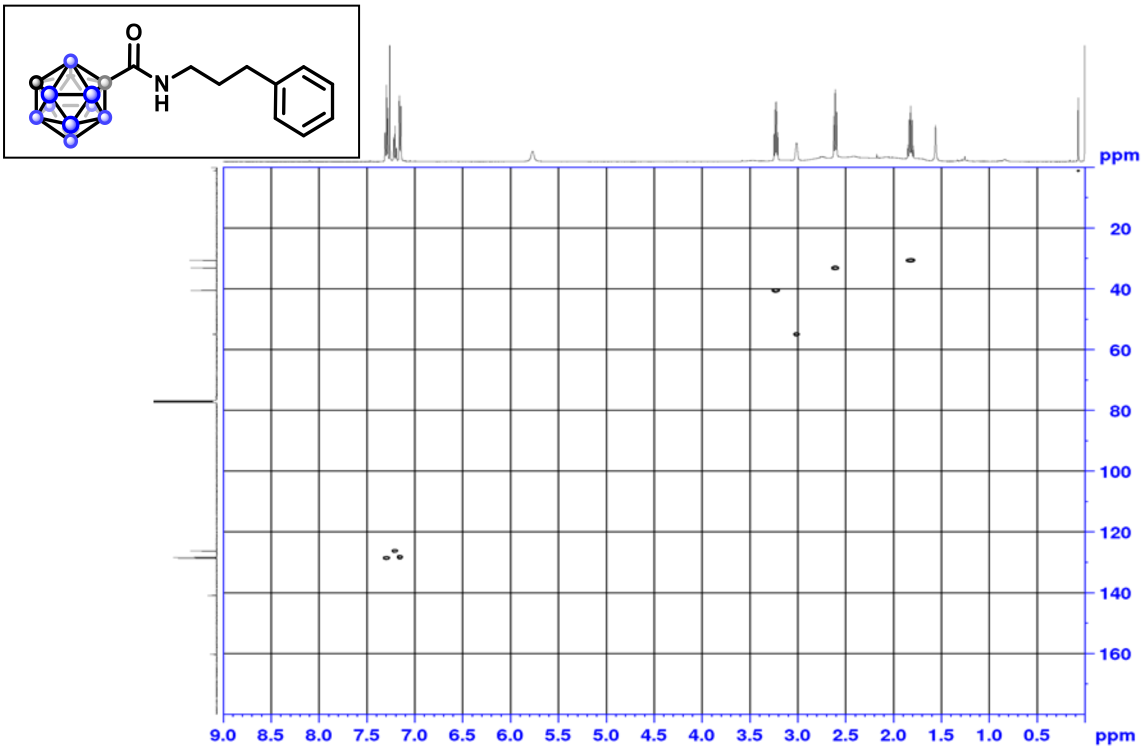

$^{1}H$-$^{13}C$ HMBC

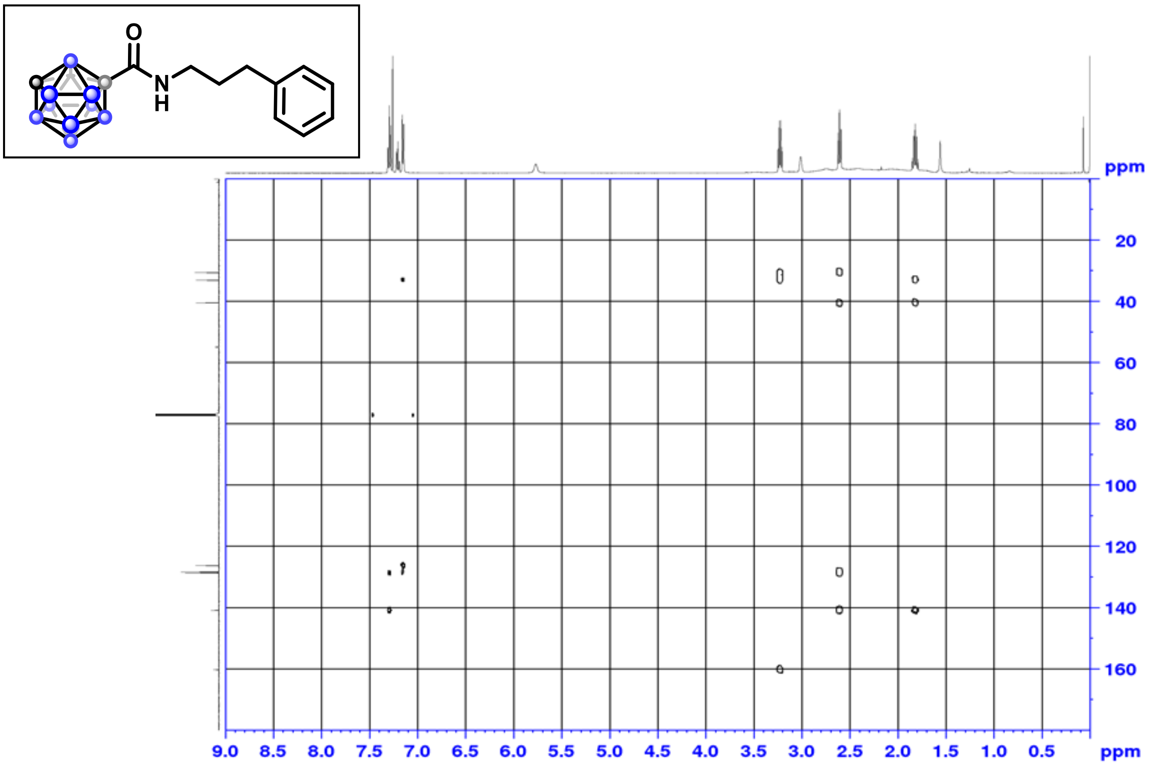

$^{1}H$-$^{15}N$ HMBC

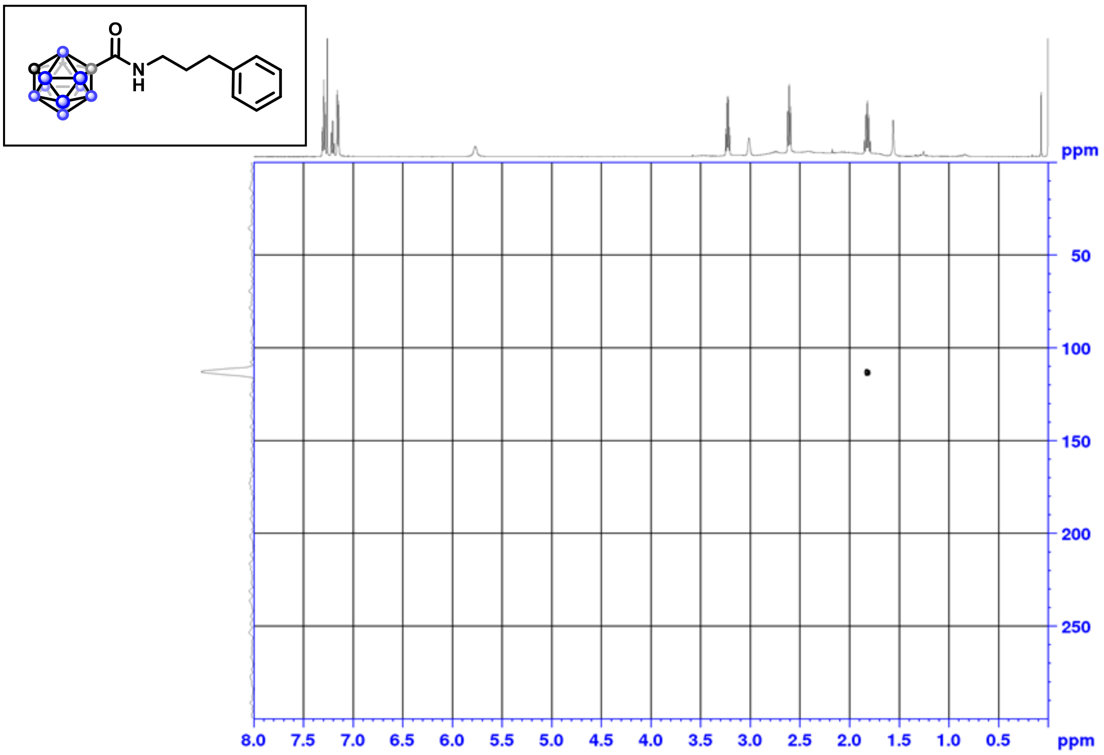

$^{1}H$ NMR (500 MHz, $CDCl_3$)

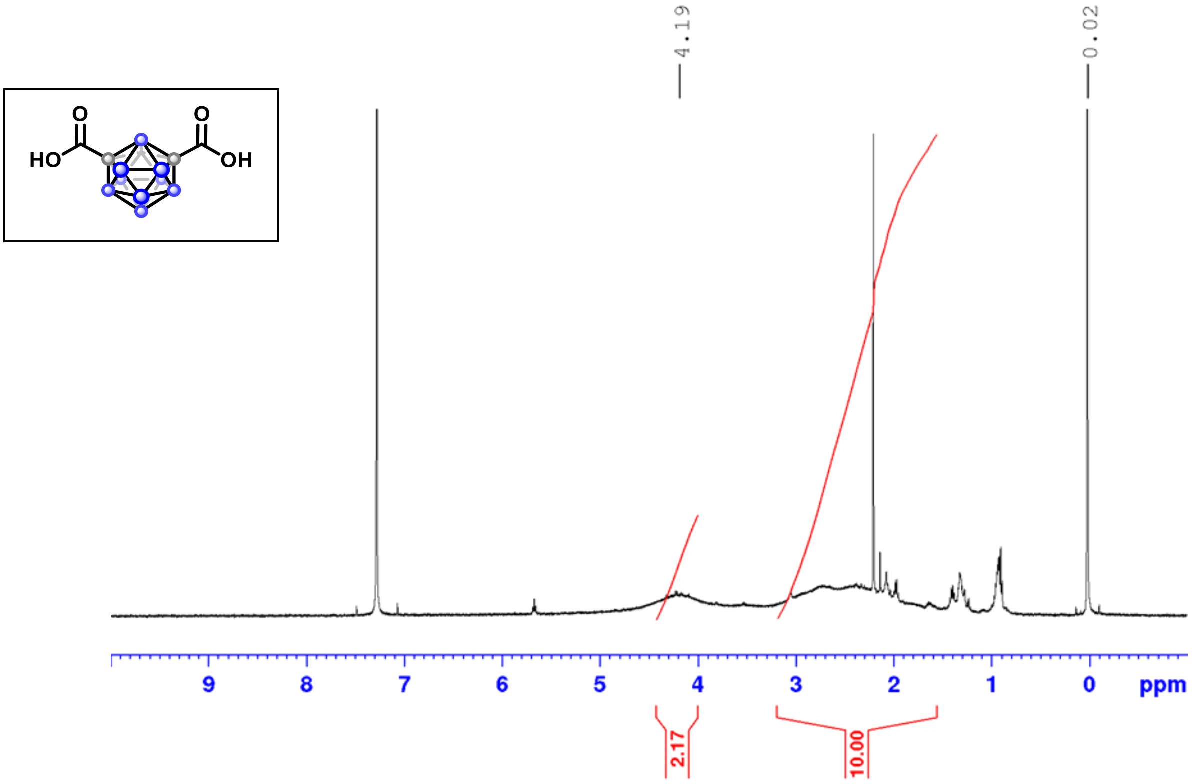

$^{1}H$ NMR (500 MHz, $CDCl_3$)

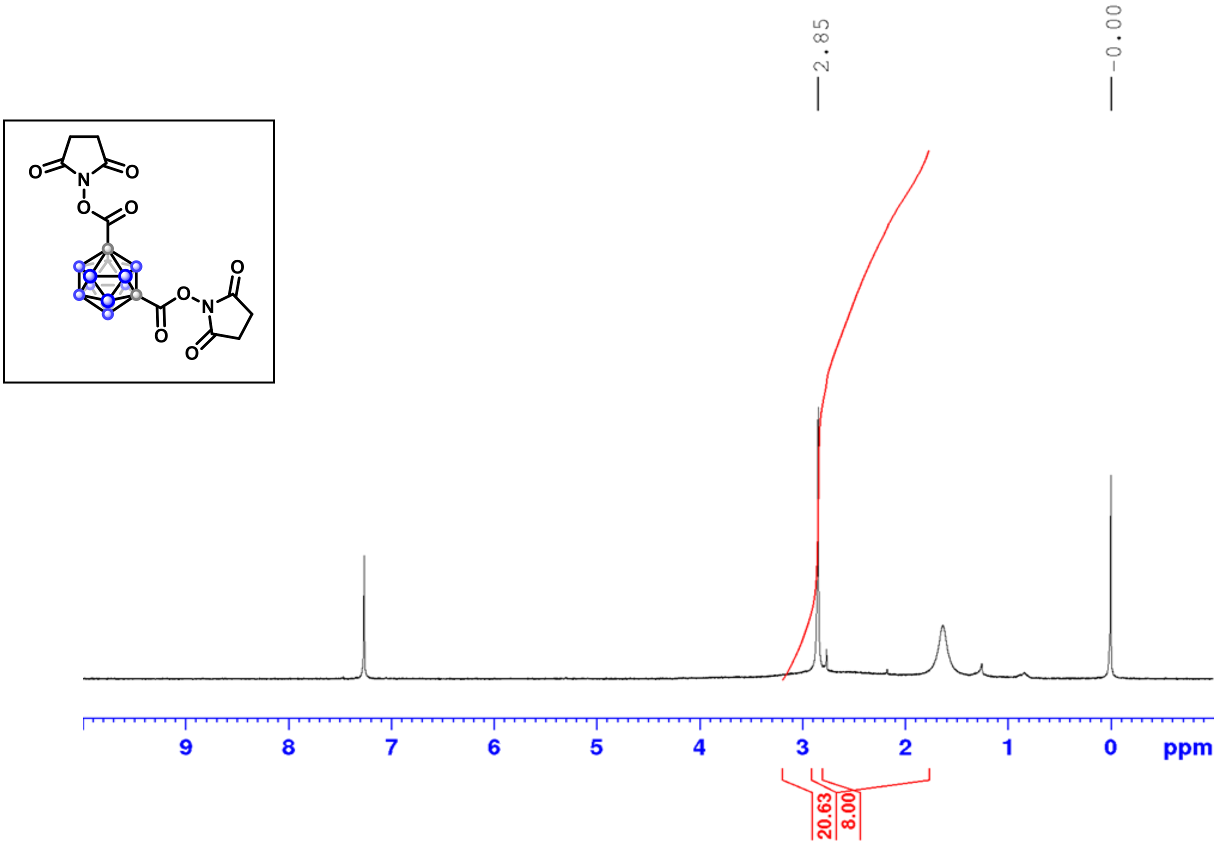

$^{13}C$ NMR (126 MHz, $CDCl_3$)

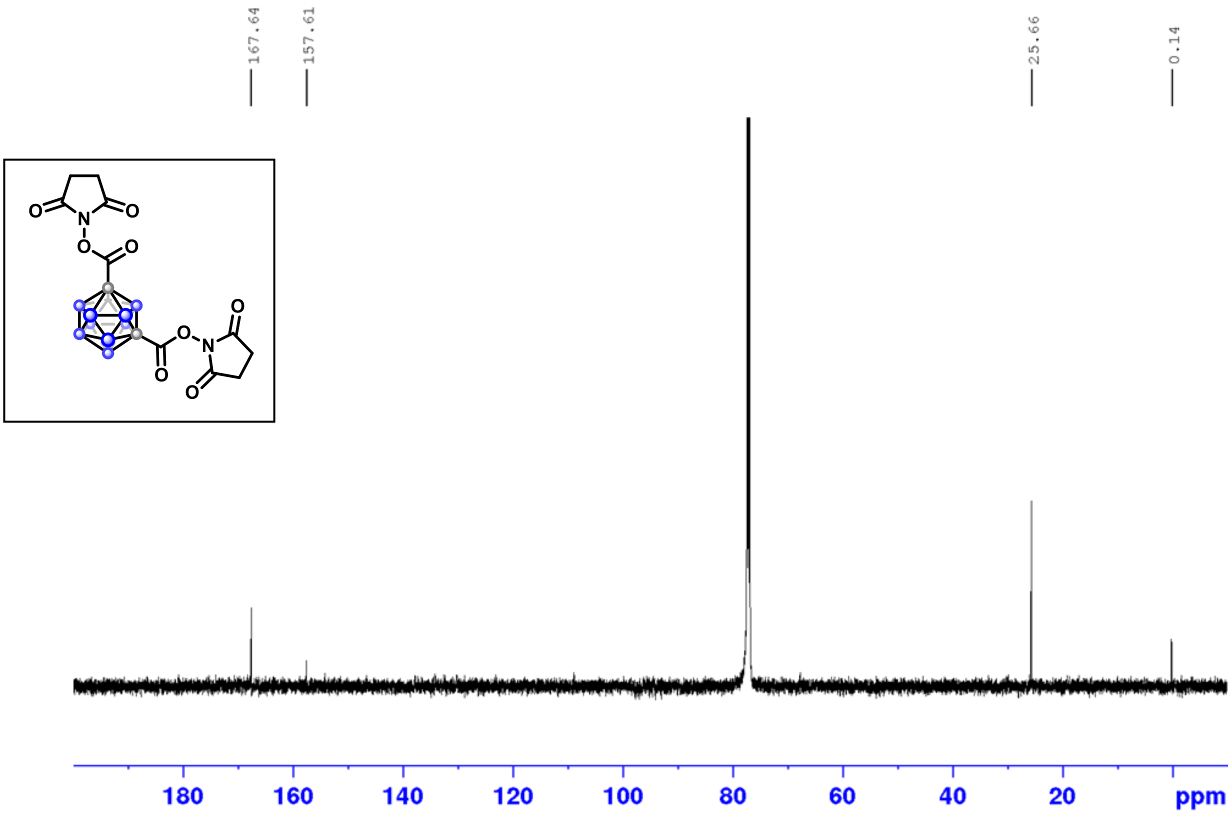

$^{1}H$-$^{13}C$ HSQC

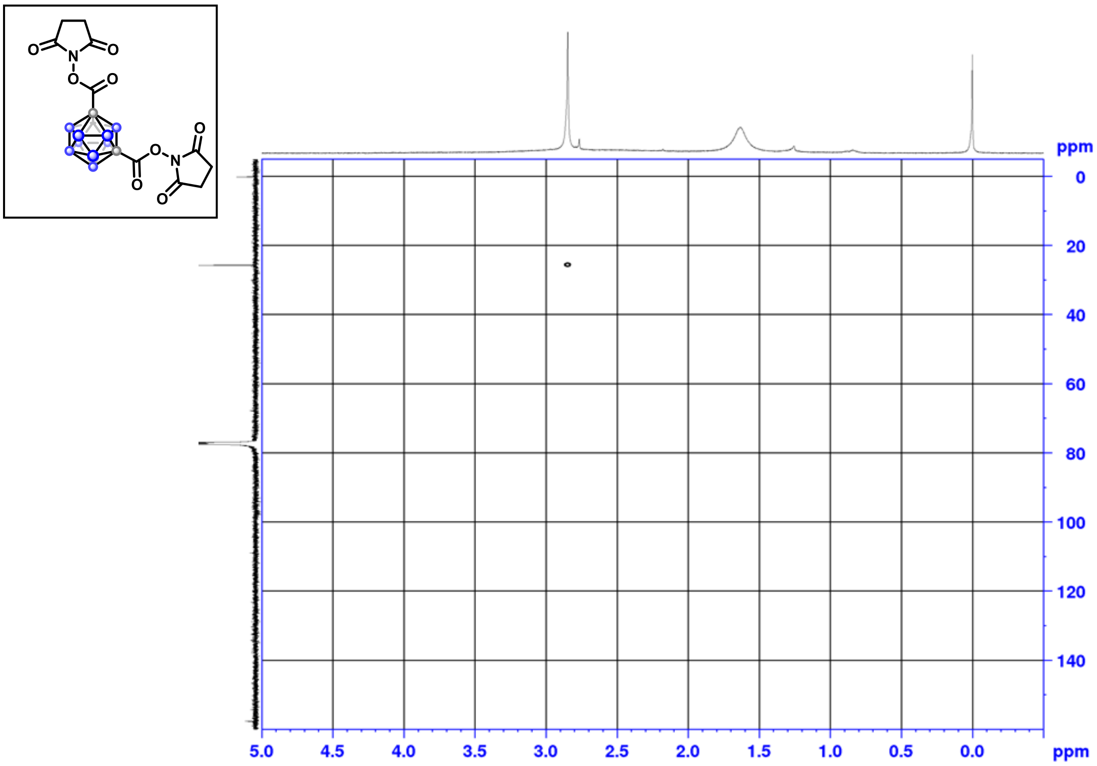

$^{1}H$-$^{13}C$ HMBC

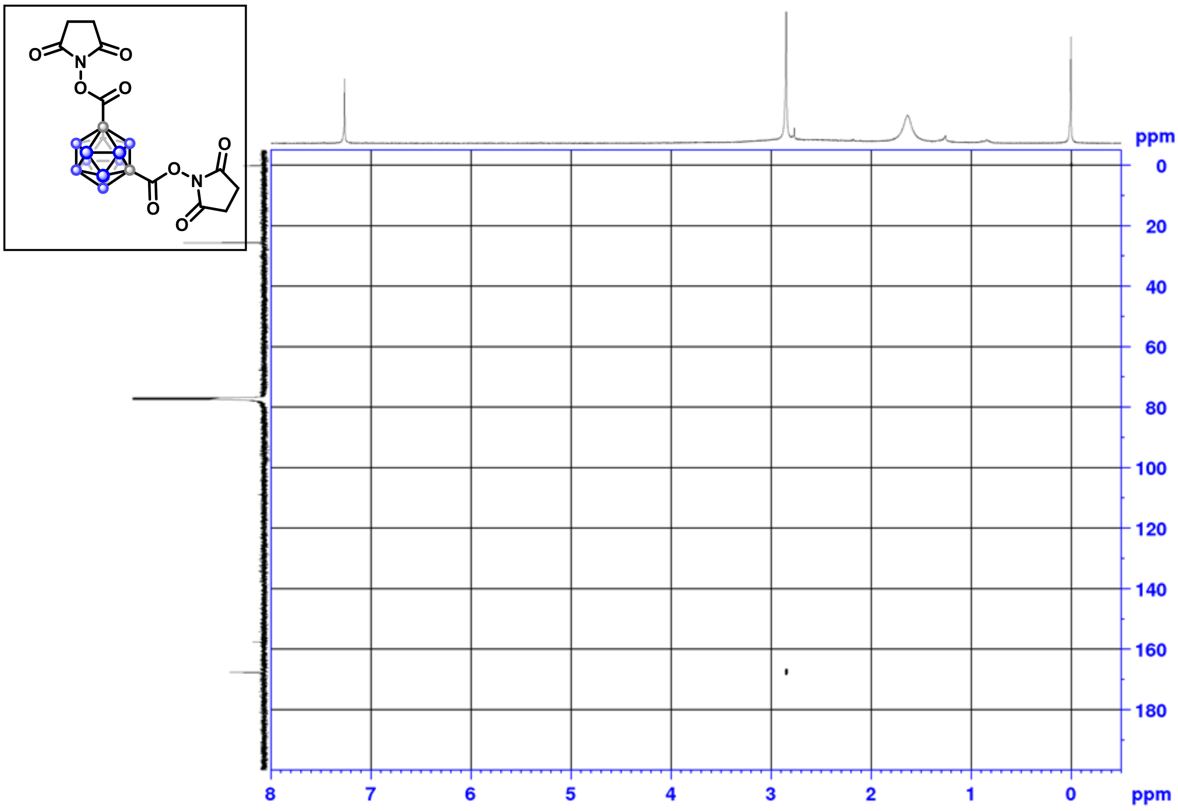

[1]H NMR (500 MHz, $CDCl_3$)

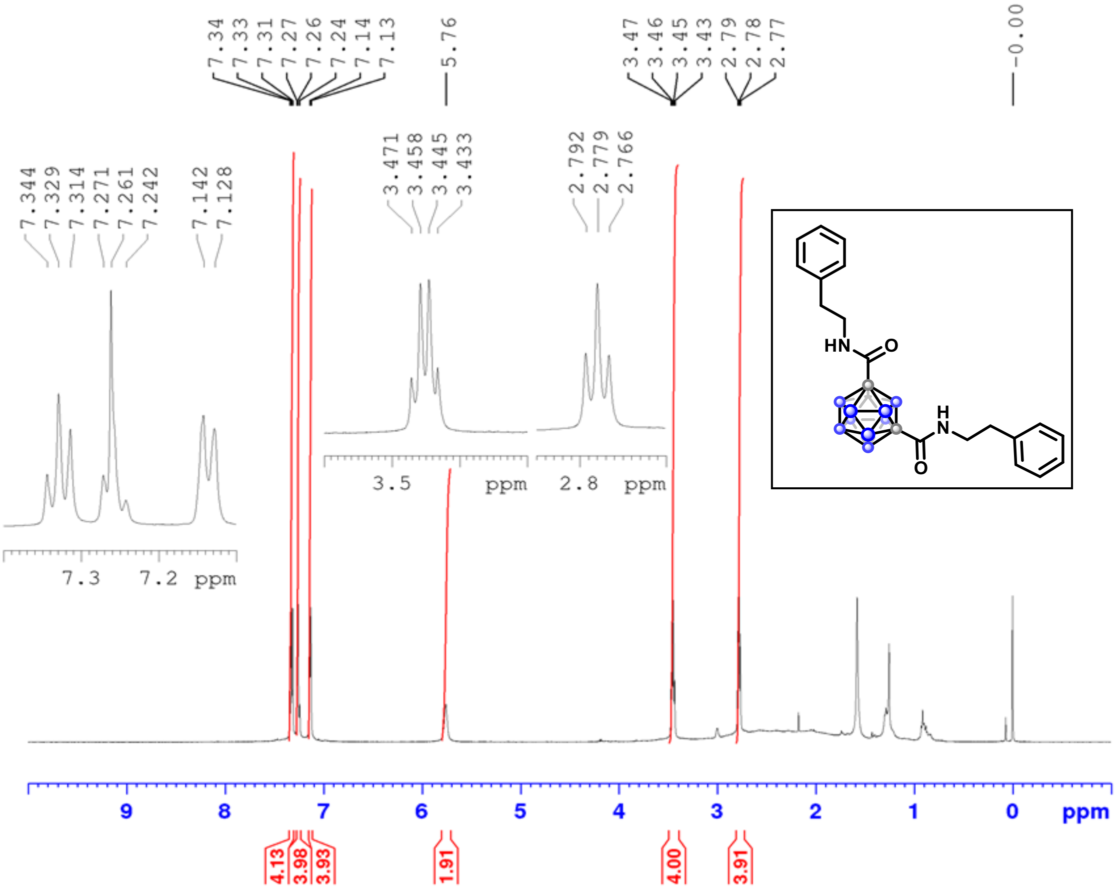

[13]C NMR (126 MHz, $CDCl_3$)

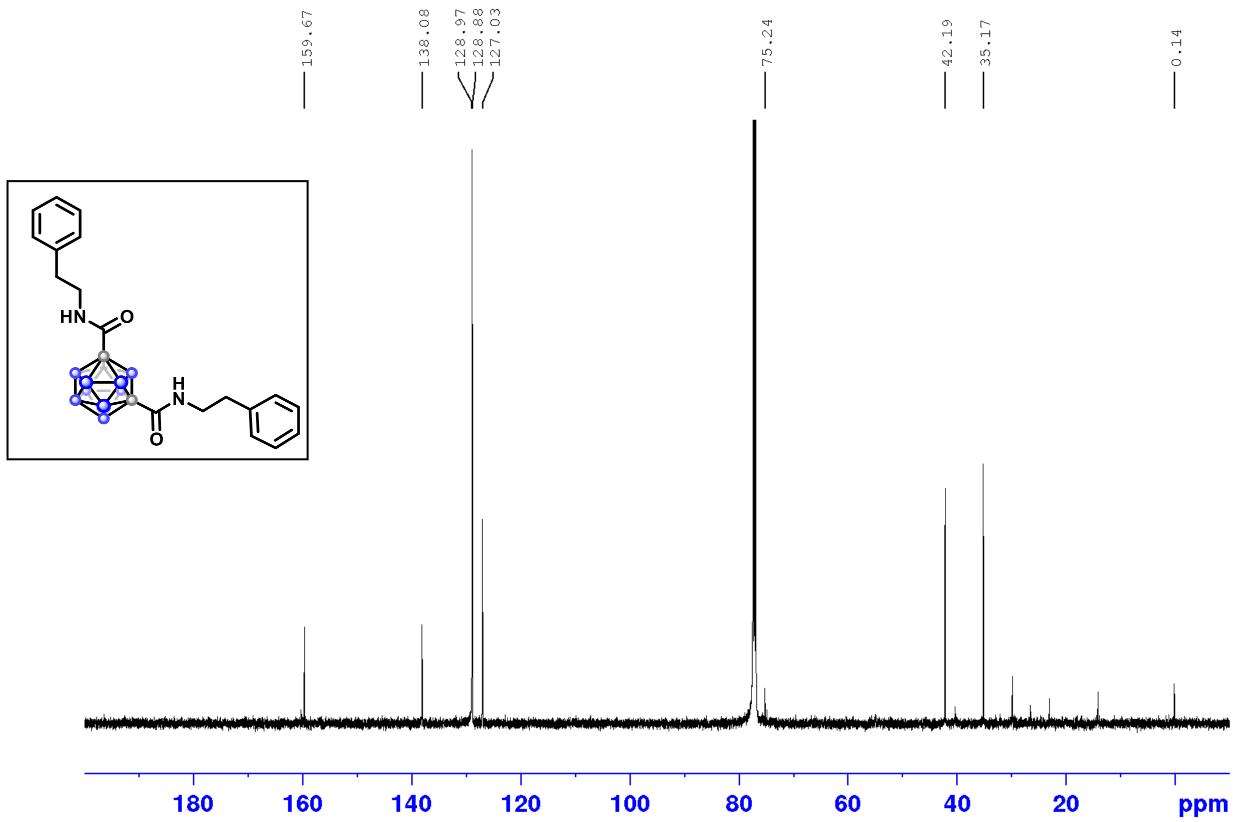

$^{11}B$ NMR (160 MHz, $CDCl_3$)

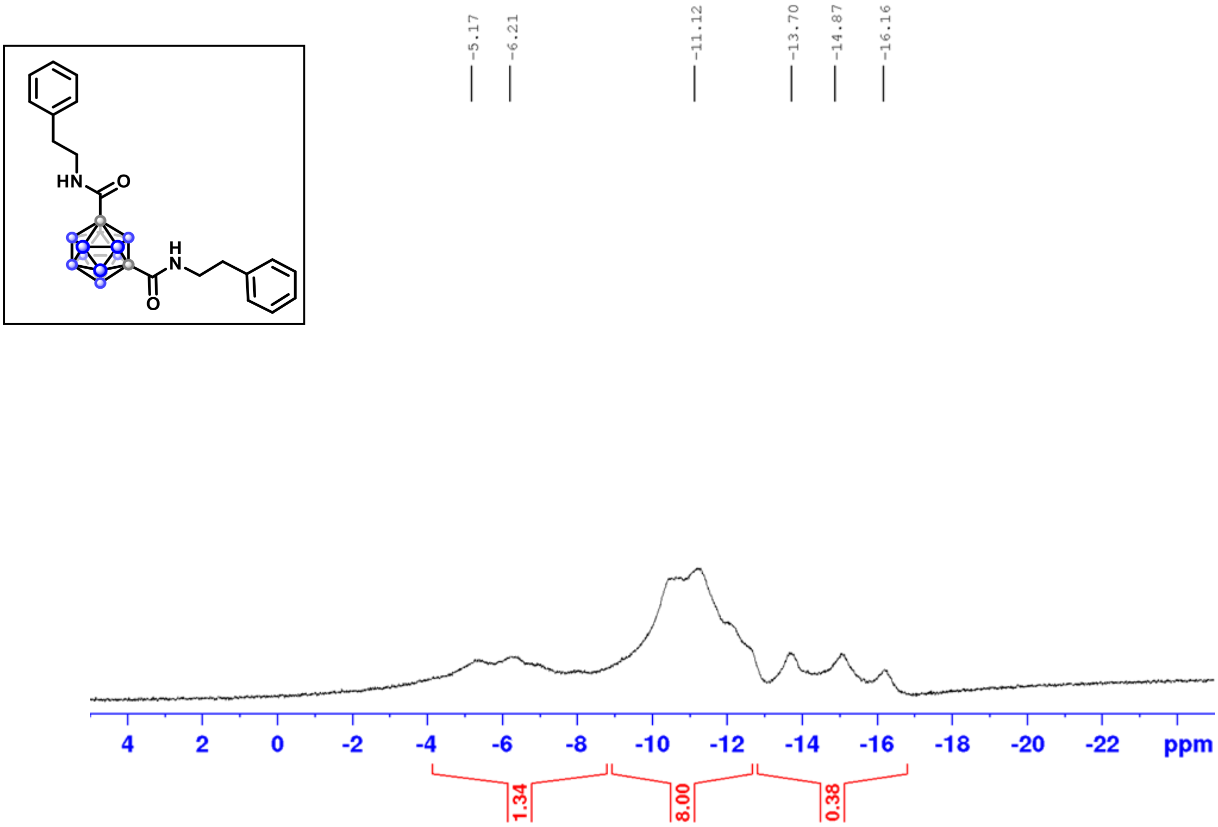

$^{1}H$-$^{1}H$ COSY

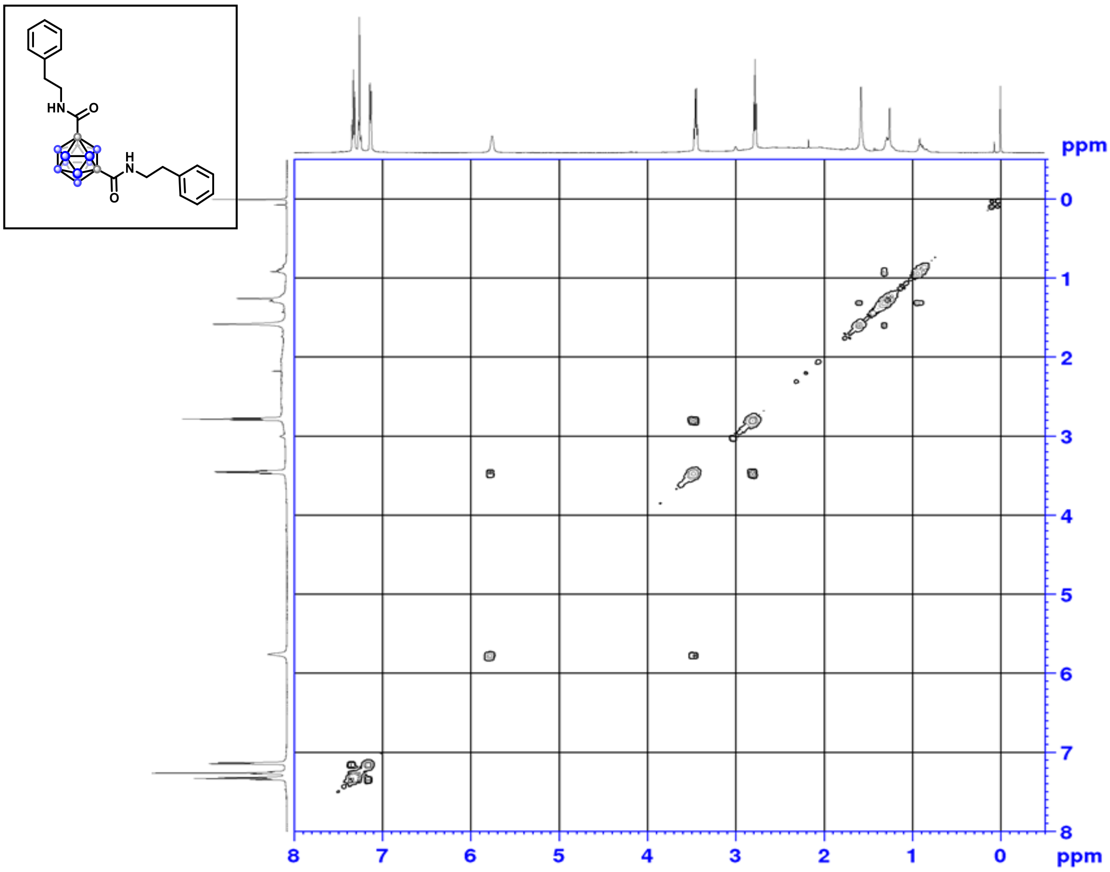

$^{1}H$-$^{13}C$ HSQC

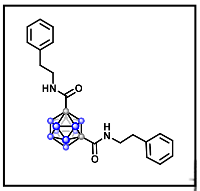

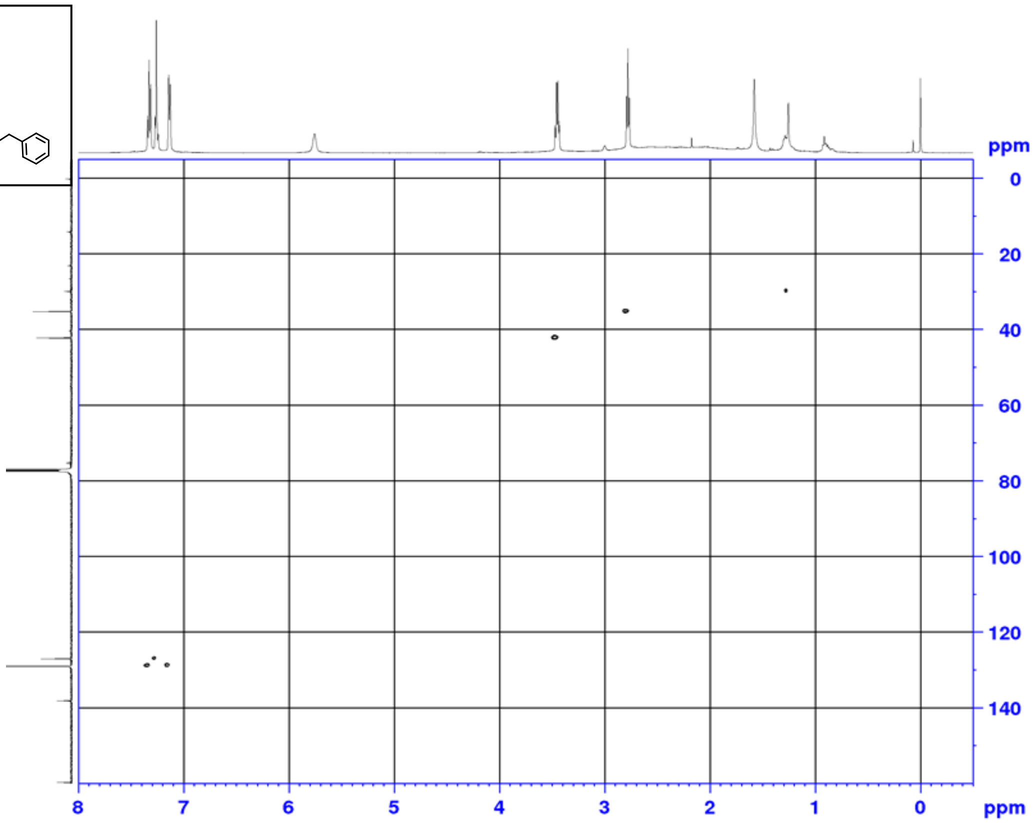

$^{1}H$-$^{13}C$ HMBC

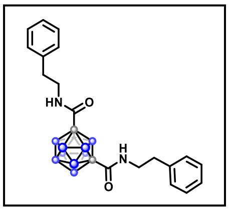

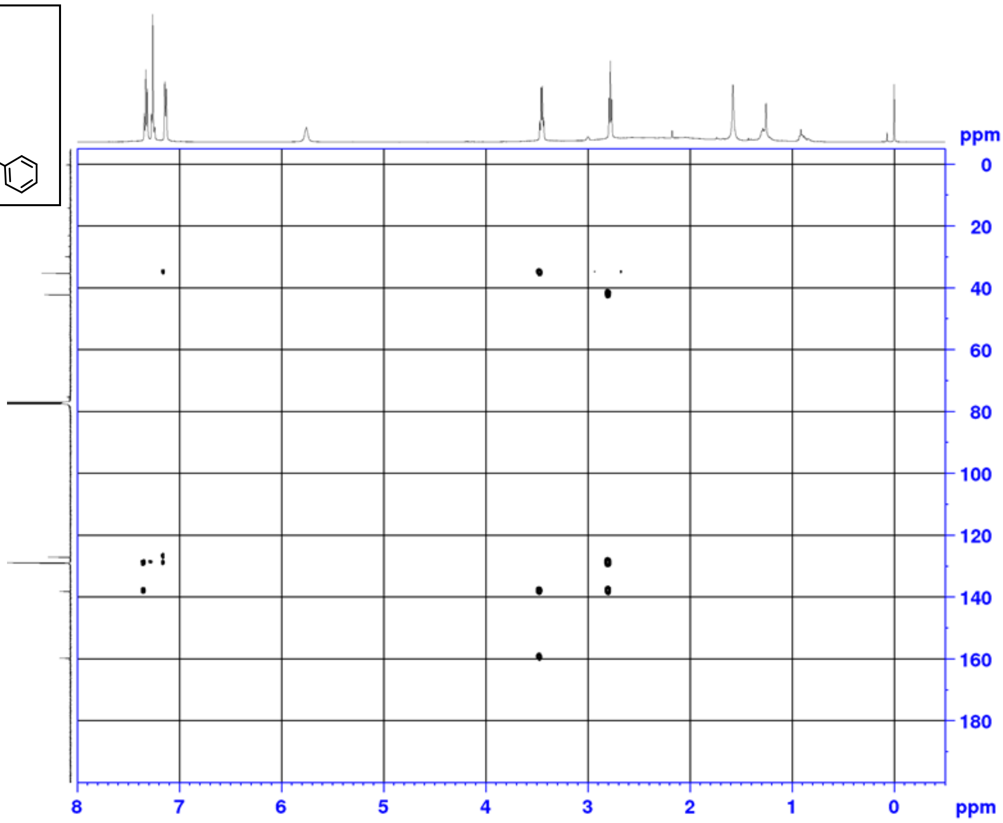

$^{1}H$-$^{15}N$ HMBC

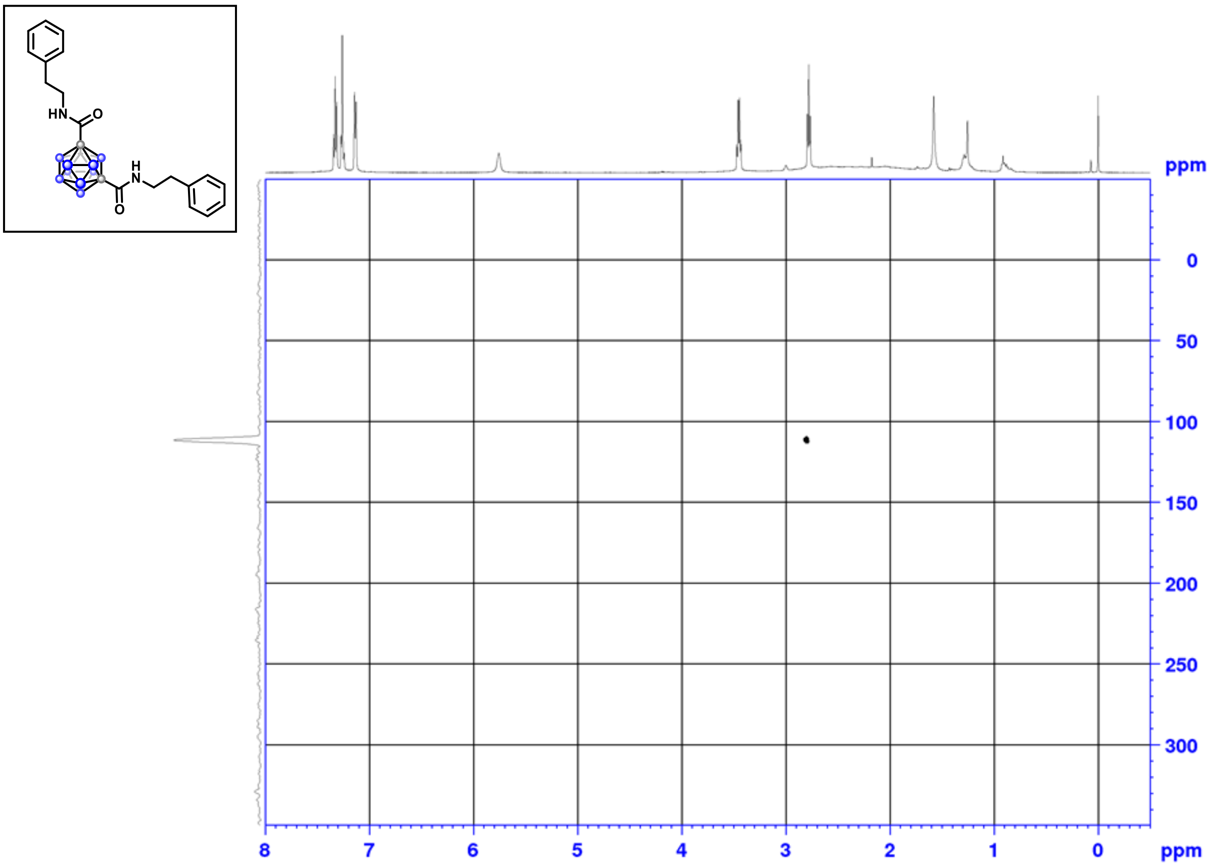

$^{1}H$ NMR (500 MHz, $CDCl_3$)

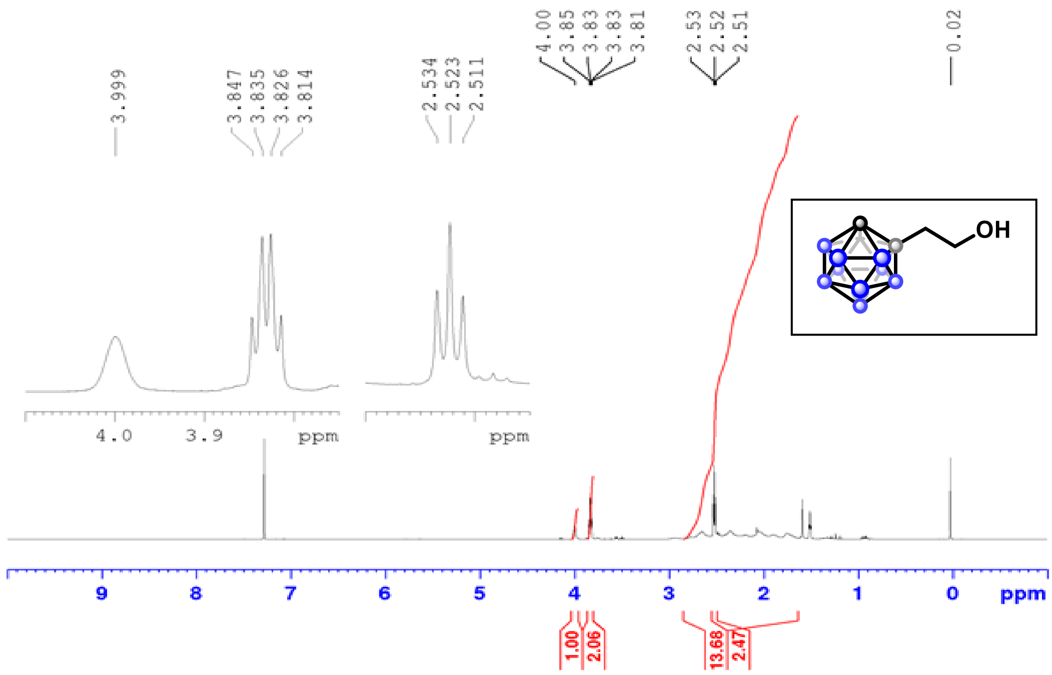

$^{1}H$ NMR (500 MHz, $CDCl_3$)

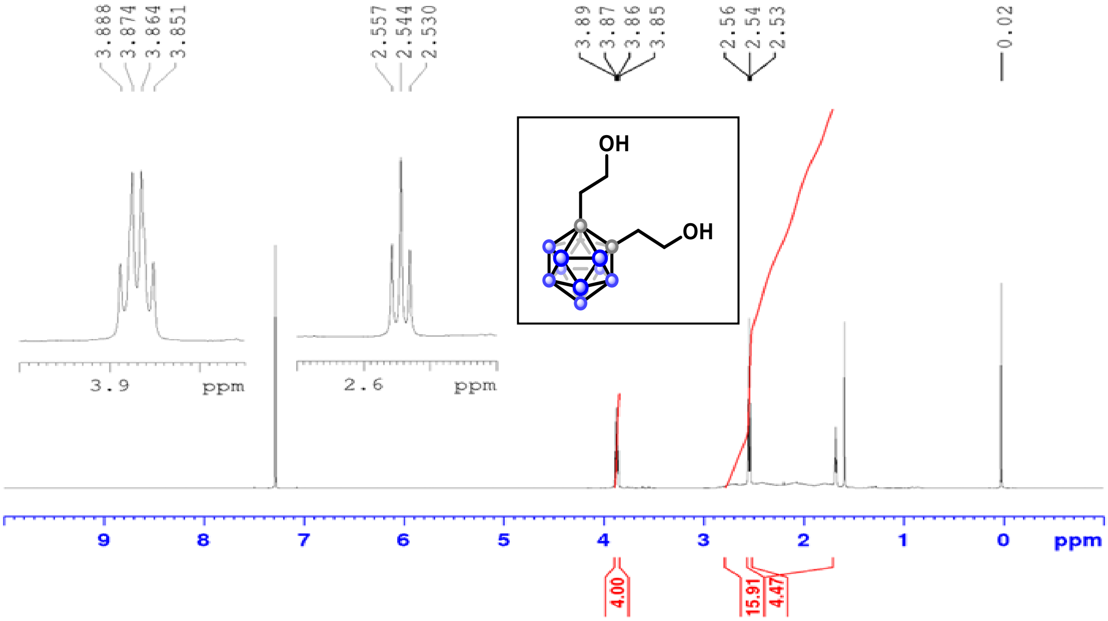

$^{1}H$ NMR (500 MHz, $CDCl_3$)

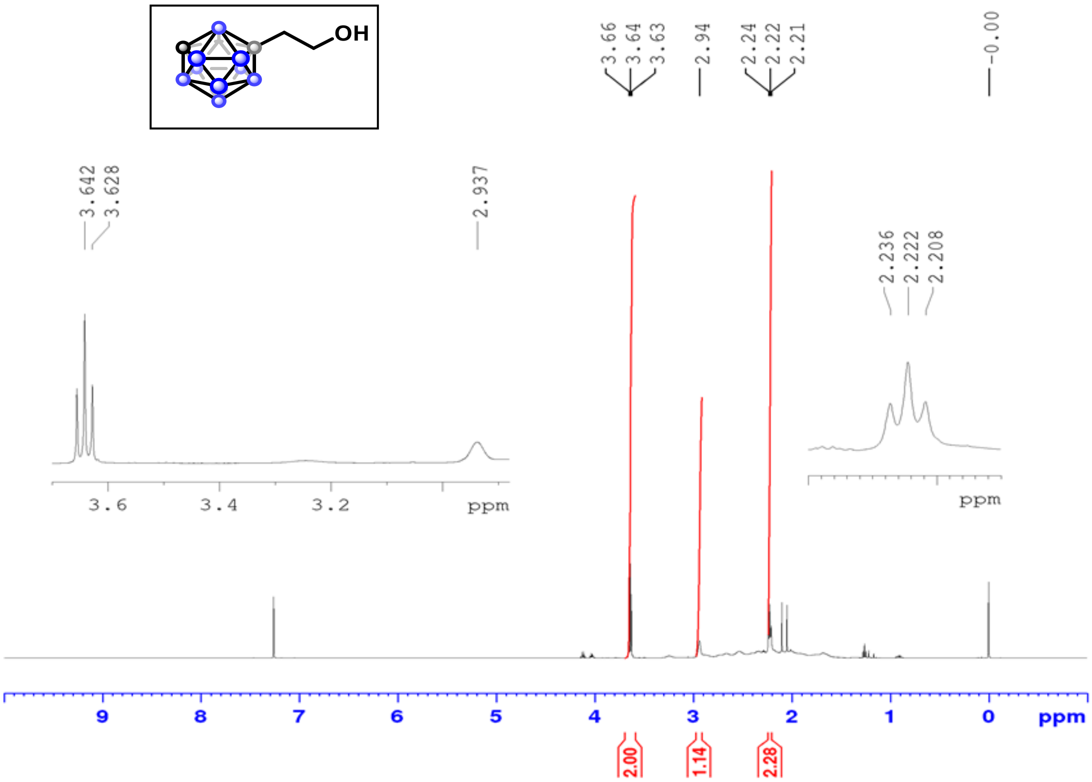

[1]H NMR (500 MHz, $CDCl_3$)

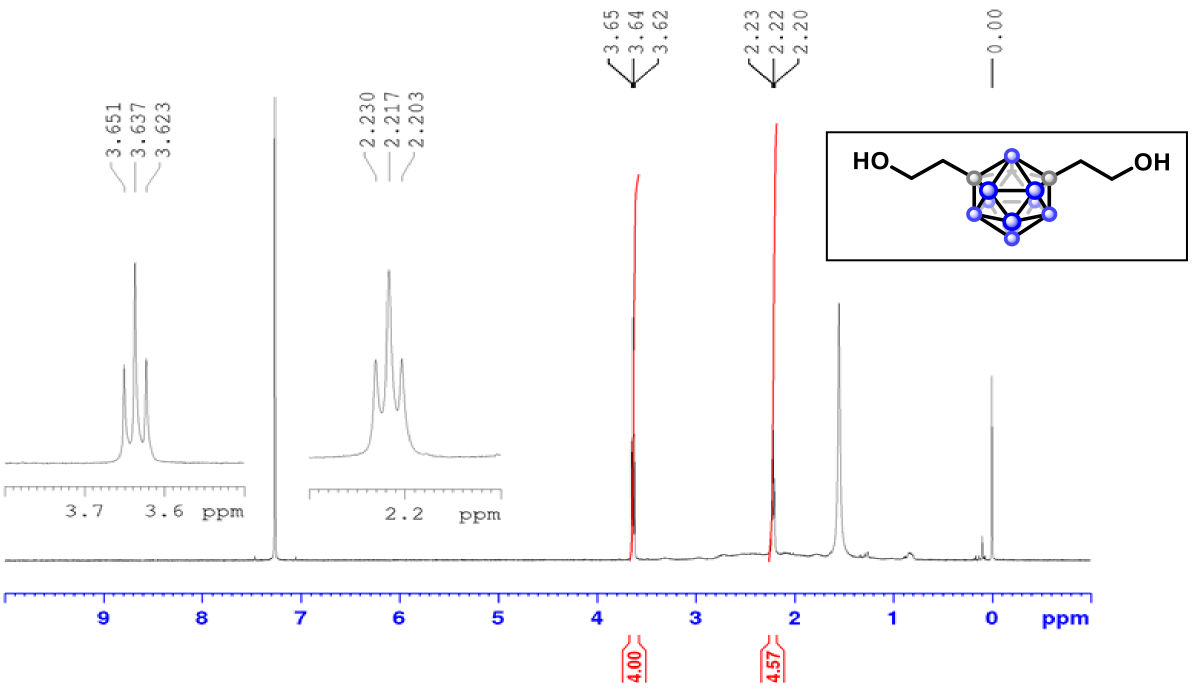

[1]H NMR (500 MHz, $CDCl_3$)

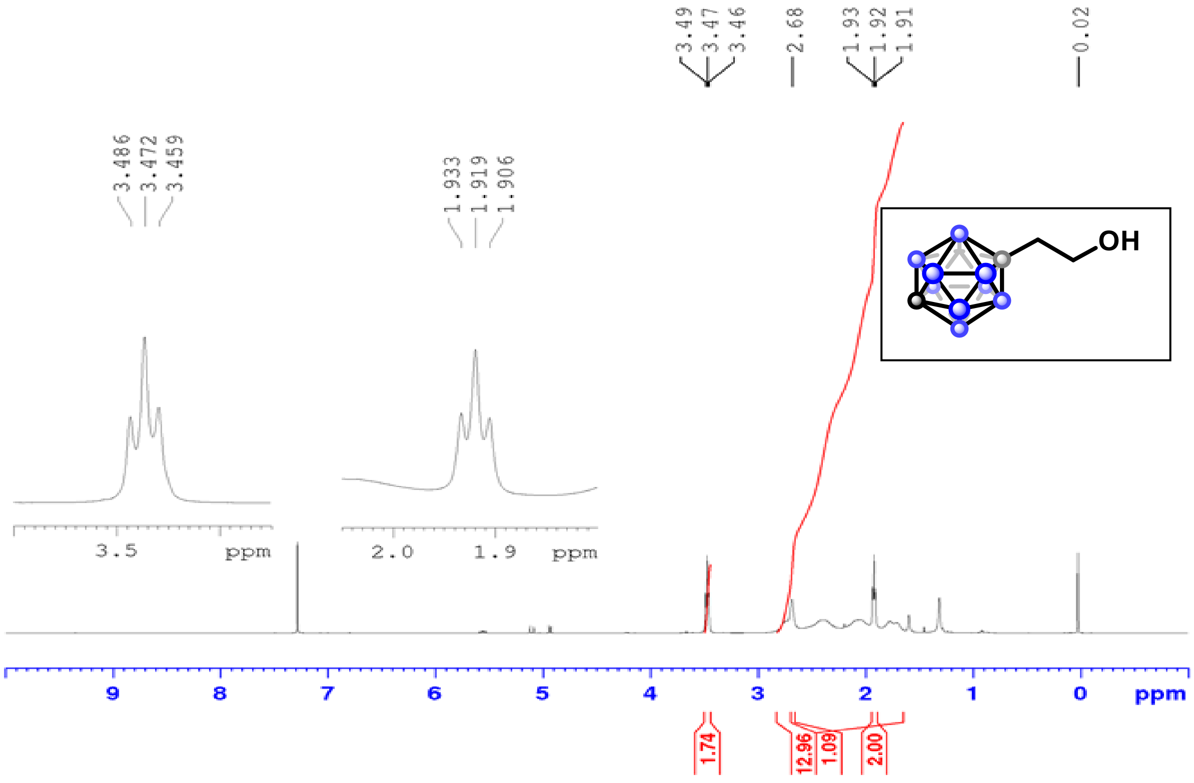

$^1$H NMR (500 MHz, $CDCl_3$)

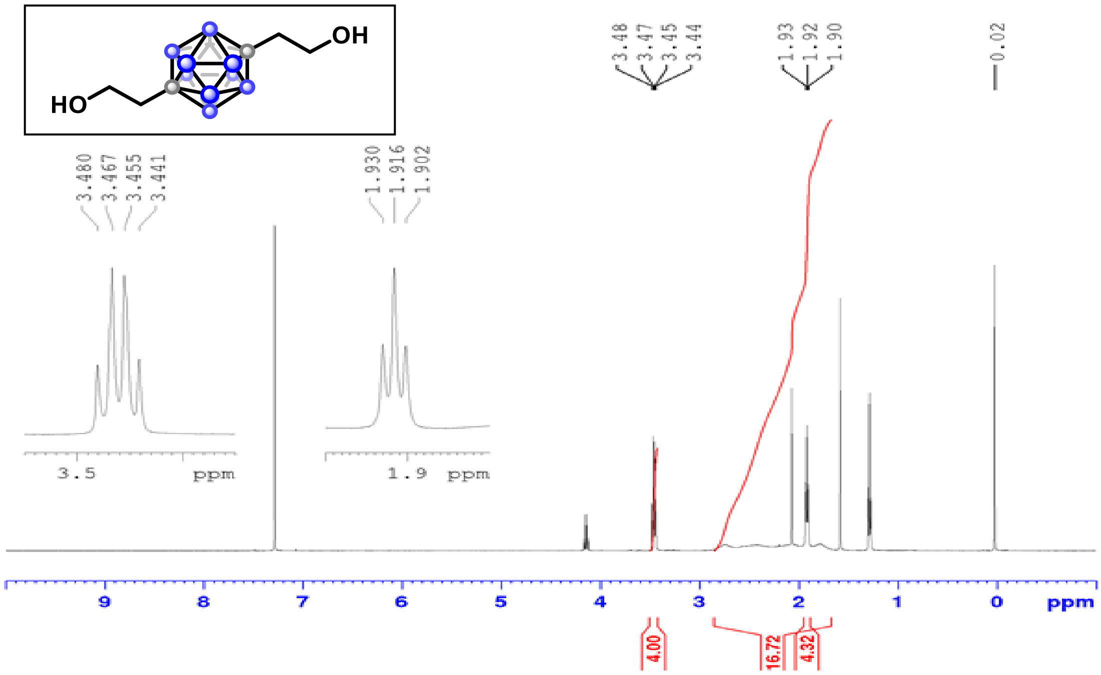

$^1$H NMR (500 MHz, $CDCl_3$)

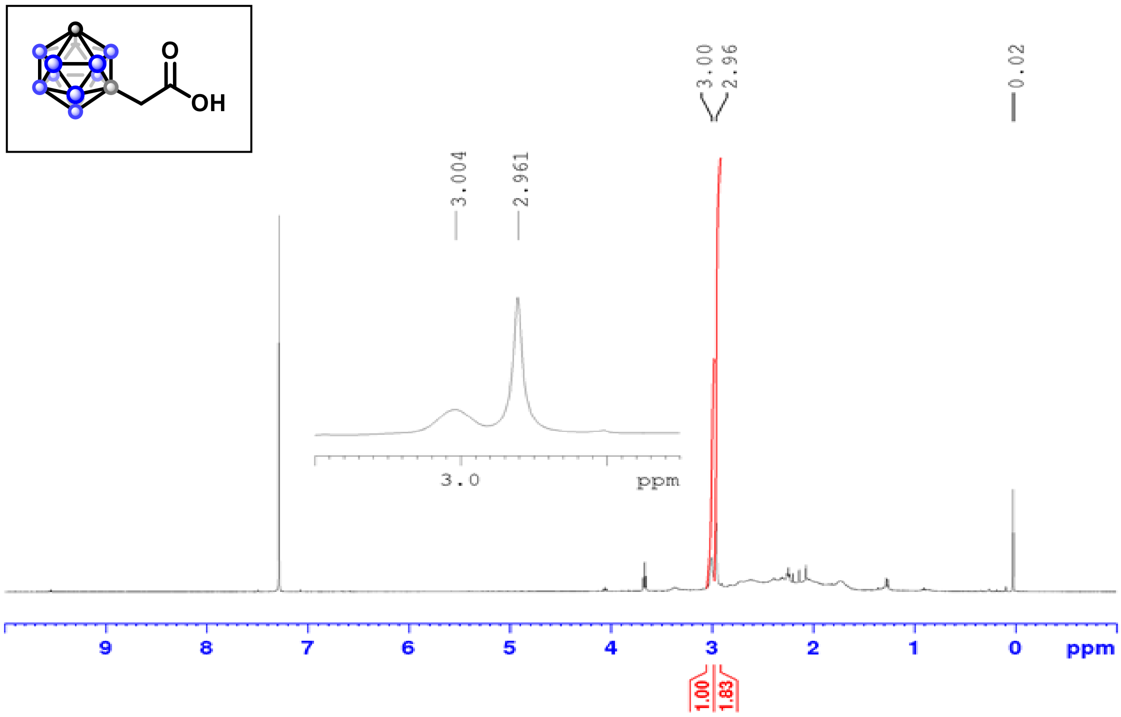

$^{1}$H NMR (500 MHz, $CDCl_3$)

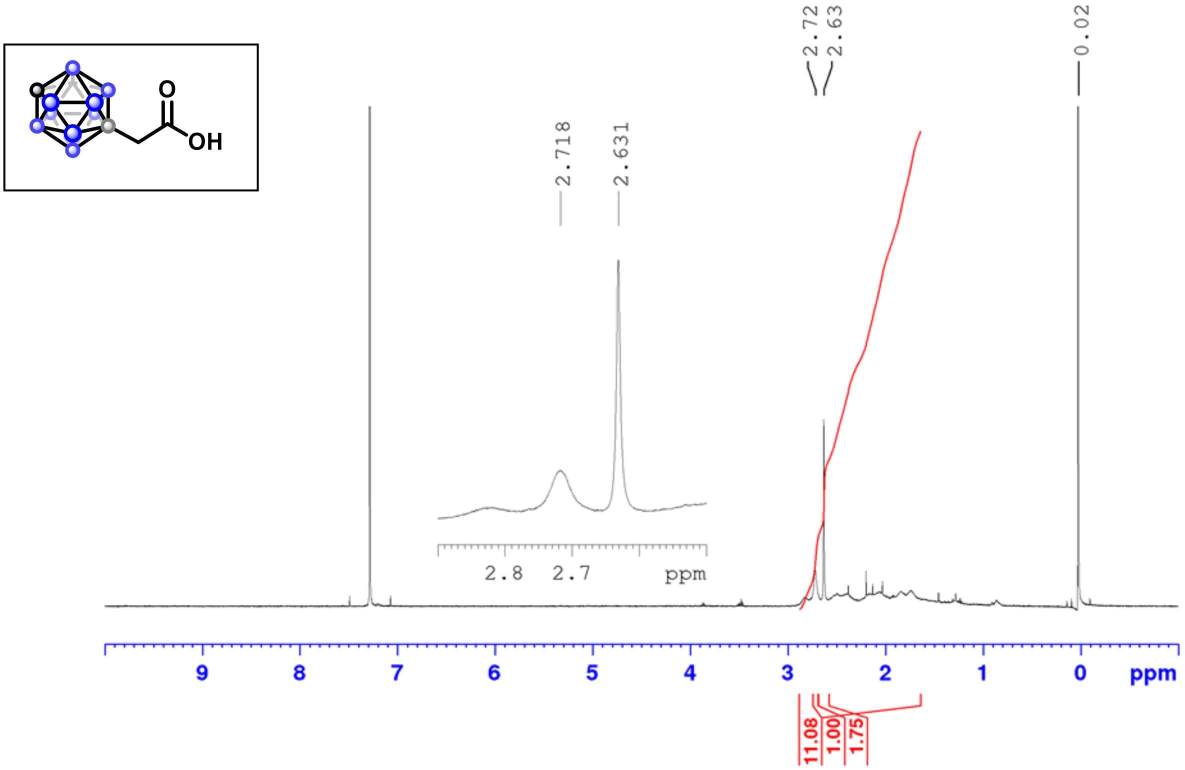

$^{1}$H NMR (500 MHz, $CDCl_3$)

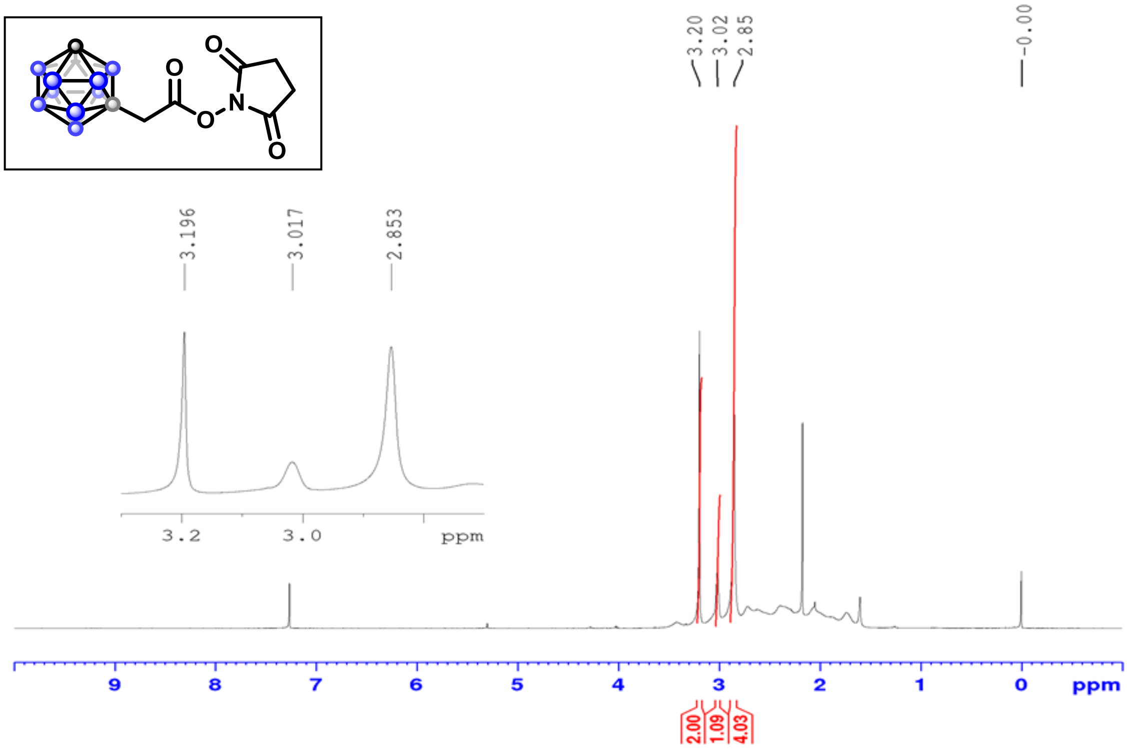

$^{13}$C NMR (126 MHz, $CDCl_3$)

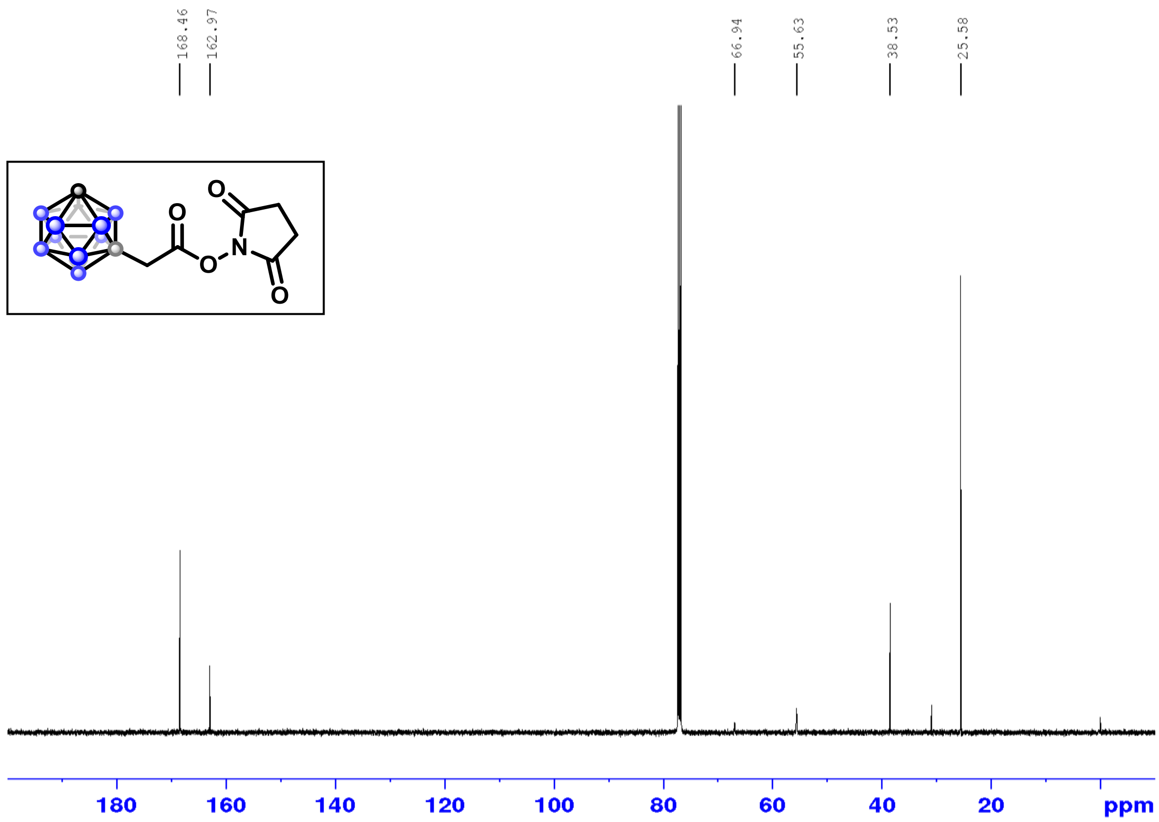

$^{11}$B NMR (160 MHz, $CDCl_3$)

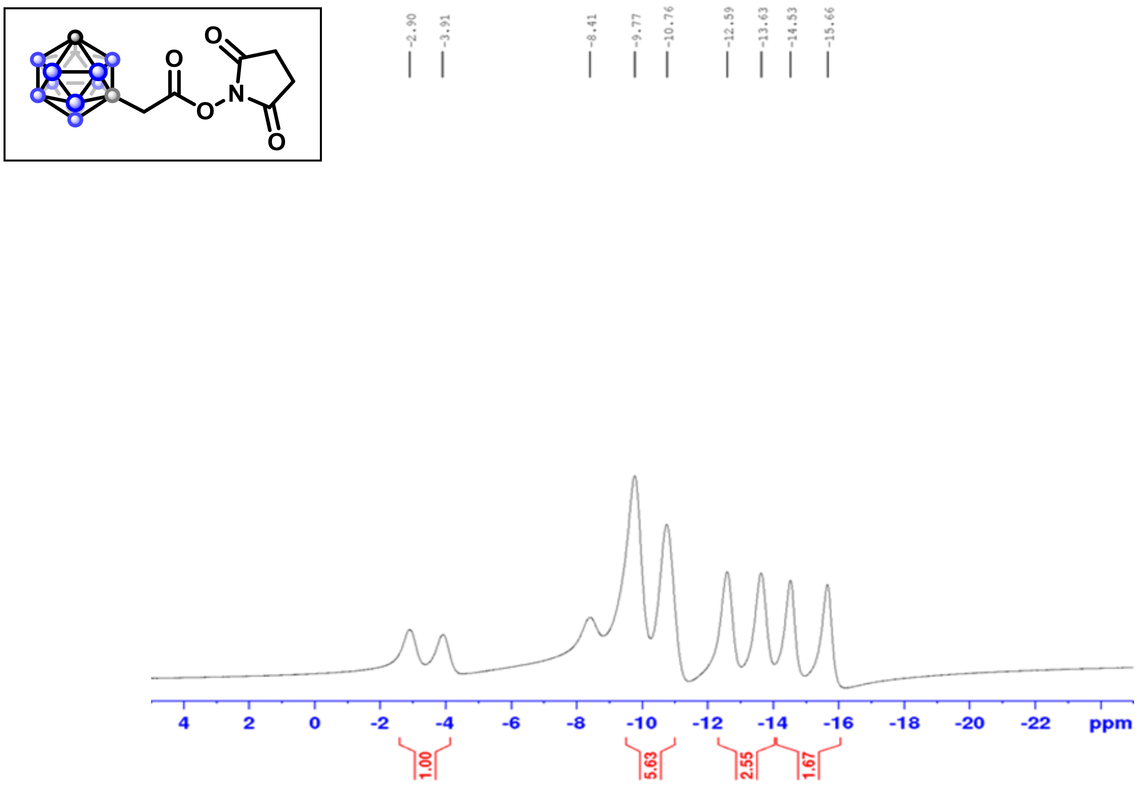

$^{1}H$-$^{1}H$ COSY

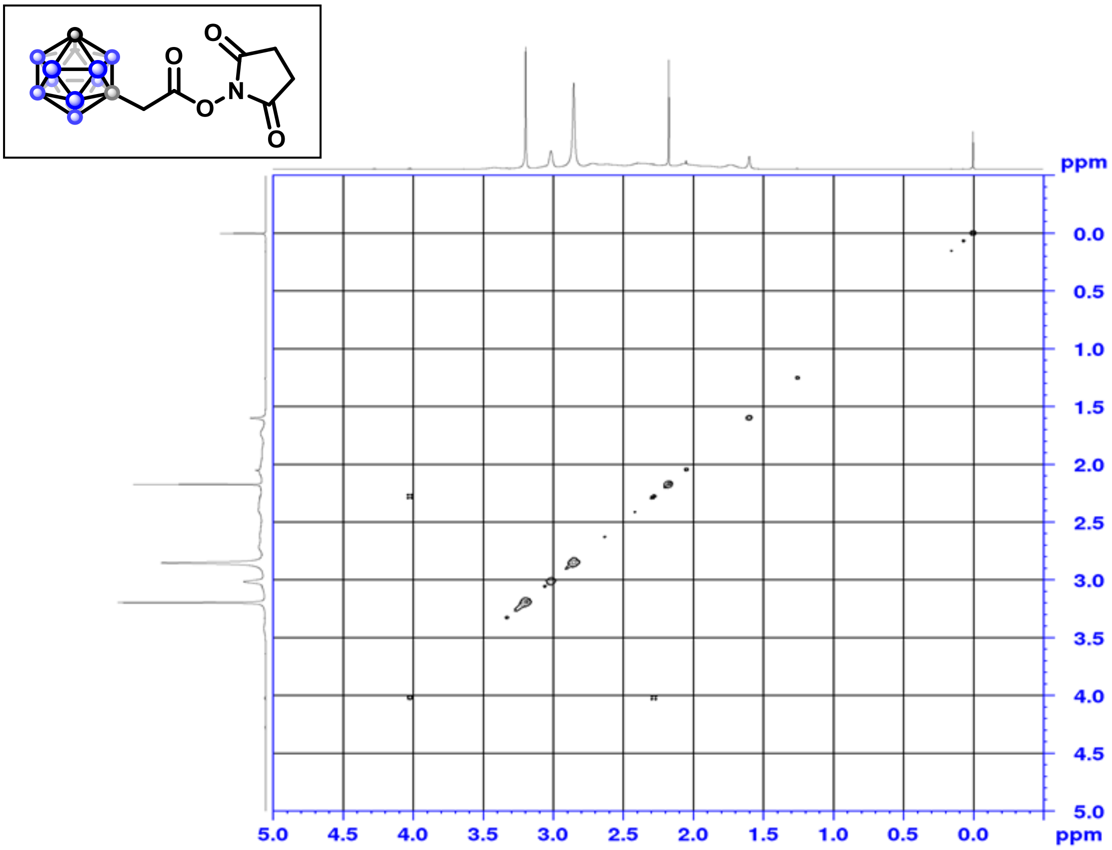

$^{1}H$-$^{13}C$ HSQC

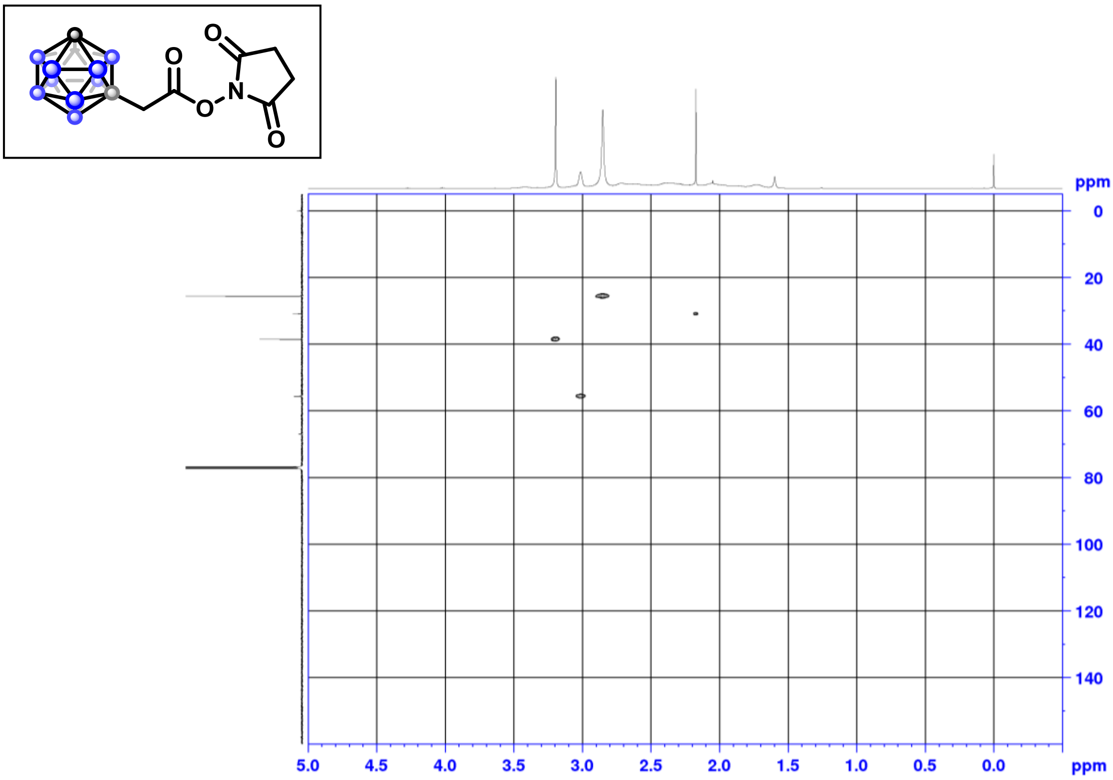

[1]H-[13]C HMBC

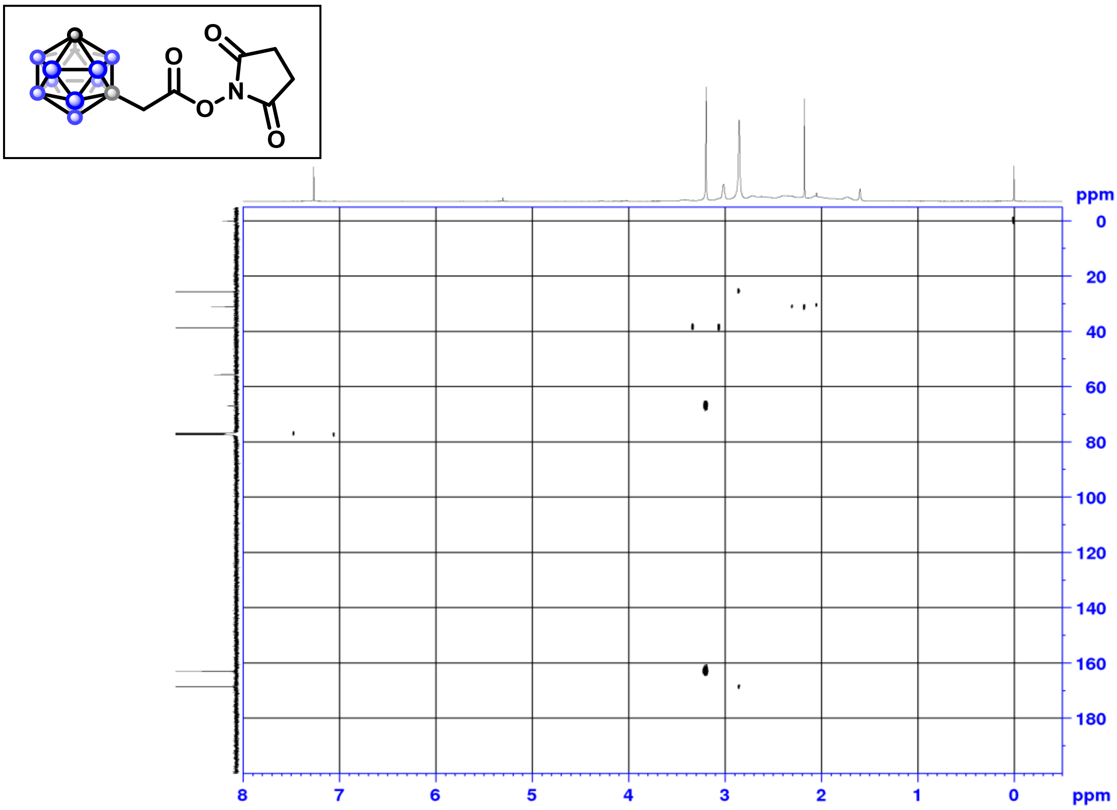

[1]H NMR (500 MHz, $CDCl_3$)

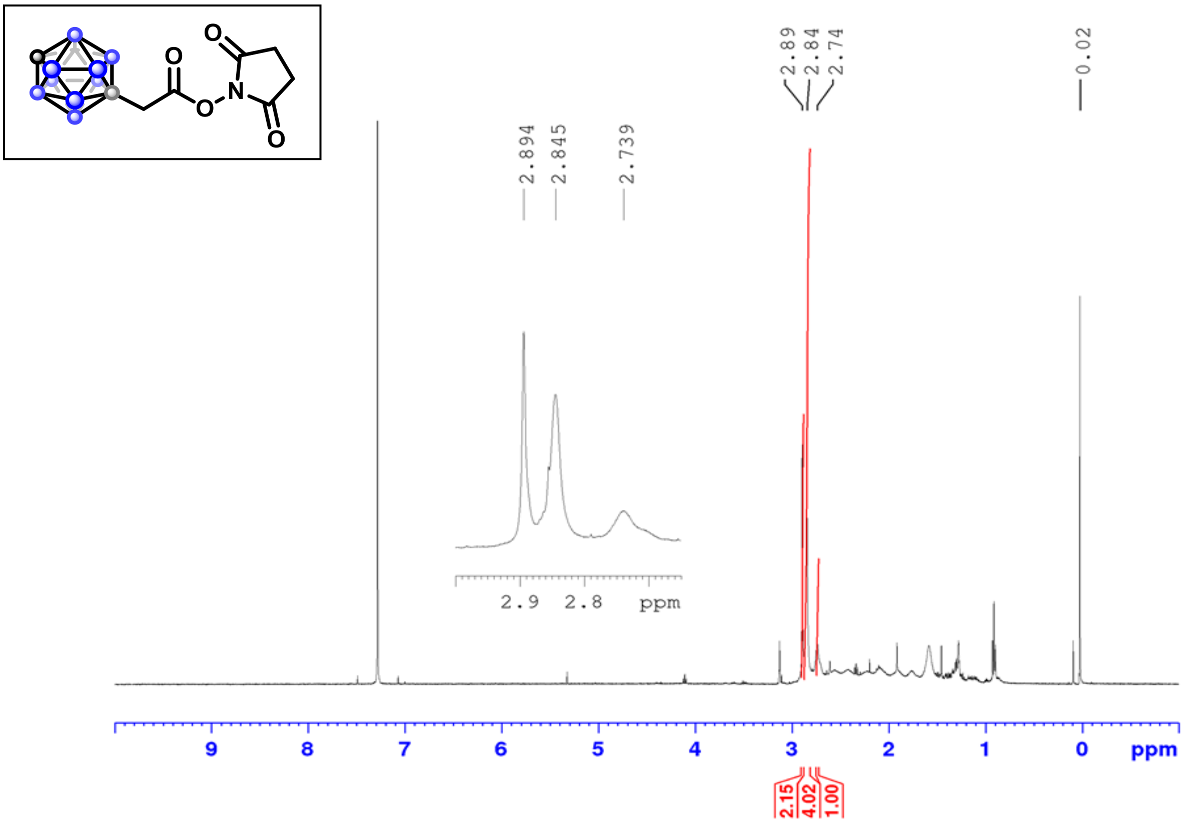

$^{13}C$ NMR (126 MHz, $CDCl_3$)

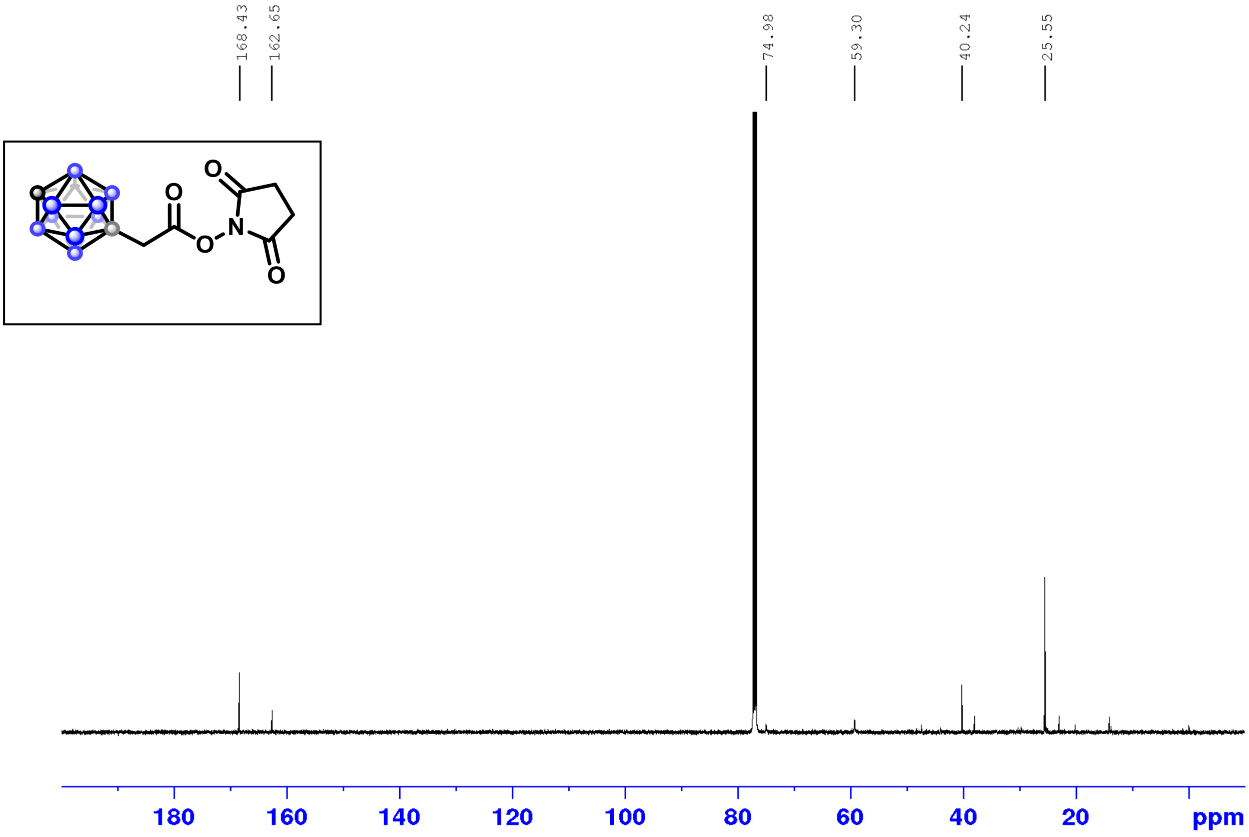

$^{11}B$ NMR (160 MHz, $CDCl_3$)

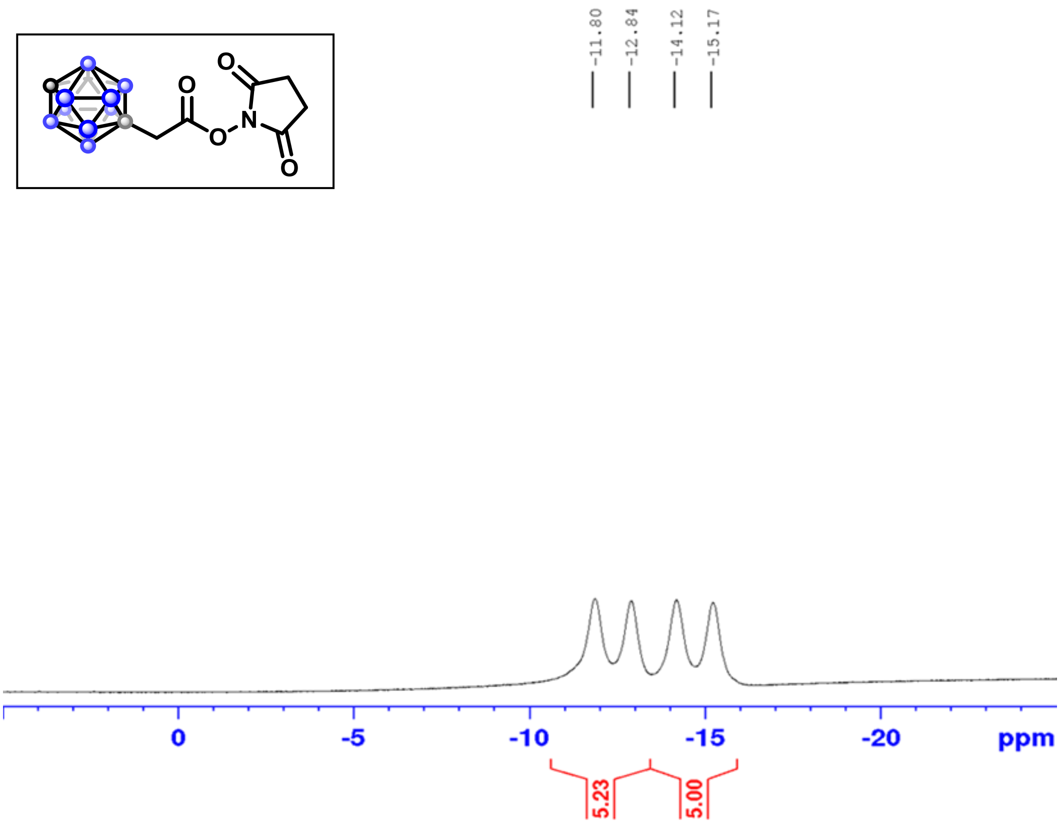

$^{1}H$-$^{13}C$ HSQC

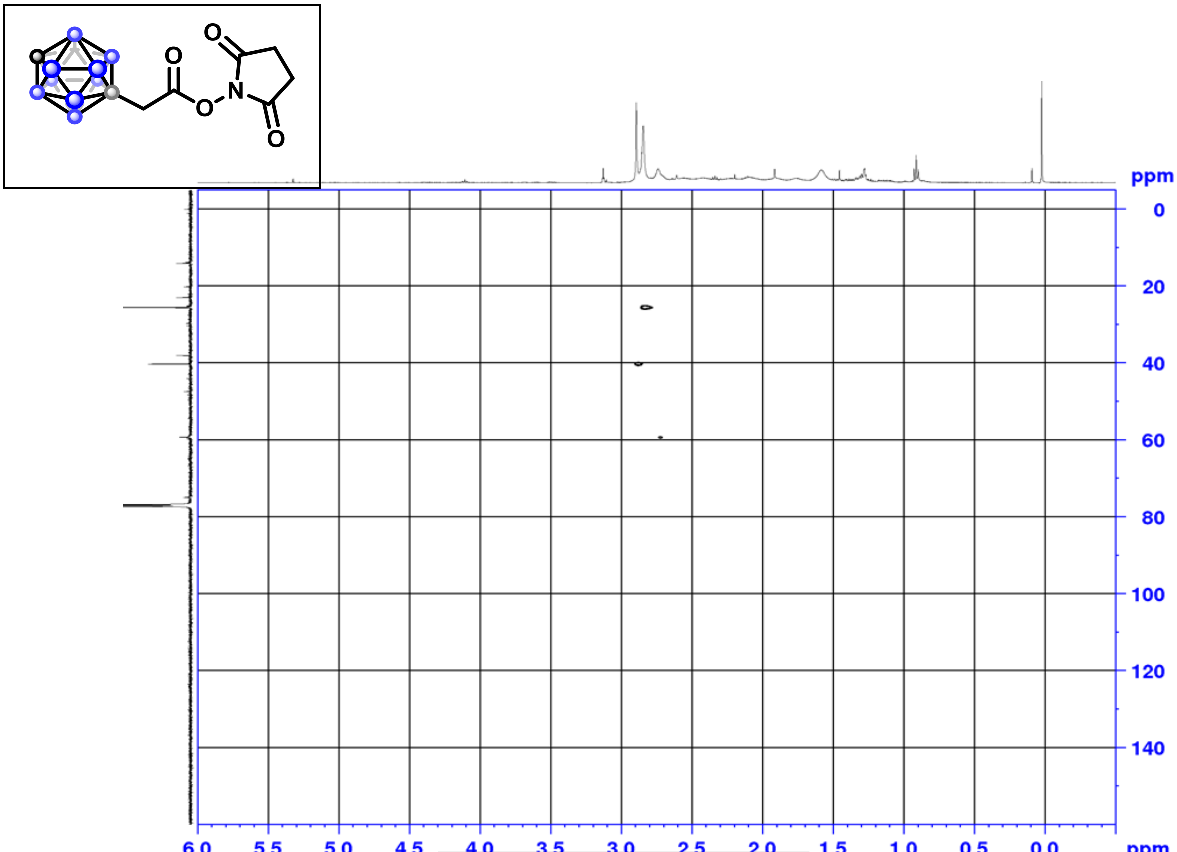

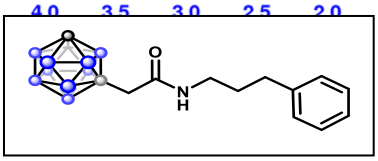

$^{1}H$ NMR (500 MHz, $CDCl_3$)

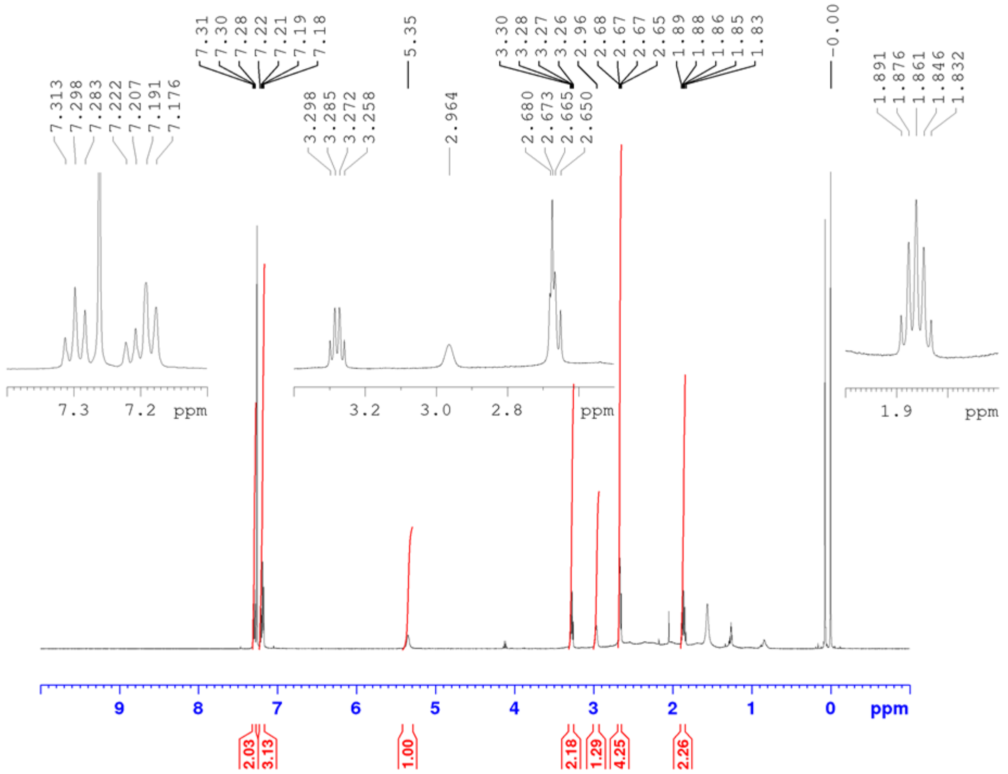

[13]C NMR (126 MHz, $CDCl_3$)

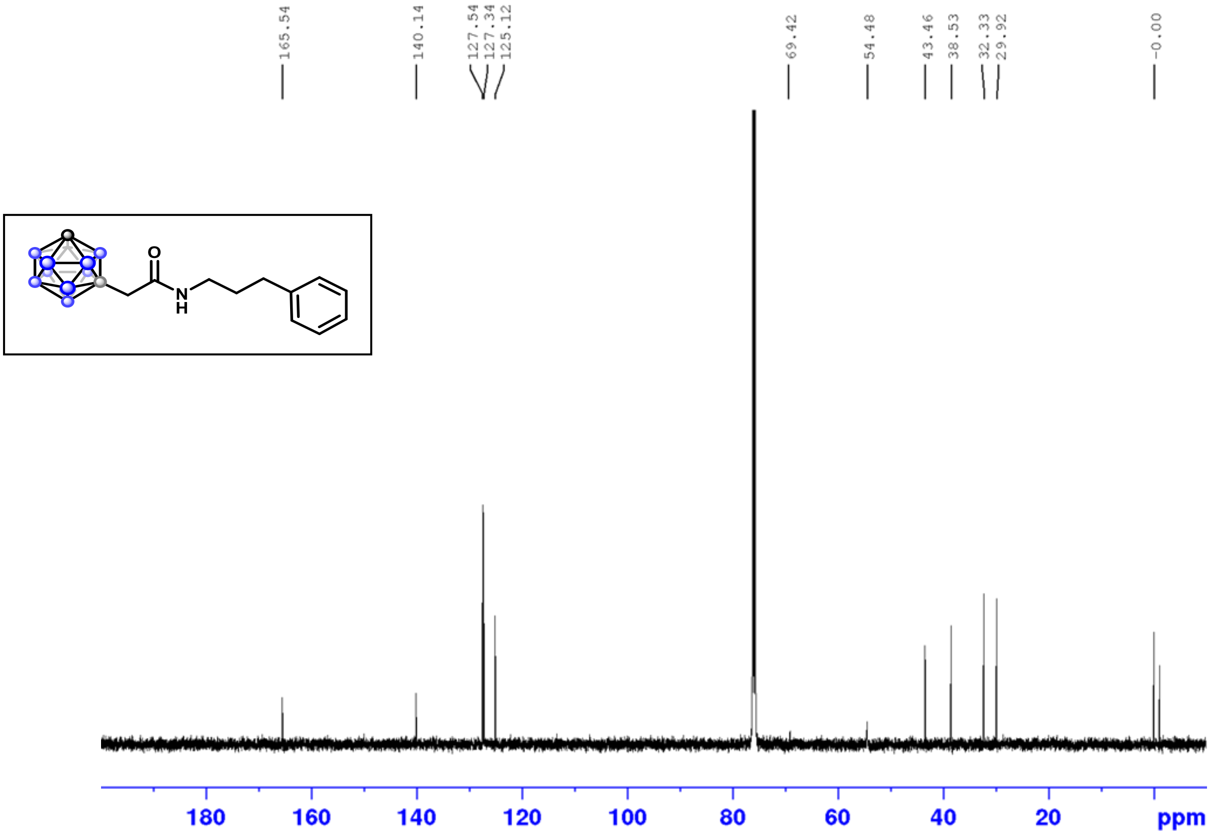

[11]B NMR (160 MHz, $CDCl_3$)

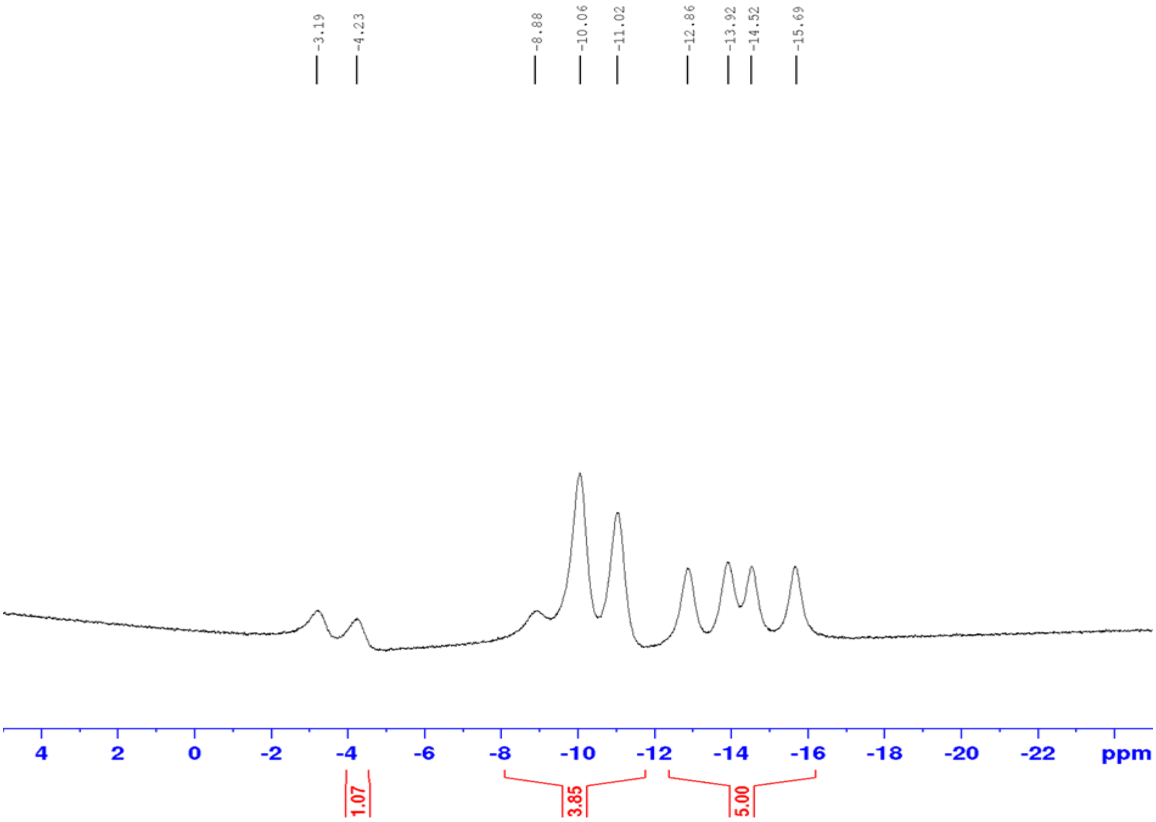

$^{1}H$-$^{1}H$ COSY

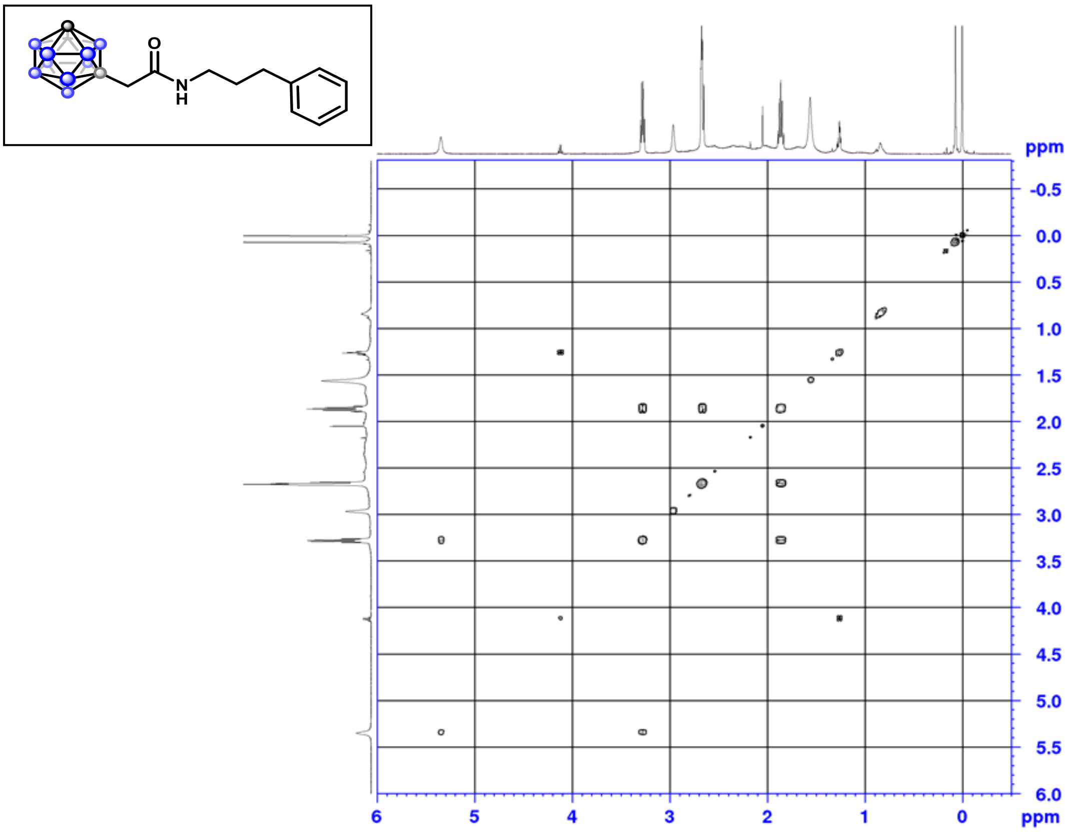

$^{1}H$-$^{13}C$ HSQC

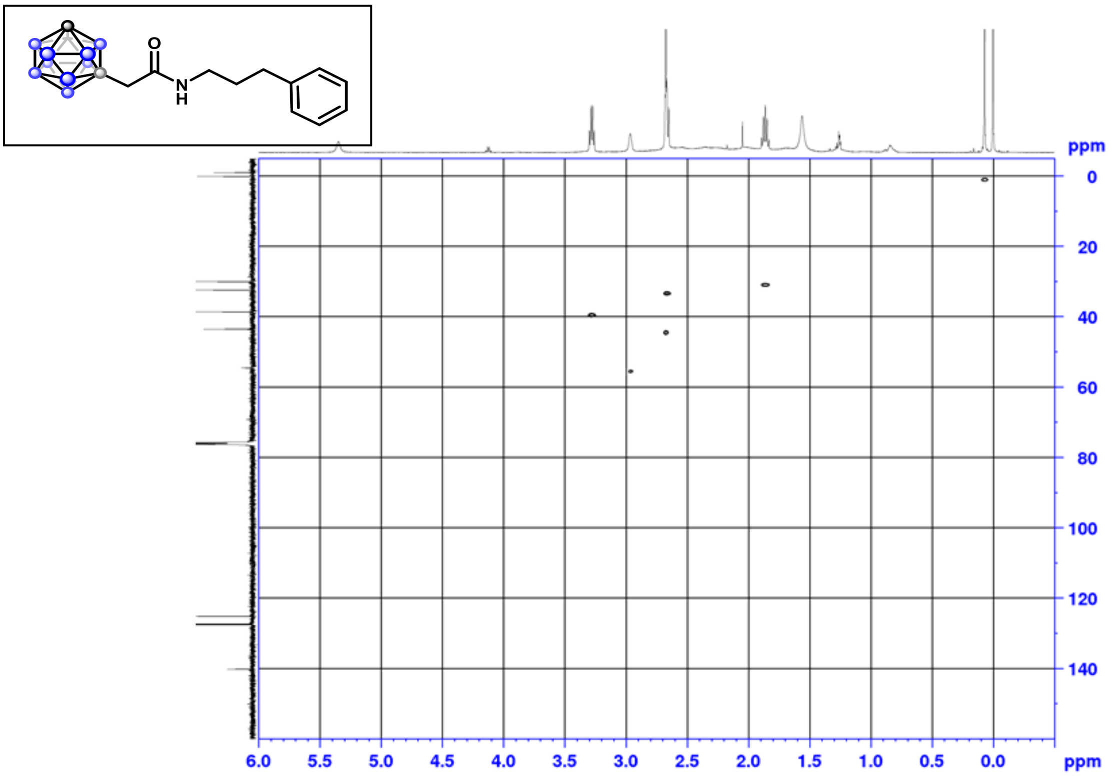

[1]H-[13]C HMBC

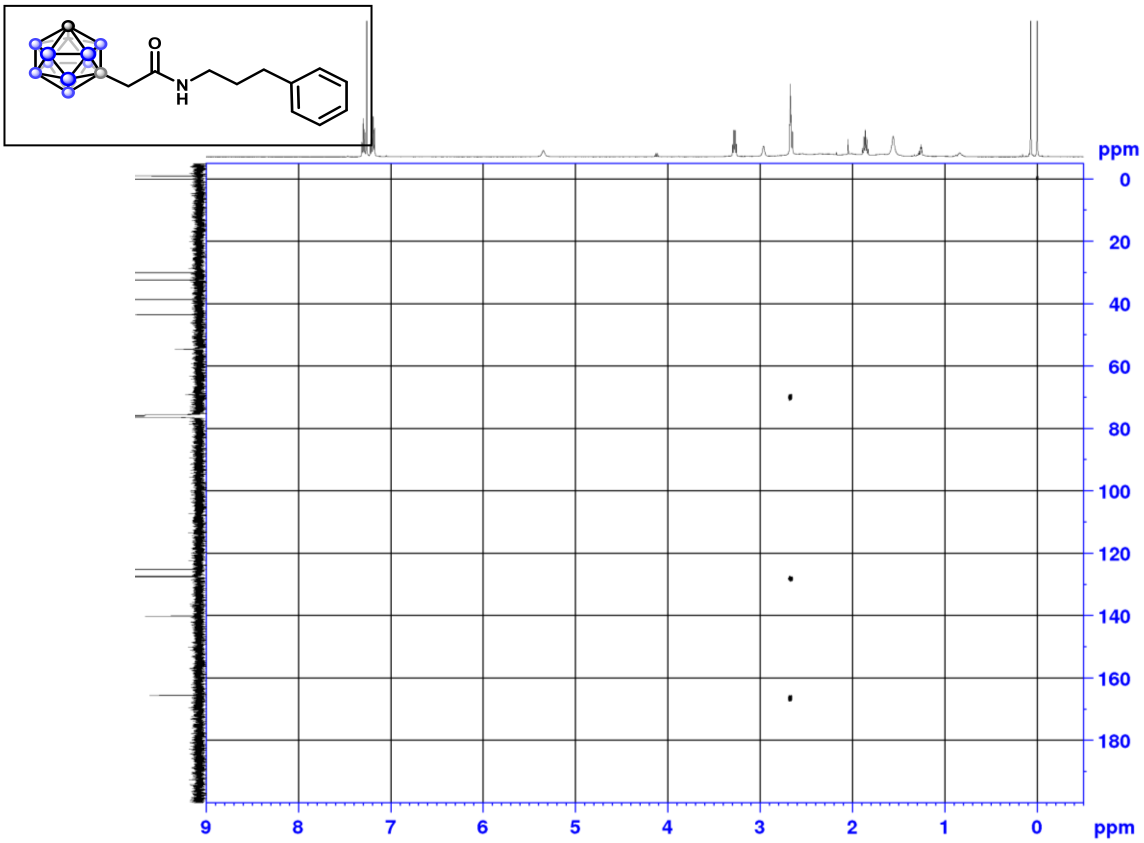

[1]H NMR (500 MHz, $CDCl_3$)

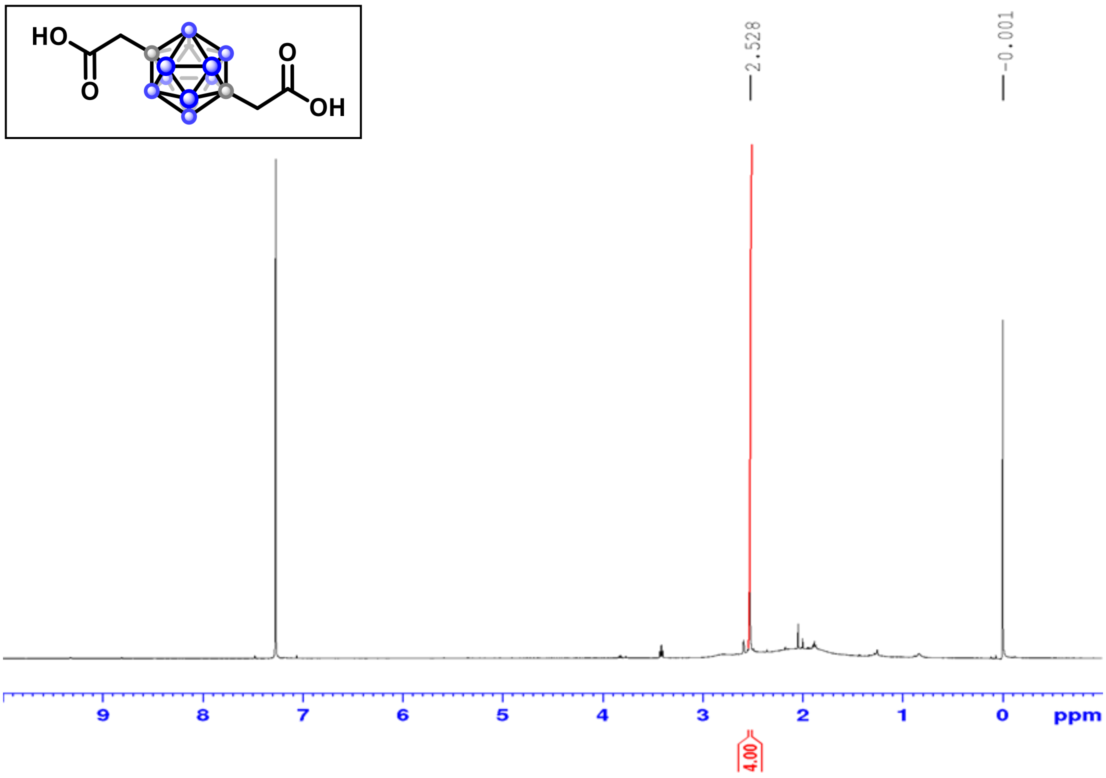

$^{1}H$ NMR (500 MHz, $CDCl_3$)

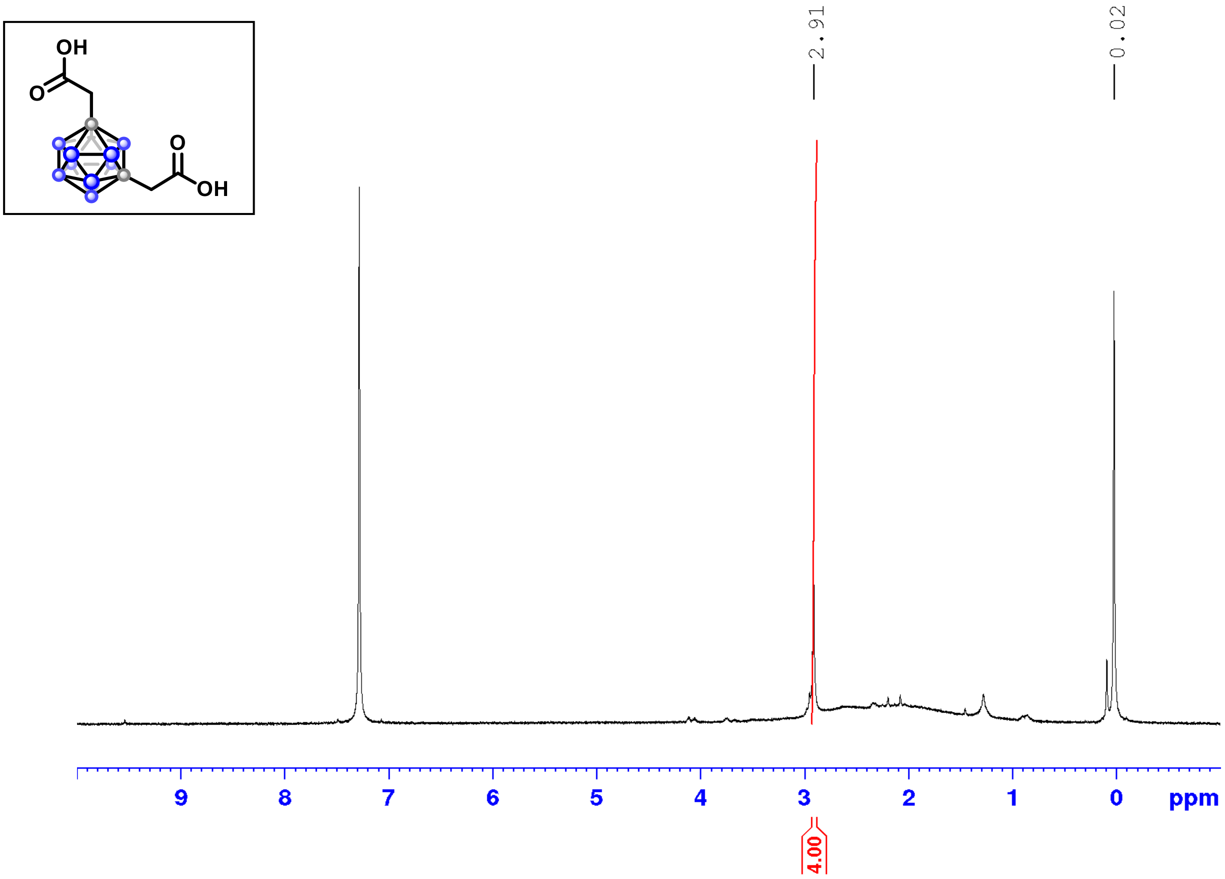

$^{1}H$ NMR (500 MHz, $CDCl_3$)

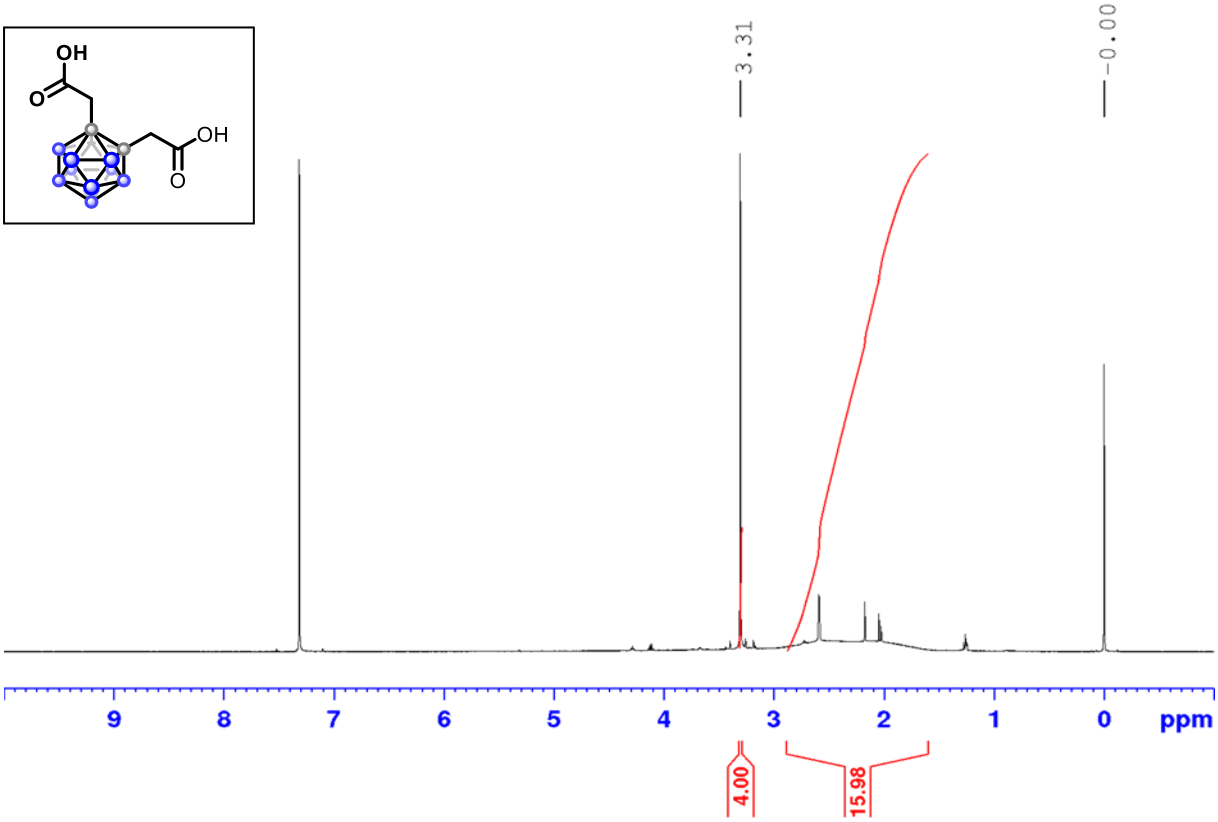

[1]H NMR (500 MHz, $CDCl_3$)

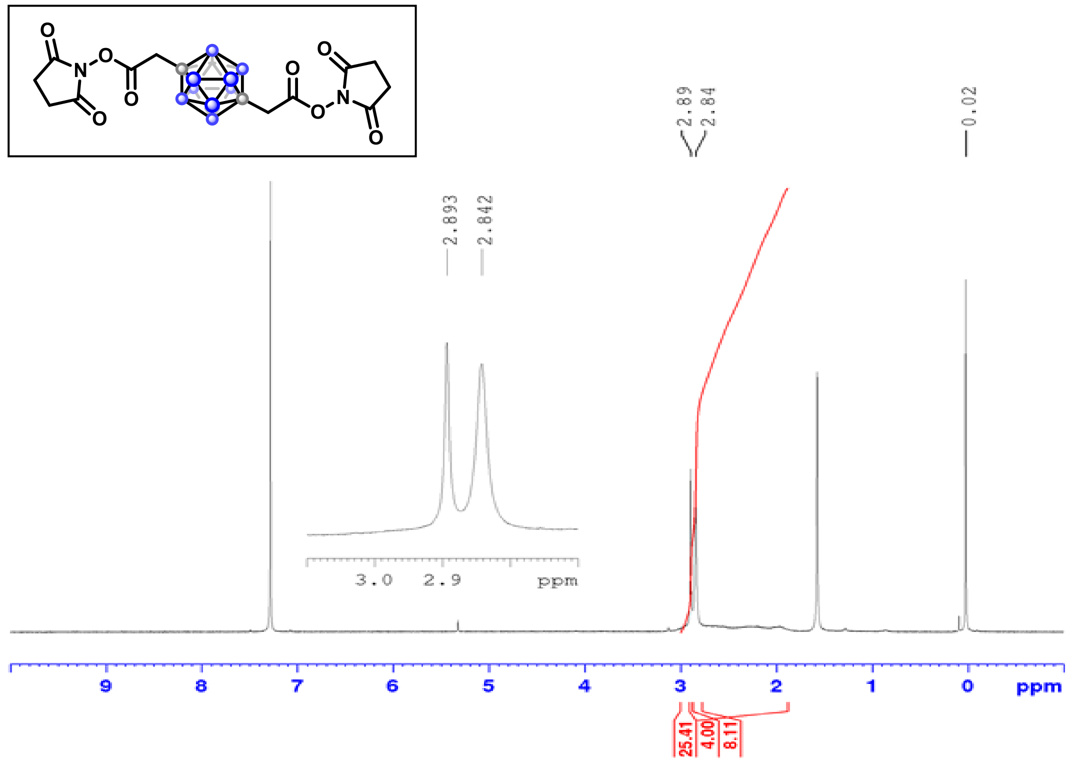

[13]C NMR (126 MHz, $CDCl_3$)

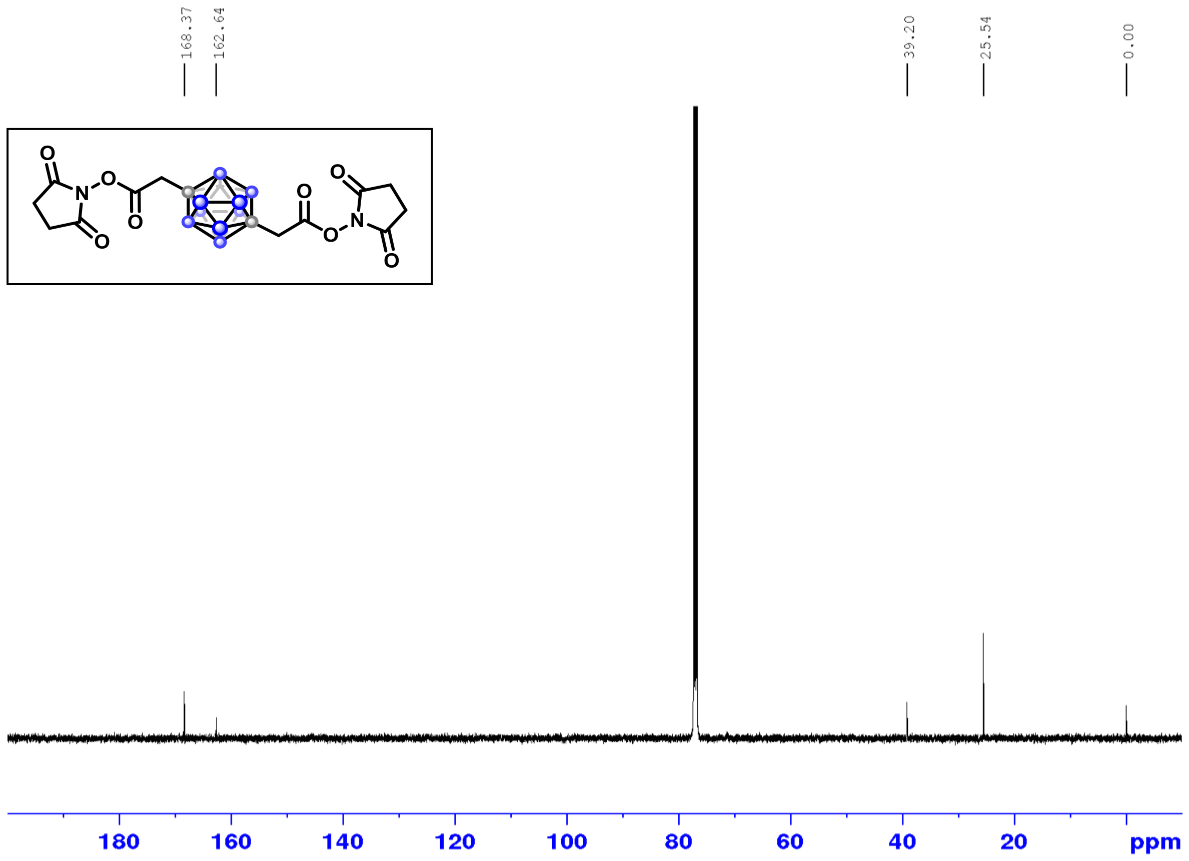

$^{1}H$ NMR (500 MHz, DMSO-$d_6$)

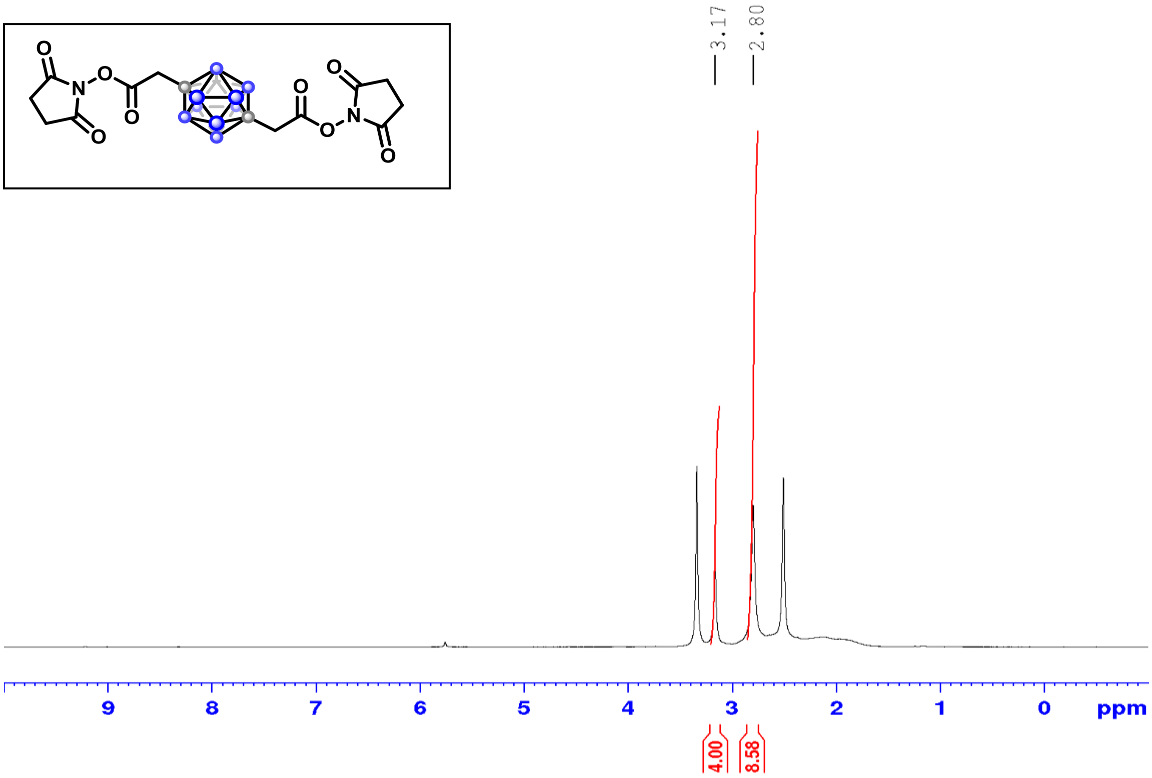

$^{13}C$ NMR (126 MHz, DMSO-$d_6$)

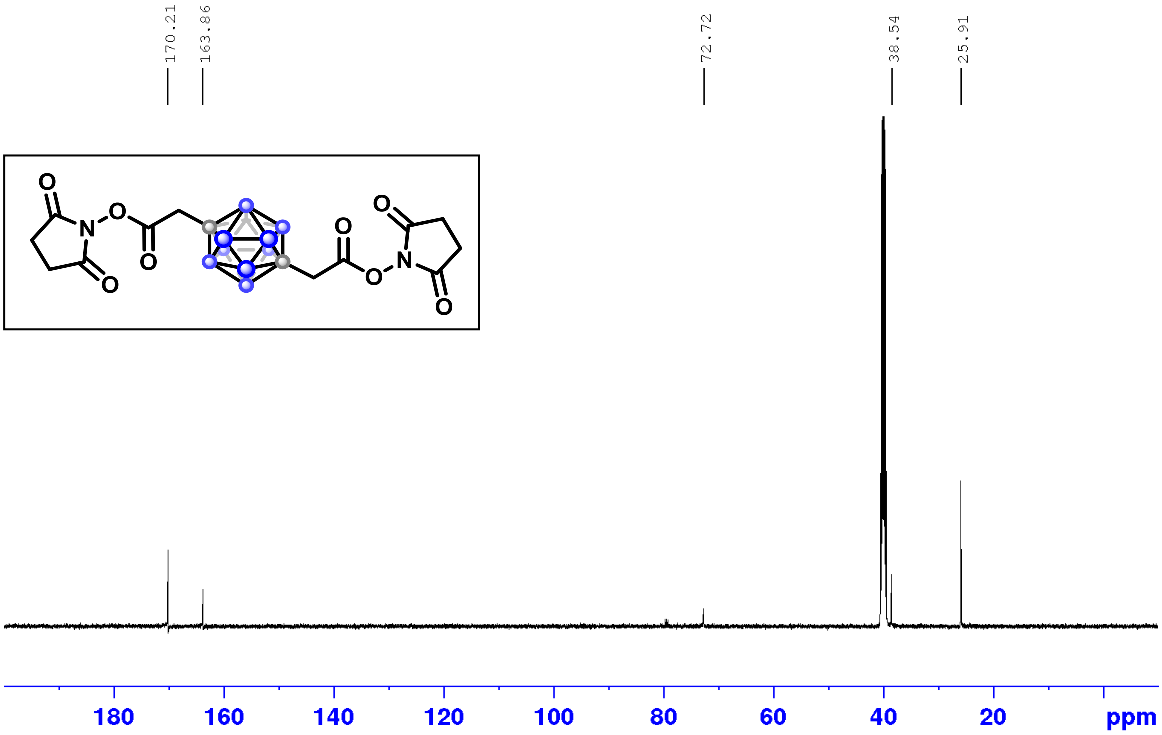

$^{11}B$ NMR (160 MHz, $CDCl_3$)

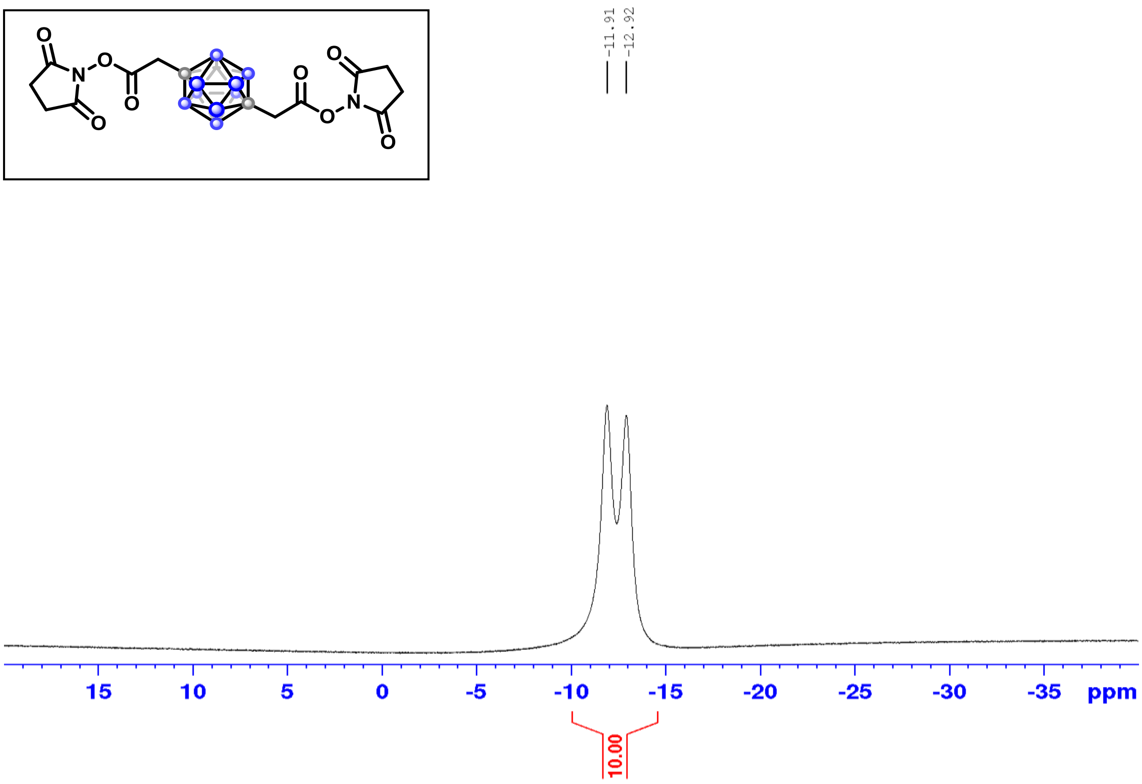

-11.91
-12.92
15
10
5
0
-5
-10
-15
-20
-25
-30
-35
ppm
10.00

$^{1}H$-$^{1}H$ COSY

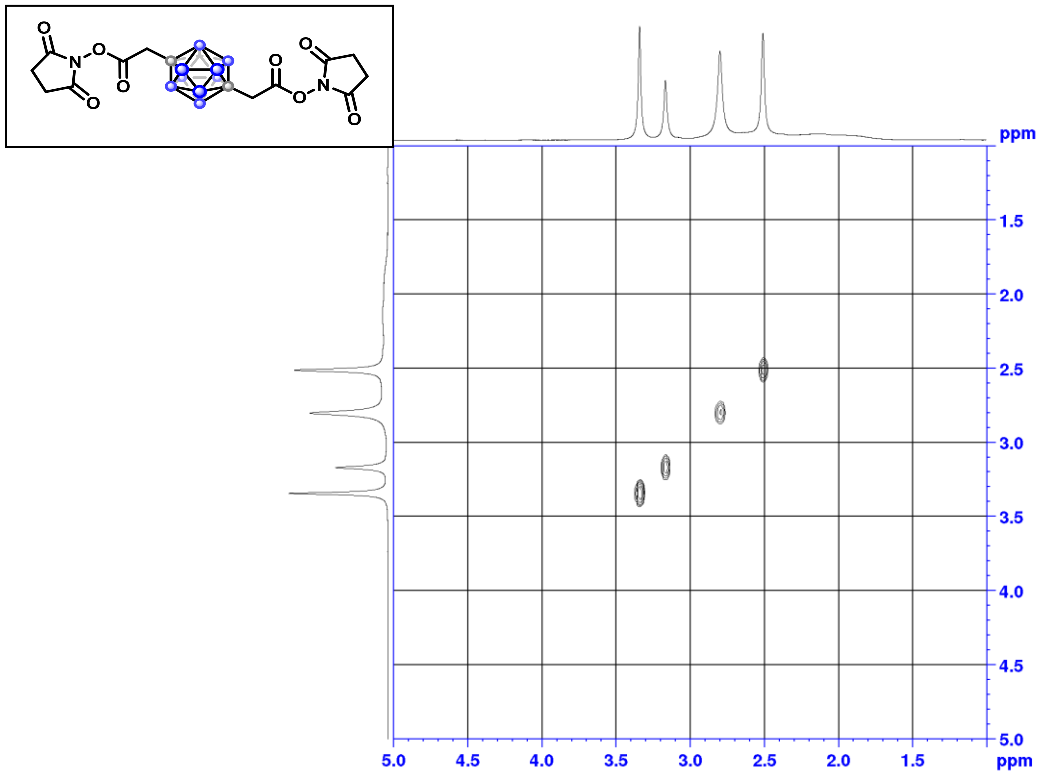

ppm
1.5
2.0
2.5
3.0
3.5
4.0
4.5
5.0
5.0
4.5
4.0
3.5
3.0
2.5
2.0
1.5
ppm

$^{1}H$-$^{13}C$ HSQC

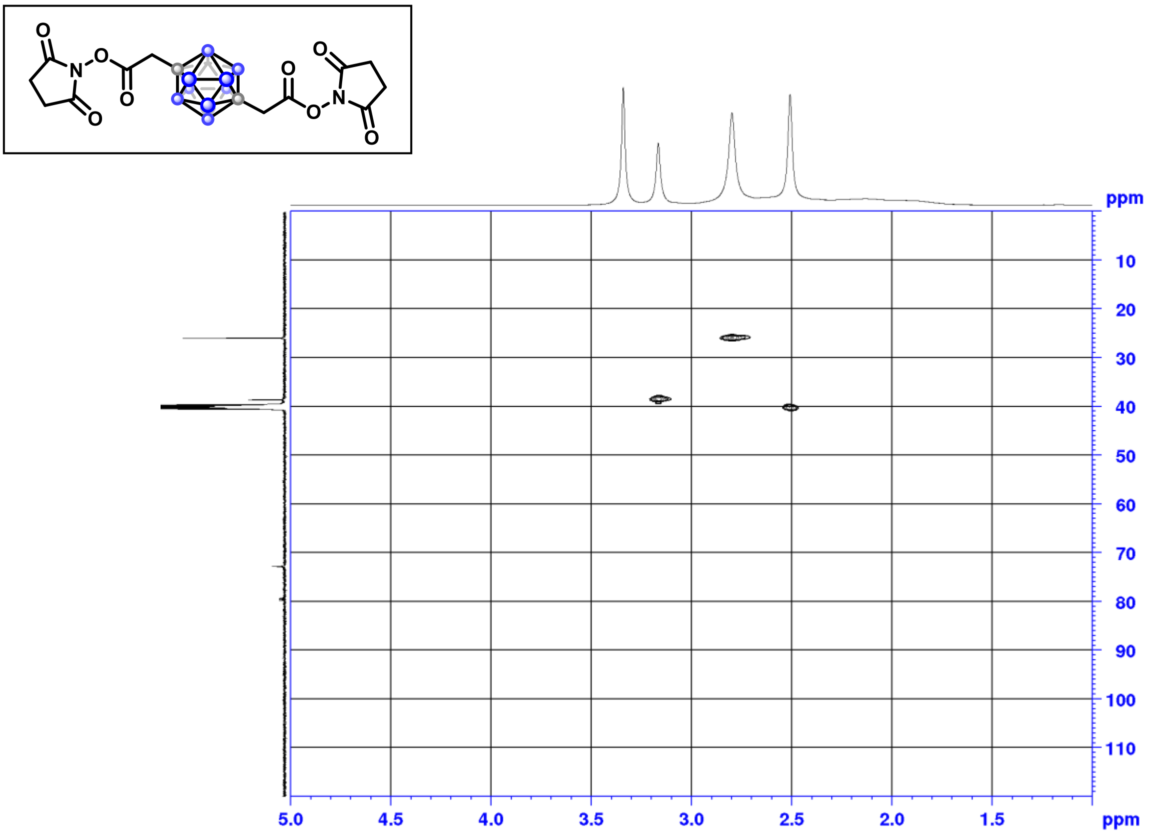

$^{1}H$-$^{13}C$ HMBC

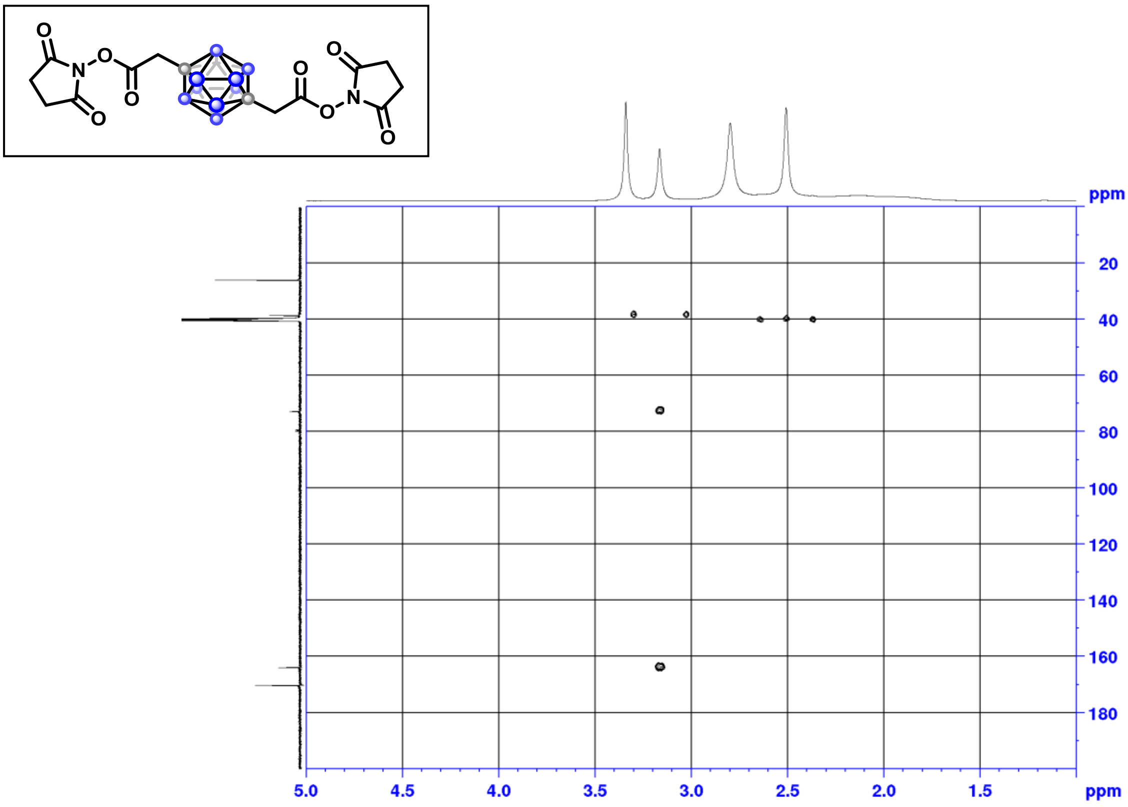

$^{1}H$ NMR (500 MHz, $CDCl_3$)

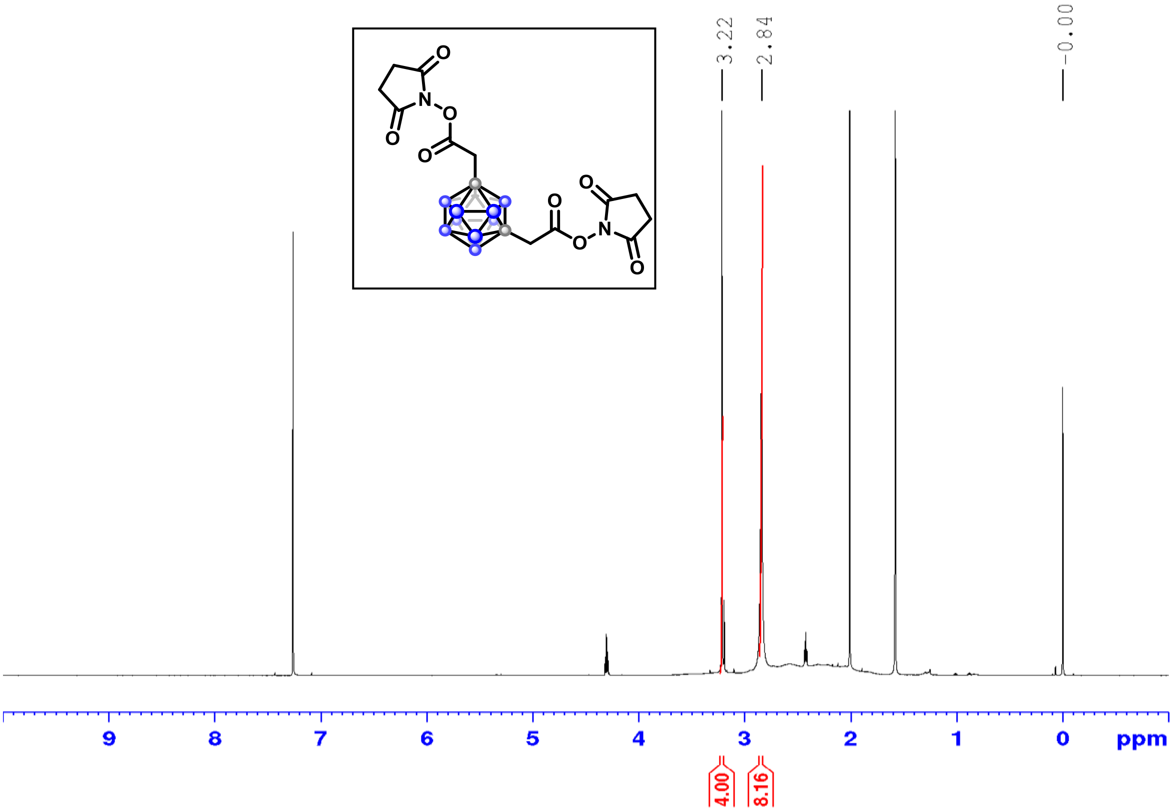

$^{13}C$ NMR (126 MHz, $CDCl_3$)

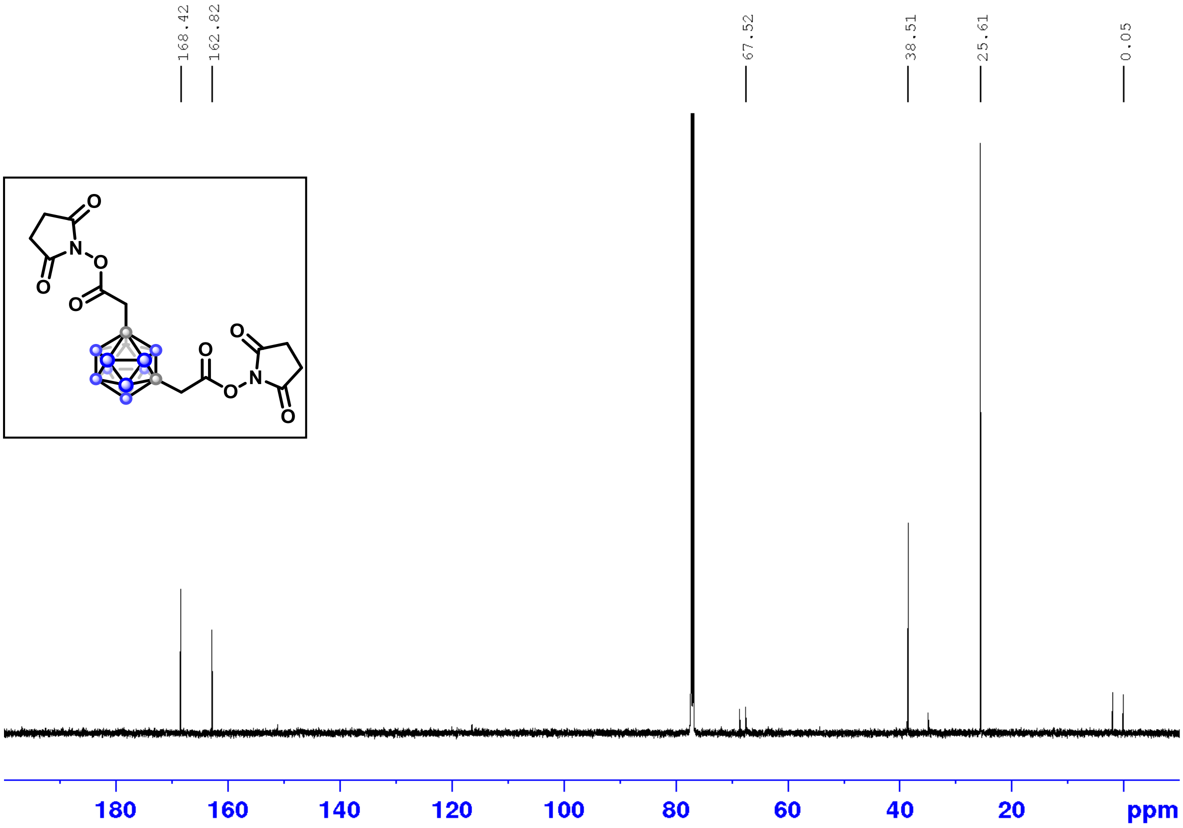

$^{11}B$ NMR (160 MHz, $CDCl_3$)

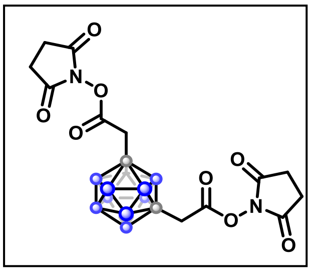

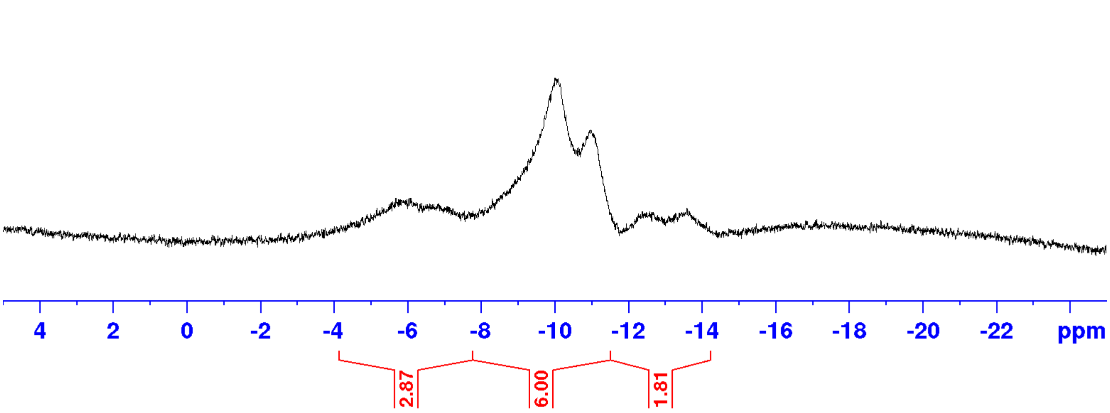

$^{1}H$-$^{1}H$ COSY

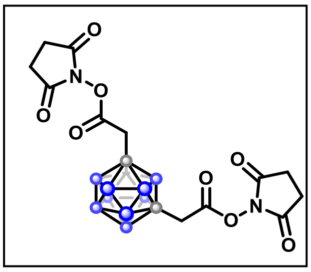

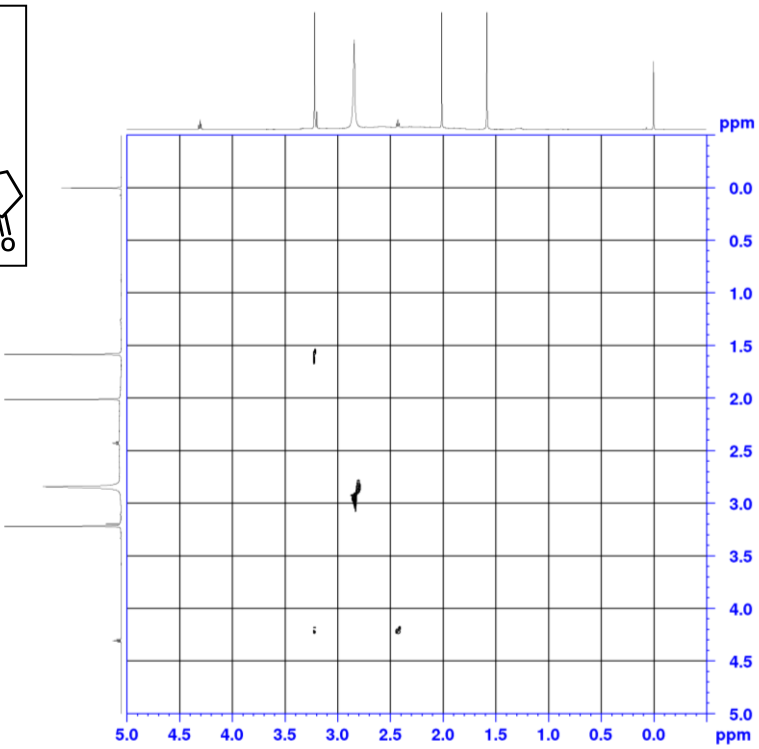

$^{1}H$-$^{13}C$ HSQC

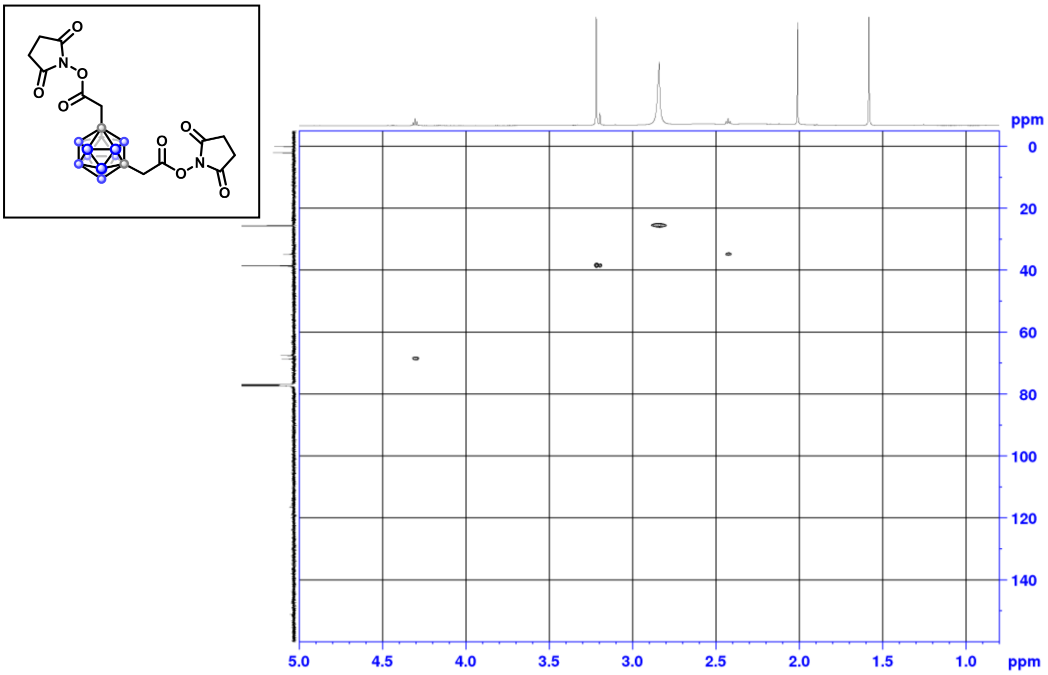

$^{1}H$-$^{13}C$ HMBC

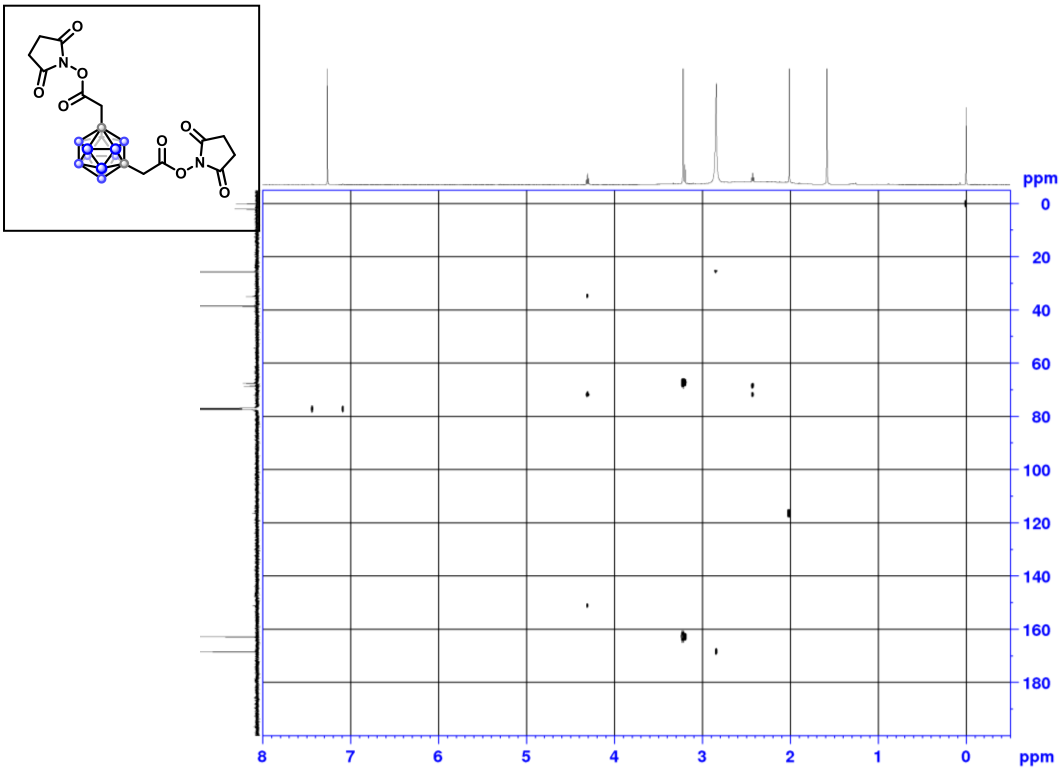

$^1$H NMR (500 MHz, $CDCl_3$)

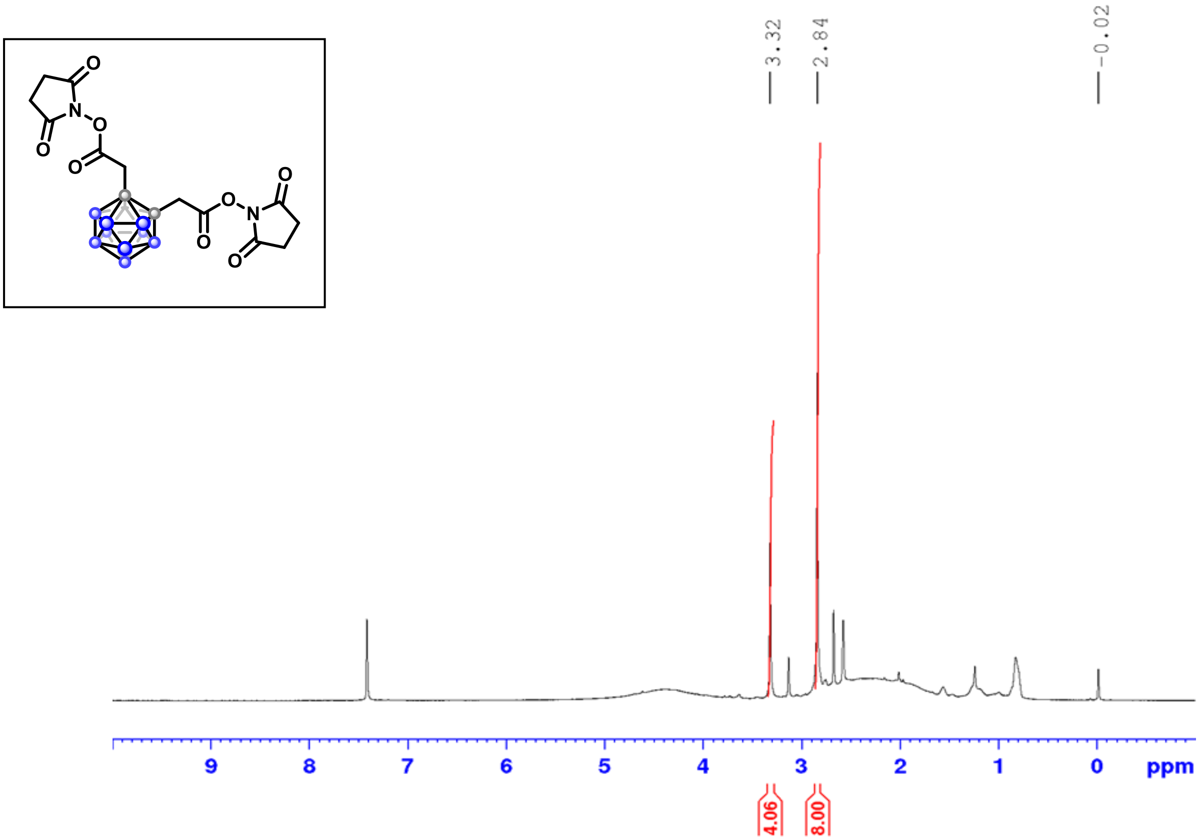

$^1$H NMR (500 MHz, $CDCl_3$) after 10 hours

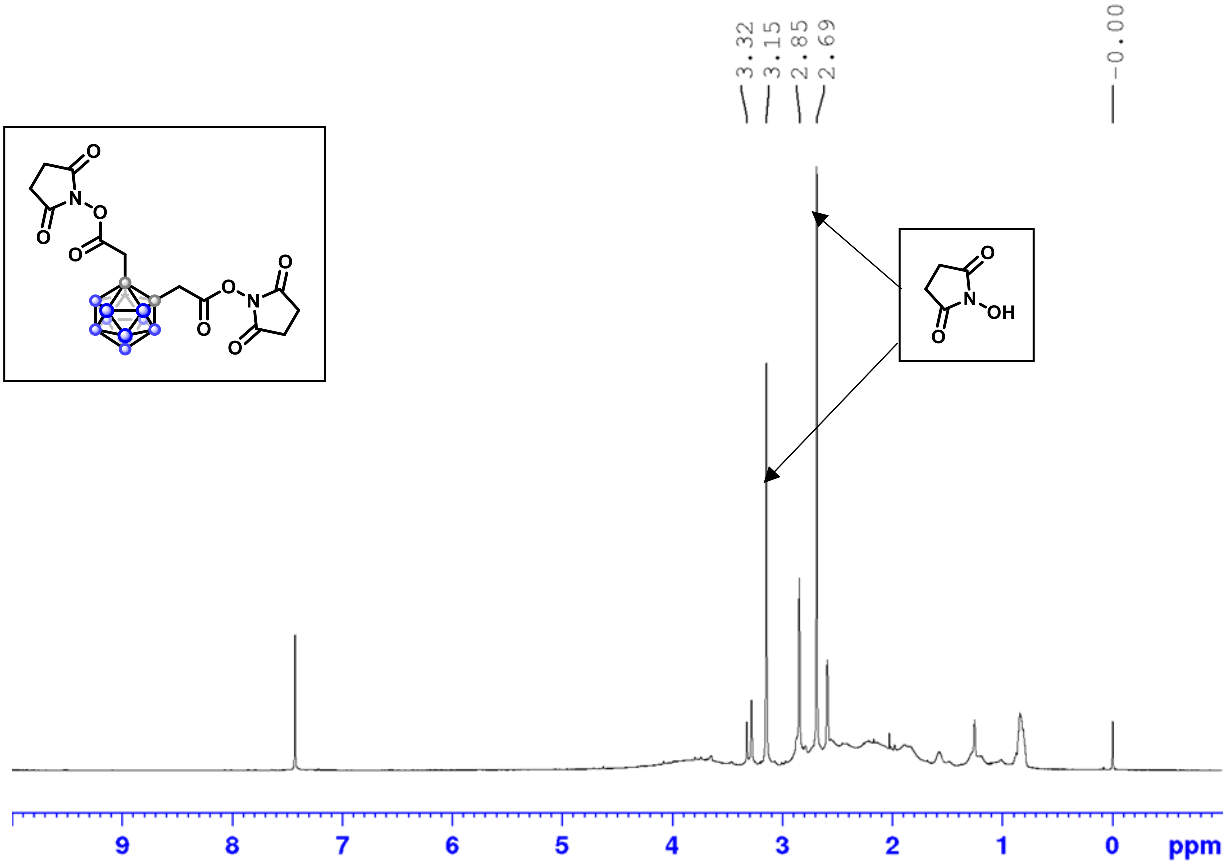

$^{13}C$ NMR (126 MHz, $CDCl_3$)

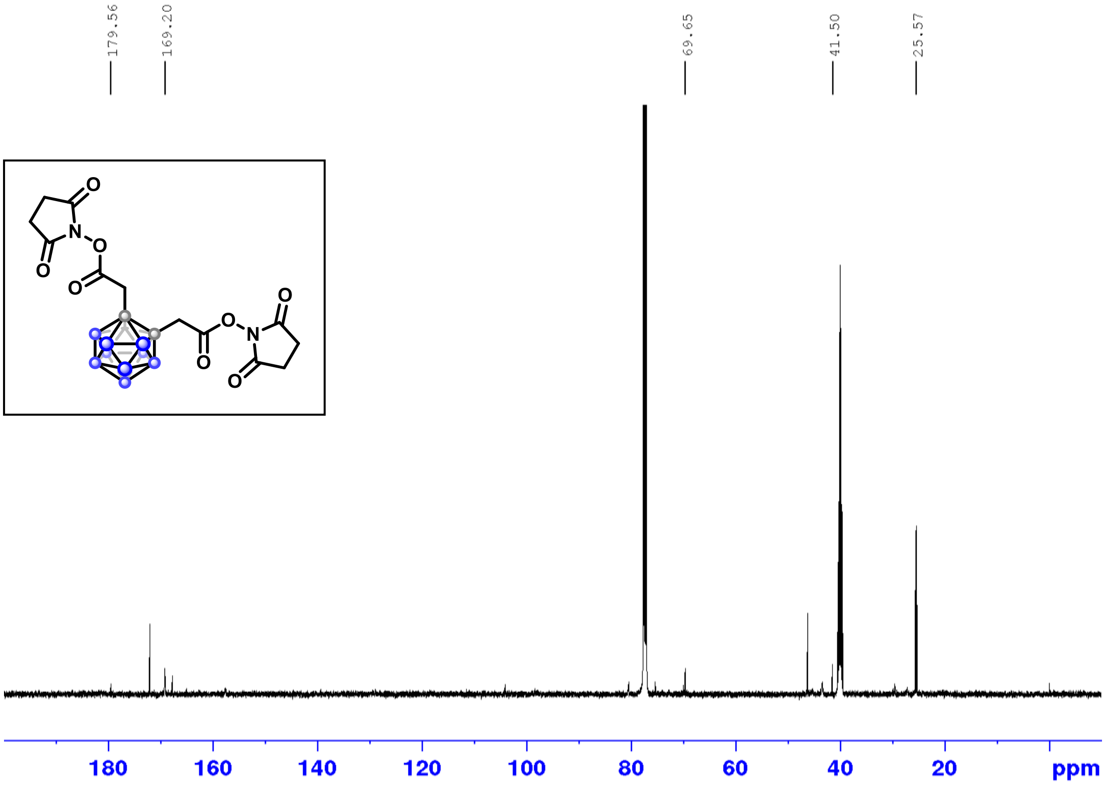

$^{11}B$ NMR (160 MHz, $CDCl_3$)

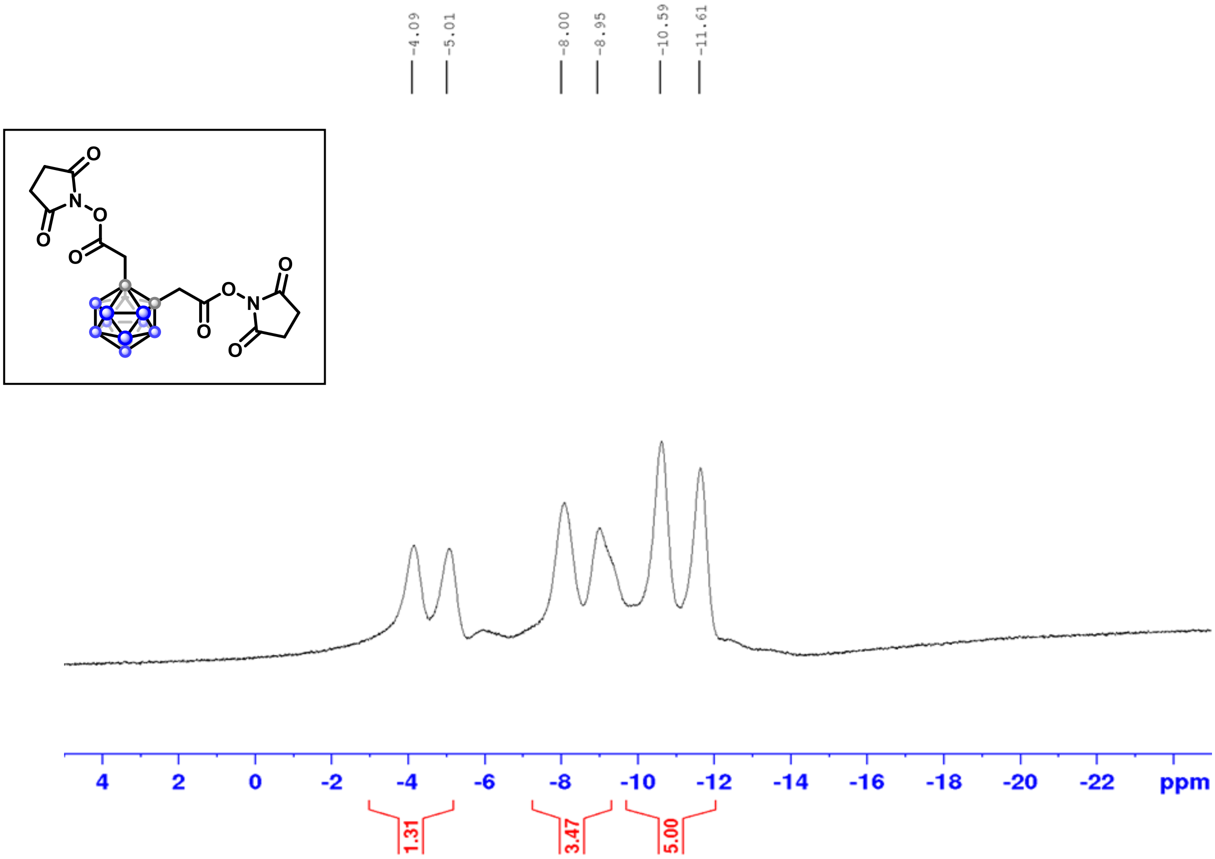

$^{1}H$-$^{1}H$ COSY

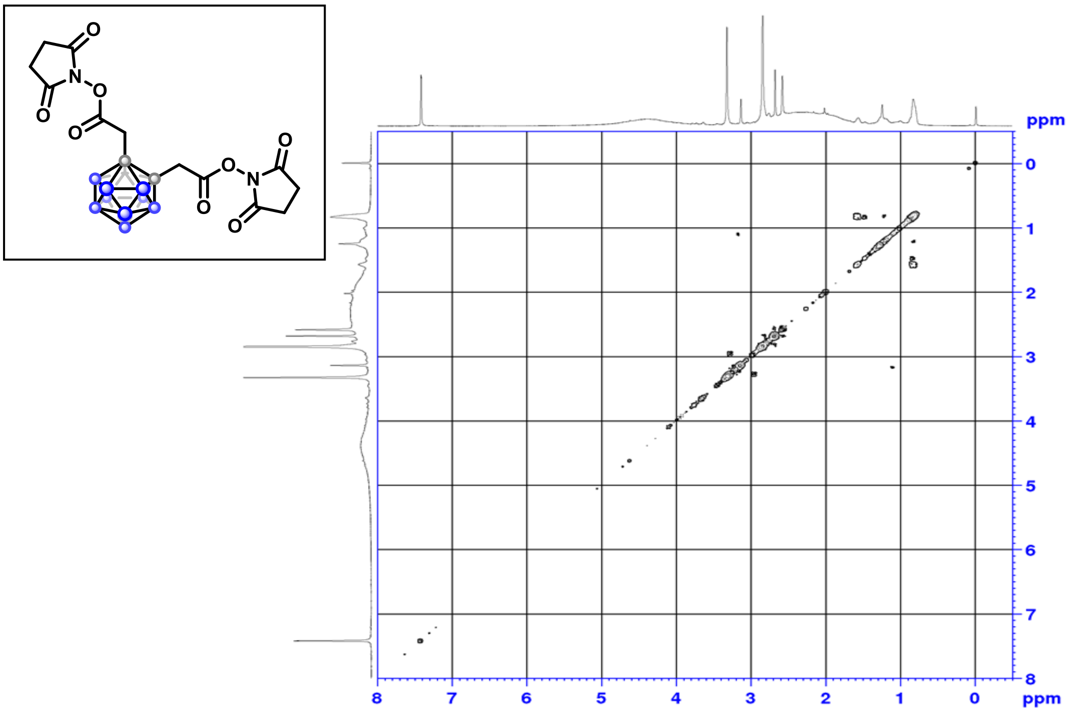

$^{1}H$-$^{13}C$ HSQC

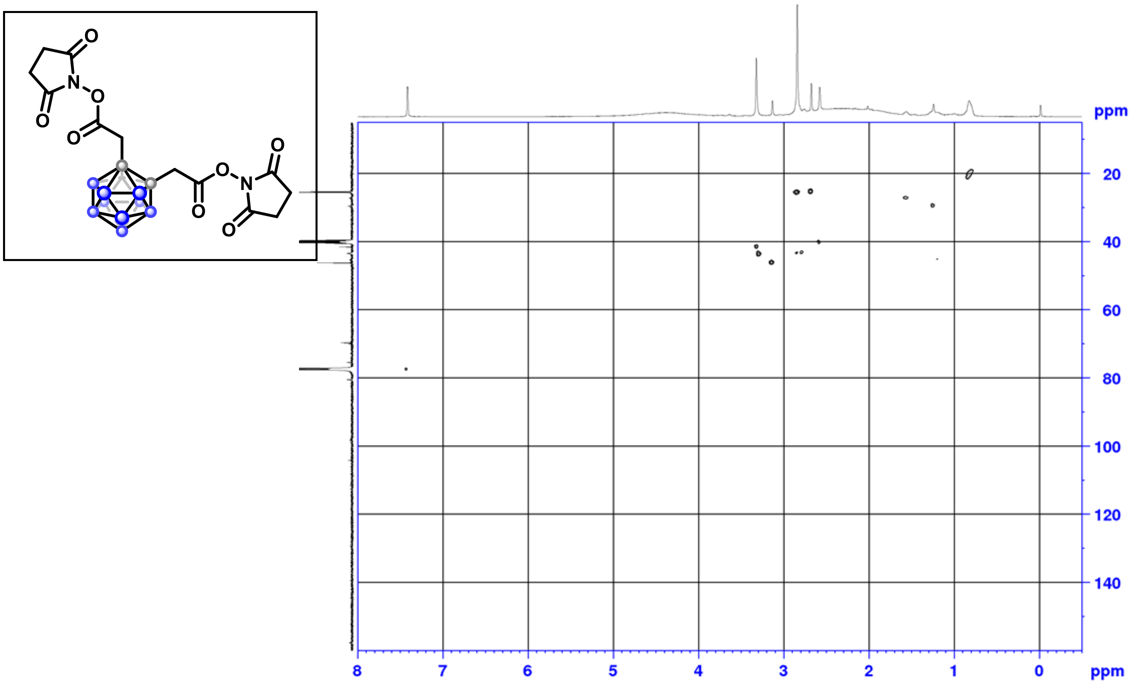

[1]H-[13]C HMBC

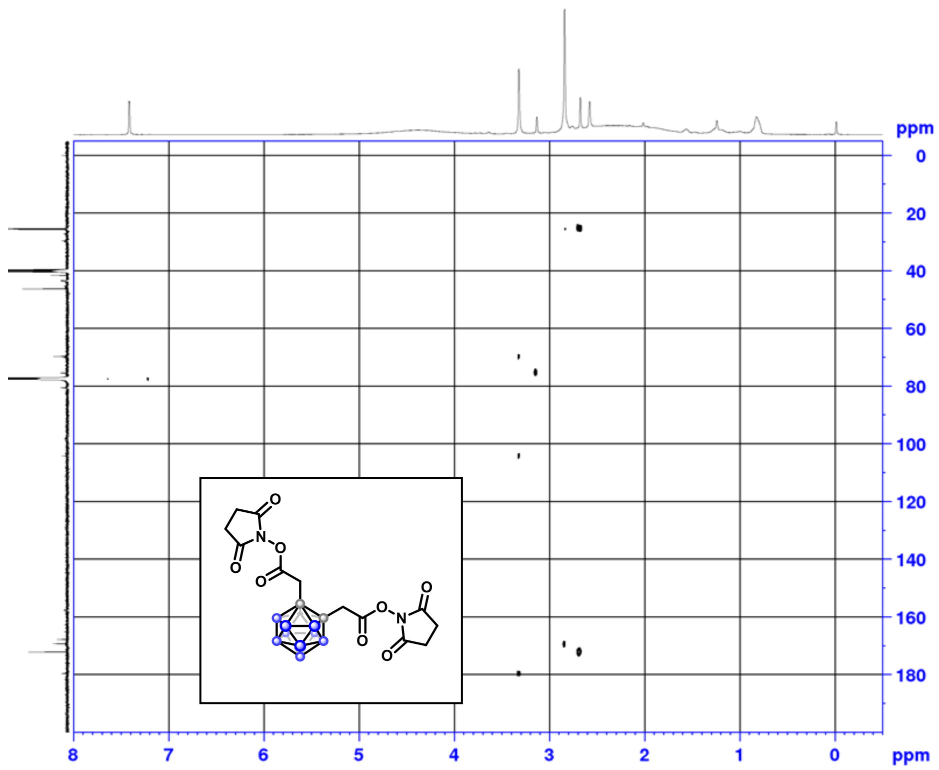

[1]H NMR (500 MHz, $CDCl_3$)

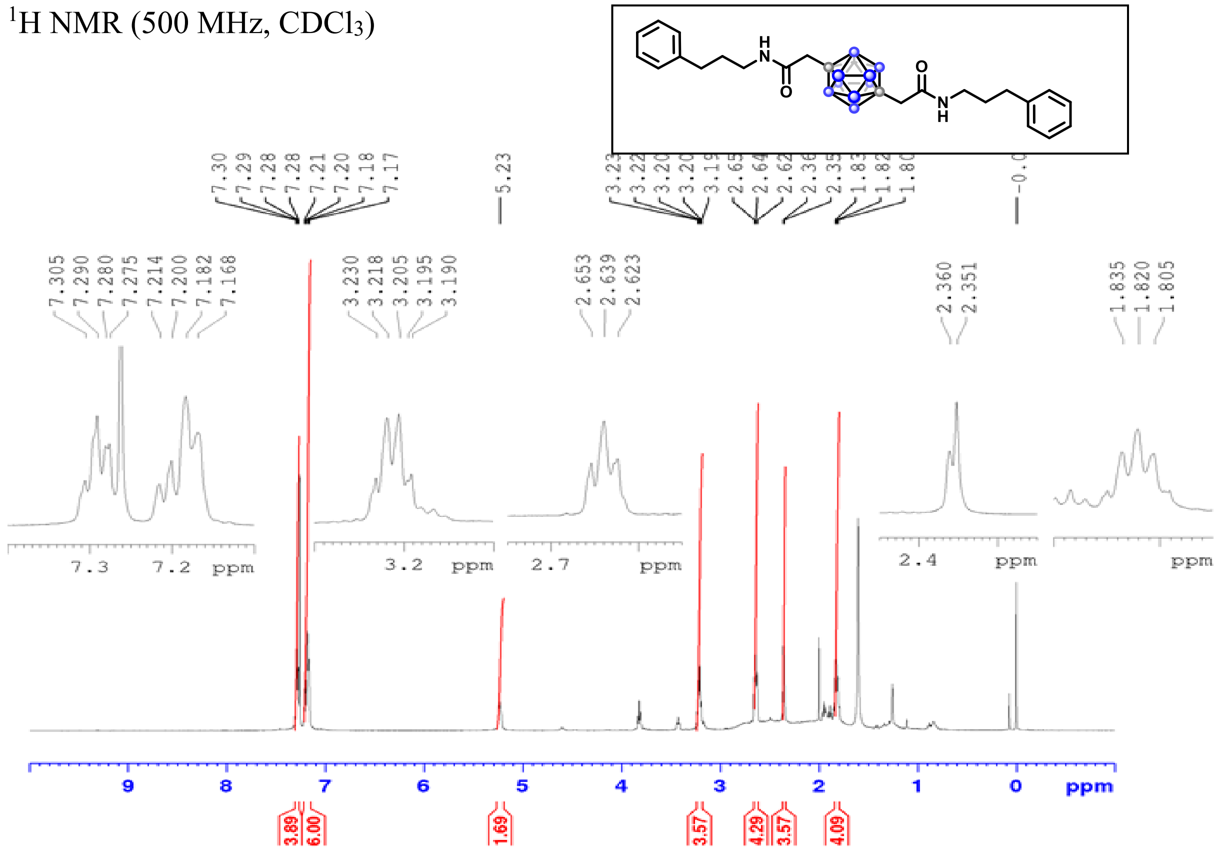

$^{13}C$ NMR (126 MHz, $CDCl_3$)

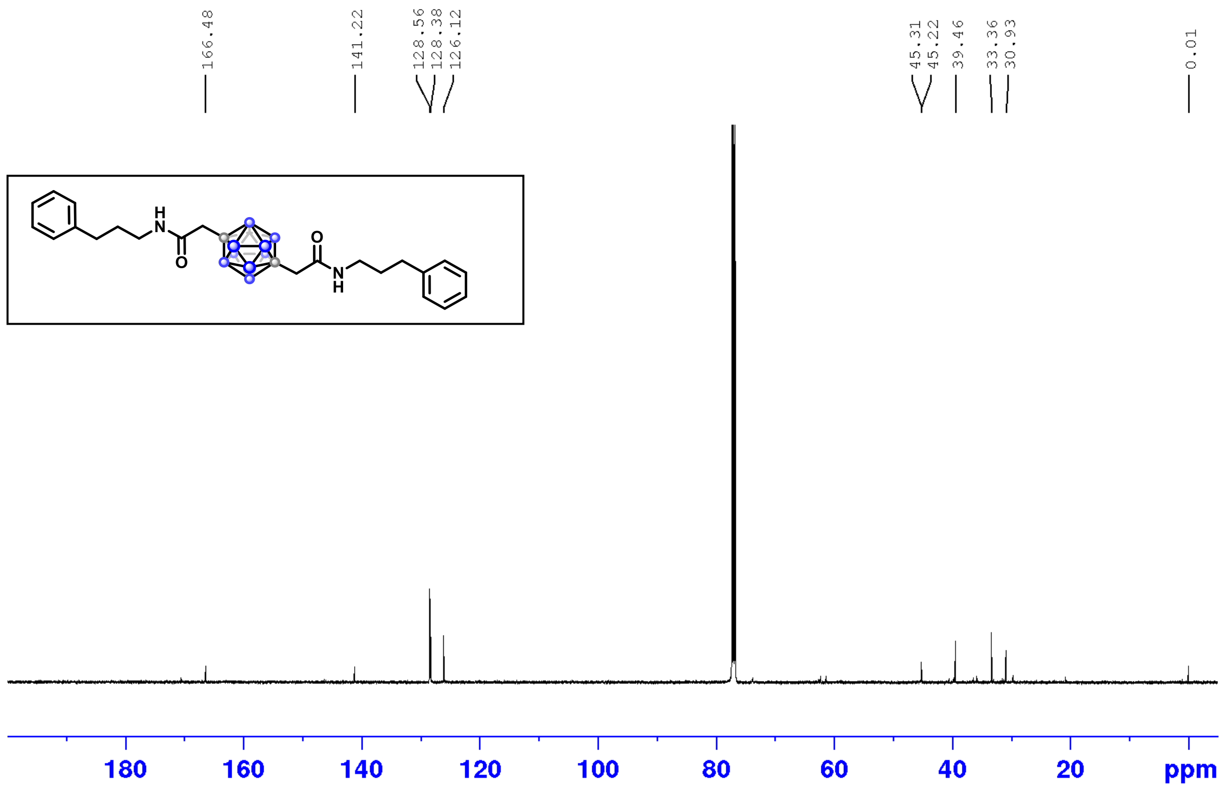

$^{11}B$ NMR (160 MHz, $CDCl_3$)

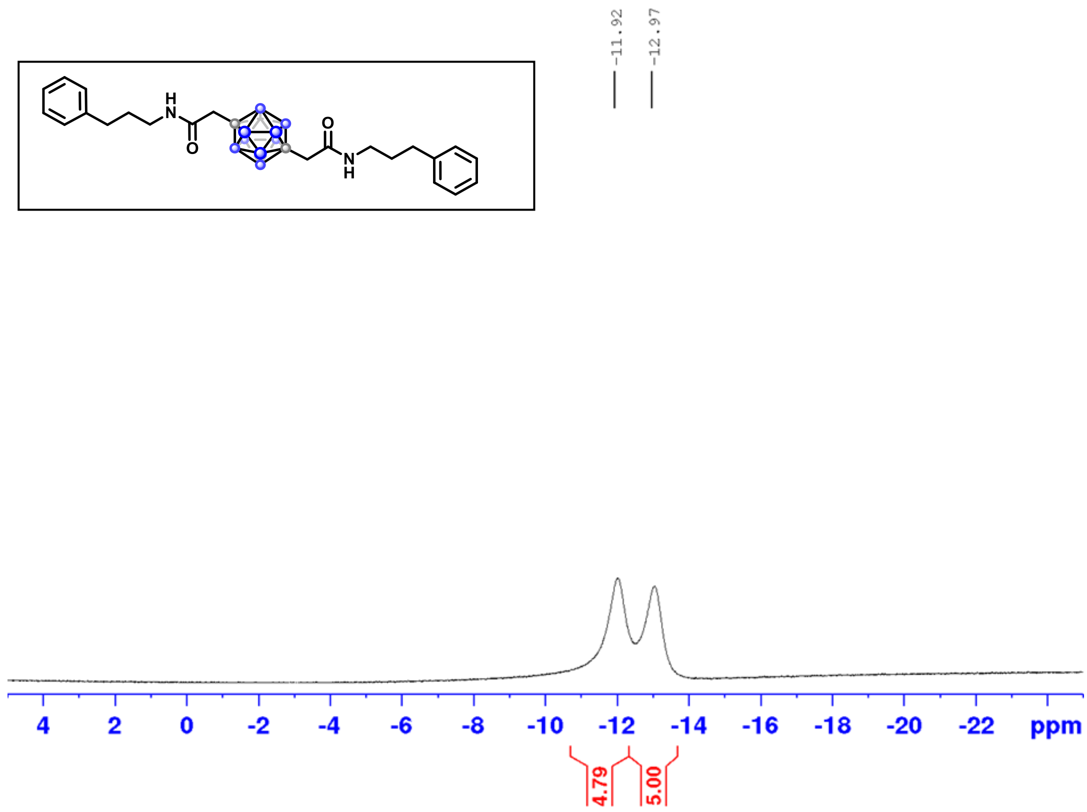

$^{1}H$-$^{1}H$ COSY

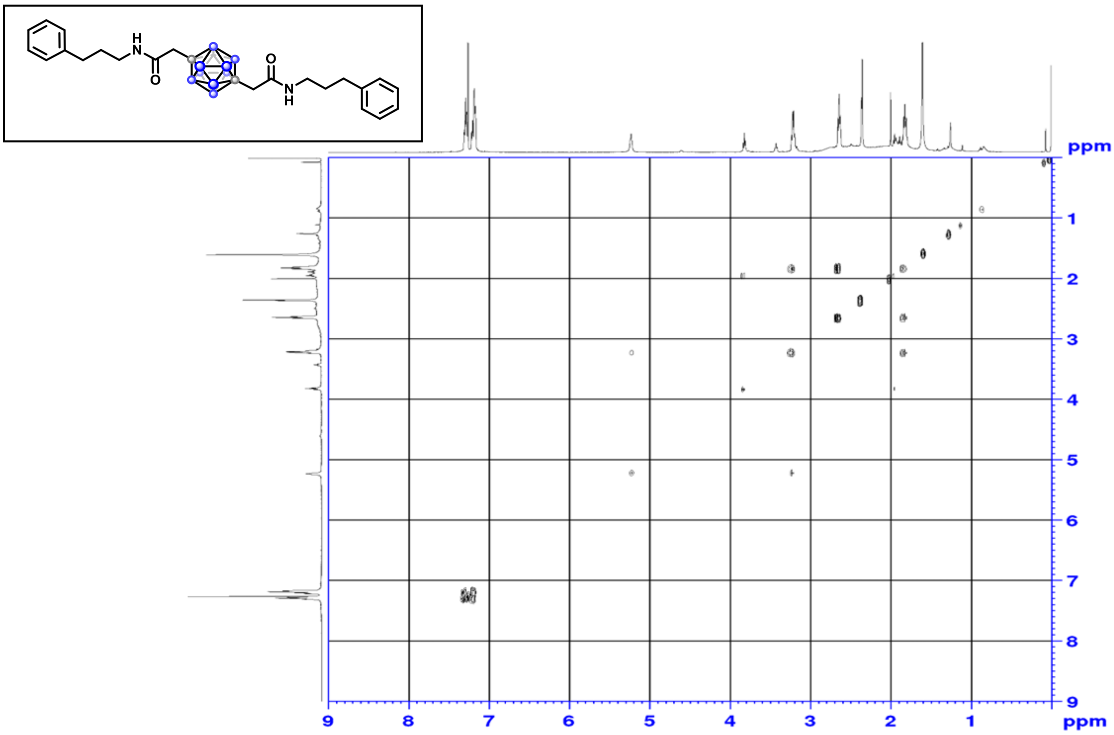

$^{1}H$-$^{13}C$ HSQC

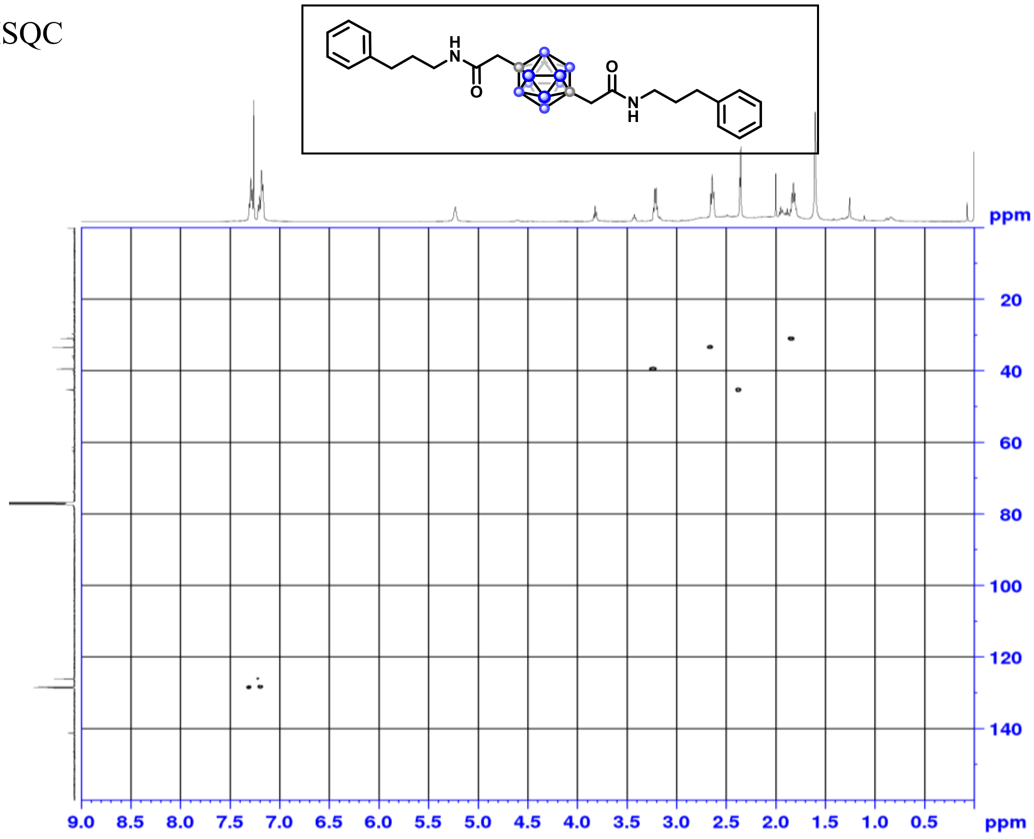

[1]H-[13]C HMBC

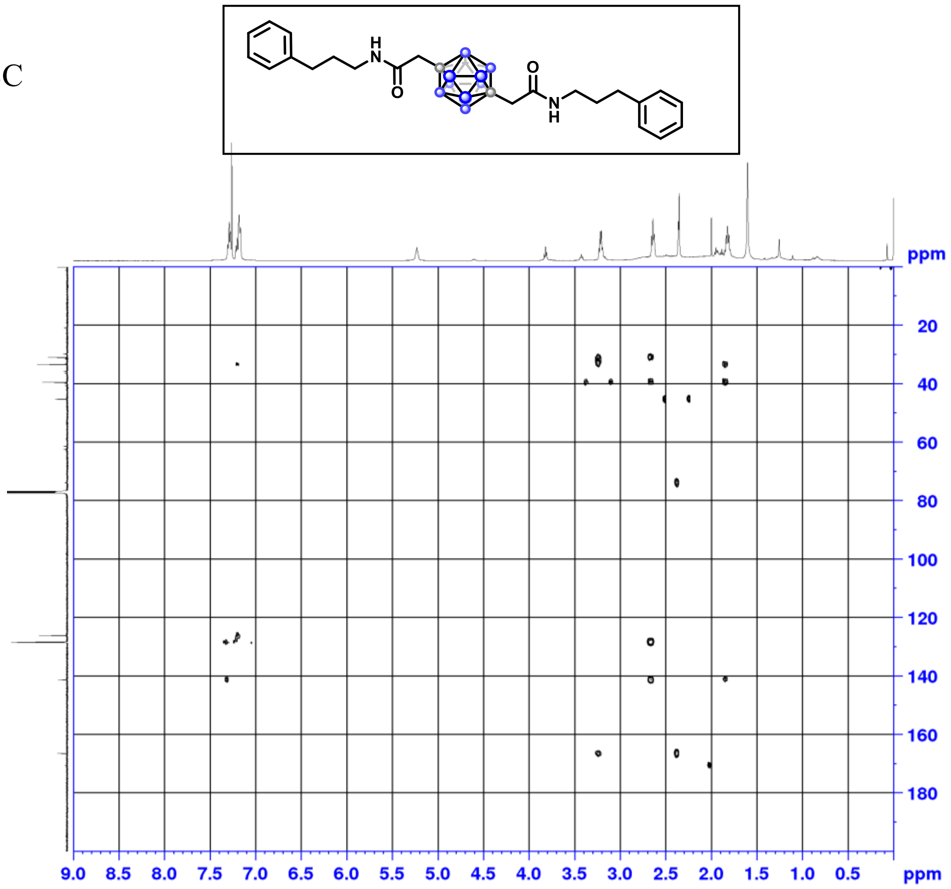

[1]H NMR (500 MHz, $CDCl_3$)

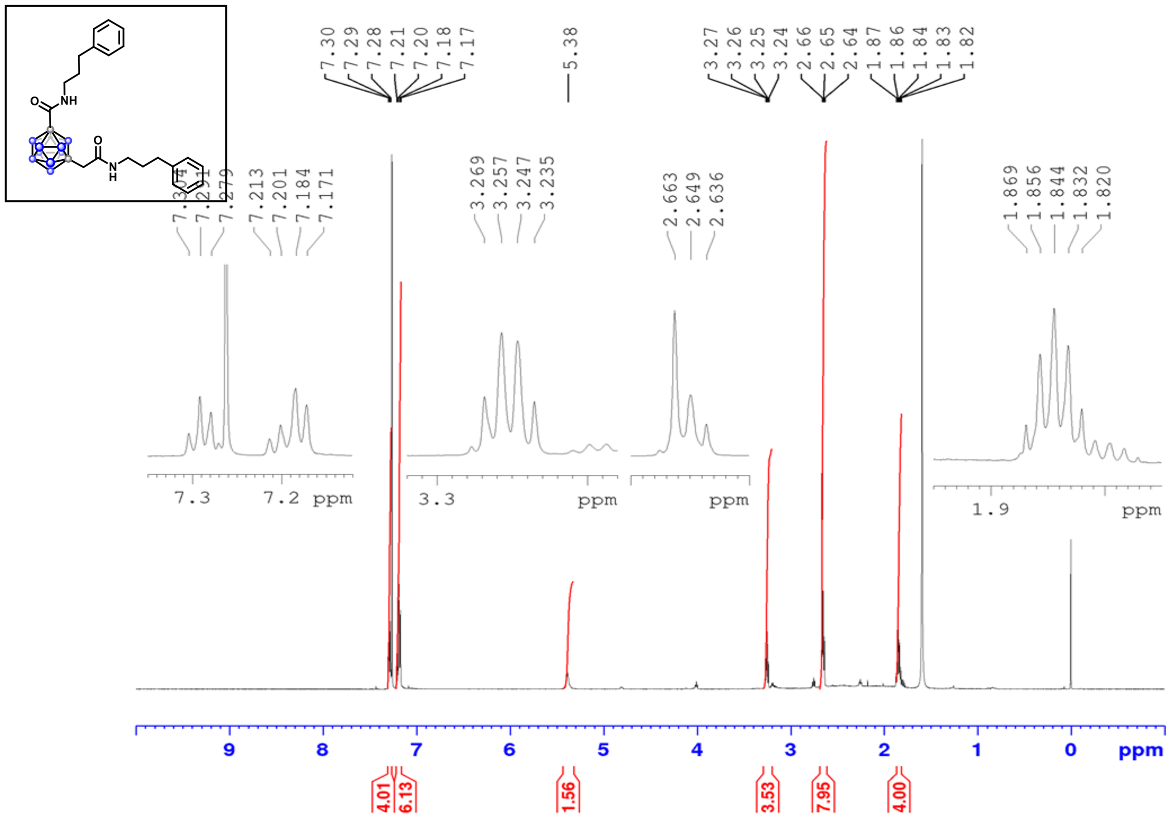

$^{13}C$ NMR (126 MHz, $CDCl_3$)

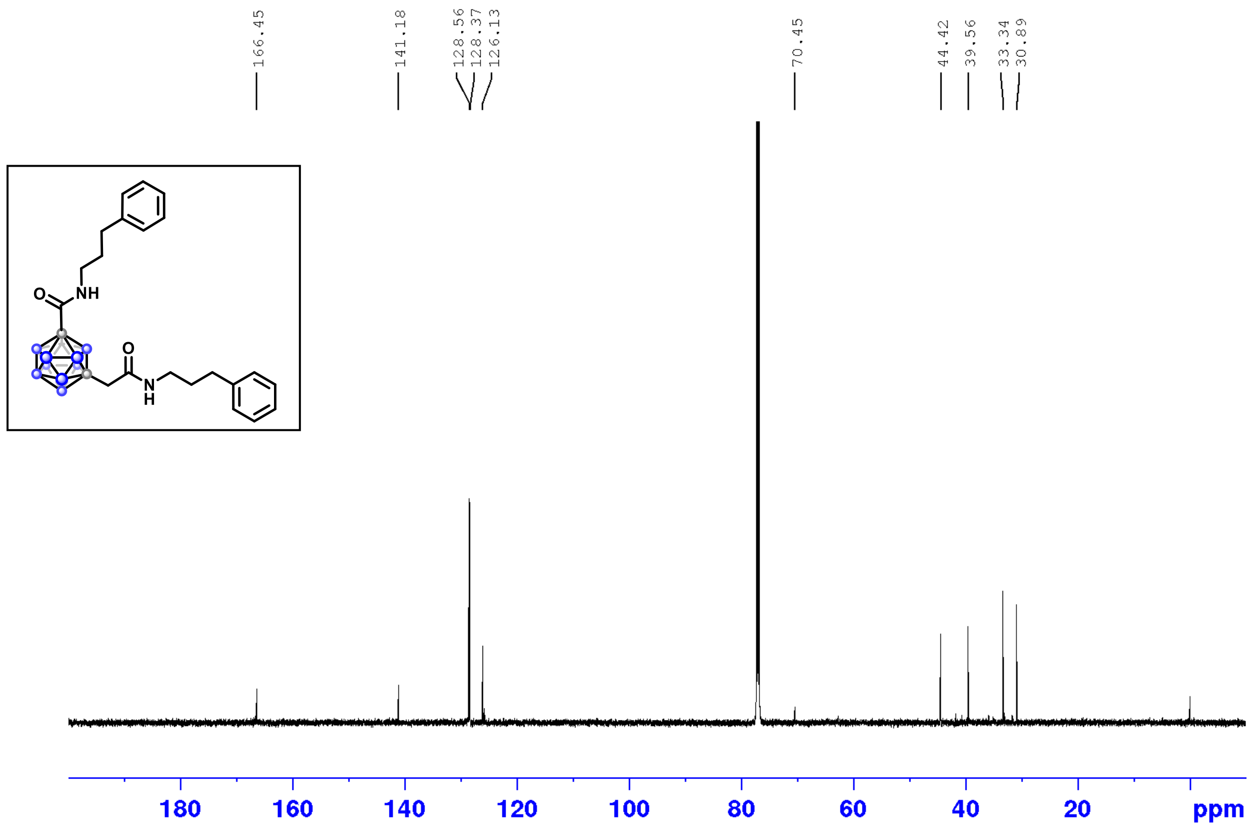

$^{11}B$ NMR (160 MHz, $CDCl_3$)

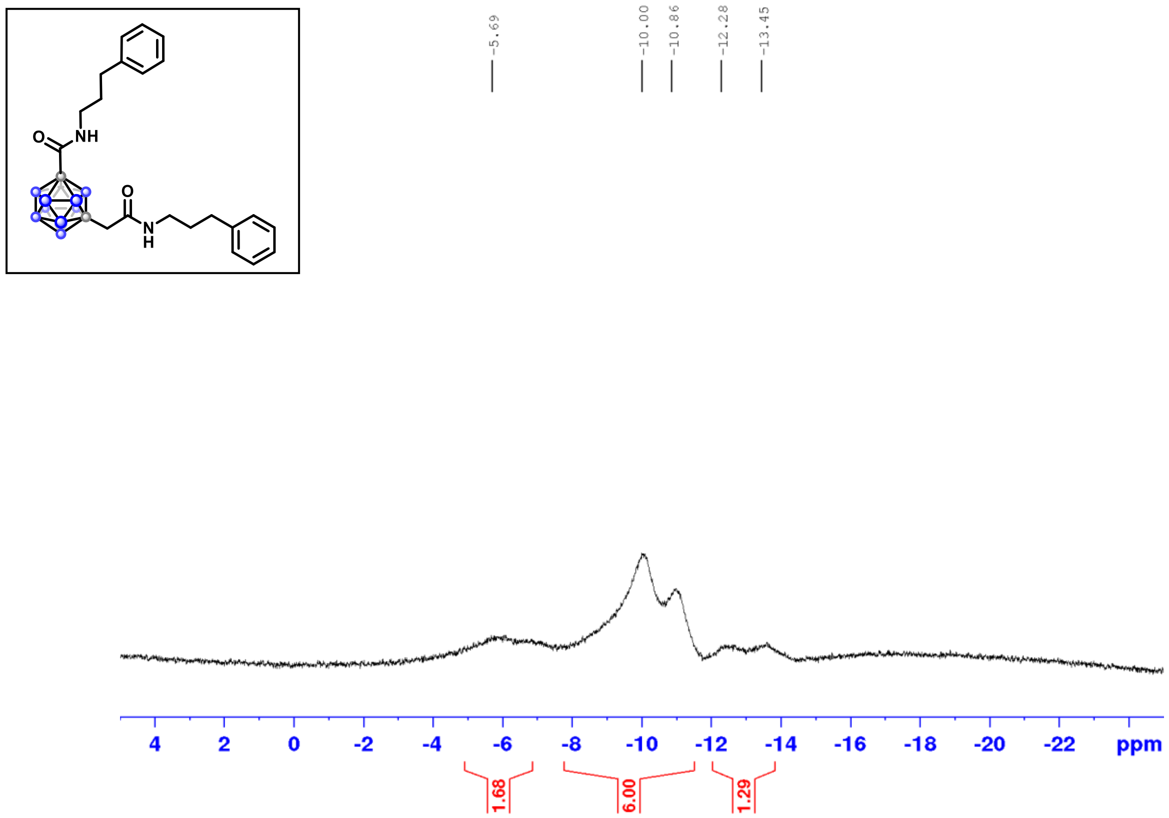

$^{1}H$-$^{1}H$ COSY

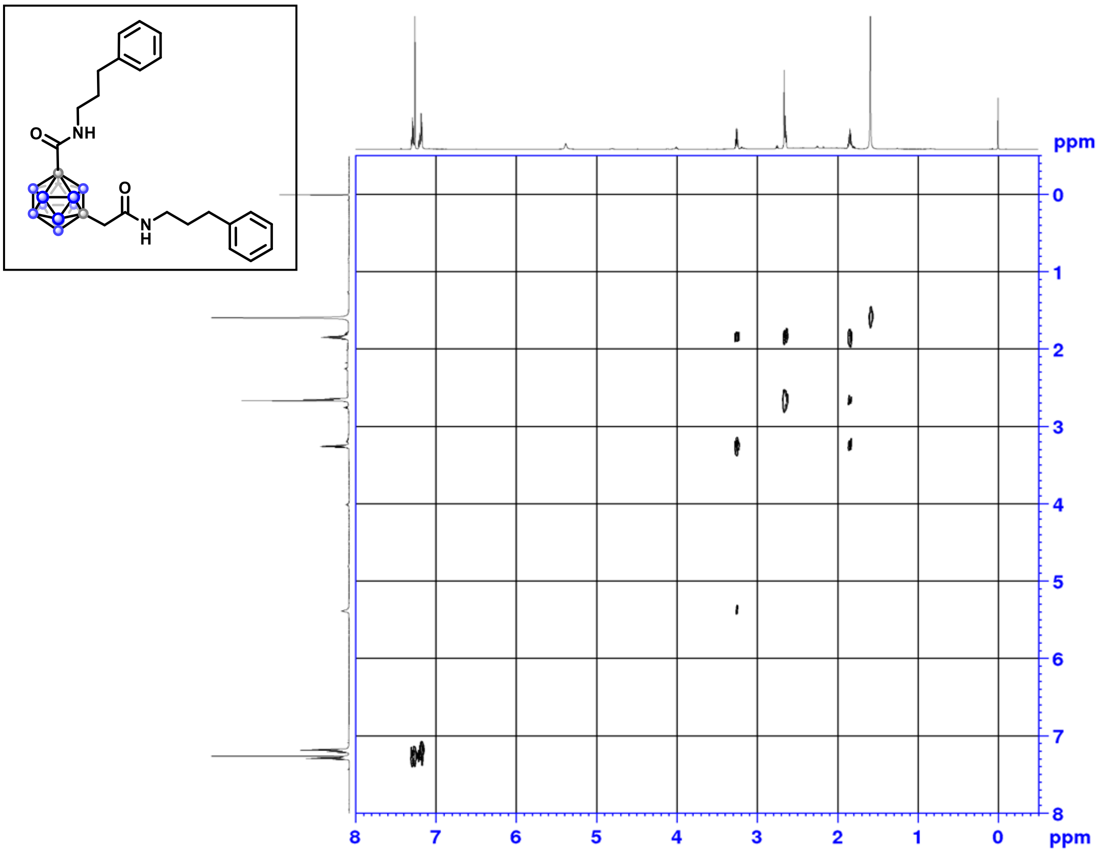

$^{1}H$-$^{13}C$ HSQC

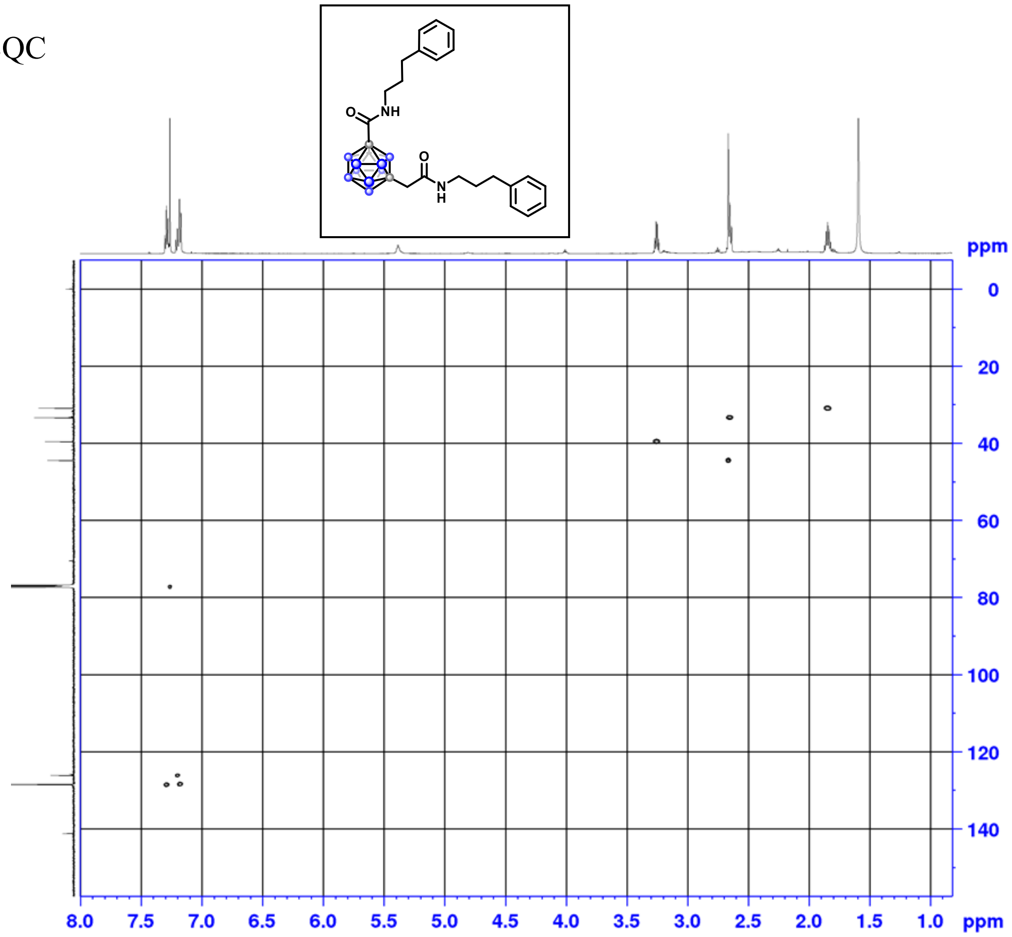

#### $^{1}H$-$^{13}C$ HMBC

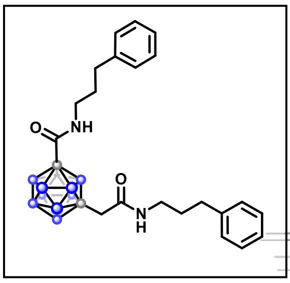

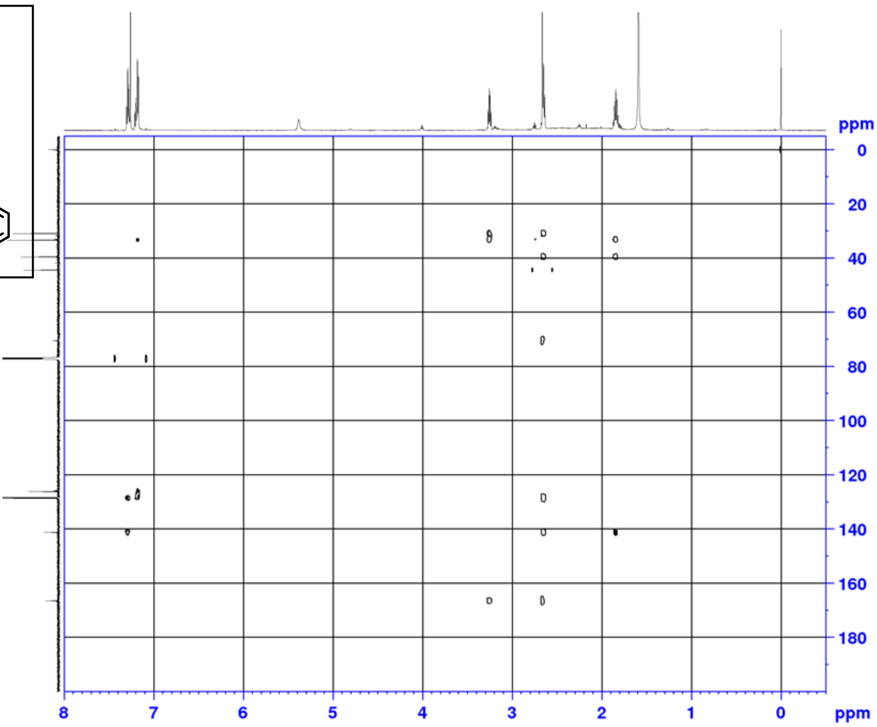

#### $^{1}H$-$^{15}N$ HMBC

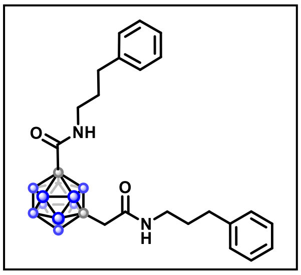

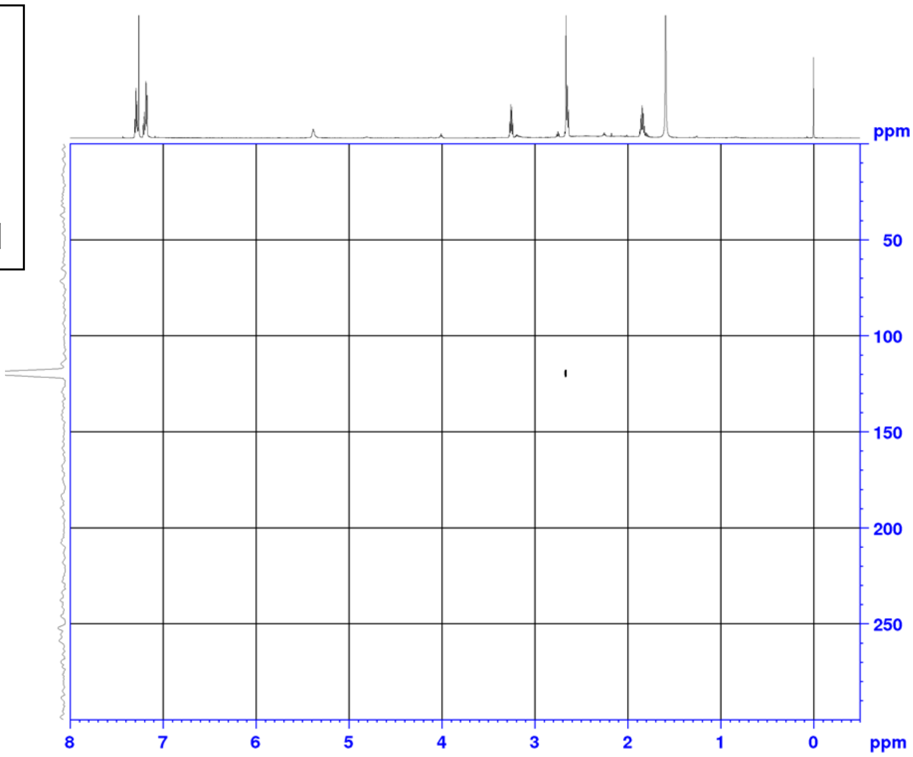

$^{1}H$ NMR (500 MHz, $CDCl_3$)

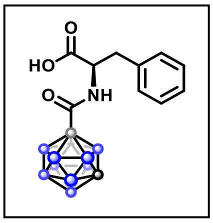

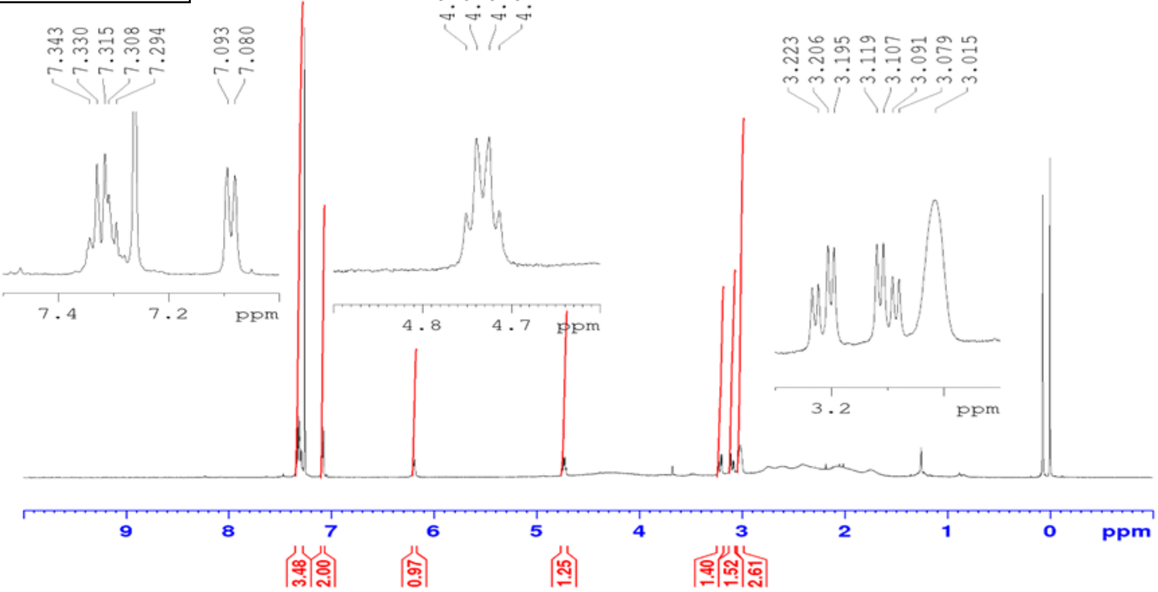

$^{13}C$ NMR (126 MHz, $CDCl_3$)

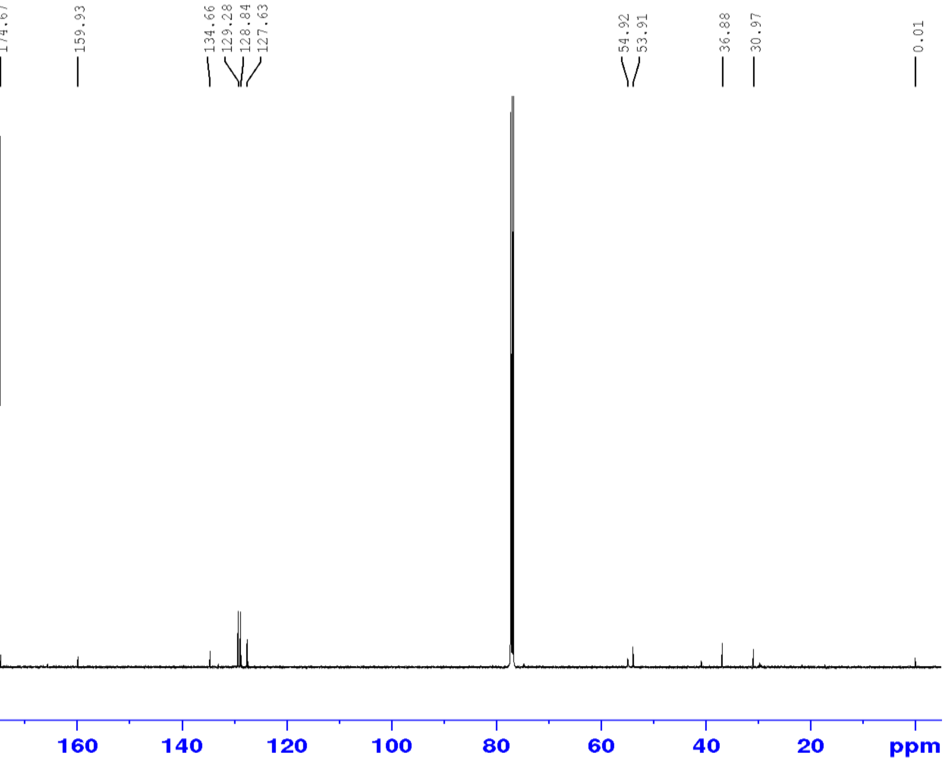

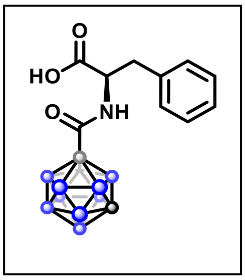

$^{11}$B NMR (160 MHz, $CDCl_3$)

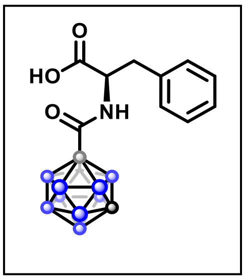

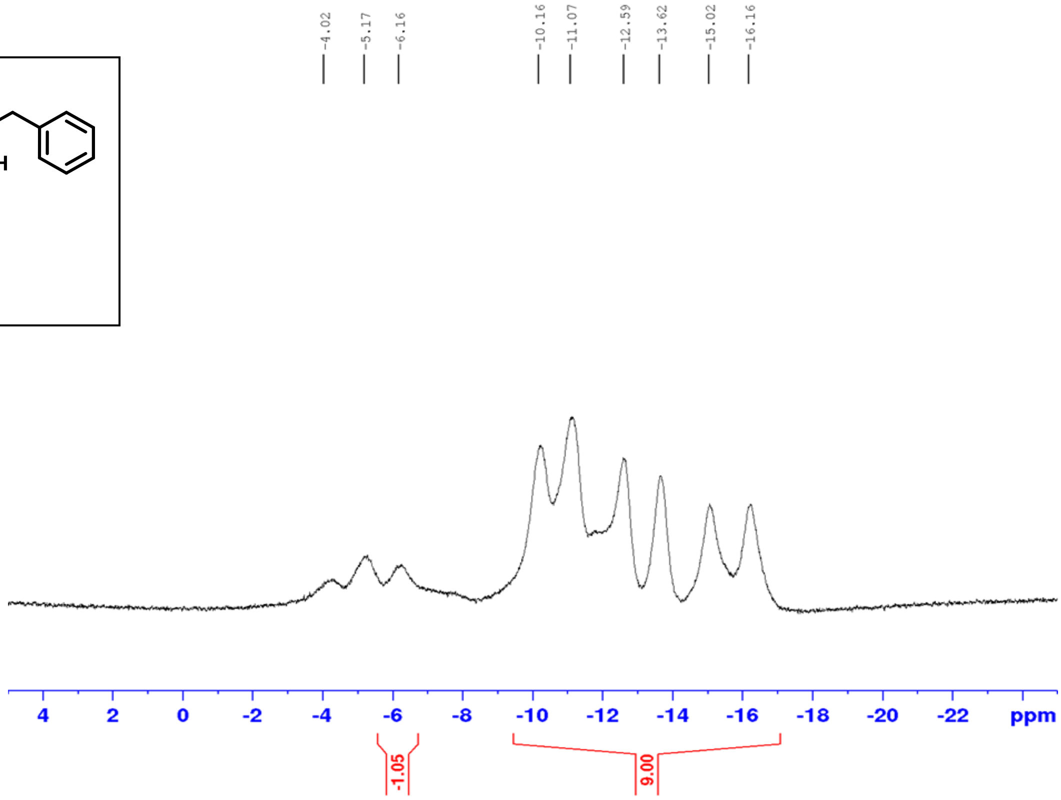

$^1$H-$^1$H COSY

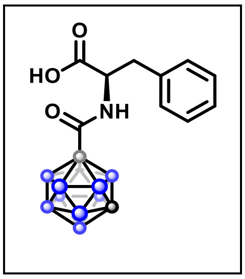

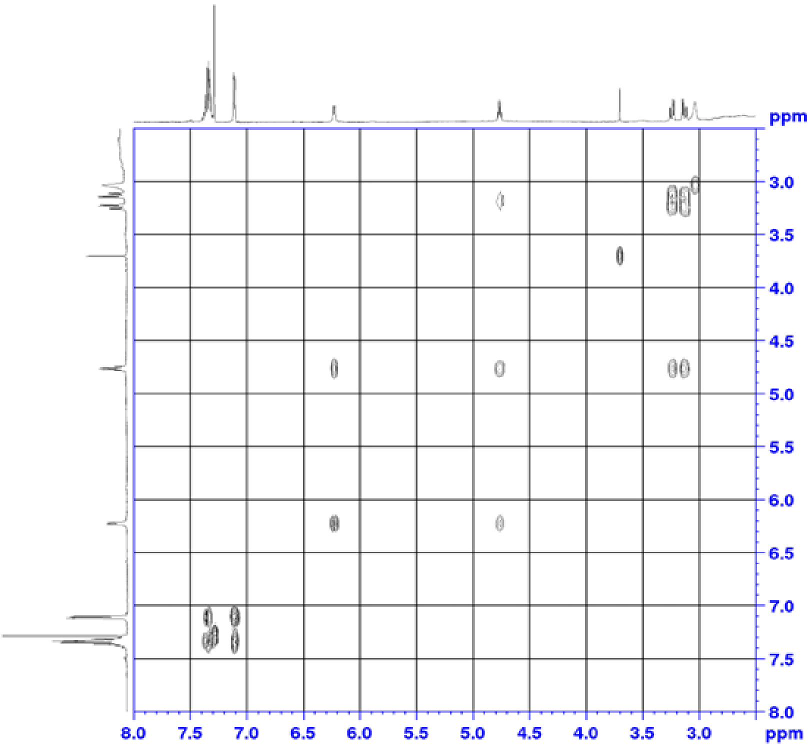

$^{1}H$-$^{13}C$ HSQC

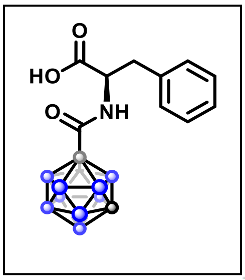

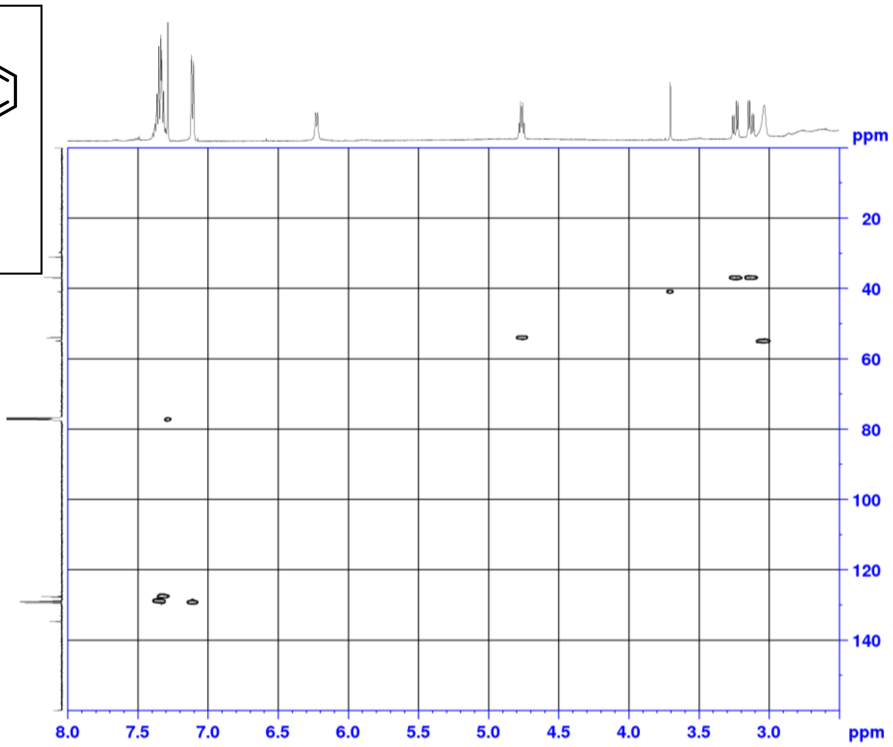

$^{1}H$-$^{13}C$ HMBC

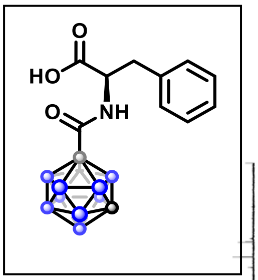

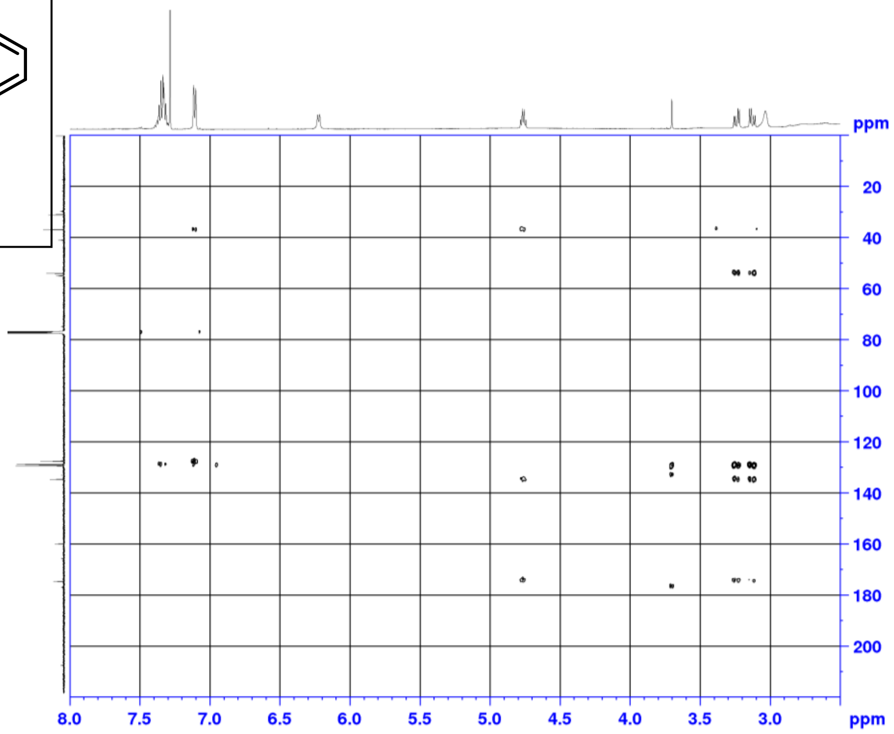

$^{1}H$ NMR (500 MHz, DMSO-$d_6$)

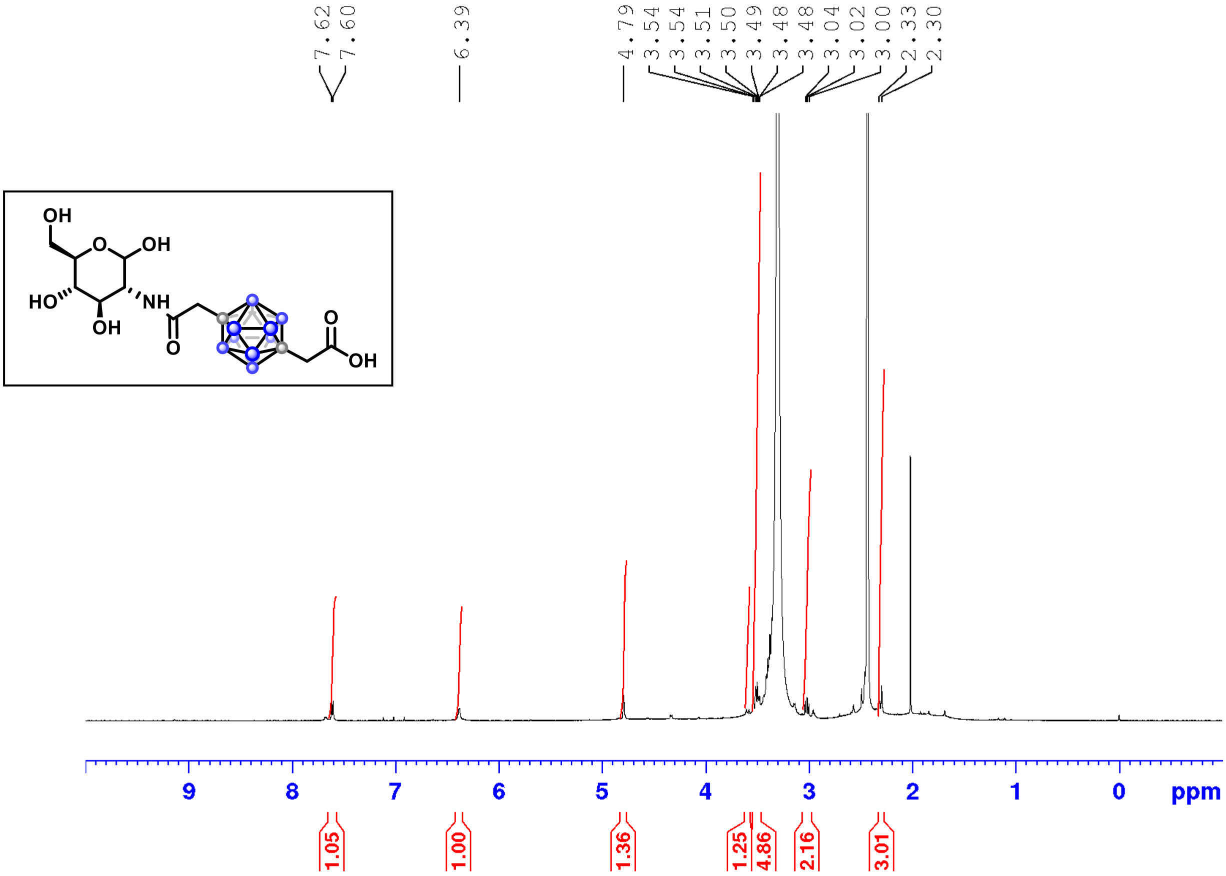

$^{11}B$ NMR (160 MHz, DMSO-$d_6$)

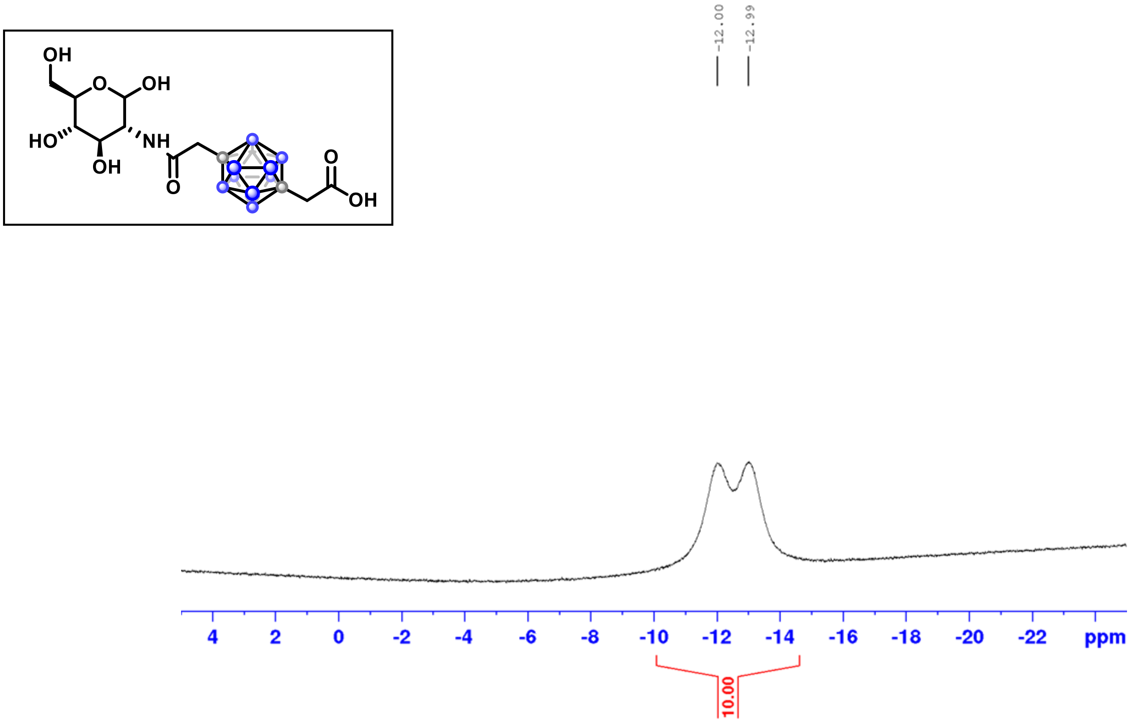

$^{13}C$ NMR (126 MHz, DMSO-$d_6$)

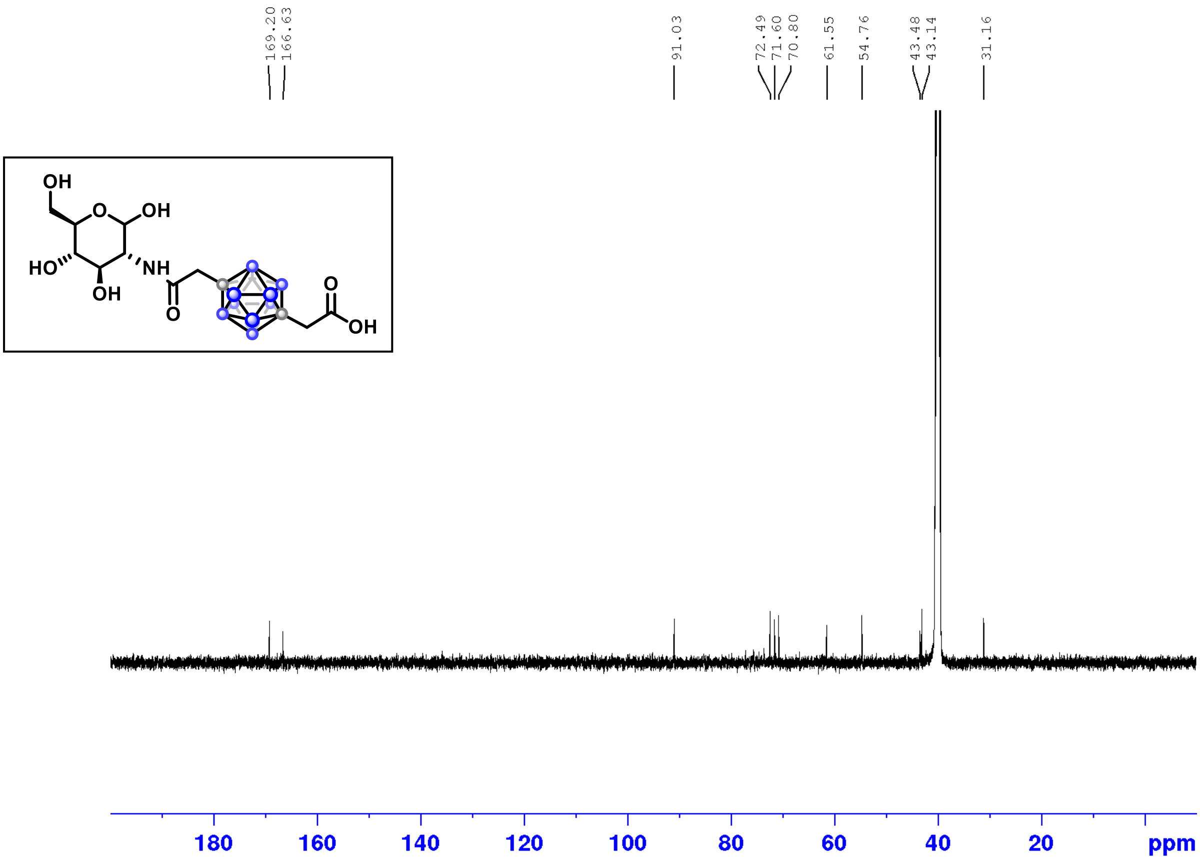

$^1H$-$^1H$ COSY

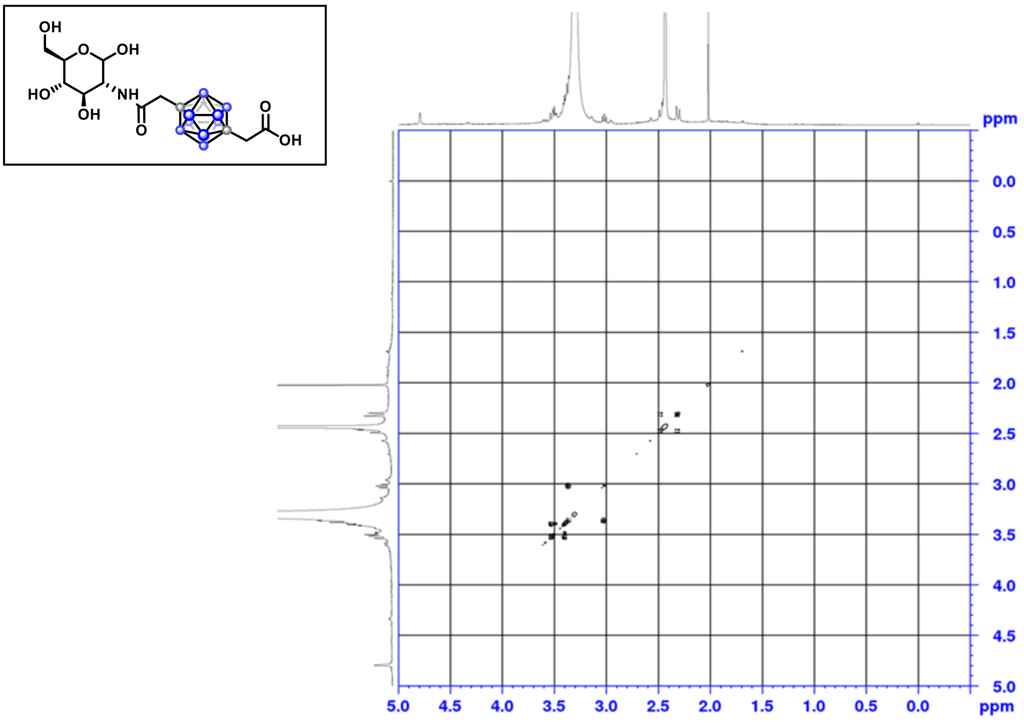

$^{1}H$-$^{13}C$ HSQC

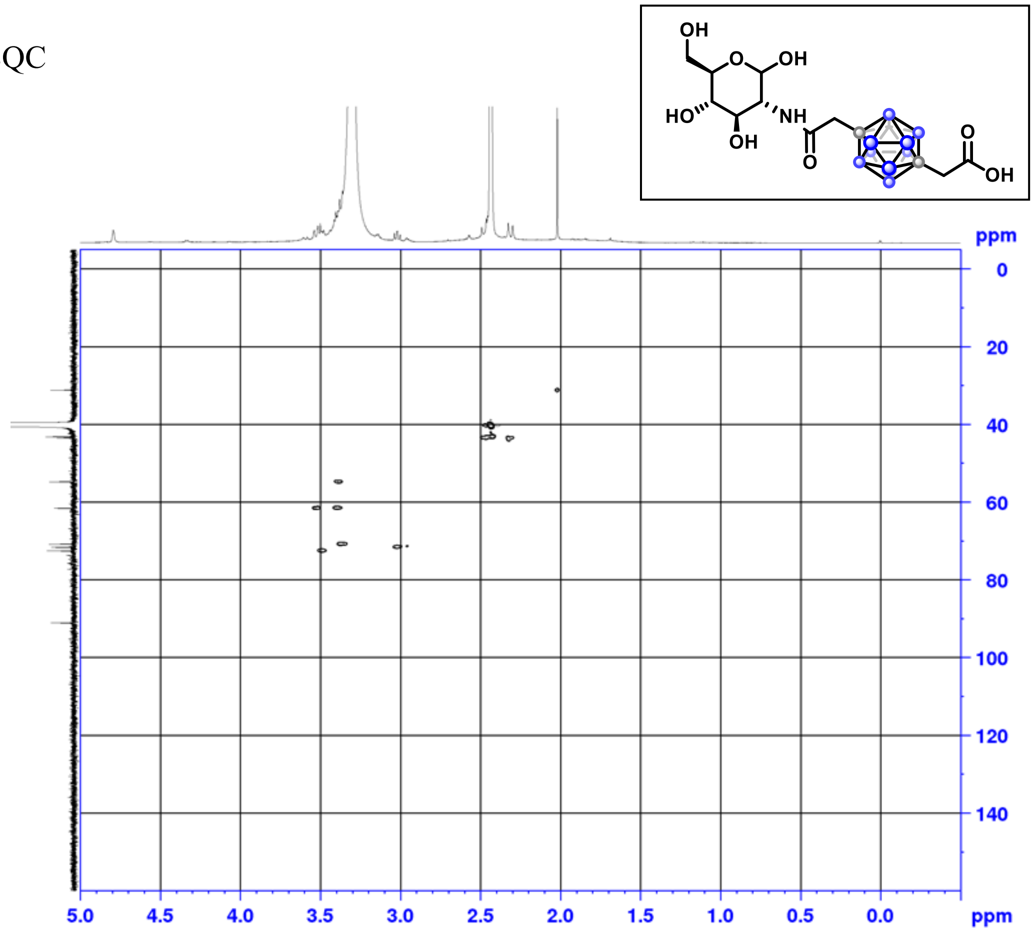

$^{1}H$-$^{13}C$ HMBC

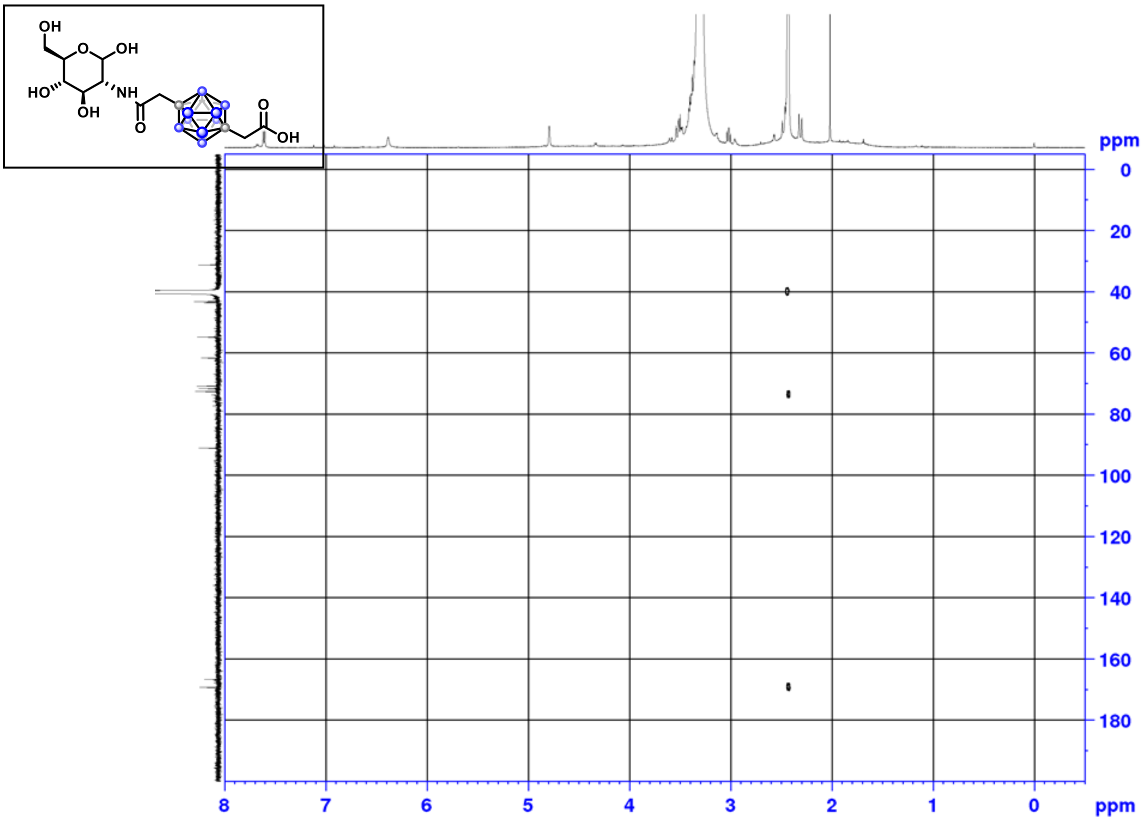

$^{1}H$ NMR (500 MHz, DMSO-$d_6$)

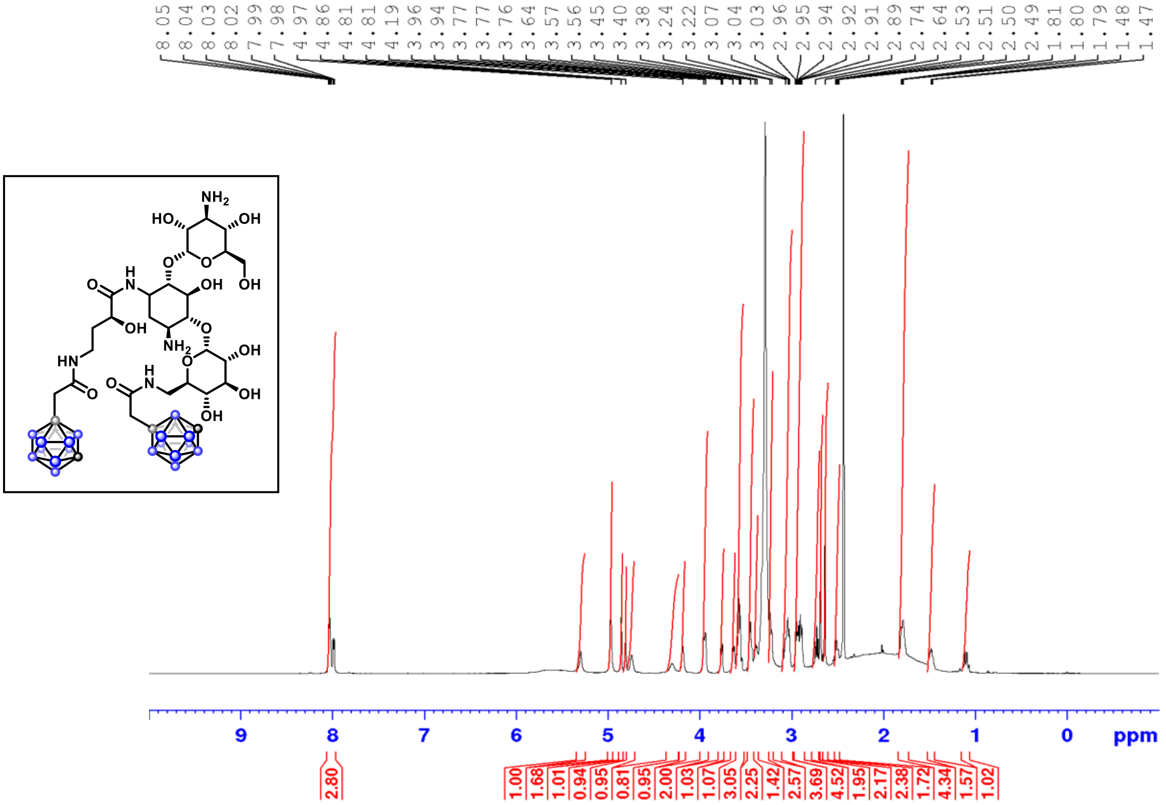

$^{13}C$ NMR (126 MHz, DMSO-$d_6$)

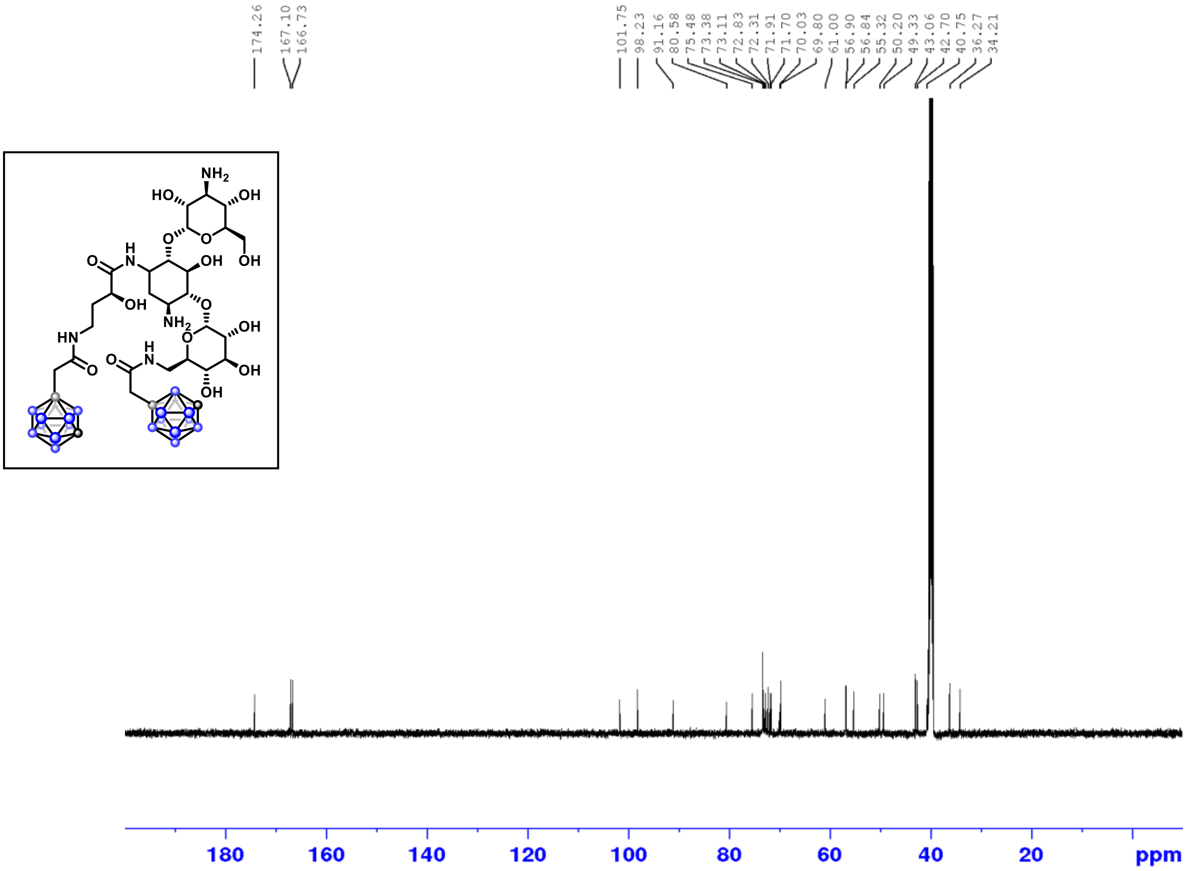

[1]H NMR (500 MHz, DMSO-$d_6$)

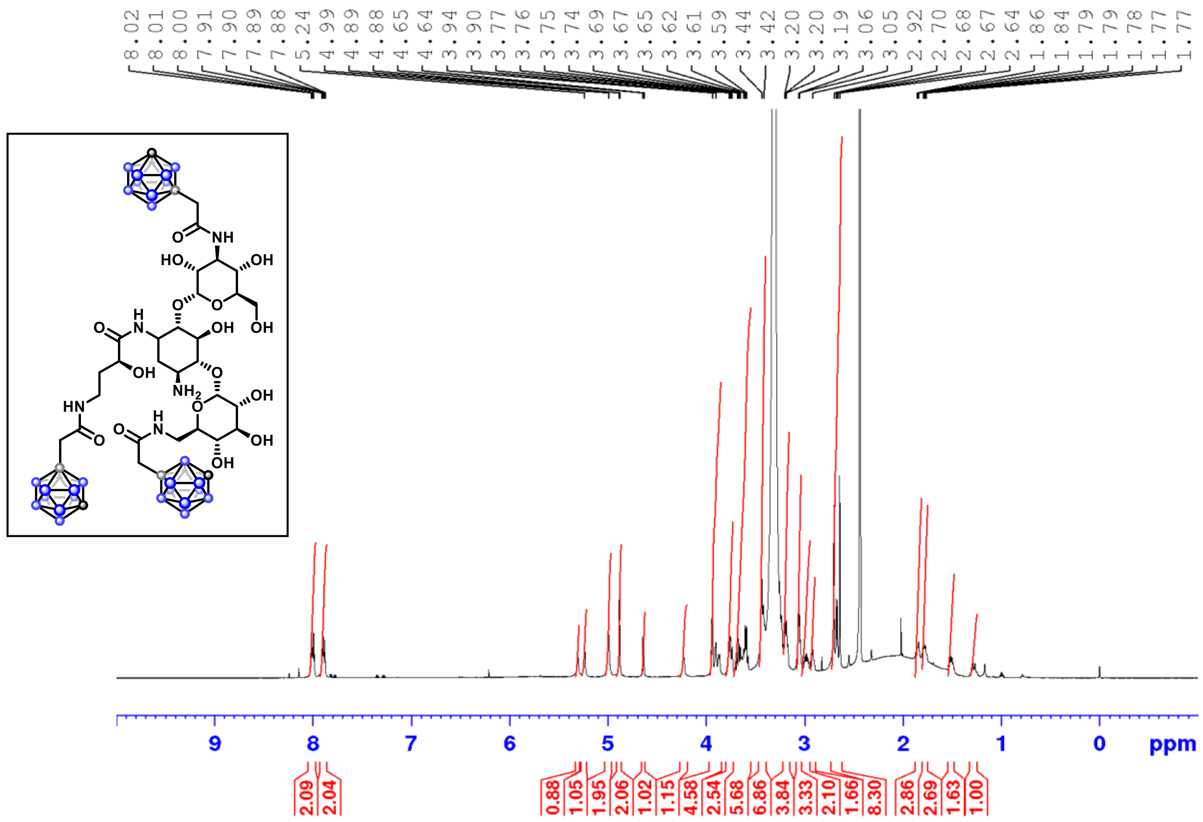

[13]C NMR (126 MHz, DMSO-$d_6$)

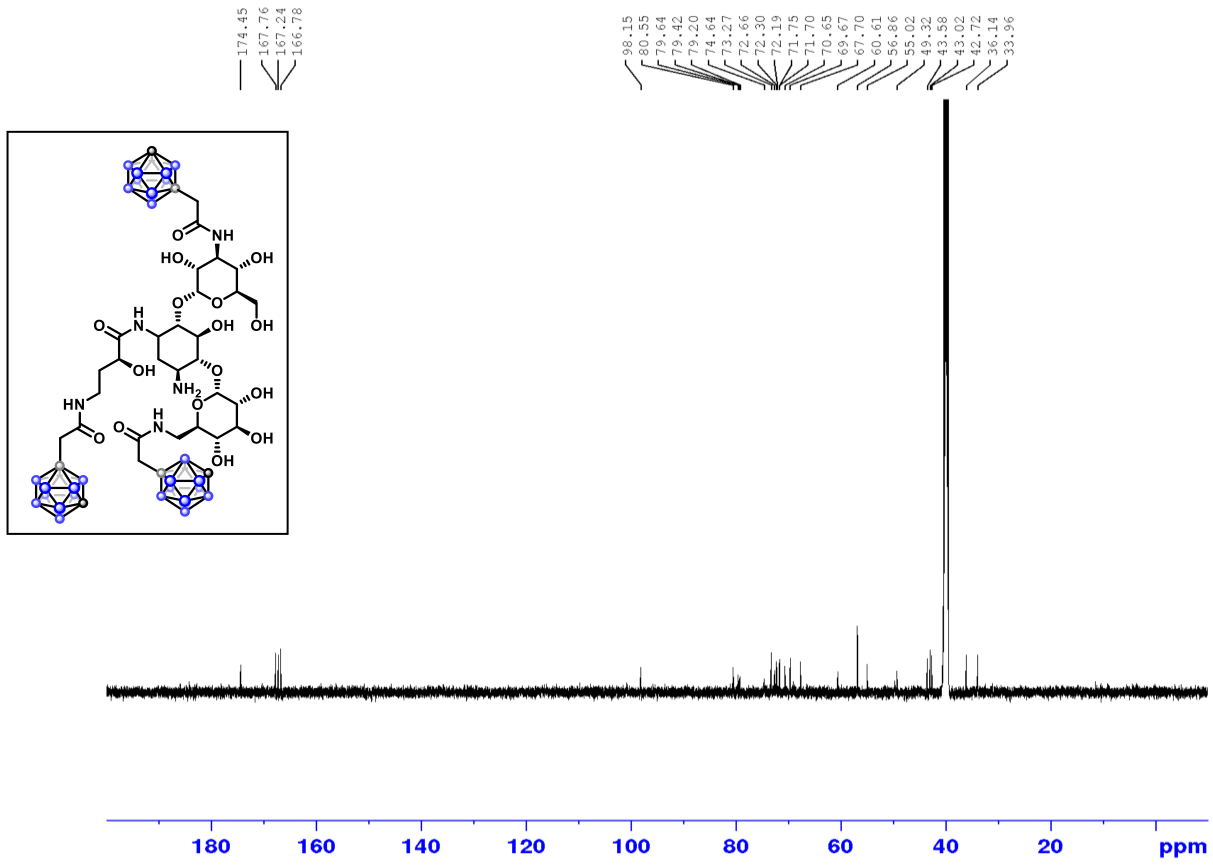

$^{1}H$ NMR (500 MHz, DMSO-$d_6$)

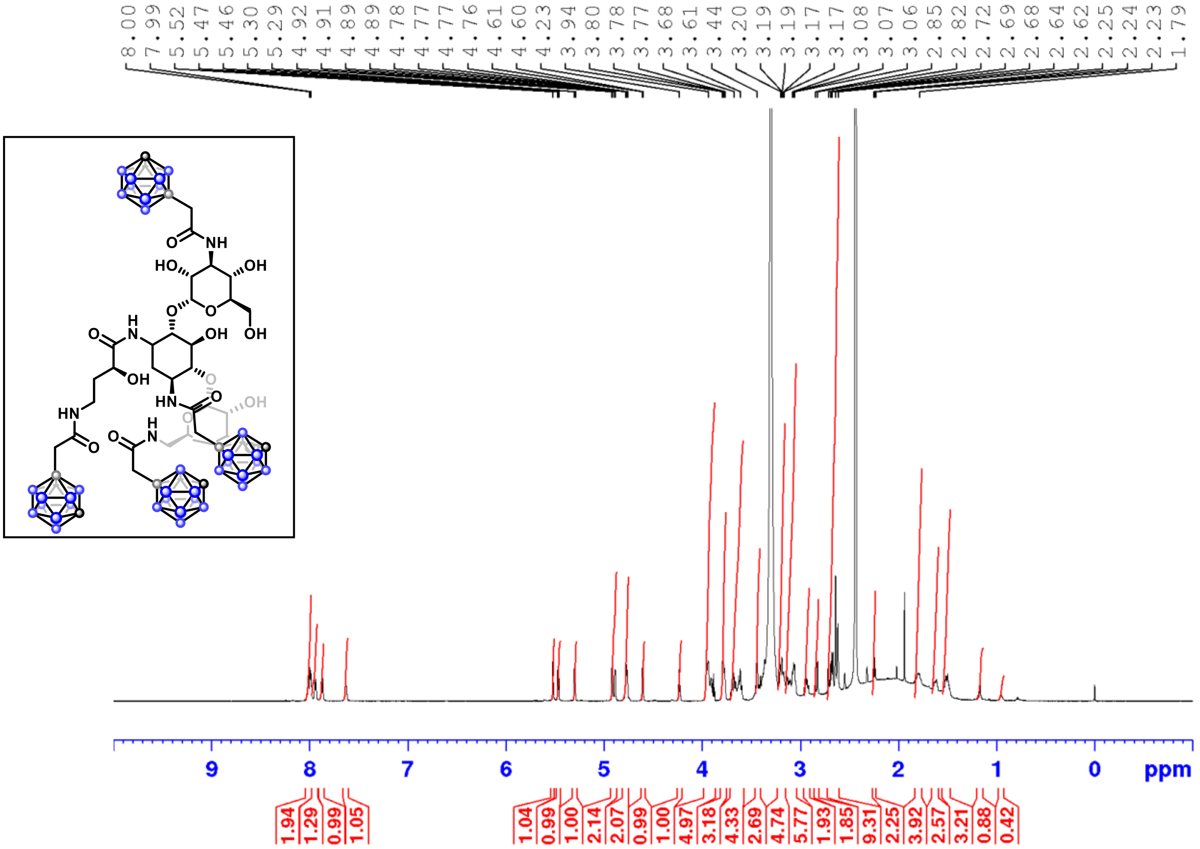

Stack of $^{1}H$ NMR spectra of amikacin, **18**, **19** and **20**. The conjugation of carborane fragments corresponds to an increase in spectral complexity, along with enhanced signal intensity (integrals) in the δ 3.5–1.8 ppm region, where the B-H proton resonances of the carborane units are observed.

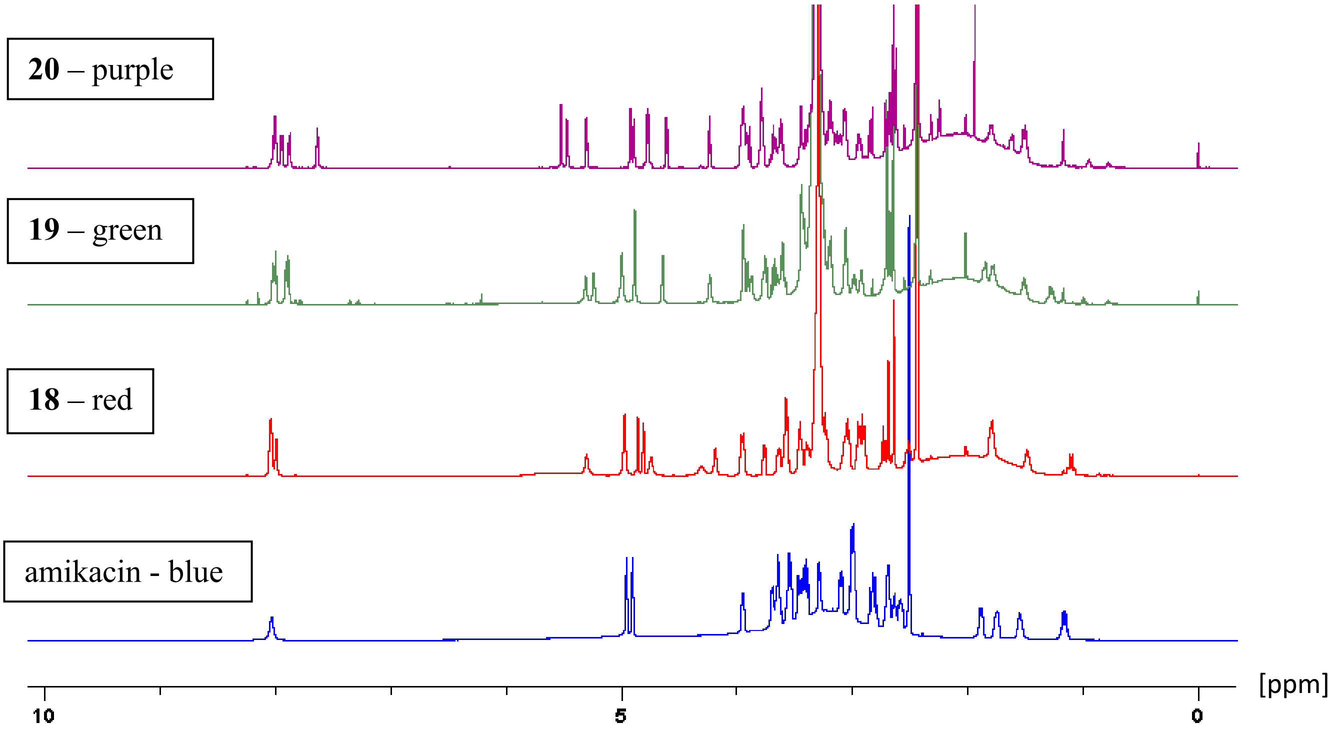

$^{1}H$ NMR (500 MHz, DMSO-$d_6$)

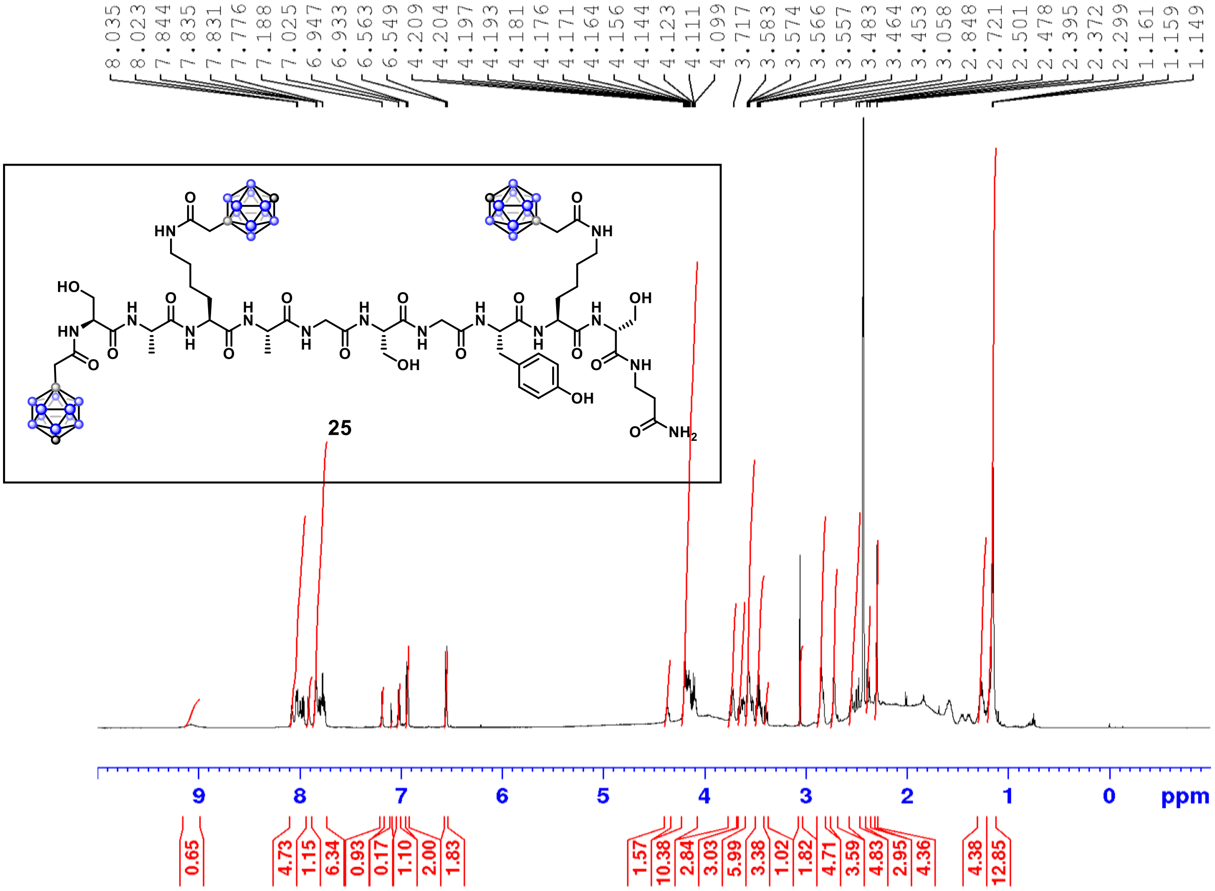

Stack of $^{1}H$ NMR spectra of Peptide E (red) and **25** (blue). Compared to the starting Peptide E, the carborane-conjugated peptide **25** exhibits increased spectral complexity, along with the clear appearance of B–H resonances corresponding to the carborane cages in the blue spectrum.

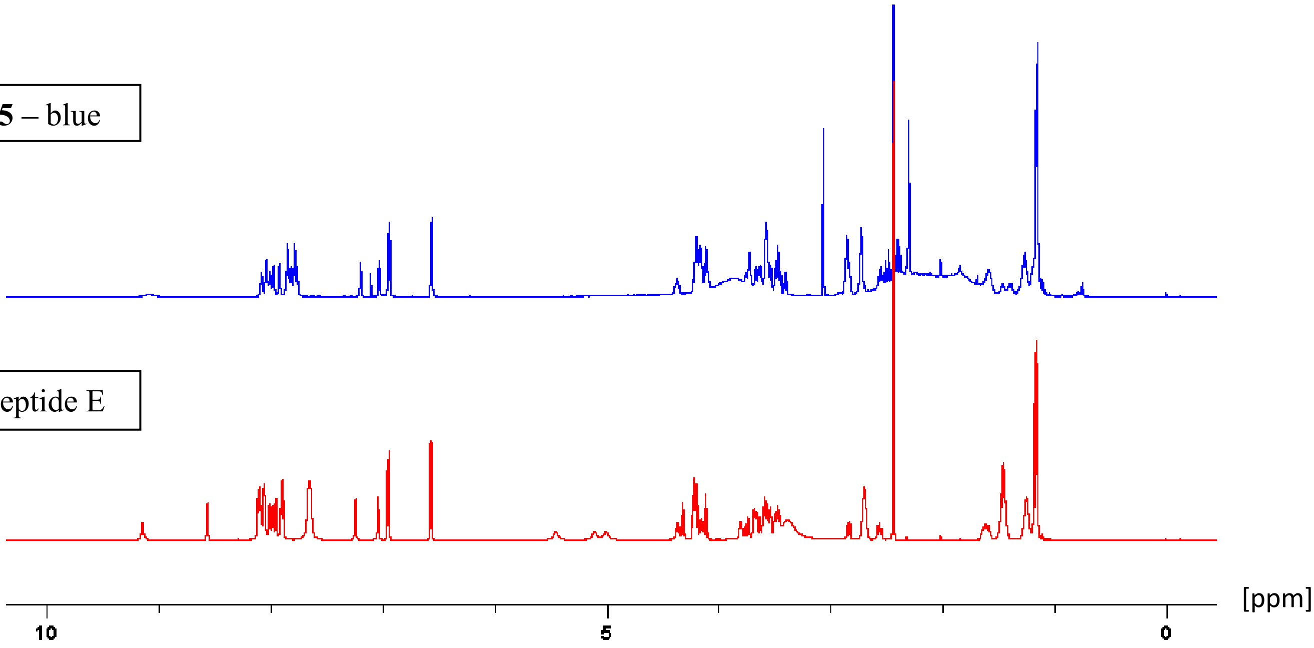

$^{13}C$ NMR (126 MHz, DMSO-$d_6$)

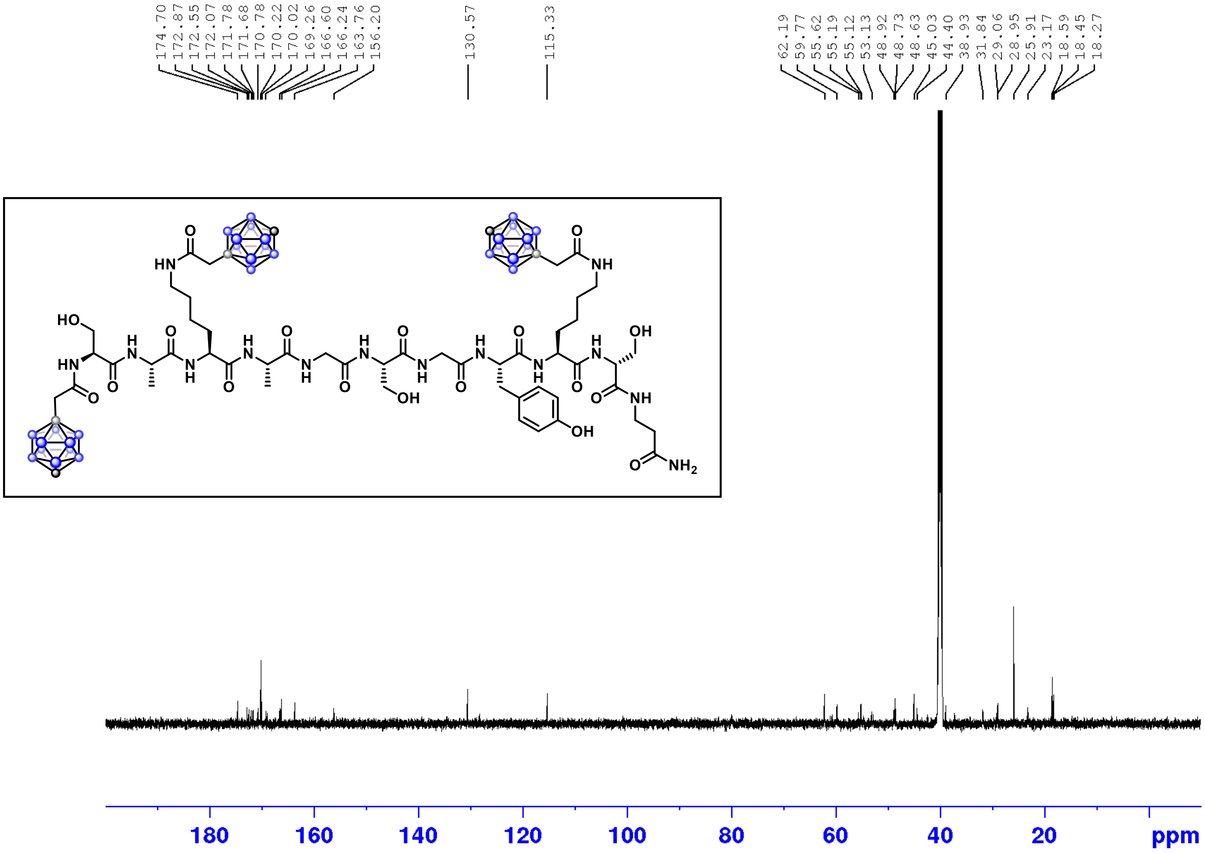

$^{11}B$ NMR (160 MHz, $CDCl_3$)

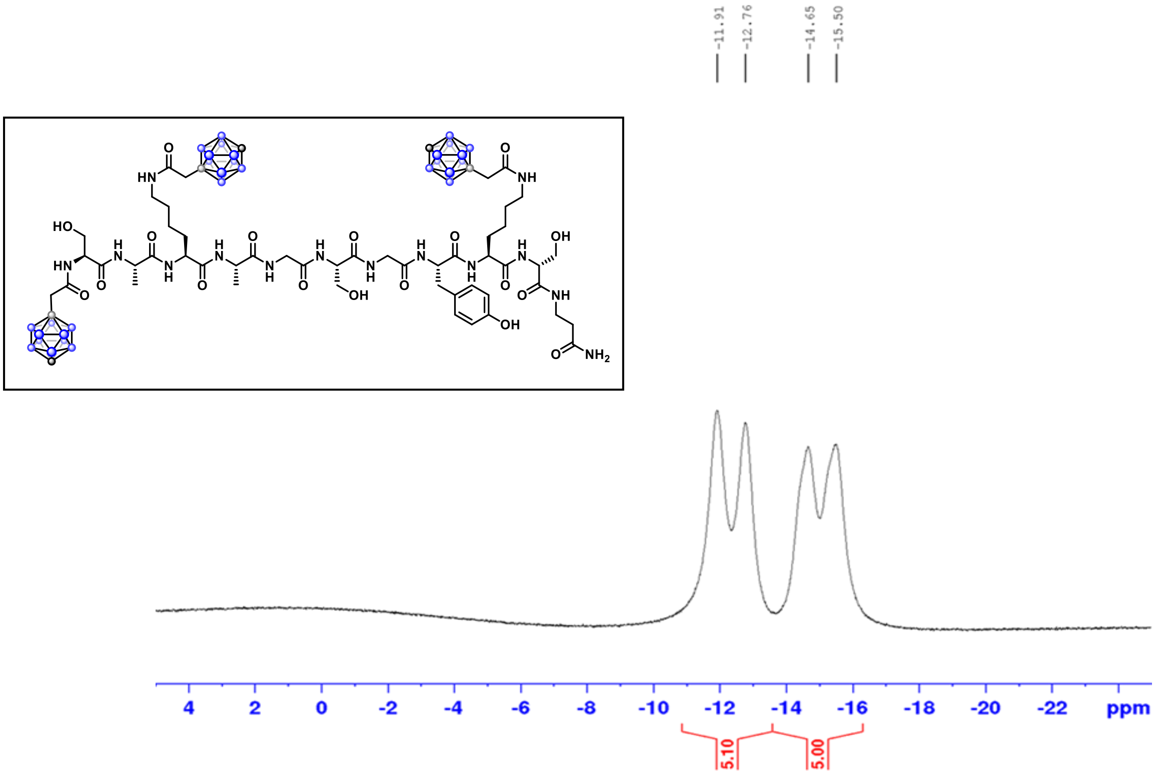

Stack of ¹H NMR spectra of Peptide B (red) and **26** (blue).

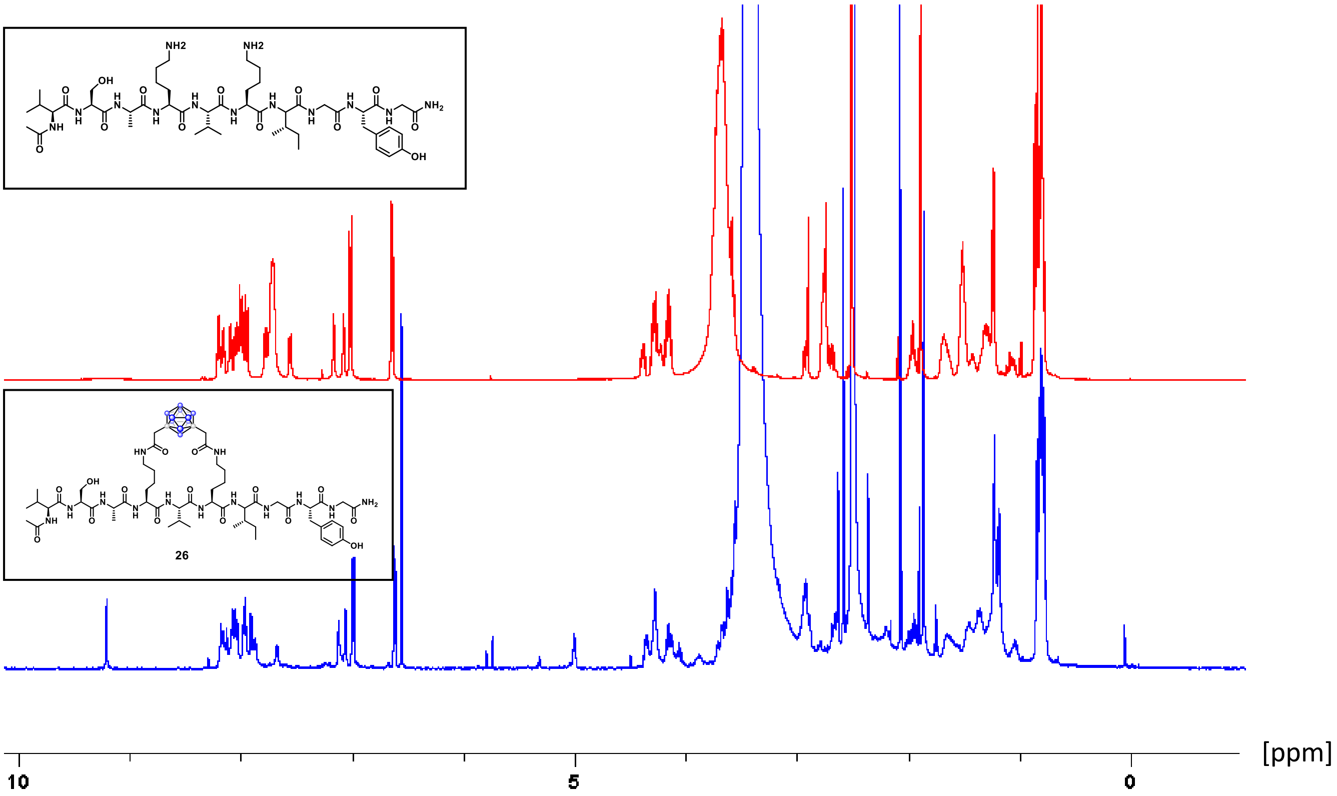

Stack of ¹H NMR spectra of Peptide B (red) and **27** (blue).

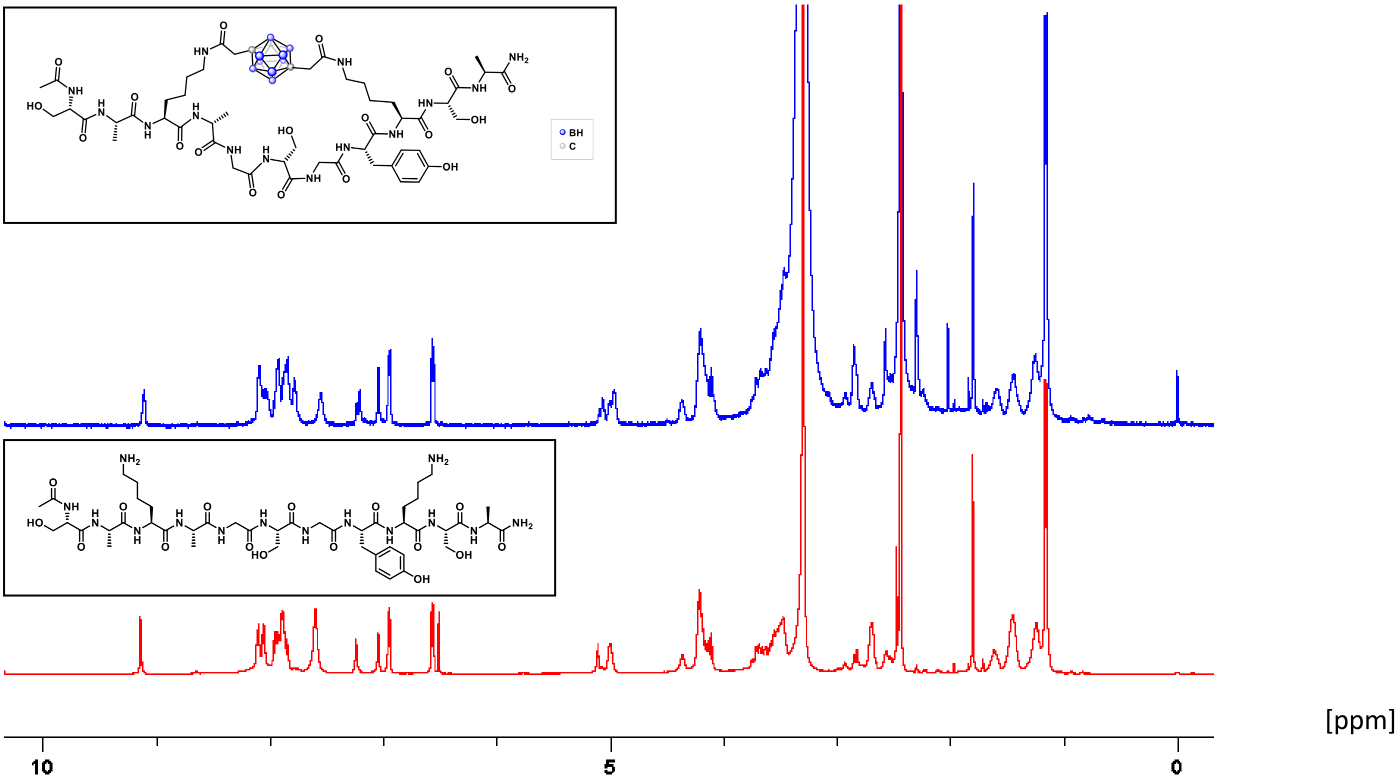

[1]H NMR (500 MHz, DMSO-$d_6$)

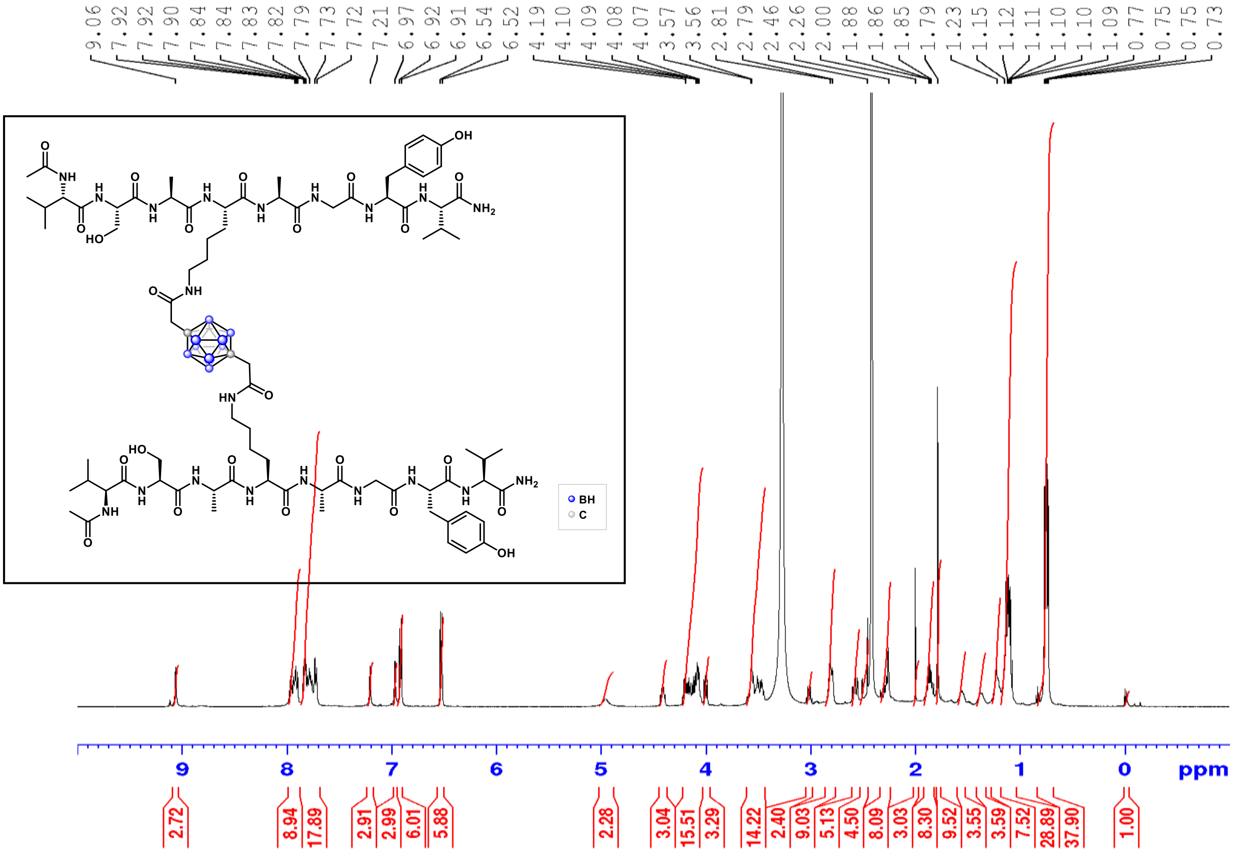

[13]C NMR (126 MHz, DMSO-$d_6$)

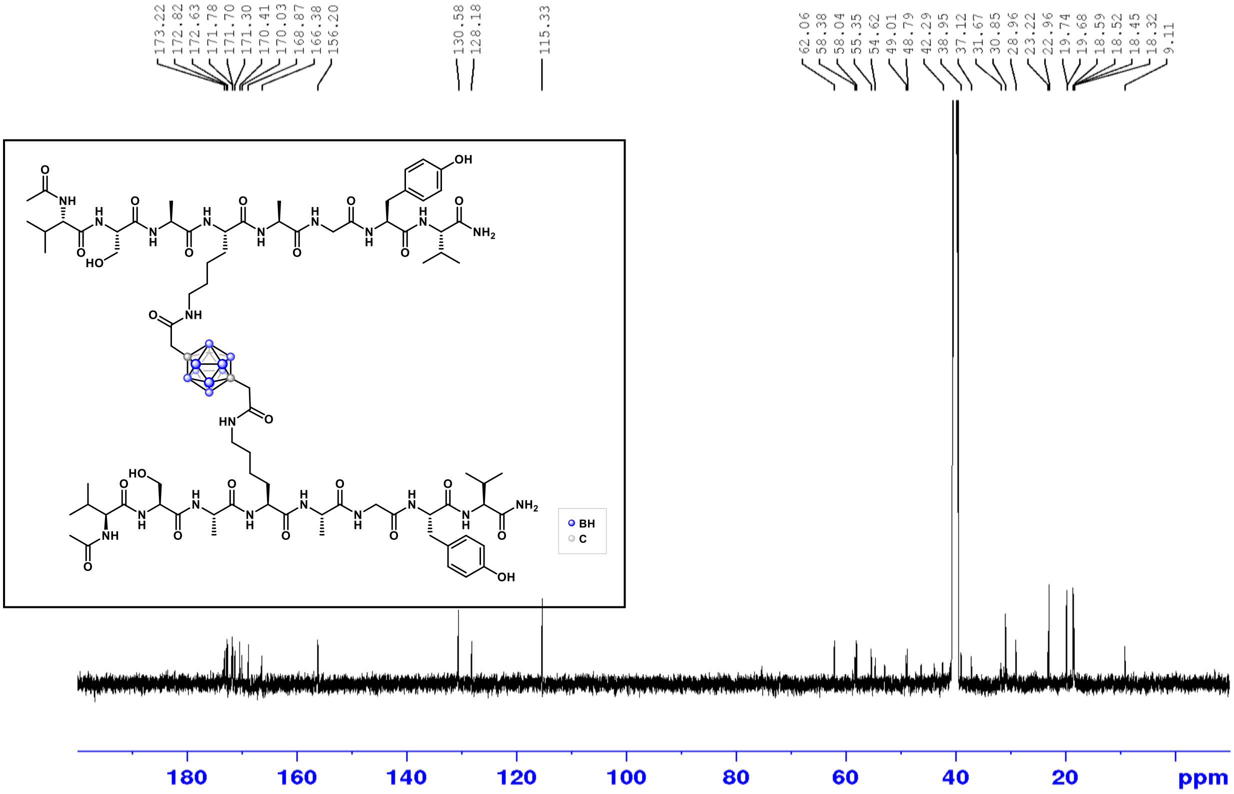

$^{11}B$ NMR (160 MHz, $CDCl_3$)

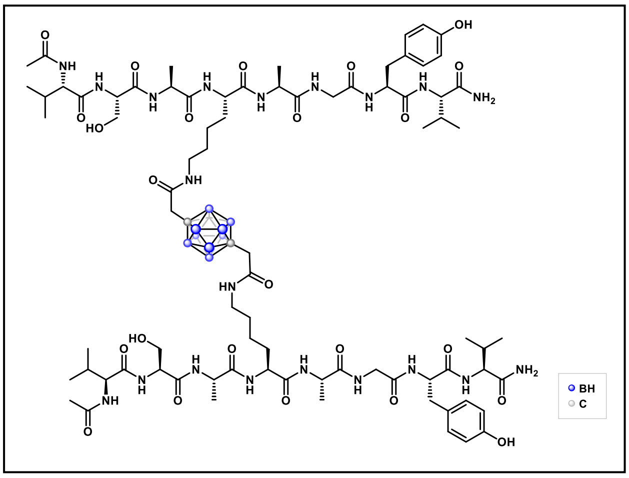

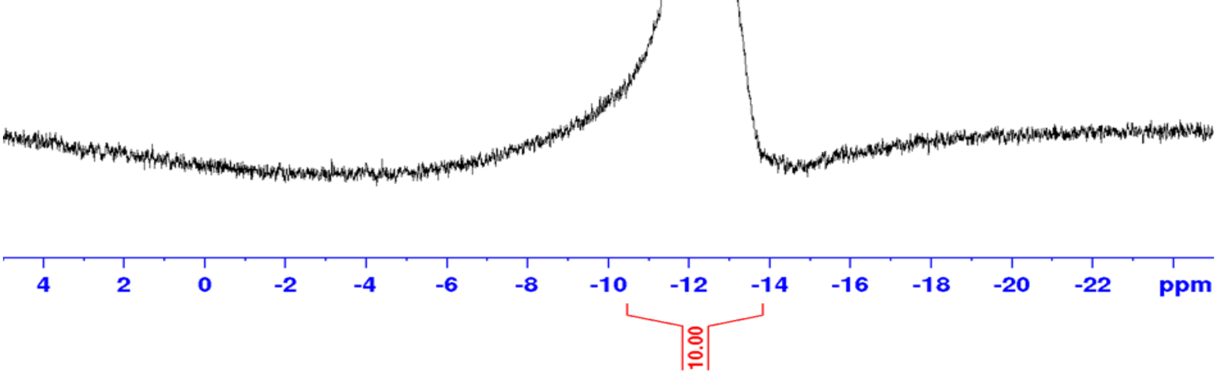

### Copies of HRMS spectra

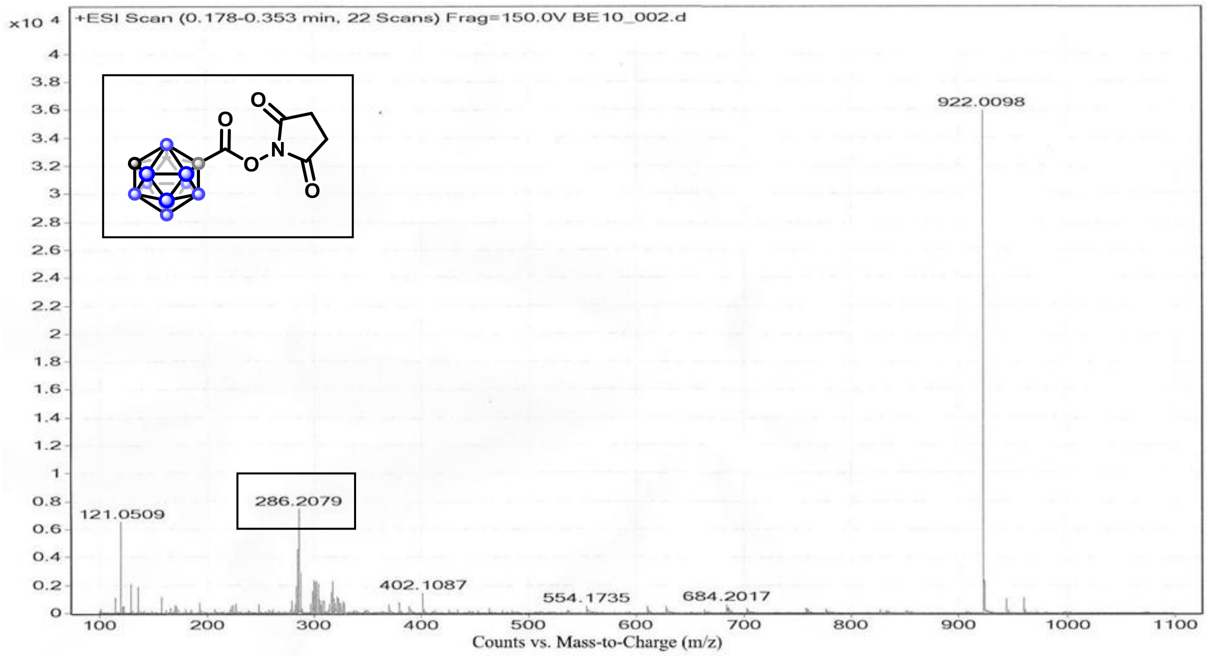

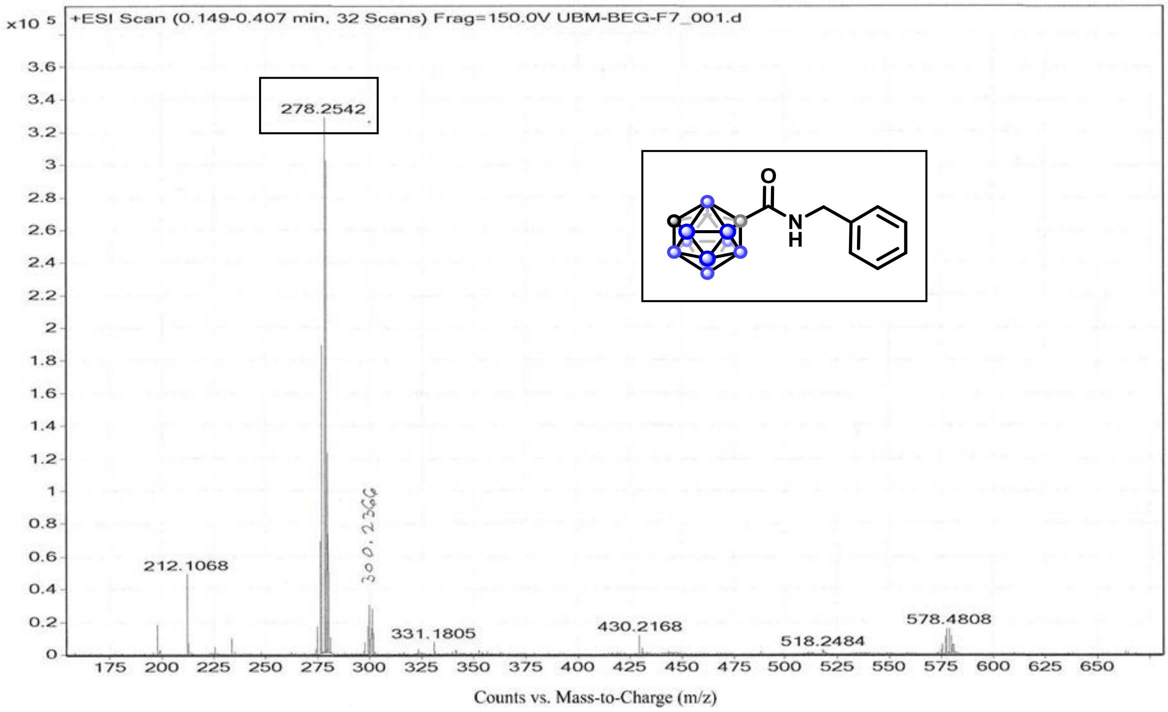

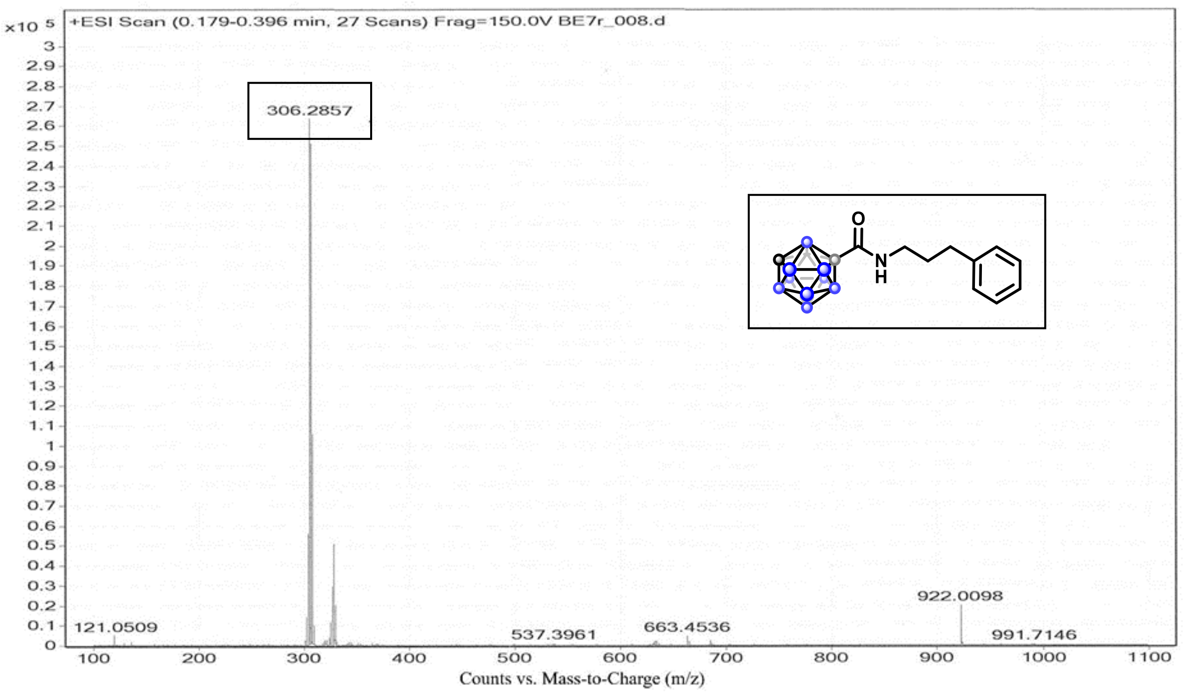
x10 5
+ESI Scan (0.179-0.396 min, 27 Scans) Frag=150.0V BE7r_008.d
306.2857
922.0098
121.0509
537.3961
663.4536
991.7146
Counts vs. Mass-to-Charge (m/z)

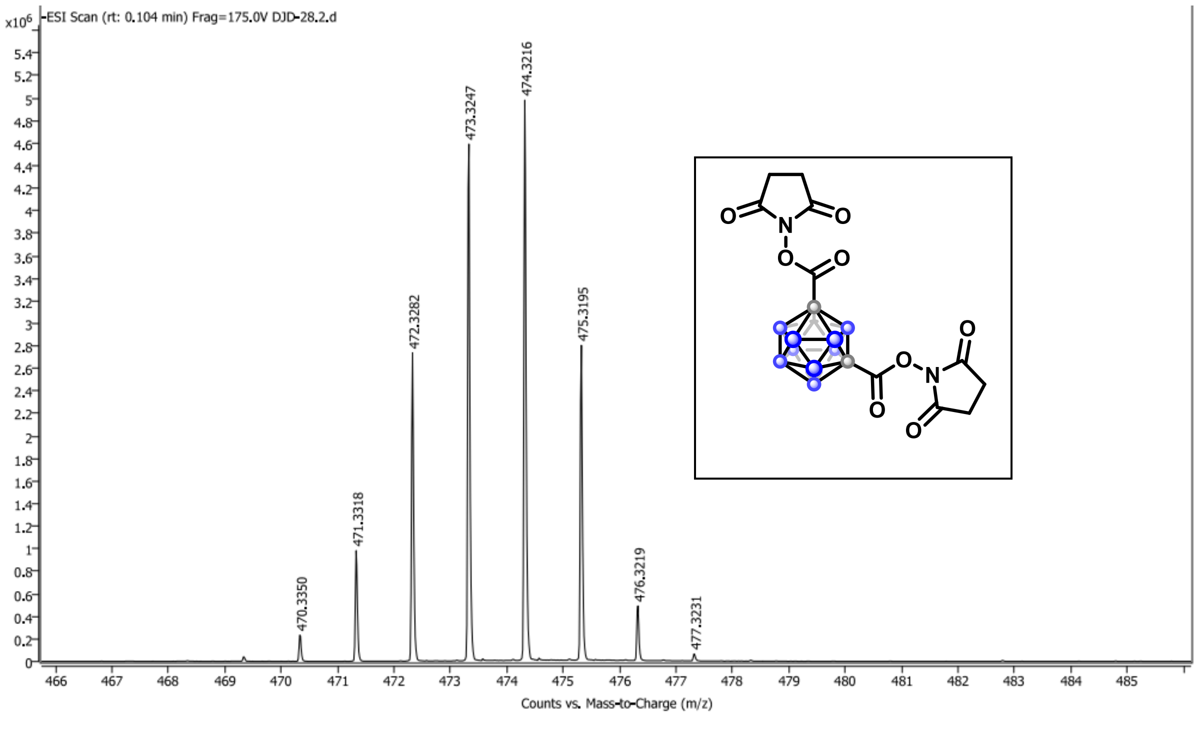
x10^6
-ESI Scan (rt: 0.104 min) Frag=175.0V DJD-28.2.d
474.3216
473.3247
472.3282
475.3195
471.3318
470.3350
476.3219
477.3231
Counts vs. Mass-to-Charge (m/z)

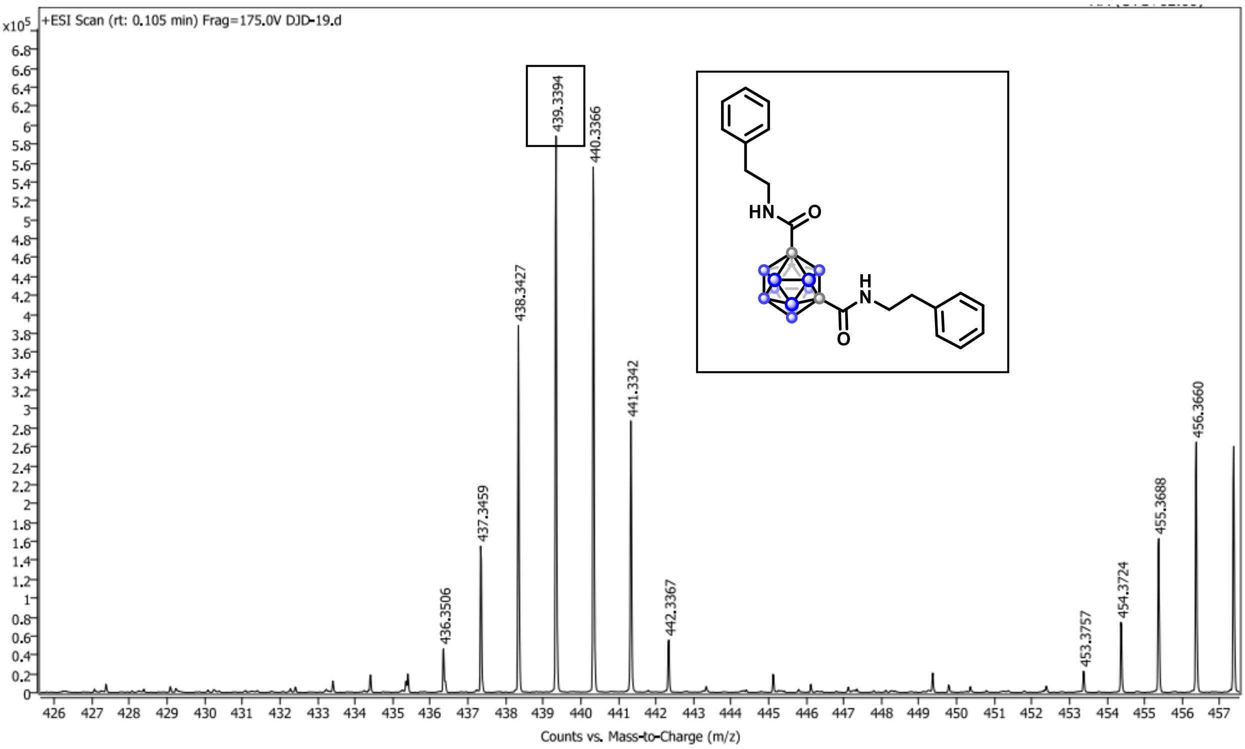
+ESI Scan (rt: 0.105 min) Frag=175.0V DJD-19.d
x10^5
439.3394
440.3366
438.3427
441.3342
437.3459
436.3506
442.3367
453.3757
454.3724
455.3688
456.3660
HN
O
H
N
O
Counts vs. Mass-to-Charge (m/z)

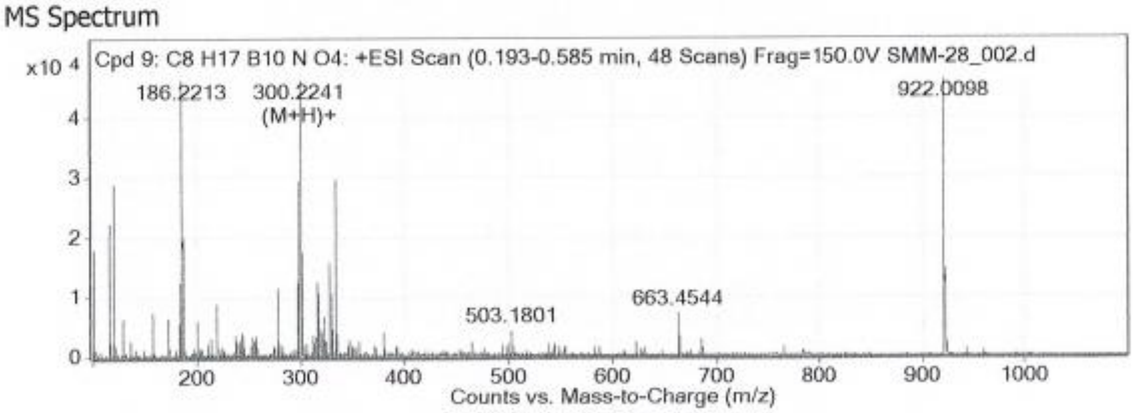
MS Spectrum
Cpd 9: C8 H17 B10 N O4: +ESI Scan (0.193-0.585 min, 48 Scans) Frag=150.0V SMM-28_002.d
x10 4
186.2213
300.2241
(M+H)+
922.0098
503.1801
663.4544
Counts vs. Mass-to-Charge (m/z)

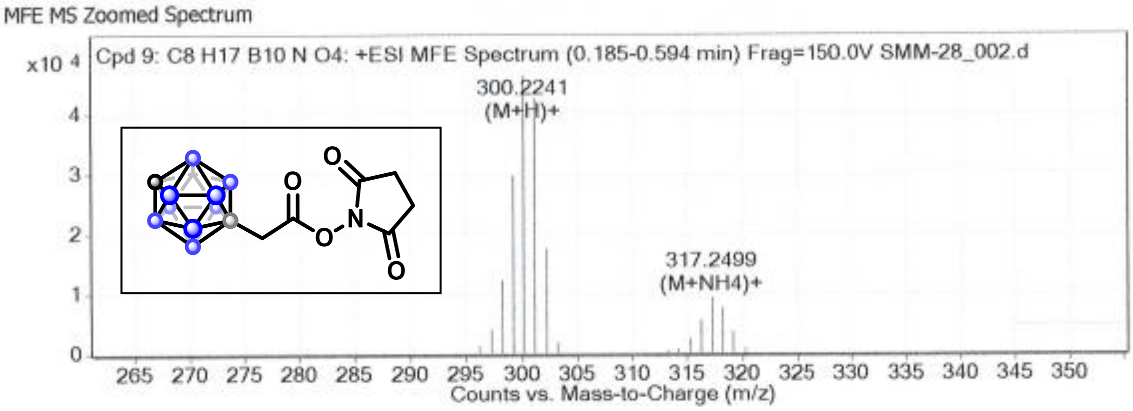
MFE MS Zoomed Spectrum
Cpd 9: C8 H17 B10 N O4: +ESI MFE Spectrum (0.185-0.594 min) Frag=150.0V SMM-28_002.d
x10 4
300.2241
(M+H)+
317.2499
(M+NH4)+
O
O
N
O
O
Counts vs. Mass-to-Charge (m/z)

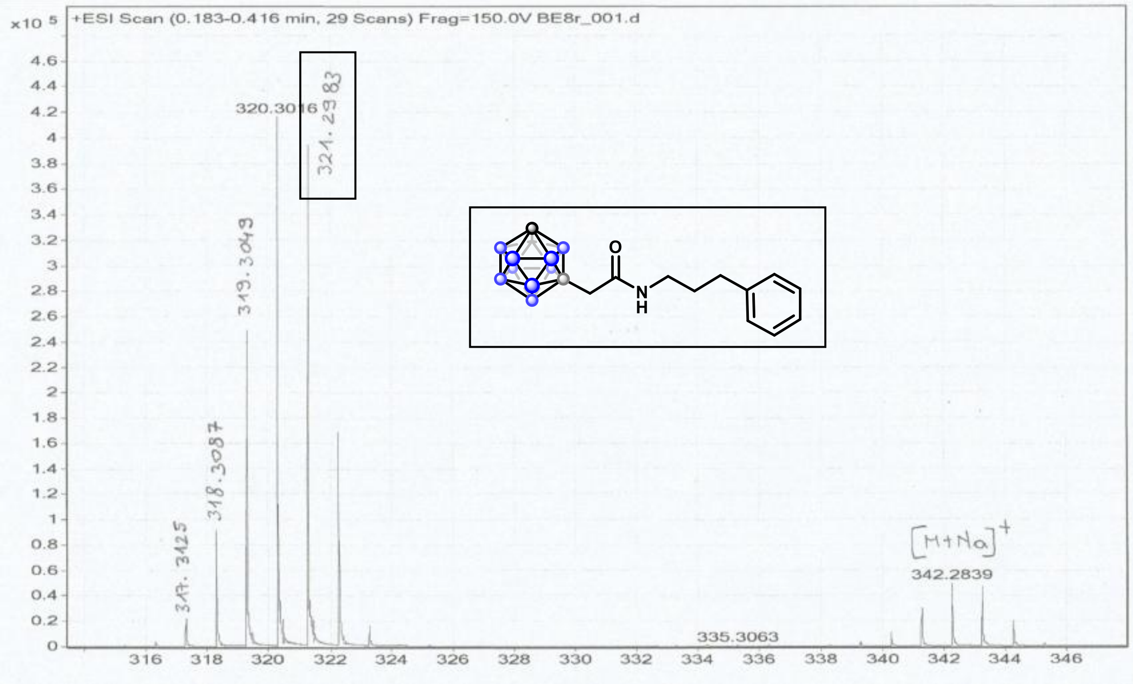

MS Spectrum

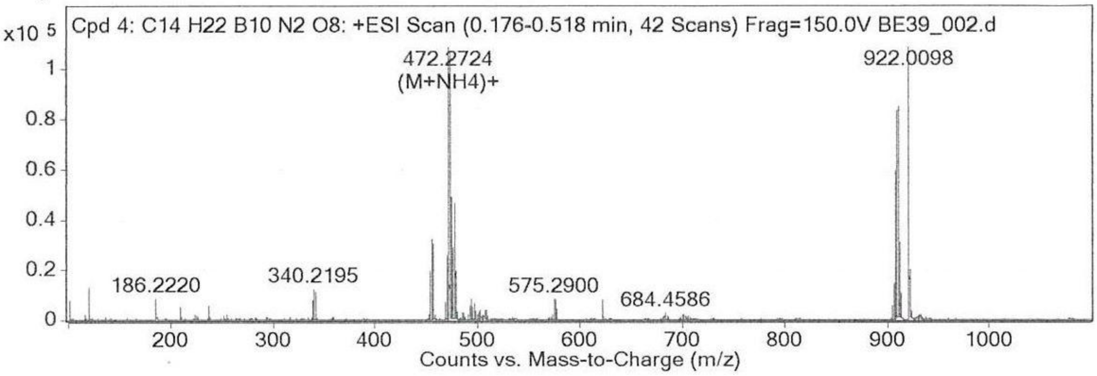

MFE MS Zoomed Spectrum

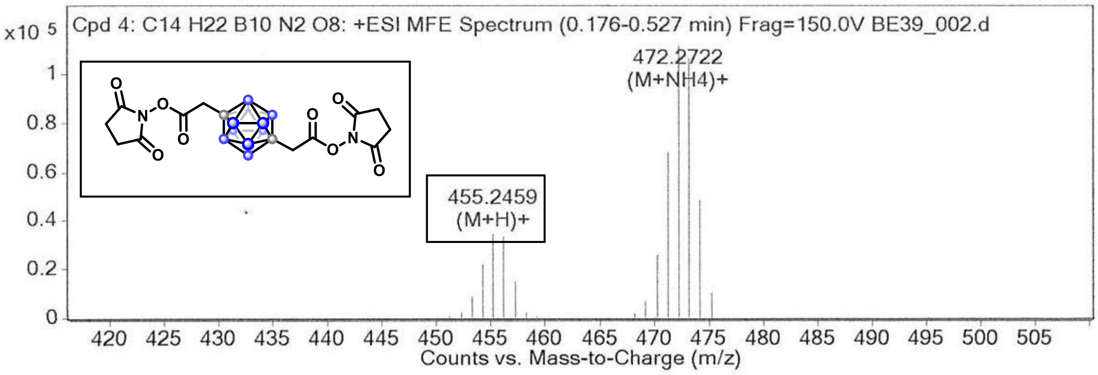

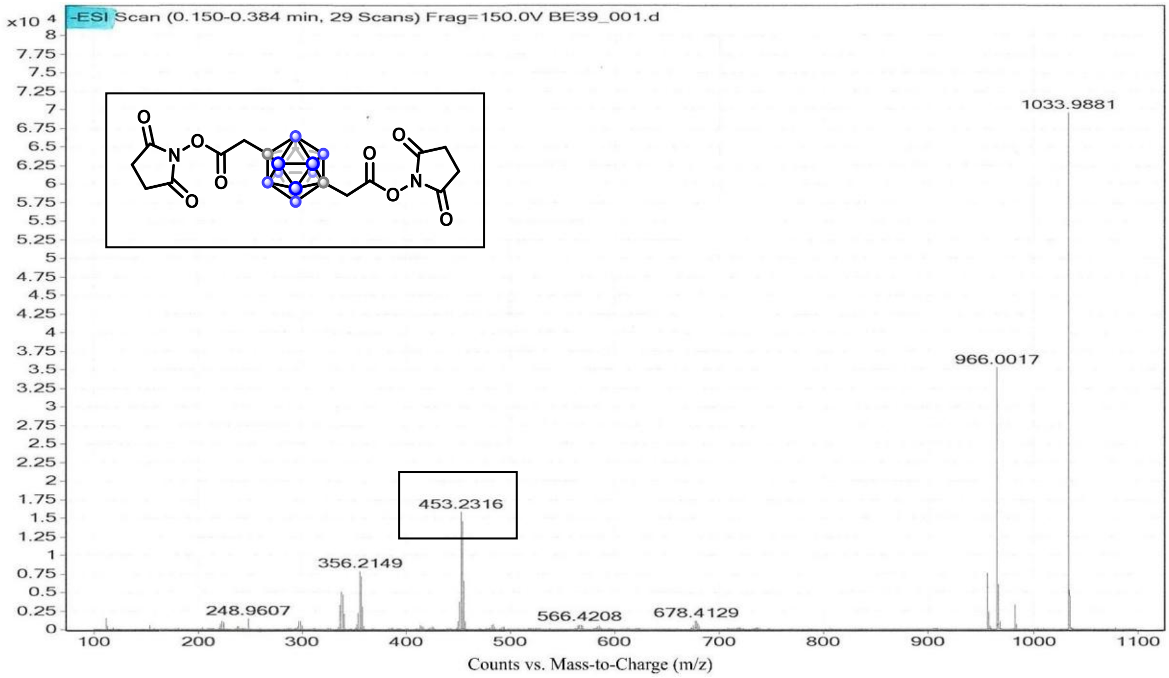
x10 4
-ESI Scan (0.150-0.384 min, 29 Scans) Frag=150.0V BE39_001.d
1033.9881
966.0017
453.2316
356.2149
248.9607
566.4208
678.4129
Counts vs. Mass-to-Charge (m/z)

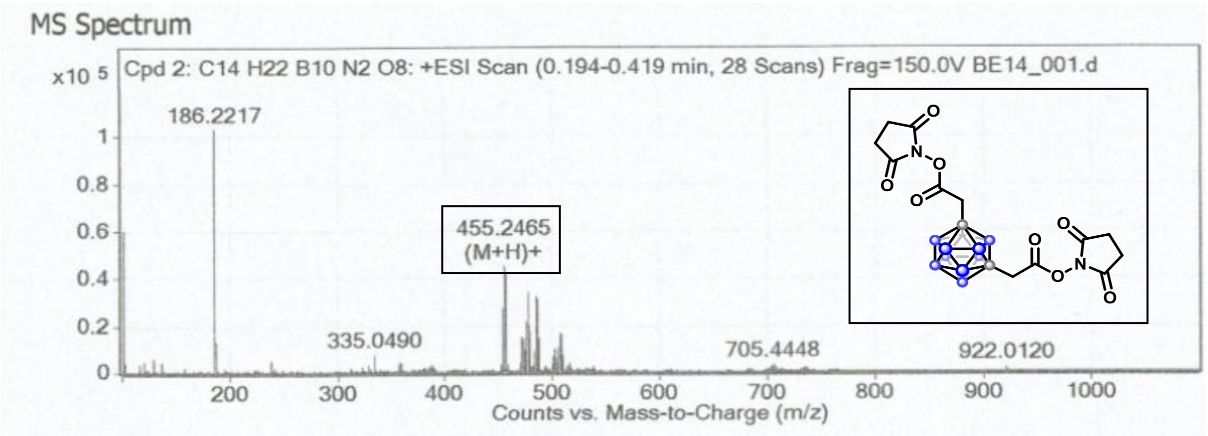
MS Spectrum
x10 5
Cpd 2: C14 H22 B10 N2 O8: +ESI Scan (0.194-0.419 min, 28 Scans) Frag=150.0V BE14_001.d
186.2217
455.2465
(M+H)+
335.0490
705.4448
922.0120
Counts vs. Mass-to-Charge (m/z)

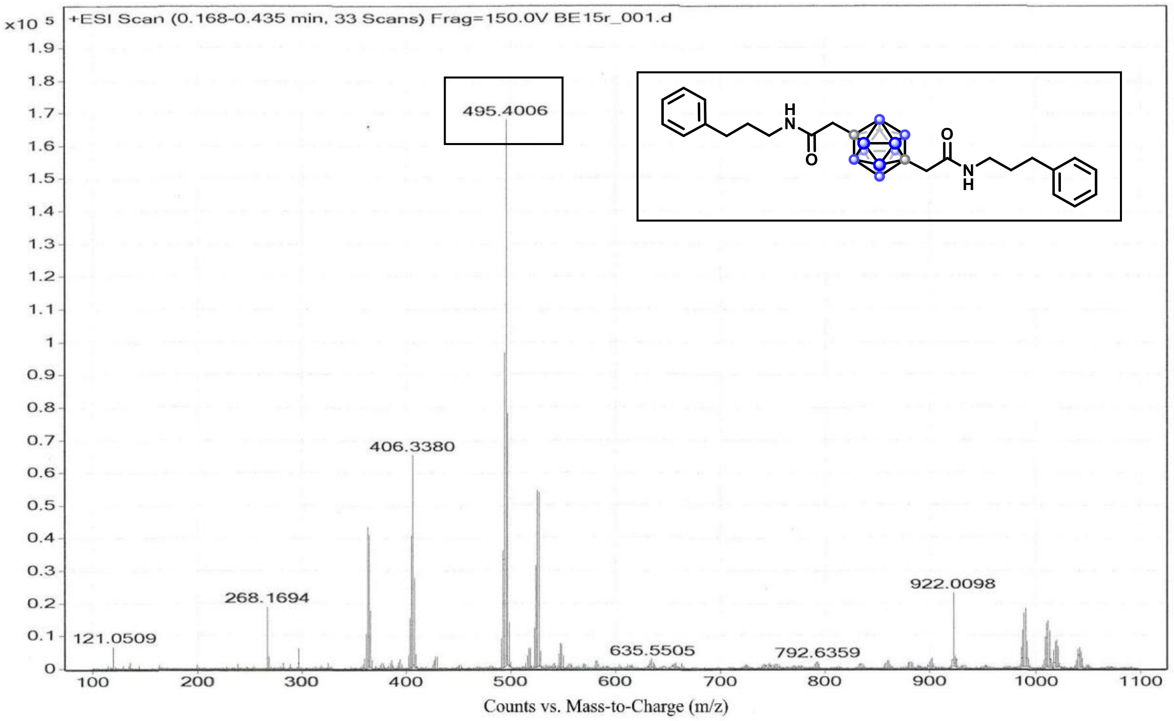

x10 5
+ESI Scan (0.168-0.435 min, 33 Scans) Frag=150.0V BE15r_001.d
495.4006
406.3380
268.1694
121.0509
922.0098
635.5505
792.6359
Counts vs. Mass-to-Charge (m/z)

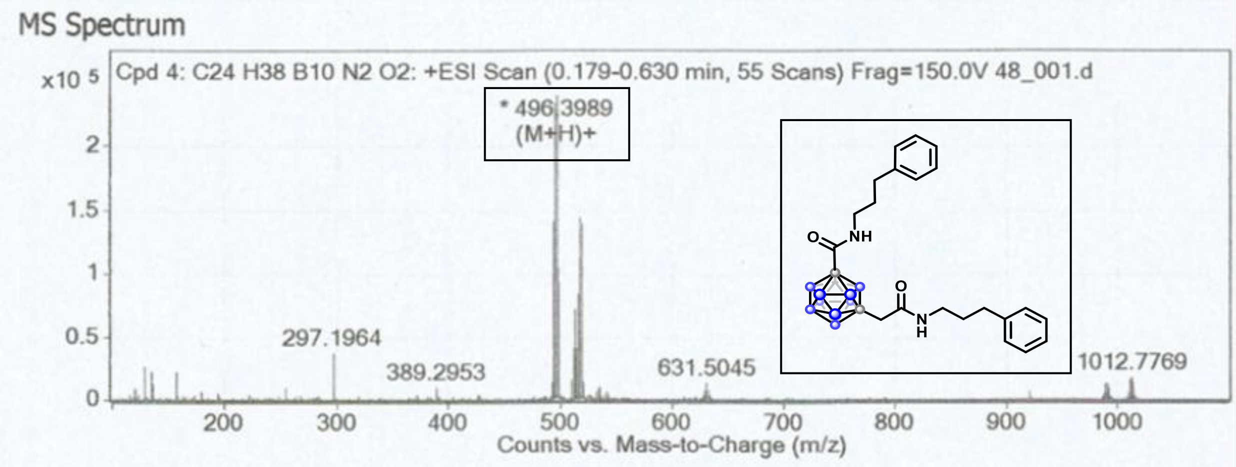

MS Spectrum
x10 5
Cpd 4: C24 H38 B10 N2 O2: +ESI Scan (0.179-0.630 min, 55 Scans) Frag=150.0V 48_001.d
* 496.3989
(M+H)+
297.1964
389.2953
631.5045
1012.7769
Counts vs. Mass-to-Charge (m/z)

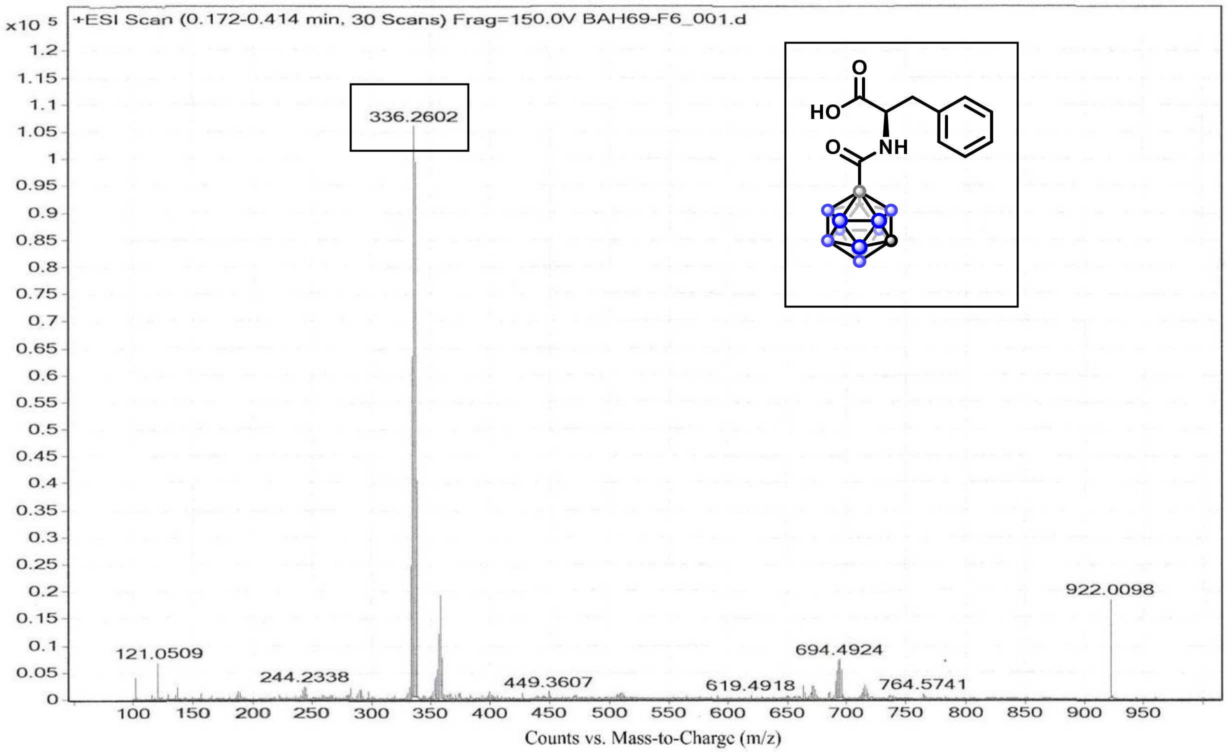

+ESI Scan (0.172-0.414 min, 30 Scans) Frag=150.0V BAH69-F6_001.d
x10 5
336.2602
922.0098
121.0509
244.2338
449.3607
619.4918
694.4924
764.5741
HO
O
NH
Counts vs. Mass-to-Charge (m/z)

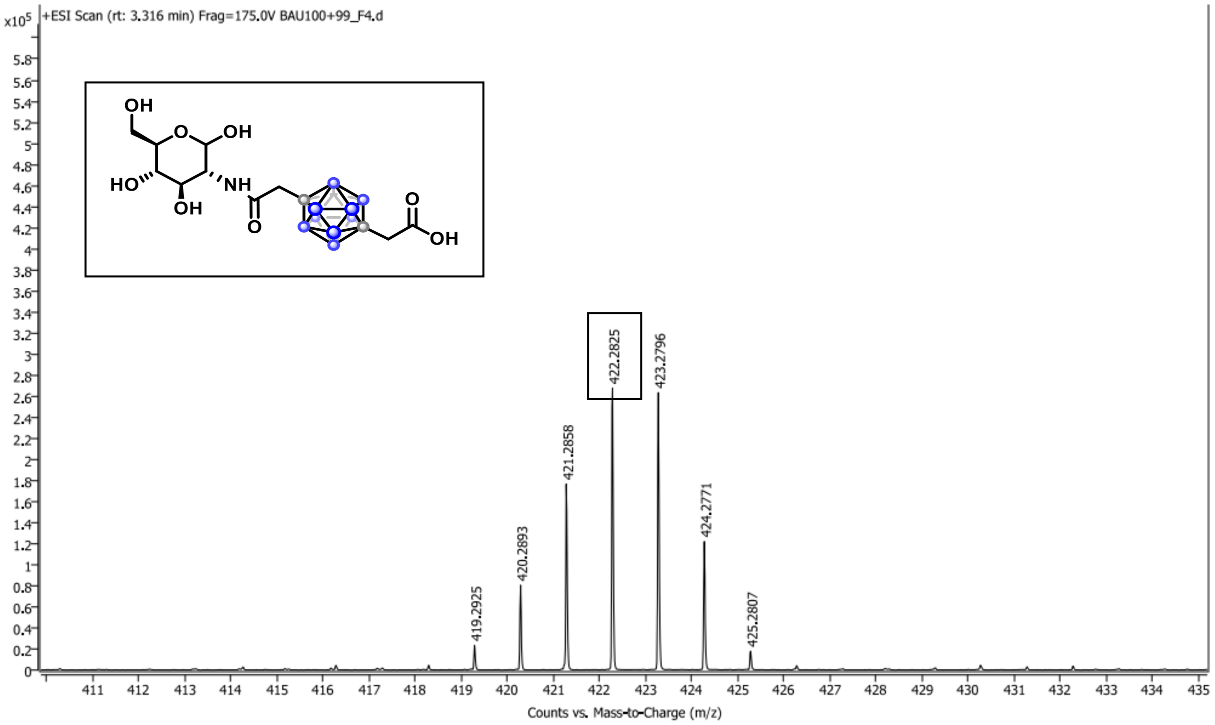

+ESI Scan (rt: 3.316 min) Frag=175.0V BAU100+99_F4.d
x10^5
OH
OH
HO
NH
O
OH
419.2925
420.2893
421.2858
422.2825
423.2796
424.2771
425.2807
Counts vs. Mass-to-Charge (m/z)

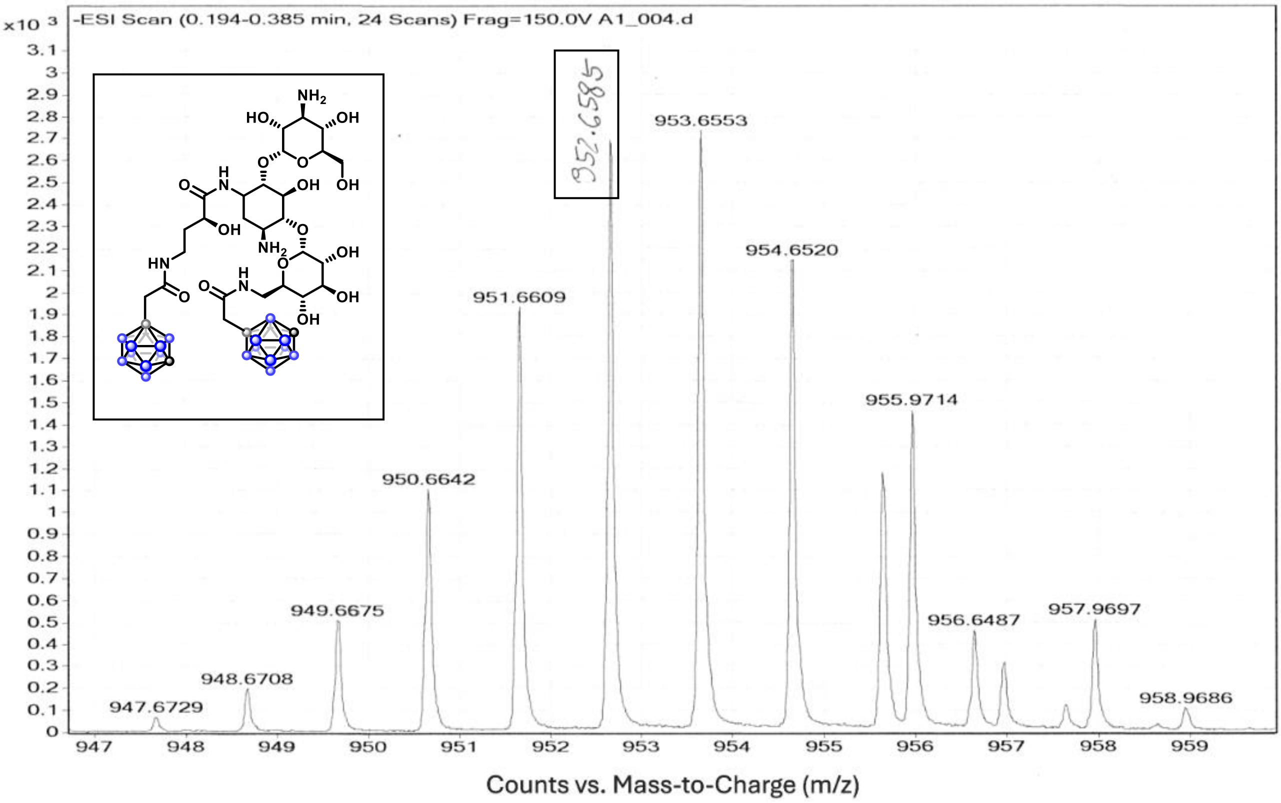
-ESI Scan (0.194-0.385 min, 24 Scans) Frag=150.0V A1_004.d
x10 3
952.6585
953.6553
954.6520
951.6609
955.9714
950.6642
949.6675
956.6487
957.9697
948.6708
947.6729
958.9686
Counts vs. Mass-to-Charge (m/z)

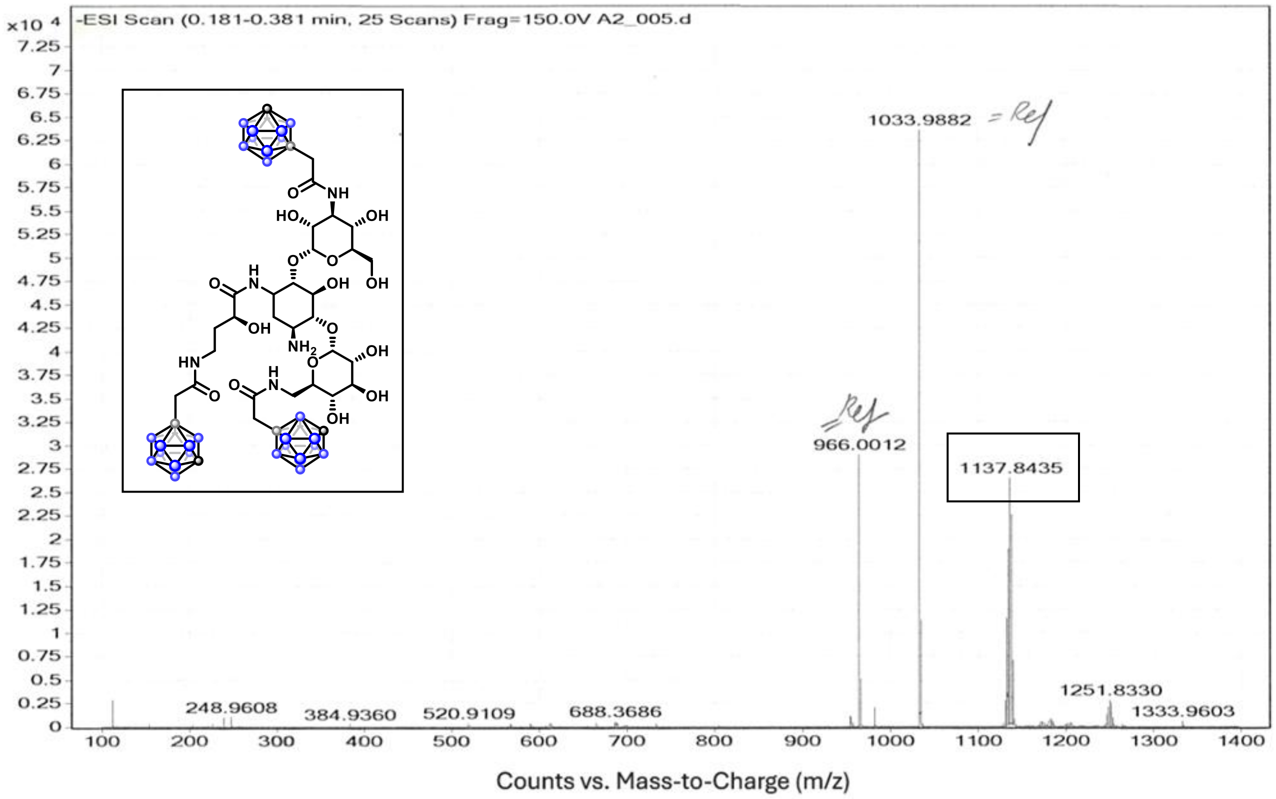
-ESI Scan (0.181-0.381 min, 25 Scans) Frag=150.0V A2_005.d
x10 4
1033.9882 = Ref
= Ref
966.0012
1137.8435
1251.8330
1333.9603
248.9608
384.9360
520.9109
688.3686
Counts vs. Mass-to-Charge (m/z)

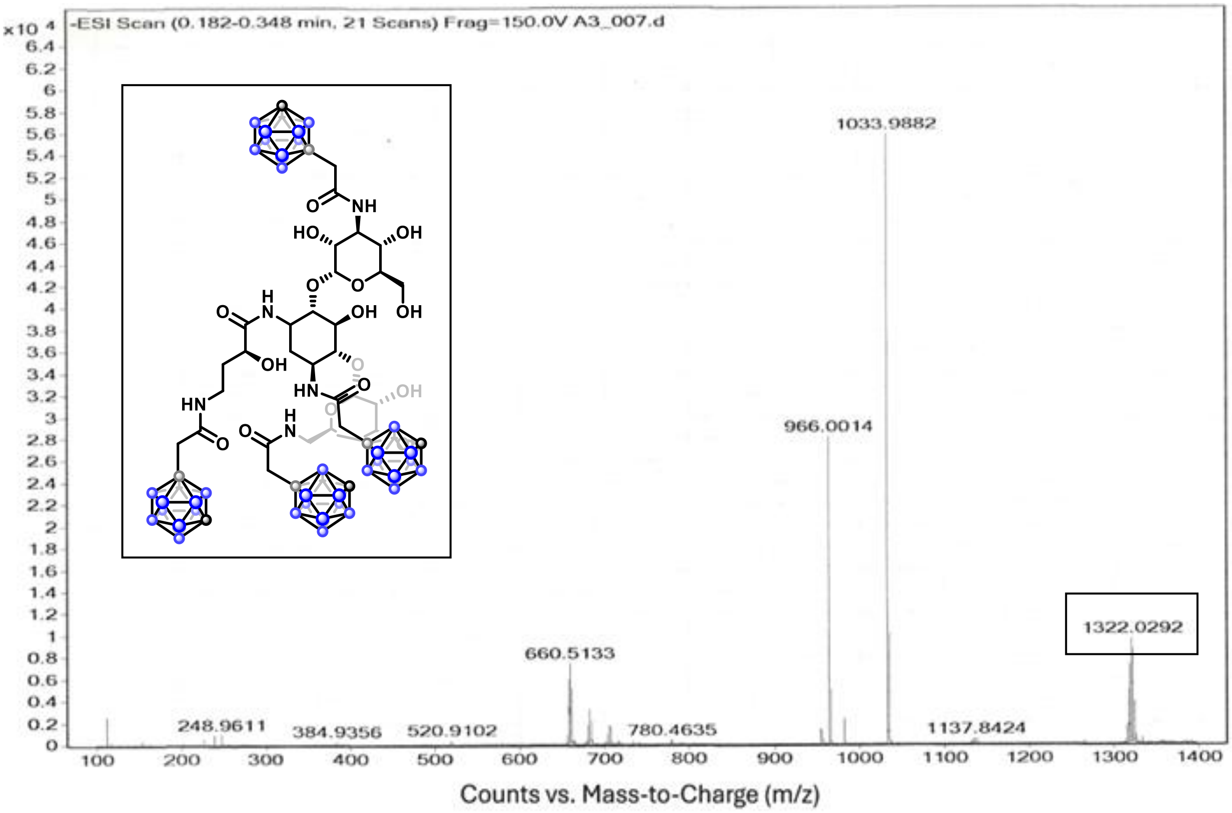
-ESI Scan (0.182-0.348 min, 21 Scans) Frag=150.0V A3_007.d
x10 4
1033.9882
966.0014
660.5133
1322.0292
248.9611
384.9356
520.9102
780.4635
1137.8424
Counts vs. Mass-to-Charge (m/z)

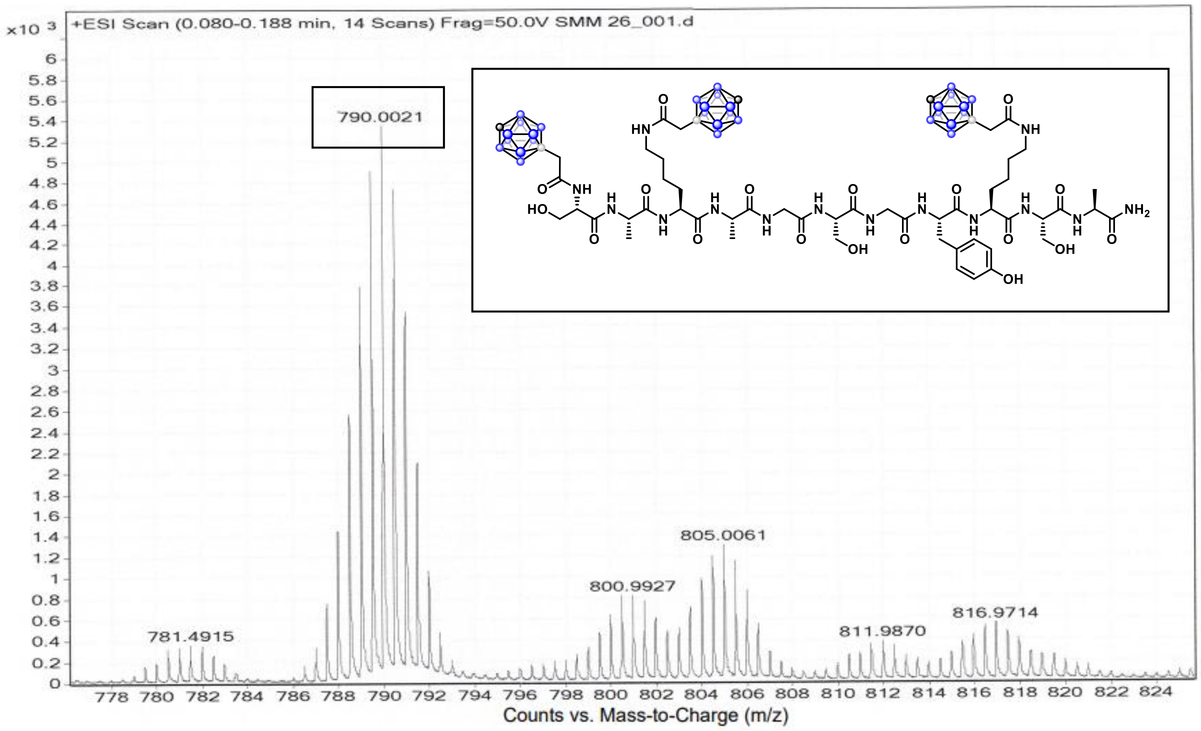
+ESI Scan (0.080-0.188 min, 14 Scans) Frag=50.0V SMM 26_001.d
x10 3
790.0021
805.0061
800.9927
781.4915
811.9870
816.9714
Counts vs. Mass-to-Charge (m/z)

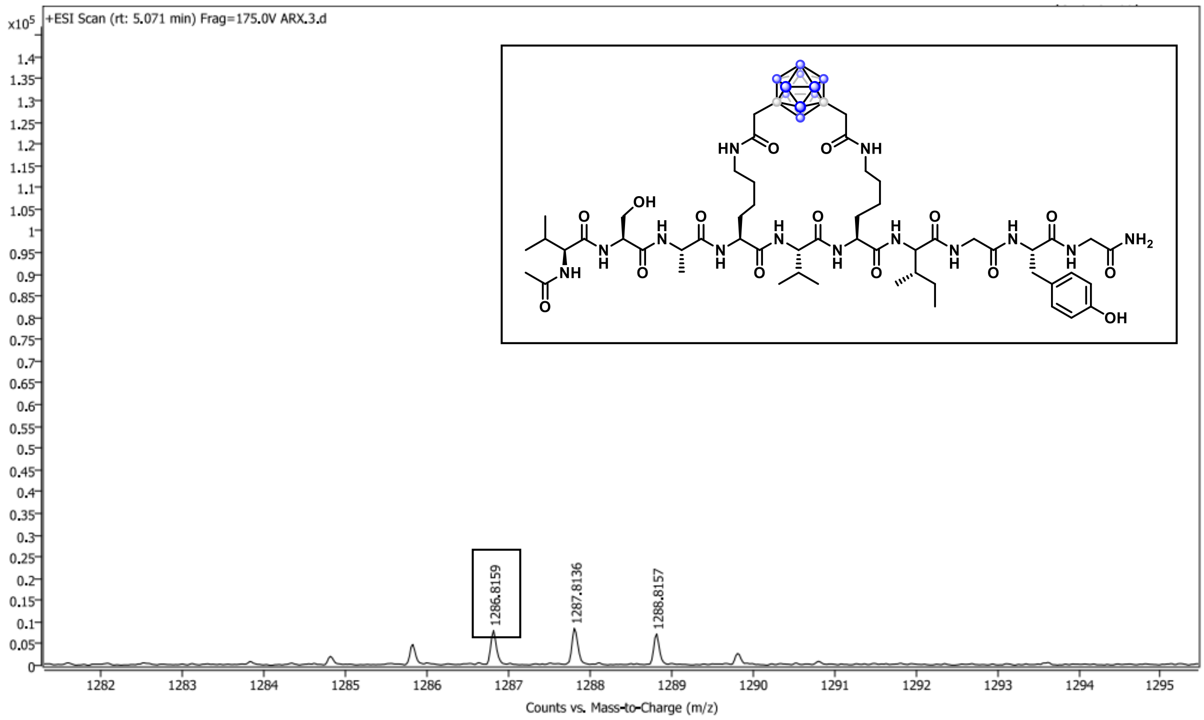
+ESI Scan (rt: 5.071 min) Frag=175.0V ARX.3.d
x10^5
1286.8159
1287.8136
1288.8157
Counts vs. Mass-to-Charge (m/z)

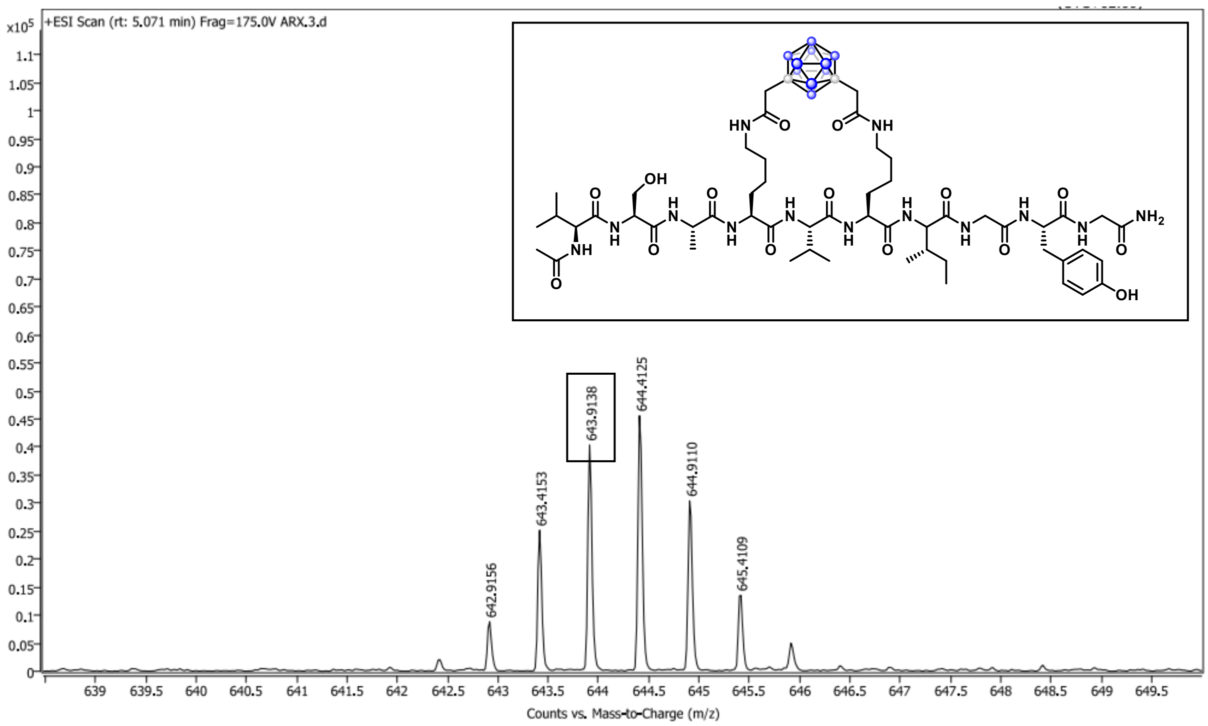
+ESI Scan (rt: 5.071 min) Frag=175.0V ARX.3.d
x10^5
642.9156
643.4153
643.9138
644.4125
644.9110
645.4109
Counts vs. Mass-to-Charge (m/z)

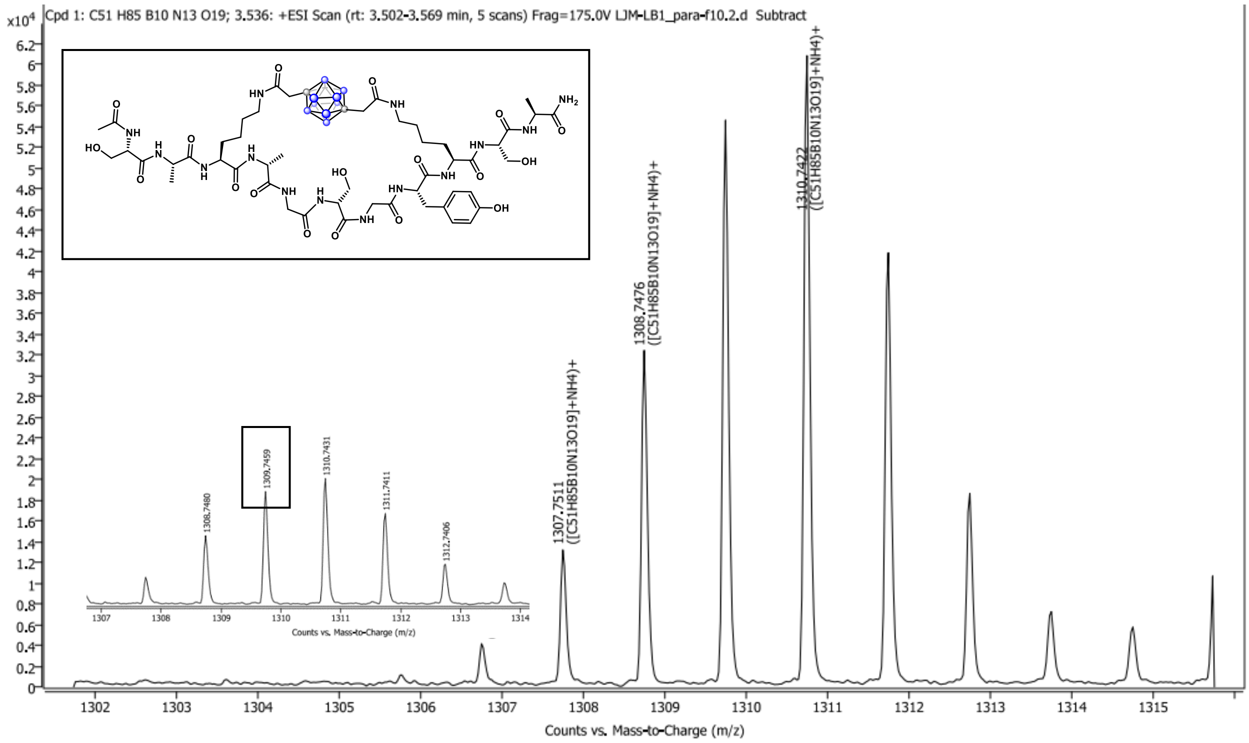
Cpd 1: C51 H85 B10 N13 O19; 3.536: +ESI Scan (rt: 3.502-3.569 min, 5 scans) Frag=175.0V LJM-LB1_para-f10.2.d Subtract
1307.7511 ([C51H85B10N13O19]+NH4)+
1308.7476 ([C51H85B10N13O19]+NH4)+
1310.7422 ([C51H85B10N13O19]+NH4)+
1308.7480
1309.7459
1310.7431
1311.7411
1312.7406
Counts vs. Mass-to-Charge (m/z)

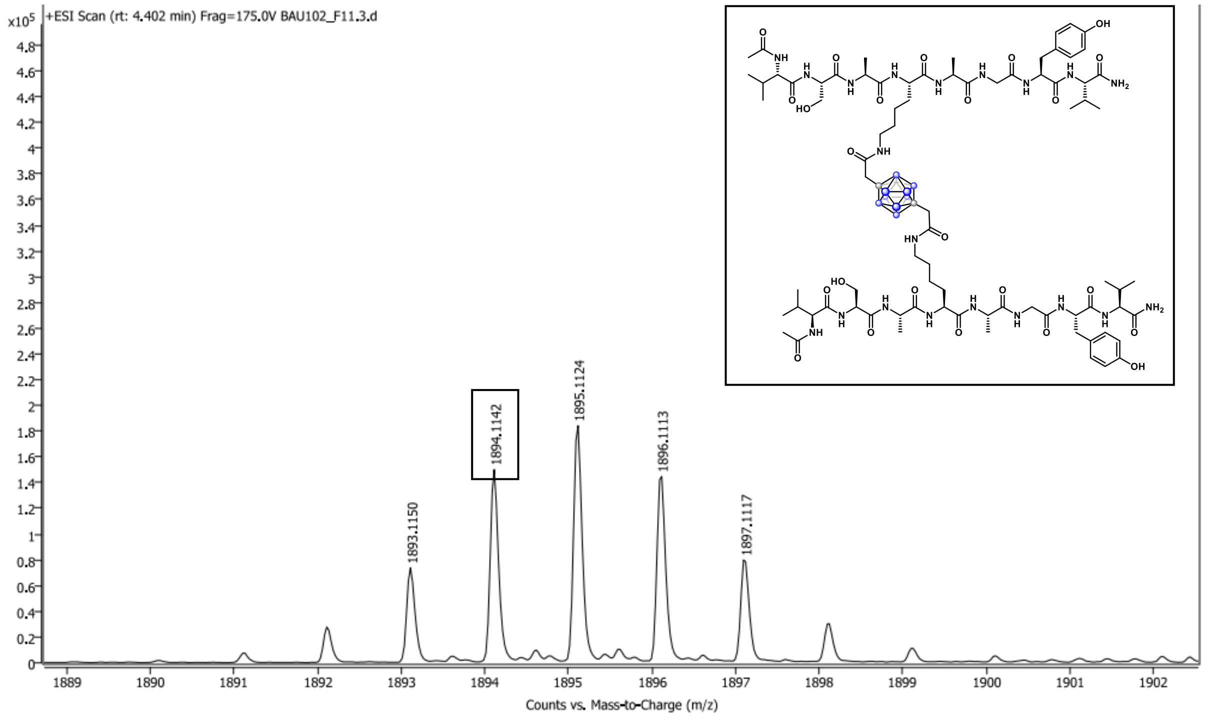
+ESI Scan (rt: 4.402 min) Frag=175.0V BAU102_F11.3.d
1893.1150
1894.1142
1895.1124
1896.1113
1897.1117
Counts vs. Mass-to-Charge (m/z)

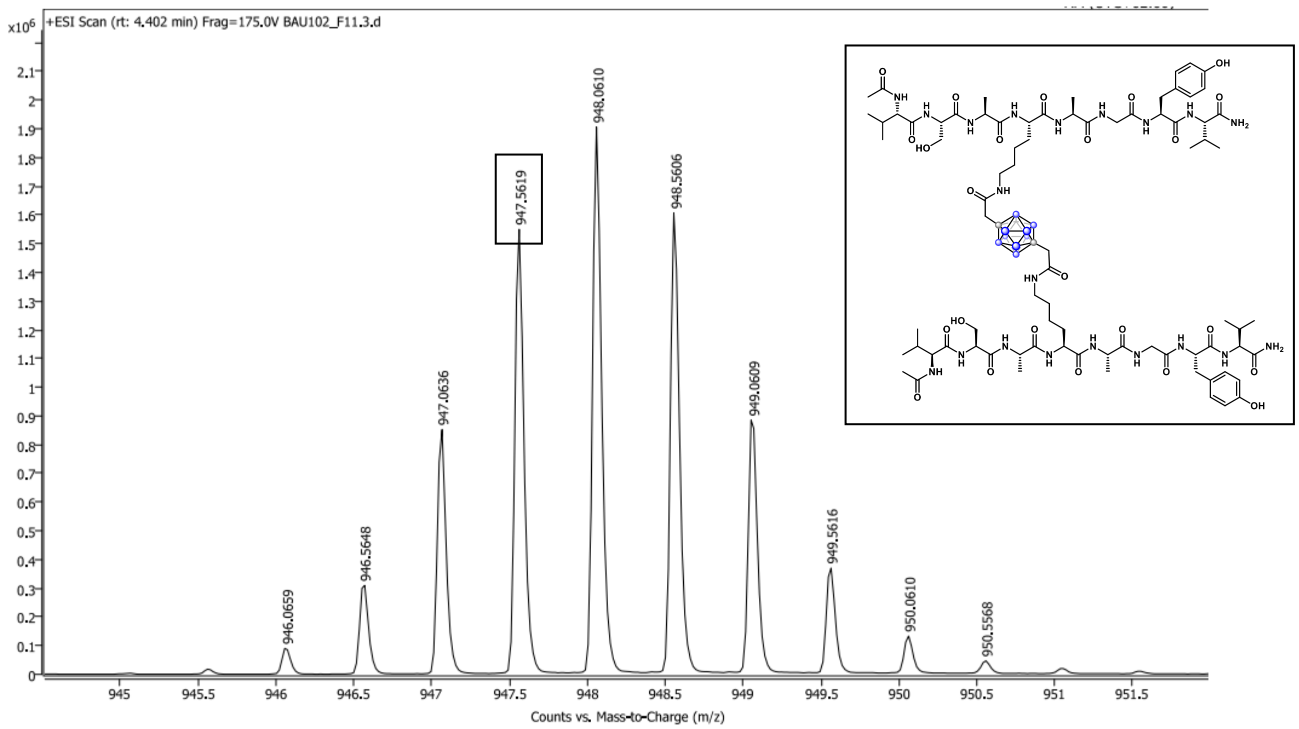

x10^6
+ESI Scan (rt: 4.402 min) Frag=175.0V BAU102_F11.3.d
946.0659
946.5648
947.0636
947.5619
948.0610
948.5606
949.0609
949.5616
950.0610
950.5568
Counts vs. Mass-to-Charge (m/z)

#### Literature references

[1] Choi, S.; Byun, Y. Synthesis of sterically-hindered 1.7-dicarba-closo-dodecarborane thiourea analogs. *J. Organomet. Chem.* **2013**, *733,* 49–52.

[2] Kasar, R. A.; Knudsen, G. M.; Kahl, S. B. Synthesis of 3-Amino-1-carboxy-o-carborane and an Improved, General Method for the Synthesis of All Three C-Amino-C-carboxycarboranes. *Inorg. Chem.* **1999**, *38,* 2936–2940.

[3] El-Zaria, M. E.; Genady, A. R.; Janzen, N.; Petlura, C. I.; Beckford Vera, D. R.; Valliant, J. F. Preparation and evaluation of carborane-derived inhibitors of prostate specific membrane antigen (PSMA). *Dalton Trans.* **2014**, *43,* 4950–4961.

[4] van den Berg, T. A.; Feringa, B. L.; Roelfes, G. Double strand DNA cleavage with a binuclear iron complex. *Chem. Commun.* **2007**, *2*, 180-182.

[5] Nekvinda, J.; Grüner, B.; Gabel, D.; Nau, W. M.; Assaf, K. I. Host-Guest Chemistry of Carboranes: Synthesis of Carboxylate Derivatives and Their Binding to Cyclodextrins. *Chem. Eur. J.* **2018**, *24*, 12970-12975.

[6] Scholz, M.; Bensdorf, K.; Gust, R.; Hey-Hawkins, E. Asborin: The Carbaborane Analogue of Aspirin. *ChemMedChem.*, **2009**, *4,* 746–748.

[7] Murray, K. K.; Boyd, R. K.; Eberlin, M. N.; Langley, G. J.; Li, L.; Naito, Y. Definitions of terms relating to mass spectrometry (IUPAC Recommendations 2013). *Pure Appl. Chem.*, **2013**, *85*, 1515–1609.

[8] Andrade, S. F.; Oliveira, B. G.; Pereira, L. C.; Ramos, J. P.; Joaquim, A. R.; Steppe, M.; Souza-Fagundes, E. M.; Alves, R. J. Design, synthesis and structure-activity relationship studies of a novel focused library of 2,3,4-substituted oxazolidines with antiproliferative activity against cancer cell lines. *Eur. J. Med. Chem.* **2017**, *138,* 13–25.

[9] Dhanjee, H. H.; Buslov, I.; Windsor, I. W.; Raines, R. T.; Pentelute, B. L.; Buchwald, S. L. Palladium–Protein Oxidative Addition Complexes by Amine-Selective Acylation. *J. Am. Chem. Soc.* **2020**, *142,* 21237–21242.

[10] Zhang, Z.; Shah, B. Characterization of variable region glycosylation in monoclonal antibodies using LC–MS. *Anal. Chem.* **2010**, 82, 10102–10110.

[11] Walsh, G.; Jefferis, R. Post-translational modifications in the context of therapeutic proteins. *Nat. Biotechnol.* **2006**, *24*, 1241–1252.

[12] Gazvoda, M.; Dhanjee, H. H.; Rodriguez, J.; Brown, J. S.; Farquhar, C.; Truex, N. L.; Loas, A.; Buchwald, S. L.; Pentelute, B. L. Palladium-Mediated Incorporation of Carboranes into Small Molecules, Peptides, and Proteins. *J. Am. Chem. Soc.* **2022**, *144*, 7852–7860.

[13] Burke, H. M.; Nicholls, B. T.; Stasiuk, G. J. NHS ester chemistry and its pitfalls: Reactions in bioconjugation and labeling. *Chem. Commun.* **2017**, *53*, 13295–13298.

[14] Gahoual, R.; Burr, A.; Busnel, J.-M.; Kühn, L.; Hammann, P.; Beck, A.; François, Y.-N.; Leize-Wagner, E. Rapid and Multi-Level Characterization of Trastuzumab Using Sheathless Capillary Electrophoresis–Tandem Mass Spectrometry. *mAbs* **2013**, *5*, 479–490.

[15] Hermanto, S.; Yusuf, M.; Mutalib, A.; Hudiyono, S. Molecular Dynamic Simulation of Trastuzumab F(ab′)$_2$ Structure in Corporation with HER2 as a Theranostic Agent of Breast Cancer. *J. Phys.: Conf. Ser.* **2017**, *835*, 012005.
